# Population sequencing enhances understanding of tea plant evolution

**DOI:** 10.1101/2020.03.19.998393

**Authors:** Xinchao Wang, Hu Feng, Yuxiao Chang, Chunlei Ma, Liyuan Wang, Xinyuan Hao, A’lun Li, Hao Cheng, Lu Wang, Peng Cui, Jiqiang Jin, Xiaobo Wang, Kang Wei, Cheng Ai, Sheng Zhao, Zhichao Wu, Youyong Li, Benying Liu, Guo-Dong Wang, Liang Chen, Jue Ruan, Yajun Yang

## Abstract

Tea is an economically important plant characterized by a large genome size and high heterozygosity and species diversity. In this study, we assembled a 3.26 Gb high-quality chromosome-scale genome for tea using the ‘Longjing 43’ cultivar of *Camellia sinensis* var. *sinensis*. Population resequencing of 139 tea accessions from around the world was used to investigate the evolution of tea and to reveal the phylogenetic relationships among tea accessions. With the spread of tea cultivation, hybridization has increased the heterozygosity and wide-ranging gene flow among tea populations. Population genetics and transcriptomics analyses revealed that during domestication, the selection for disease resistance and flavor in *C. sinensis* var. *sinensis* populations has been stronger than that in *C. sinensis* var. *assamica* populations. The data compiled in this study provide new resources for the marker assisted breeding of tea and are a basis for further research on the genetics and evolution of tea.

## Introduction

Tea [*Camellia sinensis* (L.) O. Kuntze, 2n = 30] is one of the most important and traditional economic crops in many developing countries in Asia, Africa, and Latin America, and it is consumed as a beverage by more than two-thirds of the world’s population^1,2^. Originally, tea was used as a medicinal herb in ancient China, and it was not until the Tang dynasty (A.D. 618-907) that it gained popularity as a beverage^3,4^. From that time on, tea planting expanded throughout the world through the influence of trading along the Silk and Tea Horse Roads^5,6^. Subsequent to its initial domestication, the further breeding and cultivation of tea contributed to enhancement of certain organoleptic traits, primarily taste and aroma, and biotic and abiotic stress resistance properties, including cold and disease resistances^7^. However, the genes underlying the traits that were gradually selected and expanded remain to be determined.

The majority of cultivated tea plants belong to the genus *Camellia* L., section *Thea* (L.) Dyer, in the family Theaceae, and are categorized into one of two main varieties: *C. sinensis* var. *sinensis* (CSS) and *C*. *sinensis* var. *assamica* (Masters) Chang (CSA). CSS is characterized by smaller leaves, cold tolerance, and a shrub or semi-shrub growth habit, whereas CSA has larger leaves and an arbor or semi-arbor habit^8,9^. Moreover, some *C. sinensis*-related species (CSR) belonging to the section *Thea*, such as *C*. *taliensis* (W.W. Smith) Melchior, *C. crassicolumna* Chang, *C. gymnogyna* Chang, and *C. tachangensis* F.C. Zhang, are locally consumed as tea by inhabitants in certain regions of the Indo-China Peninsula, particularly in Yunnan Province, China. Theoretically, different species are assumed to have experienced reproductive isolation; however, different tea species can readily hybridize, and thus it is difficult to accurately classify the offspring of different hybrids. Moreover, numerous morphological features are continuous, which makes it difficult to identify taxonomic groups^10^. The traditional classifications of tea have been based on morphology and sometimes contradict the more recent classifications based on molecular characterization^11–15^; however, given that tea plant taxonomy generally lacks comprehensive genomic evidence, further analyses using population resequencing are required to optimize taxonomic assignments at the whole-genome level.

With a view toward gaining a better understanding of the domestication, breeding, and classification of tea, we collected and sequenced samples of 139 tea accessions from across the world. High-quality annotated genes and chromosome-scale tea genomes were necessary for our population research. In this regard, previous elucidations of the genomes of the tea cultivars Yunkang 10 (YK10, CSA)^1^ and Shuchazao (SCZ, CSS)^16^ are considered important milestones in tea genetic research. However, these two genomes were not characterized at the chromosome scale, and scaffold N50 values were less than 1.4 Mb, thereby impeding evaluation of the phenotypic variation and genome evolution in important intergenic regions. Moreover, the core genes (Benchmarking Universal Single-Copy Orthologs^17^, BUSCO) of the SCZ and YK10 genomes were respectively only 80.58% and 68.58% complete, and accordingly this incomplete gene annotation has hampered further population selection, functional genomics analysis, and molecule breeding research. Therefore, for the purposes of *de novo* genome assembly in the present study, we focused on the ‘Longjing 43’ (LJ43) cultivar of *C. sinensis*, which is among the most widely cultivated tea cultivars in China, and it is characterized by high cold resistance, extensive plantation adaptation, early sprouting time, excellent taste and favorable aroma, etc^18^.

Herein, we describe a high-quality chromosome-scale tea genome, along with divergent selection directions in the CSS and CSA populations, and present a phylogenetic tree of tea. However, details regarding the origin of tea and the subsequent routes of expansion remain to be clarified, thus presenting opportunities for further research.

## Results

### Sequencing and assembly of the LJ43 genome

The predicted size of the LJ43 genome was approximately 3.32 Gb (Supplementary Figs. 1 and 2), which is larger than the assembled YK10 (2.90–3.10 Gb)^1^ and SCZ (~2.98 Gb)^16^ genomes. To enhance genome assembly, 196 Gb SMRT long reads (Supplementary Table 1) were initially assembled using WTDBG^19^ (Version 1.2.8; Supplementary Material), which resulted in a 3.26-Gb assembled genome containing 37,600 contigs and covering approximately 98.19% of the whole genome. To further improve the integrity of the assembled genome, contigs were scaffolded based on chromosome conformation capture sequencing (Hi-C) (Supplementary Table 1, Supplementary Figs. 3 and 4, Supplementary Material) and the final assembly of 3.26 Gb was generated with a scaffold N50 value of 144 Mb. Of the 37,600 initially assembled contigs, 7,071 (~2.31 Gb, 70.9% of the original assembly) were then anchored with orientation into 15 chromosomal linkage groups (Fig. 1b, Supplementary Fig 5, Supplementary Tables 2 and 3).

**Fig. 1.**
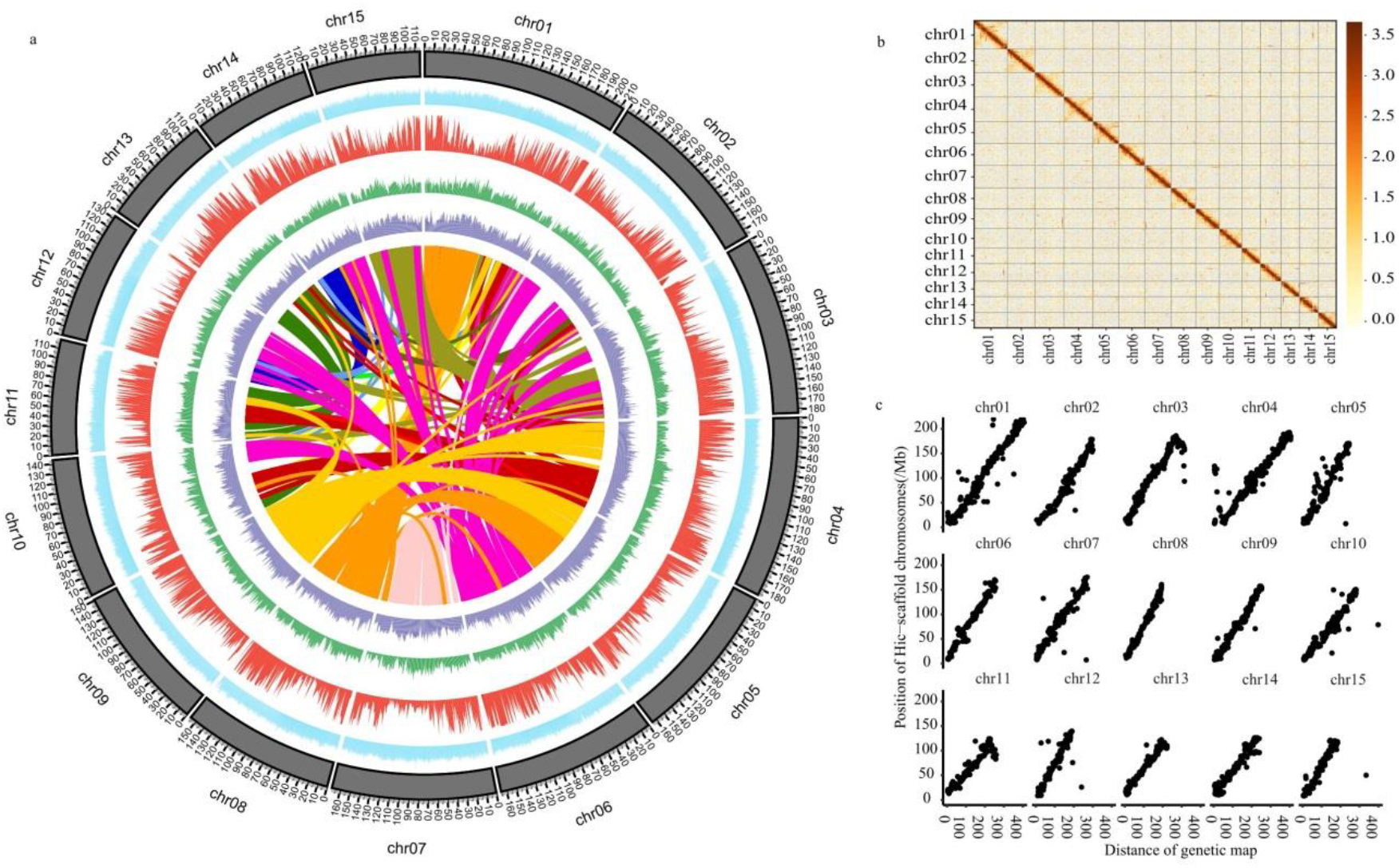
Characterization and quality of the LJ43 genome. a, The landscape of the LJ43 genome. From inside to outside: LJ43 gene collinearity; long terminal repeat density (purple); single-nucleotide polymorphism density (green); gene density (red); GC content (blue). The chromosome units of all the above-mentioned features are 1 Mbp. b, Genome-wide all-by-all Hi-C interaction. The resolution is 0.5 Mbp. c, The collinearity of the genetic map and assembled genome.

To evaluate the quality of the assembled LJ43 genome, we estimated the sequence accuracy at both the single-base and scaffold levels. The percentages of homogeneous single-nucleotide polymorphisms (SNPs) and homogeneous insertions-deletions (InDels) in the genome were 0.000224% and 0.000568%, respectively, thereby indicating a low error rate at the single-base level (Supplementary Table 4). The accuracy of the scaffolding was evaluated based on three strategies. Firstly, 5,879 (83.14%) of 7,071 connections in the Hi-C scaffolds were confirmed with at least two 10x Genomics Chromium linked reads spanning the connections. Secondly, 5,374 (76.00%) connections were confirmed by at least two BioNano Genomics (BNG) optical molecules, among which, 4,484 (63.41%) overlapped with those confirmed by the 10x Genomics Chromium linked reads. In total, 6,769 (95.73%) connections in the scaffold generated with Hi-C could be confirmed by 10x Genomics Chromium linked reads or BNG optical molecules, indicating that the scaffold was accurate. Thirdly, the collinearity of the tea genetic map^20^ with 3,483 single sequence repeat (SSR) markers and the LJ43 genome had a mean coefficient of determination (*R*^2^) of 0.93, with a maximum value of 0.98 and a minimum value of 0.84 (Fig. 1c, Supplementary Table 3). In summary, the assembly accuracy for the LJ43 genome at both the single-base and scaffold levels was high.

### Genome annotation

For genome annotation, we annotated the repetitive sequences of the genome combined with the strategies of *de novo* and homology-based prediction. We identified and masked 2.38 Gb (80.06%) of the LJ43 genome as repetitive sequences (Supplementary Table 5). Among the integrated results, 60.77% (1.98 Gb) were long terminal repeat (LTR) retrotransposons (Supplementary Table 6), with LTR/Gypsy elements being the dominant class (49.85% of the whole genome, 1.63 Gb), followed by LTR/Copia elements (7.09%, 231.27 Mb). Compared with the previously sequenced tea genomes, the LTR/Gypsy and LTR/Copia repeats were similar in SCZ (Gypsy 46%, Copia 8%)^16^ and YK10 (Gypsy 47%, Copia 8%)^1^, whereas the LTR/Gypsy and LTR/Copia repeats in tea are expanded compared with those in kiwifruit (*Actinidia chinensis*) (13.4%)^21^, silver birch (*Betula pendula*) (10.8%)^22^, and durian (*Durio zibethinus*) (29.4%)^23^, but contracted compared with those of maize (*Zea mays*) (74.20%)^24^.

LTR retrotransposons are the predominant repeat elements that tend to be poorly assembled in draft genomes^25^, and it has been reported that the LTR assembly index (LAI), which approximates to the ratio of intact LTR to total LTR, can be exploited to evaluate assembly continuity. Thus, we investigated the LTR composition of the LJ43 genome and compared this with that of the SCZ and YK10 genomes, and found that the LAI of the LJ43, SCZ, and YK10 genomes was 5.50, 3.29, and 0.98, respectively, thereby indicating that a larger number of intact LTR retrotransposons had been assembled in the LJ43 genome. We used LTR-finder to detect intact LTR retrotransposons in the three tea genomes, and then aligned the 5ʹ and 3ʹ terminal repeats using MUSCLE (version 3.8.31), and calculated the Kimura two-parameter distance for each alignment using EMBOSS (version 6.4.0). The equation Time = Ks/2μ (μ = 6.5E-9)^26^ was used to calculate the insertion time of each LTR. Unexpectedly, we found that the LTR from LJ43 accumulated less point mutations, resulting in the calculated peaks of LTR insertion in LJ43, SCZ, and YK10 at 1 million years ago (mya), 9 mya, and 9 mya, respectively (Supplementary Fig. 6). To further investigate this seemingly anomalous pattern, we performed NGS read error correction during genome assembly. We compared the genome sequences corrected by PacBio reads and NGS reads and found that 98.19% of the 5ʹ and 3ʹ terminal IR sequences were corrected no more than three bases by NGS reads (Supplementary Fig. 6d). Moreover, error correction could not change the Ks from 0.013 (the peak of LJ43) to 0.117 (the peak of SCZ and YK10). Taken together, our analyses indicate that the LJ43 genome assembly was more complete than that of the previously sequenced tea genomes, has a high LAI, and contains more recently derived LTRs, which resulted in contradictory estimates of the LTR insertion time among LJ43, SCZ, and YK10.

To assist in gene prediction, we generated a total of 340 Gb of clean RNA-seq data from 19 samples of five tissue types (bud, leaf, flower, stem, and root) collected in each of the four seasons (with the exception of flowers during summer, Supplementary Table 7). Protein-coding genes were annotated using integrative gene prediction with *ab initio* prediction, homology search, and transcriptome data. EVidenceModeler (version 1.1.1) was used to integrate all predicted gene structures. A total of 33,556 protein-coding genes with an RNA-seq coverage ratio greater than 50% were annotated with an average gene size of 10,816 bp (Supplementary Material, Supplementary Fig. 7) and a mean number of 5.3 exons per gene (Table 1). Subsequently, we assessed LJ43 genome annotation integrity using the BUSCO database^17^, and found that 1,215 (88.36%) annotations were complete, compared with the 1,108 (80.58%) and 943 (68.58%) complete annotations obtained for the SCZ and YK10 genomes, respectively.

**Table 1.** Genome assembly and annotated genes of the tea cultivars LJ43, SCZ, and YK10

|  | LJ43 | SCZ | YK10 |
| --- | --- | --- | --- |
| Genome size | 3.26 G | 3.14 G | 3.02 G |
| Contigs N50 | 271.33 kb | 67.01 kb | 19.96 kb |
| Scaffold N50 | 143.85 Mb | 1.39 Mb | 0.45 Mb |
| GC percentage | 38.67% | 37.84% | 39.62% |
| Number of genes | 33,556 | 33,932 | 36,951 |
| Number of exons | 188,681 | 191,870 | 176,616 |
| Length of exons | 40.4 Mb | 45.6 Mb | 41.6 Mb |
| Average length of exons | 226.1 bp | 237.8 bp | 235.6 bp |
| Average length of genes<br>(intron+exon) | 10,815.5 bp | 7,385 bp | 3,548 bp |
| Average number of exons per<br>gene | 5.3 | 5.7 | 4.8 |
| Average length of coding<br>sequence | 1,205 bp | 1,345 bp | 1,131 bp |
| BUSCO | 88.36% | 80.58% | 68.58% |

Using the genome annotation data, we determined the chromosomal locations of 26,561 (79.15%) annotated genes. Furthermore, we compared the protein sequences of the LJ43 genome with those of *Actinidia chinensis*, which has a high-quality reference genome sequence and belongs to the order of Ericales, and used MCScanX to detect synteny (Supplementary Fig. 8). The results revealed that the LJ43 genome comprises 690 collinear blocks containing 18,030 genes, whereas the SCZ genome has 111 collinear blocks containing 1,487 genes, and that of YK10 has 54 collinear blocks containing 393 genes. Furthermore, we found that the extent of genome synteny between LJ43 and cocoa (*Theobroma cacao*) is comparable to that with *Actinidia chinensis* (Supplementary Material).

### Gene family evolution

To gain an insight into the evolution of the tea genome, we grouped orthologous genes using OrthoMCL (Supplementary Material), and accordingly obtained 24,350 groups of orthologous gene families among nine genomes: LJ43, *Actinidia chinensis*, *Coffea canephora*, *Theobroma cacao*, *Arabidopsis thaliana*, *Oryza sativa* subsp. *japonica*, *Populus trichocarpa*, *Amborella trichopoda*, and *Vitis vinifera*. Among these, 1,034 single-copy gene families were used to construct a phylogeny tree for the tea genome using *Amborella trichopoda* as an outgroup (Supplementary Fig. 9). Gene family evolution was analyzed using CAFE, which revealed that a total of 1,936 tea gene families have undergone expansion and 1,510 tea gene families have undergone contraction. Gene Ontology (GO), InterPro (IPR), and Kyoto Encyclopaedia of Genes and Genomes (KEGG) enrichment analyses of the expanded genes indicated the expansion of gene families involved in disease resistance, secondary metabolism, and growth and development (P-value < 0.05, FDR < 0.05, Supplementary Tables 8-14). Among these families: UDP-glucuronosyl/UDP-glucosyltransferase (GO:0016758, P-value < 2.20E-16, FDR < 2.40E-14), which catalyzes glucosyl transfer in flavanone metabolism, is related to catechin content; (-)-germacrene D synthase (K15803, P-value = 8.01E-06, FDR = 0.91E-03) catalyzes the conversion of farneyl-PP to germacrene D and is related to terpene metabolism; NB-ARC (GO:0043531, P-value < 2.20E-16, FDR < 2.40E-14), Bet v I/Major latex protein (GO:0009607, P-value = 4.49E-04, FDR = 8.64E-03), RPM1 (K13457, P-value < 2.20E-16, FDR < 1.25E-13), and RPS2 (K13459, P-value = 8.88E-08, FDR = 2.51E-05) are related to disease resistance; and the S-locus glycoprotein domain (GO:0048544, P-value < 2.20E-16, FDR < 2.40E-14) is associated with self-incompatibility.

Furthermore, we used the “Branch-site” models A and Test2 to identify those genes in the tea genome that have evolved under positive Darwinian selection using codeml in the PAML (version 4.9d) package (Supplementary Material). A total of 1,031 single-copy genes from the above mentioned nine genomes were scanned to identify those genes under selection. After filtering (in Methods), we identified 74 genes that appeared to be under positive selection (FDR ≤ 0.05, Supplementary Table 15); some of these genes are involved in disease resistance, enhanced cold tolerance, and high light tolerance. In this regard, it has previously been reported that overexpression of cationic peroxidase 3 (OCP3)^27^ (Cha14.159) and Serpin-ZX^28^ (Cha9.301) is involved in the process of disease resistance, and that of beta-glucosidase-like SFR2 (SFR2, Cha5.171) is involved in freezing tolerance^29^. Other identified genes include one involved in the maintenance of photosystem II under high light conditions (MPH1^30^, ChaUn21494.1), and a photosystem II 22-kDa protein (PSBS, Cha9.807) that protects plants against photooxidative damage.

### Whole-genome duplication and divergence of tea genomes

In order to estimate the whole-genome duplication of the tea genome, we selected a total of 3,373, 3,199, and 2,992 gene families containing exactly two paralogous genes from the SCZ, LJ43, and YK10 genomes, respectively, to calculate the Ks values of the gene pairs. The results showed that the Ks peak of the three tea genomes was 0.3 (Supplementary Fig. 10), and the most recent duplication time was approximately 25 mya (Time = Ks/2μ, μ = 6.1E-9)^31^, thereby indicating that these cultivars underwent the same genome duplication event. Syntenic genes between LJ43 and SCZ and between LJ43 and YK10 were identified to calculate the Ks values of the pairs; the Ks peaks for the LJ43 and SCZ pairs were approximately 0.003 (~0.25 mya) and for the LJ43 and YK10 pairs were approximately 0.045 (~3.69 mya) (Supplementary Fig. 11), thus indicating that the divergence times of LJ43 and SCZ were more recent than those of LJ43 and YK10.

### Population genetic analysis

Tea leaves from different species or cultivars are often processed into different types of teas according to their processing suitability and local consumer preferences, e.g., CSA leaves are often processed to produce black tea, whereas CSS leaves are typically processed to produce green or oolong tea. To investigate the genetic basis of these differences, we examined the genomes of the 139 tea accessions collected from around the world, including 105 from East Asia, seven from South Asia, nine from Southeast Asia, six from West Asia, seven from Africa, and five from Hawaii (Fig. 2a, Supplementary Table 16, Supplementary Material). The specimens were sequenced at an average depth of 13.67-fold per genome (Supplementary Table 16), and given that the LJ43 genome is well annotation and a high level of continuity, we selected this as the reference genome. We accordingly achieved an average mapping rate of 99.07%, with a minimum rate of 96.95% and a maximum rate of 99.66% (Supplementary Table 16). After performing five filtering steps (described in the Methods section), we identified a total of 218.87 million SNPs among the tea populations, with a density of approximately 67 SNPs per kb (Fig. 1a, Supplementary Tables 17 and 18). We anticipate that this extensive whole-profile SNP dataset will serve as a valuable new resource for further tea genomics research and marker assisted breeding.

**Fig. 2.**
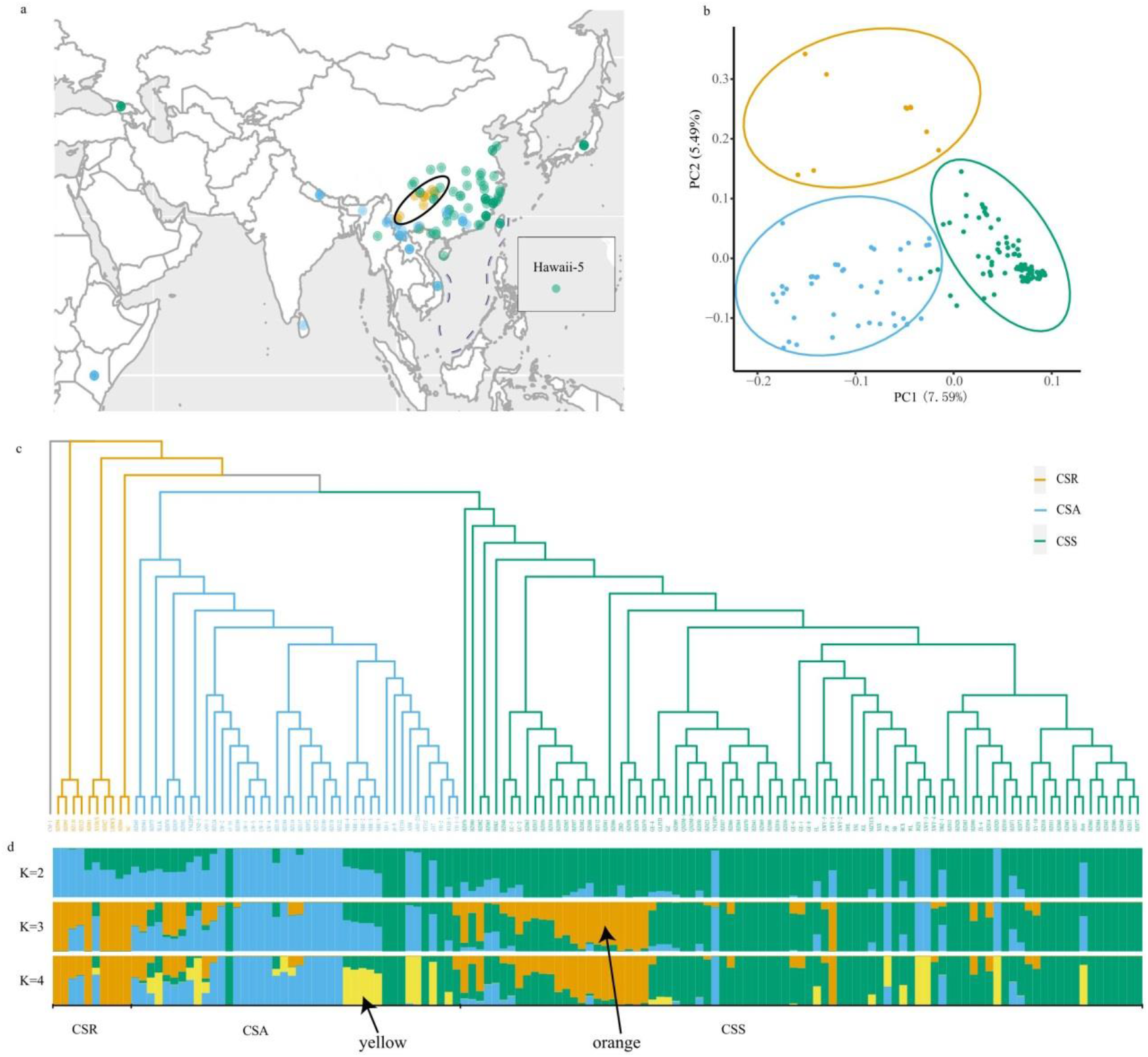
Distribution and evolution of tea. a, The distribution of tea accessions assessed in the present study. The teas within the black oval, had the highest nucleotide polymorphisms. b, Principal component analysis of the tea populations. PC1 and PC2 split the tea populations into three clusters. The *Camellia sinensis* var. *sinensis* (CSS) samples were found to cluster more tightly than the *C*. *sinensis* var. *assamica* (CSA) samples. c, A phylogenetic tree of tea. *Camellia sasanqua* Thunb. was used as the outgroup, and the tea samples closest to the outgroup were *C. sinensis*-related species (CSR). d, The structure of the tea populations. The green, blue, and yellow represent CSS, CSA, and CSR populations, respectively. The yellow and orange are marked with arrows.

To further investigate the phylogenetic relationships among these accessions, we constructed a maximum likelihood phylogenetic tree based on SNPs filtered from the total SNP dataset (Methods) using *Camellia sasanqua* as an outgroup (Fig. 2c). We found that all samples were clustered into one of three independent clades (Fig. 2c, Supplementary Fig. 12) corresponding to the CSR, CSS, and CSA populations; this result is consistent with the morphology-based classical taxonomy of CSA and CSS.

Principal component analysis (PCA) was used to investigate the relationships and differentiation among populations and consistently revealed the presence of three clusters corresponding to the CSA, CSS, and CSR teas (Fig. 2b). The first two principal components accounted for 13.08% of the total variance, with PC1 reflecting the variability of the CSA and CSS groups, and PC2 differentiating CSR plants from CSA and CSS. CSS had better aggregation than CSA and CSR, while the juncture accessions of CSA and CSS were also close to CSR in the phylogenetic tree. When K was 3, the CSA, CSS and CSR could be distinguished (Fig. 2d, Supplementary Fig. 13, Supplementary Material), this was consistent with the PCA result (Fig. 2b). When k ranged from 3 to 4, most of new accessions collected from China arose from CSA and CSS (yellow color, marked with arrow in Fig. 2d), indicating their high diversity.

On the basis of the phylogenetic and population structure results (Fig. 2c, Supplementary Figs. 12, 14, and 15), we further investigated the individual and population heterozygosities among the populations (Supplementary Table 16). We accordingly found the heterozygosity of CSR (6.37E-3) to be significantly higher than that of CSA (6.29E-3) and CSS (5.69E-3) (both P < 0.05, Supplementary Fig. 16). We also calculated linkage disequilibrium (LD) decay values based on the squared correlation coefficient (*r*^2^) of pairwise SNPs in two groups, which revealed that for the CSA and CSS groups, the average *r*^2^ among SNPs decayed to approximately 50% of its maximum value at approximately 41 kb and 59 kb, respectively. These values thus indicate that the tea genomes have relatively long LD distances and slow LD decays (Supplementary Fig. 17).

### Selective sweeps of the two major tea populations

It is generally stated that the differences between CSS and CSA teas lie primarily in their flavor, leaf and tree type, cold tolerance, and processing suitability. Among the accession assessed in the present study, the CSA population comprised three green tea accessions and 34 black tea accessions, whereas the CSS population contained 45 green tea accessions, 19 oolong tea accessions, and 11 black tea accessions (Fig. 3a). To determine the potential genetic foundation of these differences, we used SweepFinder2 (version 1.0) to scan for the selective sweep regions and selected those regions with the top 1% of composite likelihood ratio (CLR) scores and the genes overlapping with the final sweep regions (≥300 bp). On the basis of this analysis, we identified a total of 1,336 genes bearing selection signatures in the CSA populations, and 1,028 genes bearing selection signatures in the CSS populations (Supplementary Tables 19 and 20, Supplementary Fig. 18).

**Fig. 3.**
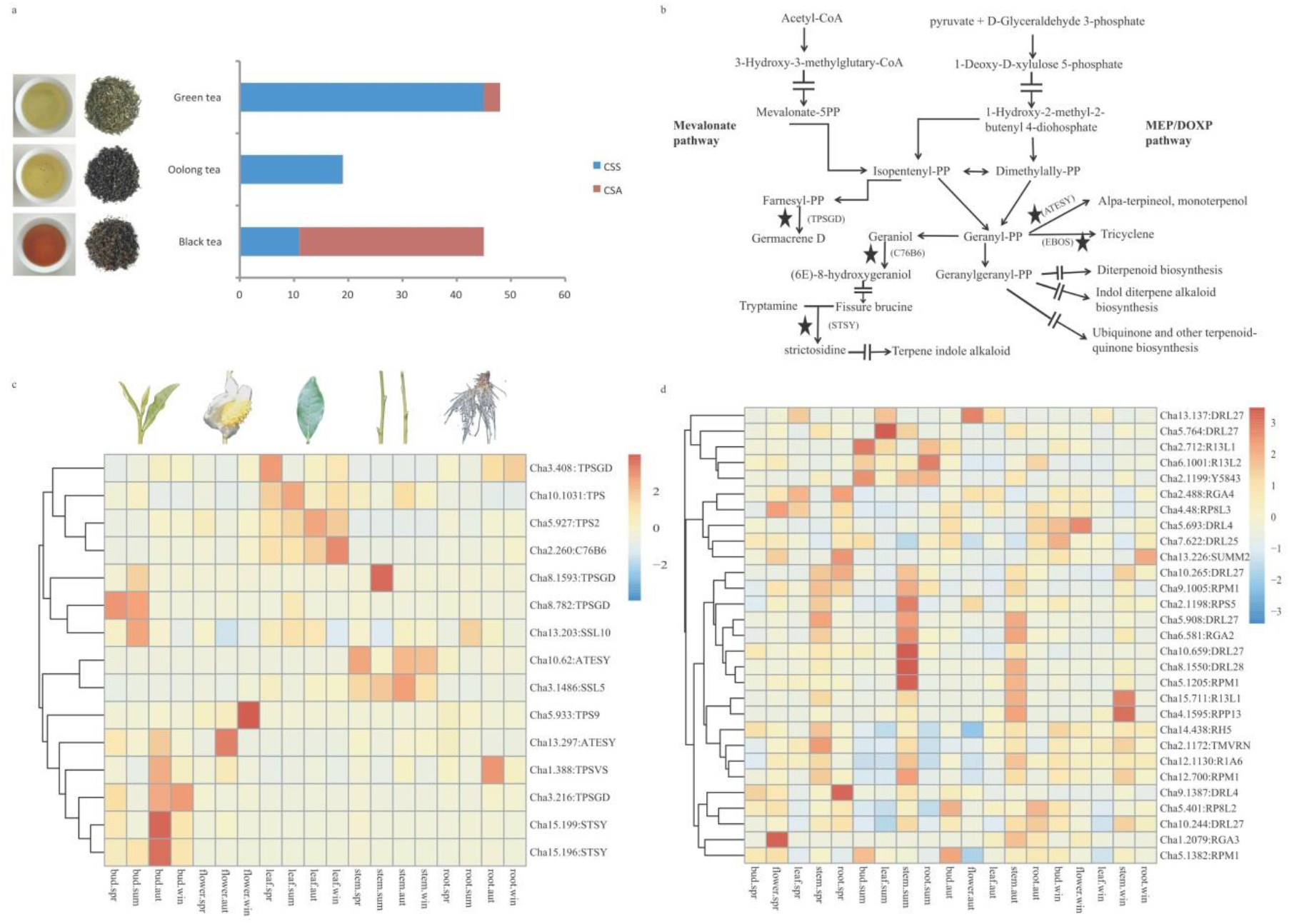
Sweep gene sets in *Camellia sinensis* var. *assamica* (CSA) and *C. sinensis* var. *sinensis* (CSS) show the different directions of domestication. a, The tea types were used to analyze the SweepFinder results of CSS and CSA. b, The pathway of terpene metabolism. The selective sweep genes are indicated by stars. The arrows bisected by equals symbols indicate hidden processes. c, The expression of terpene-related genes in different tea tissues. d, The expression of *NBS-ARC* genes in different tea tissues.

Based on GO analysis, enriched genes (P-value < 0.05, FDR < 0.05) were selected from the candidate selective sweep genes of the CSA and CSS populations (Supplementary Tables 21 and 22, Supplementary Fig. 19); we found that volatile terpene metabolism genes, such as cytochrome P450s (e.g., geraniol 8-hydroxylase) and terpene synthases, including alpha-terpineol synthase (*ATESY*), (-)-germacrene D synthase (*TPSGD*), and strictosidine synthse (*STSY*), were significantly selected in the CSS population but not in the CSA population (Fig. 3b, Supplementary Tables 21 and 22). The functionalization of core terpene molecules requires cytochrome P450s^32^, of which geraniol 8-hydroxylase catalyzes the conversion of geraniol (6E)-8-hydroxygeraniol (Fig. 3b), which may affect the accumulation level of geraniol. Alpha-terpineol is a monoterpene found in tea, which is generated by the ATESY-mediated catalysis of geranyl-PP, whereas TPSGD catalyzes the conversion of farneyl-PP to the sesquiterpene germacrene D. Strictosidine is the precursor of terpenoid indole alkaloids, and STSY is a key enzyme in the synthesis of these alkaloids (Fig. 3b). Moreover, we found that 80% of the selected terpene-related genes showed relatively high expression in buds or leaves, while 33% of the selected terpene-related genes showed significant high expression in buds or leaves (Fig. 3c, Supplementary Table 23).

Compared with CSA accessions, we also observed the selection of a larger number of *NBS-ARC* (nucleotide-binding site domain in apoptotic protease-activating factor-1, R proteins and *Caenorhabditis elegans* death-4 protein) genes in CSS accessions, the *Arabidopsis* homologs of which, including *RPS3* (also known as *RPM1*)^*33*^, *RPS5^34^*, and *SUMM2^35^*, have been shown to be involved in resistance to *Pseudomonas syringae* (*RPS*), (Supplementary Tables 21 and 22). The expression profiles of these genes revealed that 69% of the *NBS-ARC* genes subject to selection are highly expressed in spring, autumn, or winter, while 24% of the *NBS-ARC* genes are significant highly expressed in spring, autumn, or winter (Fig. 3d, Supplementary Table 24). However, among the 214 genes under selection in both CSS and CSA populations, we were unable to detect the enrichment of any genes related to flavor synthesis or abiotic and biotic stress resistance in the CSA population (Supplementary Tables 19 and 20).

## Discussion

This study represents the most comprehensive tea genome sequencing project conducted to date, and we present the first chromosome-scale genome sequence of tea and resequenced data of 139 tea accessions collected world. On the basis of our analyses, we have generated new resources, which will prove valuable for future tea-related genomics research and molecular breeding. These data reveal the genome-wide phylogeny of tea and the divergent selection direction between the two main tea varieties, namely CSS and CSA. In CSS, genes involved in flavor metabolism and cold tolerance were subjected to stronger selection than that in CSA; both traits were consistent with the fact that tea accessions from the east and north of China, like green and oolong tea, have a distinct aroma, and are cold tolerant. Our data also showed that the CSR population was the ancestral of the CSS and CSA, though this was a critical step toward the detail scenarios of the origin and domestication of CSS and CSA, the remain untold chapter need the identification of the closest ancestor of tea as well as the collection of more CSR in the future.

The first important criterion in a genome sequencing project is to obtain a high-quality reference genome and call an SNP set with high confidence from well-mapped resequencing data. In this regard, the inherent nature of the tea genome, notably its large size, high heterozygosity (Supplementary Table 25), and large number of repetitive sequences (Supplementary Tables 5 and 6), have previously led to difficulties in genome assembly. Although prior to the present study, the genomes of the YK10 and SCZ tea cultivars have been reported, these are characterized by relatively low continuity compared with that of the major currently assembled genomes (Mb scale) at both the contig and scaffold levels. Moreover, the associated BUSCO scores indicated that only approximately 80% of predicted genes could be identified in these genomes. Taking advantage of recent advances in sequencing and assembling technologies, we were able to sequence the genome of the LJ43 tea cultivar at the chromosome scale, generating an assembly characterized by a scaffold N50 value of 144 Mb, 88% gene completeness, and a base accuracy of 99.999%. There still needs improvement for the genome annotation in the future considering the complex of the tea genome. Combined with other analyses, our results showed that the quality of the LJ43 genome is higher than that of the previously published tea genomes^1,16^. Furthermore, our whole-genome sequencing of 139 worldwide tea accessions generated 6,272.74 Gb of short reads and 218.87 million high-confidence SNPs, and overall, the datasets obtained in the present study provide the richest genomic resource for tea researchers compiled to date.

*Camellia* is ranked as one of the most taxonomically and phylogenetically challenging plant taxa^12^, and we noted many disparities between assignments based on traditional taxonomic systems, which rely primarily on morphology, and our phylogenetic tree based on whole-genome sequencing analysis. Gene flow was widespread among tea accessions (Supplementary Material, Supplementary Table 26-28, Supplementary Fig. 19), and this presents challenges for the determination of the origin and evolution of tea. For example, *C. taliensis* (HZ122, HZ114) and *C. gymnogyna* (HZ104) have previously been assigned to the CSA population. Bitter tea, a hybrid progeny of CSS and CSA teas^36^, is a transitional type of large-leaved tea with a growth habit ranging from tree-like to shrub-like, and is mainly distributed in areas with mixed growth of CSS and CSA. In our phylogenetic tree, bitter teas (HZ039, HZ092, HZ080, and HSKC) were closely clustered with transitive teas in CSS and CSA, thereby supporting the fact that bitter tea is a hybrid progeny of CSS and CSA. It is expected that further worldwide sampling and more comprehensive data analysis will reduce current debates concerning tea taxonomy.

Unlike annual crops or perennial self-compatible crops, tea has not experienced severe domestication bottlenecks between wild progenitors and cultivated varieties^37^ (Supplementary Material, Supplementary Figs. 20 and 21), which can be attributed to the fact that the breeding of tea plants has largely been determined by environmental influences rather than human behavior, based on multiple generations of screening. During the expansion and domestication of tea, cultivated teas have been crossed with wild relatives, and this has contributed to the current genetic complexity of tea populations. This interbreeding is reflected by our observation that many cultivars and wild resources clustered together in the phylogenetic tree, with ancestral wild relatives appearing in the CSS cluster when a K value of 3 is used in the structural analysis (Supplementary Material, Supplementary Tables 16, 26-28, Supplementary Figs. 13 and 19). Although China has the longest tea cultivation history and the oldest written literature^4,38,39^ to support the hypothesis that tea plants originated in this country, there is still a lack of consensus regarding the events associated with the domestication of tea. In this regard, Meegahakumbura et al (2016) have suggested that the origins of CSS and CSA in China and CSA in India can probably be traced to three independent domestication events in three separate regions across China and India^40,41^, however, the lack of the convinced closest ancestor of both CSS and CSA in their analysis made the speculation doubtful. Our data showed that the CSR population was the ancestral of the CSS and CSA, though this was a critical step toward the detail scenarios of the origin and domestication of CSS and CSA, the remain untold chapter need the identification of the closest ancestor of tea as well as the collection of more CSR in the future. .

In the present study, we identified two interesting selection signatures in the CSS population, one of which is associated with genes involved in the terpene synthesis pathway. Terpene volatiles play essential roles in defining the characteristic aroma of tea, and the compositions and concentrations of theses volatiles are controlled at the genetic level^42^. Different species or varieties of tea plants are characterized by differences in terpene profiles, and in this regard, Takeo et al. found that the contents and ratios of linalool and its oxides are high in CSA, whereas the contents and ratios of geraniol and nerolidol are high in CSS^43–45^. The main terpenoids determining the aroma of black tea are linalool and its oxides, whereas geraniol and nerolidol contribute to the aroma of green tea and oolong tea^46^. These distinctions are consistent with the findings of our population selection analysis, which revealed that the terpene metabolism genes geraniol 8-hydroxylase, *ATESY*, *TPSGD*, and *STSY* have been significantly selected. In addition, our KEGG enrichment analysis of expanded gene families revealed that *TPSGD* is expanded in the LJ43 cultivar at the genomic level. Moreover, the flavor of different tea types has been influenced to a certain extent by consumer predilection and culture. On the basis of the processing suitability of CSA and CSS and the population selection analysis of the two populations, we can conclude that terpenoid metabolism is more closely related to the aroma of green and oolong tea than it is to that of black tea.

The second selection signature of interest identified in the present study relates to the *NB-ARC* genes in the CSS population. Most of these genes are associated with resistance to ice nucleation active (INA) bacteria. In *Arabidopsis*, *RPS3*/*RPM*^33^, *RPS5*^34^, and *SUMM2*^35^ have been shown to confer resistance to *Pseudomonas syringae*, which is one of the most well-studied plant pathogens that can infect almost all economically important crop species. In addition, *Pseudomonas syringae* is a prominent INA bacterium and has been proposed to be an essential factor contributing to frost injury in agricultural crops^47^. Mutants characterized by alterations in the aforementioned genes have also been found to show sensitivity to chilling temperature compared with the corresponding wild-type plants^33–35^. Similarly, in wild potato (*Solanum bulbocastanum*), the *RGA2*^48^ and *R1A6* are involved in resistance to *Phytophthora infestans*, a further factor related to INA bacteria. Moreover, significant differences have been detected in the expression of *RPS3* and *SUMM2* in cold-resistant and cold-susceptible cultivars^49^. Taken together, the results of these studies tend to indicate that *NB-ARC* genes might play an important role in endowing CSS cultivars with cold tolerance. Tea grown along the Yangtze River Basin and in eastern China is typically subjected to low temperatures in early spring and winter, and most CSA cultivars, which are characterized by large leaves, cannot survive in these areas. Some CSS adapted to cold environments survived during the expansion and domestication in eastern and northern China and after the separation of CSS and CSA, the direction of the domestication of these two varieties is assumed to have diverged. With the increase in tea consumption, humans began to select tea plants, and during domestication, selection for flavor and cold tolerance has been stronger in CSS than that in CSA. This is also reflected at the genomic level, as illustrated by the KEGG enrichment of expanded gene families, in which the disease resistance proteins RPS2 and RPS3 were found to be expanded in LJ43. Although in the present study, we found that 214 genes had undergone selection in both the CSS and CSA populations, we were unable to detect enrichment of any of the genes associated with flavor and resistance in the CSA population (Supplementary Table 21). It indicates that the selection for INA bacterial resistance and flavor during domestication has been stronger in CSS than in CSA.

## Methods

### Materials and sequencing

We collected samples of 139 tea accessions from around the world (detailed information is presented in Supplementary Table 16). Among these, 93 samples were collected from China, with the remaining 46 samples being collected from the other main tea-producing countries. For the purpose of analyses, we selected *Camellia sasanqua* Thunb. as an outgroup. DNA was extracted from the leaf tissues of all samples using the CTAB method^50^. Libraries for Illumina truseq, 10x Genomics, and PacBio analyses were prepared according to the respective manufacturer’s instructions. The detailed sequencing information is presented in Supplementary Material.

### Genome assembly and annotation

The detailed information of genome size and genome assembly is presented in Supplementary Material. Assembly of the LJ43 genome was performed based on the Hi-C-Pro pipeline and full PacBio reads using WTDBG (version 1.2.8). The final Hi-C assisted genome assembly was commissioned by Annoroad Gene Technology. Tigmint (version 1.1.2)^51^ was used to find errors using linked reads from 10x Genomics Chromium. The reads were first aligned to the Hi-C scaffolds, and the extents of the large DNA molecules were inferred from the alignments of the reads. For larger-scale gaps, we mapped optical maps from BioNano Genomics to the Hi-C scaffolds using the BioNano Solve 3.3 analysis pipeline. A high-density genetic linkage map^20^ was used to carry the genomic synteny analysis. The markers were first aligned to the Hi-C scaffolds using “bwa mem (version 0.7.15).” Properly mapped alignments with mapping quality >1 were extracted (3,483). Dot plots were plotted and correlations were calculated with the extracted alignments using R (version 3.4). Repeat sequences were identified using *de novo* and homology-based methods. Augustus^52^ and GlimmHMM^53^ were used to analyze *ab initio* gene prediction with parameters trained using unigenes. For homology-based predictions, we used the homologous proteins proposed for the genomes of *Arabidopsis thaliana*^54^, *Oryza sativa* subsp. *japonica*^55^, *Coffea canephora*^56^, *Theobroma cacao*^57^, and *Vitis vinifera*^58^. RNA was extracted from five tissue types (bud, leaf, flower, stem, and root) at four time points (except for flowers during summer). Three biological replicates were set for each sample (Supplementary Table 7), and the transcript reads were assembled using Cufflinks (version 2.2.1). All of the predicted gene structures were integrated using EVidenceModeler (version 1.1.1)^59^. Protein-coding genes with both of their CDS length shorter than 300 nt and with stop codons were filtered (except stop codons at the end of a sequence). Then, we mapped RNA-seq reads against the predicted coding regions by SOAP2^60^, and selected the predicted gene regions by RNA-seq data (regions with >50% coverage). The method of gene annotation is described in detail in Supplementary Material. The method of functional annotation is described in detail in Supplementary Material. The protein sequences of LJ43 and *Actinidia chinensis*^21^ were analyzed by blastp with the parameters -evalue 1e-5 -num_alignments 5. Then syntenic blocks were identified by MCScanX^61^ with the parameters –e 1e-20. SCZ and YK10 were analyzed with the same pipeline and parameters. The genome synteny between *Theobroma cacao* ^57^ and LJ43, SCZ and YK10, respectively was also analyzed (Supplementary Material).

### Positive Darwinian selection analyses

The species tree was constructed as described in Supplementary Material, without SCZ and YK10. We identified 1,031 single-copy gene families. The protein sequences of single-copy genes were aligned by clustalw2^62^, and then the out of clustalw2 was transformed to nuclear format according to alignment protein sequences using our own Perl script. Gblocks^63^ was used to cut the nuclear alignment sequences by t=c parameter. “Branch-site” models A and Test2 were chosen to test positive selection by codeml of PAML. The significant sites were dropped if 5 bp around the site sequences was cut by Gblocks. The False Discovery Rate (FDR) was used to filter the results (FDR ≤ 0.05).

### SNP calling and filtering

Quality controlled reads were mapped to the unmasked tea genome using bwa (version 0.7.15)^64^ with the default parameters. SAMtools (version 1.4)^65^ was used for sorting and Picard (v.2.17.0) was used for removing duplicates. The HaplotypeCaller of GATK (version 3.8.0)^65^ was used to construct general variant calling files for the tea group (139 accessions) and outgroup (*C. sanqua*, CM-1) by invoking the options of -ERC:GVCF. The gVCF files in the tea group were combined using GenotypeGVCFs in GATK to form a single variant calling file, whereas the gVCF file in the outgroup was called with the option “–allSites” to include all sites. The final single variant calling file was merged using bcftools (version 1.6), with only the consistent positions retained in both groups. To obtain high-quality SNPs, we initially used the GATK Hard-filter to filter the merged VCF with the options (QD ≥ 2.0 && FS ≤ 60.0 && MQ ≥ 40.0 && MQRankSum ≥ −12.5 && ReadPosRankSum ≥ −8.0). Thereafter, we performed strict filtering of the SNP calls based on the following criteria: (1) sites were located at a distance of least 5 bp from a predicted insertion/deletion; (2) the consensus quality was ≥40; (3) sites did not have triallelic alleles or InDels; (4) the depth ranged from 2.5% to 97.5% in the depth quartile; and (5) SNPs had minor allele frequencies (MAF) ≥0.01.

### Population genetic analyses

We selected high-quality SNPs with a maximum of 20% missing data, and to eliminate the potential effects of physical linkage among variants, the sites were thinned such that no two sites were within the same 2,000-bp region of sequence. Phylogenetic analysis was conducted with the final SNP set using iqtree (version 1.6.9)^66–68^. A maximum likelihood (ML) phylogenetic tree was calculated using the GTR+F+R5 model, and 1,000 rapid bootstrap replicates were conducted to determine branch confidence values. The best-fitting model was estimated by ModelFinder implemented in IQTree after testing 286 DNA models. GTR+F+R5 was chosen according to BIC (Bayesian Information Criterion). The ML phylogenetic tree was constructed by inter gene region SNP. The ML phylogenetic trees were also constructed using the final SNP set and 4DTV SNP. Principal component analysis (PCA) was performed using PLINK (version 1.90) on the final SNP set, with the principal components being plotted against one another using R 3.4 to visualize patterns of genetic variation. We also used the final SNP set for population structure analysis using ADMIXTURE (version 1.3)^69^, which was run with K values (number of assumed ancestral components) ranging from 1 to 10.

Population heterozygosity at a given locus was computed as the fraction of heterozygous individuals among all individuals in a given population. The average heterozygosity was then calculated for each 40-kb sliding window, with a step size of 20 kb. Individual heterozygosity was computed as the fraction of loci that are heterozygous in an individual. Average heterozygosity was also calculated using the same method. Windows with an average depth <1 were filtered out.

In order to eliminate the influence of the difference in sample number, eight samples of the CSR/CSA/CSS populations were randomly selected to calculate the nucleotide diversity. We repeated 20 times for each population to reduce the sampling error. Vcftools (version 0.1.16) with the window size 50kb and the stepping size 10kb was used to calculate the nucleotide diversity of the sample population. For each population, all the 20 results were collected, and the boxplot was plotted with R.

### Selective sweep analysis

Treetime 0.5.3^70^ was used to infer the ancestral state based on ML using the generated evolutionary tree. Sites lacking a reconstructed ancestral state in a population were folded in the SweepFinder2 analysis. We excluded sites that were neither polymorphic nor substitutions, as recommended by the SweepFinder2 manual^71^. To reduce false positives, the chromosome-wide frequency spectrum was calculated as the background for each chromosome and for each population. SweepFinder2 was run with a grid size of 100. The CLR scores from the SweepFinder2 results were extracted, and scores were merged into sweep regions when the neighboring score(s) exceeded a certain threshold, which was set as the top 1% of CLR scores. To obtain regions with greater continuity, we merged regions into a single region with a certain size threshold between regions; the threshold was set to 50% of the size in adjacent sweep regions. The final score for each sweep region was the sum of the CLR scores of the sites in the sweep region. The final sweep regions were filtered based on a minimum size of 300 bp. Gene overlap within the sweep regions was extracted as the candidate selective sweep genes. The GO-enriched (P-value < 0.05, FDR < 0.05) candidate selective sweep genes were selected, and the *Fst, θ*_*π*_ and Tajima’s D values were calculated using vcftools, with a window size of 50,000 bp and a step size of 10,000 bp..

### Gene expression

Transcript-level expression was calculated using HISAT2, StringTie, and Ballgown with the default parameters^72^. The genes identified in the selection results were selected for expression analysis, and an expression heat map was plotted using the Heatmap package in R 3.4. The average expression of selection genes in Fig. 3d was calculated by seasons, while the average expression of selection genes in Fig. 3c was calculated by tissues. Student’s T-test was used to identify the significantly different genes (P-value < 0.05).

### Data availability

The raw sequence data, genome sequence data, and genes sequence data reported in this paper have been deposited in the Genome Sequence Archive^73^ in BIG Data Center^74^ (Nucleic Acids Res 2019), Beijing Institute of Genomics (BIG), Chinese Academy of Sciences, under accession numbers PRJCA001158, PRJCA001158 that are publicly accessible at https://bigd.big.ac.cn/gsa.

## Acknowledgements

This work was funded by Agricultural Science and Technology Innovation Program of the Chinese Academy of Agricultural Sciences (CAAS-ASTIP-2017-TRICAAS and CAAS-ASTIP-2017-AGISCAAS), the Agricultural Science and Technology Innovation Program Cooperation and Innovation Mission (CAAS-XTCX2016), the Major Science and Technology Special Project of Variety Breeding of Zhejiang Province (2016C02053), the Shenzhen Science and Technology Research Funding (JSGG20160429104101251), and the National Youth Talent Support Program and the Program for the Innovative Research Team of Yunnan Province. We thank Hualing Huang (Tea Research Institute of Guangdong Agricultural Academy of Sciences), Haitao Zheng (Rizhao Tea Research Institute), and Lizhe Lv (Xinyang Tea Research Institute) for supplying tea plant samples. We thank Xiujuan Shao for analyzing the gene annotation of LJ43.

## Author contributions

X.W., Y.C., G.W., L.C., J.R., and Y.Y. designed the experiments and managed the project. X.W., F.H., Y.C., C.M., L.Y.W., X.H., and A.L., wrote the manuscript with input from all authors. X.W., F.H., Y.C., C.M., X.H., A.L., H.C., J.J., L.W., K.W., X.B.W, C.A., Z.W., S.Z., P.C., Y.L., B.L., G.W., L.C., J.R., and Y.Y. collected the samples, extracted genetic material, analyzed the data, and performed the experiments. X.W., Y.C., C.M., X.H., and S.Z. performed experiments and the genomic and RNA-sequencing. J.R. performed the genome assembly analyses. H.F. and X.B.W. performed the gene annotation analyses. H.F., X.H., A.L., and C.A. performed transcriptomic analyses. X.W., H.F., A.L., and G.W. performed population analyses. X.W., Y.C., P.C., L.C., G.W., J.R., and Y.Y. revised the manuscript.

## Competing interests

The authors declare no competing interests.

