## Supplementary material for "Population sequencing enhances understanding of tea plant evolution"

[1] Key Laboratory of Tea Biology and Resources Utilization, Ministry of Agriculture and Rural Affairs, National Center for Tea Plant Improvement, Tea Research Institute, Chinese Academy of Agricultural Sciences, Hangzhou, China. [2] Lingnan Guangdong Laboratory of Modern Agriculture, Genome Analysis Laboratory of the Ministry of Agriculture and Rural Affairs, Agricultural Genomics Institute at Shenzhen, Chinese Academy of Agricultural Sciences, Shenzhen, China. [3] Tea Research Institute, Yunnan Academy of Agricultural Sciences, Menghai, China. [4] State Key Laboratory of Genetic Resources and Evolution, Kunming Institute of Zoology, Chinese Academy of Sciences, Kunming, China. [5]Center for Excellence in Animal Evolution and Genetics, Chinese Academy of Sciences, Kunming 650223, China. [6] These authors contributed equally: Xinchao Wang, Hu Feng, Yuxiao Chang, Chunlei Ma, Liyuan Wang, Xinyuan Hao. *

### Supplementary Note

### 1 Sequencing and Assembly of the Longjing43 genome

#### 1.1 Plant material

The tea cultivar Longjing 43 (LJ43, *Camellia sinensis var. sinensis cv.* Longjing 43), a very popular and famous cultivar for preparing West Lake Longjing green tea in China, was selected for *de novo* assembly sequencing. LJ43 is a line selected from the populations of "Longjing Quntizhong", an old population land race growing in Hangzhou (TRI, CAAS, N 30°10', E 120°5') tea production areas. Since LJ43 showed excellent tea agronomic traits including early sprouting, high tea quality, and cold-resistance, it is one of the largest acreages of tea plant cultivars in China and its total cultivation area exceeds 150,000 ha in more than 10 provinces in China. Moreover, it supplies sustainable income of more than ten billion RMB Yuan for millions of farmers per year in China.

#### 1.2 Genomic DNA preparation and sequencing

For LJ43 Illumina shotgun library preparation, DNA was isolated from fresh leaves by 2% cetyltrimethylammonium bromide (CTAB) according to a previously published protocol[1], Truseq library was prepared using the KAPA Hyper Prep Kit (Illumina® platforms, KAPA BIOSYSTEMS, Boston, MA, USA. Cat No. KK8504) following the manufacturers' manual. All the libraries were sequenced by Hiseq 4000 or HiSeq X® (Illumina®, San Diego, CA, USA) according to the manufacturer's instructions.

##### 1.3 Pacific Biosciences single molecular long-read sequencing

For Pacific Biosciences (PacBio) single molecular long-read sequencing, high molecular weight genomic DNA was isolated by the CTAB method[1]. SMRTbell 25 kb needle sheared library was constructed, size-selected with 0.375x SPRI beads, and sequenced using P6/C4 chemistry in RSII (180-min movie) according to the manufacturer's instructions. We generated 196 Gb (approximately 60-fold) raw data with the read N50 of 12.5 kb and an average length of 9.1 kb.

##### 1.4 Bionano genomics optical mapping data generation

For Bionano genomics optical mapping data generation, megabase-containing genomic DNA from tender shoot cultivated under dark was prepared by Bionano Prep™ Plant Tissue DNA Isolation Kit (Bionano Genomics, Inc. San Diego, CA, USA, Cat No. RE-014-05) according to the manufacturers' manual. The genomic DNA was fluorescently labeled using the nicking endonuclease Nt.BspQI, and stained according to the manual of the IrysPrep Reagent Kit (BioNano Genomics, Inc. San Diego, CA, USA). Then the stained DNA sample was loaded onto the nanochannel array of IrysChip and imaged by the Irys system (BioNano Genomics, Inc. San Diego, CA, USA).

##### 1.5 RNA isolation, RNA-seq library preparation and sequencing quality control

Total RNA from tissues of LJ43 (Supplementary Table 7) was isolated with the RNAprep Pure Plant Kit (TIANGEN Biotech CO., LTD, Beijing, China, Cat No.

DP432), and the RNA-seq library was prepared using the KAPA RNA Hyper Prep Kit (Illumina® platforms, KAPA BIOSYSTEMS, Boston, MA, USA. Cat No. KK8541) according to the manufacturers' manual. All the RNA-seq libraries were sequenced by HiSeq X® (Illumina®, San Diego, CA, USA).

##### 1.6 Sequencing Quality control

DNA and RNA sequencing reads were trimmed and filtered using Trimmomatic (version 0.36.5)[2] after the first round of quality control using FastQC (version 0.11.5). The adapters and low-quality bases (Phred score <20) were removed from the leading and trailing of the reads. The reads were then scanned with a 4-base wide sliding window, and cut when the average quality per base within the window dropped below 15. Reads with a length of <75 bp were dropped and a second round of quality control was performed using FastQC to ensure the quality of the trimmed data.

##### 1.7 Genome assembly

###### 1.7.1 Estimation of genome size

The genome size of LJ43 was estimated by three approaches. First, using the Angiosperm DNA C-values Database (http://data.kew.org/cvalues/release 8.0, Dec 2012), where the DNA C-values of *Camellia sinensis* Kuntze were estimated by Feulgen microdensitometry and the 1C DNA amount in megabase pairs was 3,824 Mb [3]. Second, using an optimized DNA flow cytometry method for genome size estimation according to Huang's method[4,5] in BD FACSCalibur (Becton Dickinson, San Jose, CA,

USA) flow cytometer (Supplementary Fig. 1). Third, exploiting KmerGenie (version1.7051)[6] to estimate genome size with 214 Gb Illumina short reads. KmerGenie was run with different k-mer lengths ranging from 17 to 127 with a step size of 10. K-mer abundance histogram was computed and the best possible k-mer length was chosen. The predicted genome size was about 3.32G with the best k-mer length 97 (Supplementary Fig. 2).

##### 1.7.2 LJ43 genome assembly

About 196 Gb (approximately 60-fold) PacBio reads were used for the LJ43 genome assembly with WTDBG (version 1.2.8). WTDBG was ran with the parameters (-fo dbg --load-alignments dbg.alignments --edge-min 3 --rescue-low-cov-edges). The assembly genome was corrected with PacBio reads and around 214 Gb (approximately 66-fold) Illumina PE 150 reads. Firstly, PacBio reads were used to correct the genome by arrow (version 2.1.0) with default parameters**.** Following that, Illumina short reads were mapped to previous step corrected genome by bwa (version 0.7.15)[7] with default parameters, variant was called using bcftools (version 1.6) and all of the homozygous mutation sites were removed by the in house developed script. Our principle was that polishing stops when the number of corrected bases reached plateau. After 7 rounds of Illumina reads correction, a version of 3.2 Gb tea genome with a contig number of 37,600, contig N50 of 271.33 kb, and GC content of 38.67% was obtained. We compared the genome with published Yunkang10 (YK10) and Shuchazao (SCZ) (Table 1).

##### 1.7.3 Hi-C library preparation and sequencing

The leaves of LJ43 were treated with formaldehyde to fix nuclear chromatin. The fixed chromatins were digested with the MboI enzyme. The free blunt ends were ligated by biotinylated nucleotides. Then, the DNA was purified, and the fragments with biotinylated nucleotides were extracted. Sequencing libraries were generated by the manufacturer's instructions (Illumina). After PCR enrichment, three libraries were sequenced and produced 263 Gb PE150 clean data.

##### 1.7.4 Hi-C assisted genome assembly

The 10x reads were first processed by Long Ranger v2.2.2 to create an interleaved file of barcoded pair-end reads. Then, we aligned the barcoded pair-end reads to the genome with BWA-MEM (version 0.7.15) with '-pC' parameters. The alignment file was processed using ARCS (version 1.0.6) + LINKS (version 1.8.6) pipeline ARCS was used to create a Graphviz Dot file (.gv) with contig head/tail length for masking alignments set to 50 kb. LINKS was then used to join nodes in the graph produced by ARCS with default parameters8,9. Both of the contig versions obtained by the PacBio assembly and the scaffold version obtained by the PacBio assembly with 10x scaffolding were used as input for Hi-C scaffolding. Clean paired-end reads were aligned to the genome using BWA (version 0.7.15). Then the mapping results were filtered with a mapping quality (≧20) and edit distance (NM≦5). We also filtered the alignment file to only keep reads aligned to the region within 500bp around a restriction site. The final alignment file was fed to Lachesis. To arrive at the final set of Lachesis parameters, we randomly varied the parameters through 10,000 scaffolding iterations. These randomized parameter sweeps varied within the following bounds: CLUSTER_MIN_RE_SITES between 1 and 5,000; CLUSTER_MAX_LINK_DENSITY between 1 and 30; ORDER_MIN_N_RES_IN_TRUNK between 1 and 5000; ORDER_MIN_N_RES_IN_SHREDS between 1 and 5000. The result of the contig version of the genome shown fraction of sequences in orderings with high orientation quality: 17,715 (76.47%), with length 2,801,038,355 (93.56%). The result of the scaffold version of the genome shown fraction of sequences in orderings with high orientation quality: 12,288 (59.31%), with length 2,840,497,749 (92.13%). The interaction heatmap of the scaffold version showed more errors than the contig version (Supplementary Fig. 3 and 4). Moreover, when we compared collinear protein blocks with *Actinidia chinensis* using MCScanX. The results showed that the contig version also showed better results (3,205 genes in block of the contig version, vs 3,053 of the scaffold version), indicating that incorrectly joined scaffolds were probably by 10x data. Considering this, we finally used the contig version for HiC anchoring. After the preliminary test above, we entrusted Annoroad Gene Technology for further polishing for the limited computing resources.

The final Hi-C assisted genome assembly was commissioned by Annoroad Gene Technology. Around 1,266,516,127 clean paired-end reads were used to improve the LJ43 genome assembled by the PacBio reads using HiC-Pro (version 2.7.8)[10]. Firstly, the reads were mapped to genome (PacBio assembly genome) by bowtie2. Then the results were filtered by extracting the unique mapped paired-end reads. HiC-Pro was used to locate the unique mapping paired-end reads to contigs. Lachesis[11] was used to scaffold the contigs to 15 chromatin clusters by agglomerative hierarchical clustering (Supplementary Tables 2 and 3, Fig. 1b, Supplementary Fig. 5). A total of 7,071 contigs with 2,311,549,792 bp (70.9%) were ordered with orientation. The resulted scaffold N50 was 143,847,529 bp.

##### 1.8 Evaluation of the LJ43 genome assembly quality

A total of 49,529 ESTs of tea were downloaded from NCBI, and mapped to the tree tea genome by gmap (2017-10-30)[12]. We filtered the results by coverage ≥ 90% and identity ≥ 90%. 35,240 (about 71.15%) ESTs were mapping to LJ43, 35,202 (about 71.07%) ESTs were mapping to SCZ, 31,303 (about 63.20%) ESTs were mapping to YK10. The results shown the LJ43 genome were more complete.

We evaluated the completeness at genome level by benchmarking universal single-copy orthologs (BUSCOs) , and found that the completeness value of LJ43 was 90.3% with 3.2% fragment and 6.5% missing. Afterwards, we also checked whether the completeness of the gene annotation was caused by incompleteness assembly of genome. The missing part position of fragement genes were extracted from the alignment results of BUSCO and compared to annotation file. If the fragment genes were due to the break of contig, the missing parts would be located at the either end of contig. We found that 24 (out of 90) fragment genes of LJ43 were due to the break of contig, that was due to the complexity and heterozygosity of the tea genome. Though we attempted to increase the number of complete genes, the break of contigs resulted in 24 fragmented genes.

#### 2 Genome Annotation

##### 2.1 Repeat sequences and Transposable Elements

Repeat sequences were identified by combining the *de novo* annotation and homology-based methods. For the *de novo* repeat sequences prediction, RepeatModeler (1.0.4, http://www.repeatmasker.org/RepeatModeler.html) was used to search for repetitive sequences in the genome, and then the results were used to build a repeat sequences library. After that, RepeatMasker (v. 2.1, http://www.repeatmasker.org) was applied to identify repeat sequences by the repeat sequences library (Supplementary Table 5). For the homology-based prediction, the genome assembly was compared to the Repbase of RepeatMasker and RepeatProteinMask. Then, the predicted transposable elements (TEs) were combined by removing redundant TEs. TEs repeat annotation was revealed to be up to ~2.30 Gb, and comprised about 70.44% of the tea genome (Supplementary Table 6).

We used LTR-finder (version 1.05)[13] to search the LJ43 genome, and 35,380 intact LTR retrotransposons were obtained. Then, the 5' and 3'-LTR sequences were aligned with Muscle (version 3.8.31)[14] and the Kimura two-parameter distance was calculated using EMBOSS (version 6.4.0) for each intact LTR. The insertion time between varieties was calculated according to the formula of Time=Ks/2μ ($\mu = 6.5\times10^{-9}$ mutations per site per year). The SCZ and YK10 were analyzed in the same way.

Comparison of the results of the three tea genomes showed that LJ43 had more recently inserted LTR retrotransposons (Supplementary Fig. 6). We compared the PacBio reads corrected genome and NGS reads corrected genome to verify that the different 5' and 3' terminal IR sequences of LTR were real and not caused by NGS reads correction (Supplementary Fig. 6d). The presence of more recent LTR retrotransposons in the LJ43 genome, indicated that its genome is more complete compared to SCZ and YK10. LTR-retriever[15] was used to identify long repeat retrotransposons, then LAI[16] was used to evaluate the LTR assembly index.

#### 2.2 Protein coding gene prediction

The protein coding genes were annotated using the strategy of combining the *ab initio* prediction and the homology-based predictions. To assist the protein coding gene prediction, we generated a total of approximately 340 Gb RNA-seq clean data from 19 samples including 5 tissues (bud, leaf, flower, stem, and root) in four seasons (except for flower during summer), and three biological replicates for each sample (Supplementary Table 7). First, we used PASA (version 2.0.0)[17] to build a comprehensive transcriptome library. Then unigenes with length of CDS longer than 900 bp and all vs all identity less than 70% were selected to train Augustus (version 3.3)[18] and GlimmHMM (version 3.0.4)[19]. Afterwards, Augustus and GlimmHMM with parameters were trained by the selected unigenes for *ab initio* prediction. For the homology-based predictions, we used the homologous proteins annotated in the genomes of arabidopsis[20], rice[21], coffee[22], coca[23], and grape[24]. First, GenblastA[25] was used to cluster the adjacent HSPs (high-scoring pairs) from the same protein alignments, and GeneWise (version 2.4.1)[26] was used to identify the accurate gene structures. Then, RNA-seq clean reads were mapped to the LJ43 genome by Tophat2[27]. Subsequently, Cufflinks (version 2.2.1) was used to predict gene models. All of the above individual results were integrated with EVidenceModeler (version 1.1.1)[28] and protein coding genes with both of their CDS length shorter than 300 nt and with stop codon were filtered (except stop codon at the end of the sequence). Then, RNA-seq reads were mapped against the predicted coding regions by Soap2[29], and the predicted gene regions were selected by RNA-seq data (the coverage >50%). Finally, a total of 33,556 genes supported by transcription reads were identified in the annotation. To calculate the transcript-level expression, RNA-seq reads were mapped to genome by HISAT2 (version 2.1.0) with default parameters, and transcript-level expression was analyzed by StringTie (version 1.3.3b) and Ballgown with default parameters[30].

The average length of LJ43 (10,816 kb) was longer than shuchazao[31] (SCZ, 7,386 kb) and Yunkang 10[32] (YK10, 3,549 kb). The completeness of gene set of LJ43 was higher than SCZ and YK10, and this may explain why the average gene length of LJ43 was longer than SCZ and YK10. We also compared the genes length and genes length distribution of LJ43 to six species (SCZ[31], YK10[32], citrus[33], *Amborella trichopoda*[34], *Actinidia Chinensis*[35], *and Gingko biloba*[36]). The average gene length of LJ43, SCZ, YK10, citrus, *Amborella trichpooda*, *Actinidia Chinensis*, and *Gingko biloba* were 10,816 bp, 7,386 bp, 3,549 bp, 4,061 bp, 11,053 bp, 5,388 bp, and 25,619 bp, respectively. LJ43 and SCZ had more genes with length between 10-50 kb, LJ43,

*Amborella trichpooda* and *Ginkgo biloba* had more genes with length ≥ 50kb (Supplementary Fig. 7). The average length of LJ43 was similar to *Amborella trichpooda*.

If the long gene among our gene annotation was reliable, the exon-exon conjunction in the longest and shortest gene set should have similar ratio supported with RNA-seq. Thus, we selected genes with highly expressed (FPKM values >= 20) in any tissue or period with more than two exons (14,698 genes). Among them, the top 1,000 longest genes contain 11,446 exon-exon conjunctions, and 4,796 were supported by RNA-seq reads. The 1,000 shortest genes contained 1,270 exon-exon conjunctions, and 631 were supported by transcription reads. The supported and unsupported exon-exon of longest genes and shortest genes were tested by Chi-square test of R, the P-value was 0.5345, indicating that there is no significant difference in the reliability between the longest genes and shortest genes. The gene annotation of LJ43 was reliable.

##### 2.3 Evaluation of the genome annotation

For further quantitative assessment of the annotation completeness, the three genome annotations were evaluated by BUSCO[39] with the same default parameters. The database was embryophyta ortholog data base 10 (embryophyta_odb10, https://busco.ezlab.org/datasets/prerelease/embryophyta_odb10.tar.gz). LJ43, SCZ and YK10 had 88.4%, 80.6% and 68.6% complete genes, respectively (Table 1). The detailed results of LJ43 were 88.4% complete (single:80.9%, duplication:7.5%), 6.9% fragment,and 4.7% missing; The results for SCZ were 80.6% complete (single:73.3%, duplication:7.3%], 9.5% fragment, and 9.9% missing; The results for YK10 were 68.6% complete (single:64.2%, duplication:4.4%), 16.7% fragment, and 14.7% missing. Total BUSCO gene number is 1,375.

##### 2.4 Homology search and functional annotation of the LJ43 genome

All of the predicted genes were functionally annotated according to homologous alignments with BLASTP (e-value ≤1e-5) against the Swiss-Prot and TrEMBL databases. InterProScan (version 5.21)[40] was further used to predict gene ontologies (GO terms) and domain information. Kyoto encyclopedia of genes and genomes (KEGG) automatic annotation server (KAAS) was used to assign the putative gene function to the KEGG pathway. Using homologous alignments and domain scanning, integrated with pathway annotation, 93.69% (31,437) of the protein-coding genes had significant similarities in functional protein databases, among them, in 78.59% (26,373), 91.35% (30,655), 59.71% (20,035), and 25.76% (8,643) could be assigned functions by Swissprot, InterPro, GO, and KEGG, respectively (Supplementary Table 8).

#### 3. Comparative Genomics

##### 3.1 Syntonic blocks analysis

The protein sequences of LJ43 and *Actinidia chinensis*[35] were analyzed by blastp with the parameters -evalue 1e-5 -num_alignments 5. Then, syntenic blocks were identified by MCscanX[43] with the parameters –e 1e-20. SCZ and YK10 were analyzed with the same pipeline and parameters. The genome synteny between *Theobroma* *cacao*[23] to LJ43, SCZ and YK10, was also analyzed. Compared to *Actinidia chinensis*[35], the genome of LJ43, SCZ and YK10 contained 690, 111 and 54 colinear blocks, respectively. A total of 18,030, 1,487, and 393 genes were involved in above collinear blocks, respectively. Compared to *Theobroma cacao* L., LJ43 had 413 colinear blocks with 14,661 genes; SCZ had 233 colinear blocks with 3,047 genes; YK10 had 0 colinear blocks with 0 genes.

##### 3.2 The construction of Phylogenetic tree and the estimation of gain and loss of gene families

To characterize the gene families experienced gene gain and loss in the tea genome, a phylogenetic tree was constructed for LJ43, *Actinidia Chinensis*[35], *coffea*[22], *Theobroma cacao*[23], *Arabidopsis thaliana*[20], *Oryza sativa* subsp. *japonica*[21], *Populus trichocarpa*[45], *Amborella trichopoda*[46], and *Vitis vinifera*[24]. A total of 1,031 single-copy gene families were identified among 9 genomes. The longest alternative splicing genes were chosen to reconstruct the phylogeny. The OrthoMCL[47] was used to cluster the genes family. The coding sequences of the single-copy genes were concatenated to a super-gene for each species. The super-genes were aligned by MAFFT[48]. The alignment sequences were used for phylogenetic analyses by raxml-HPC-MPI-AVX[49]. *Amborella trichopoda* was chosen as the out-group in our analysis. The output of OrthoMCL and phylogenic tree structure were used for a computational analysis of changes in gene family size with the software CAFE.

##### 3.3 Whole genome duplication (WGD) of LJ43 and diversity of three teas

We selected SCZ, YK10 and 9 additional species in 3.2 to build gene families by OrthoMCL. To estimate the divergence time of the LJ43 paralogs, we selected gene families consisting of exactly 2 tea genes to calculate the Ks of the pairs. We obtained 3,233 tea 2-member gene clusters. Yn00 of PAML[50] was used to calculate the ks value. The peak of Ks distribution of gene pairs was about 0.31. The divergence time was calculated according to the formula of Time=Ks/2μ, and was based on the molecular clock (μ) of a substitution rate of $6.1 \times 10^{-9}$ mutations per site per year for eudicots[51].

MCXscanX was used to detect the syntenic genes of LJ43 and SCZ. We selected the orthologous genes to calculate the Ks by Yn00 of PAML. This approach was used also for YK10. The peak of Ks distribution of LJ43 and SCZ gene pairs was about 0.003. The peak of Ks distribution of LJ43 and YK10 gene pairs was about 0.045 (Supplementary Fig. 11).

#### 4 Population Genomics

##### 4.1 Samples and DNA extraction and sequencing for population data

The 139 tea accessions with a wide range of species distribution were collected from all the world, 105 from East Asia, 7 from South Asia, 9 from Southeast Asia, 6 from West Asia, 7 from Africa, and 5 from Hawaii (Fig. 2a, Supplementary Table 16). Average sequenced depth was approximately 13-fold (Supplement Table 16).

DNA was isolated from fresh leaves by 2% CTAB according to a previously published protocol[1]. Truseq library was prepared using the KAPA Hyper Prep Kit (Illumina platforms, Kappa Biosystems, Boston, USA. Cat No. KK8504) according to the instructions of the manufacturer. DNA from each sample was randomly fragmented by nebulization to an average size of 400 bp, and processed by the Illumina DNA sample preparation protocol including end-repair, add-tailing, paired-end adaptor ligation and PCR. Paired-end sequencing libraries of each sample were built with an insert size of 400 bp and sequencing was performed on a HiSeq 2000 platform with a read length of 150 bp.

Among the identified single nucleotide polymorphism (SNP), a total of 188,323,429 (89.37%) were located in intergenic regions, followed by 20,506,072 (9.73%) in introns and 1,756,690 (0.834%) in CDS regions. For the SNP in CDS region, 995,686 (0.47%) were missense variants, and may have strong effect on the related genes function. Furthermore, 18,186 (54.20%) annotated genes were affected at least in one accession. In addition, 734,138 (0.35%) synonymous SNPs occurred in CDS regions (Supplementary Table 18).

#### 4.2 Admixture of tea population

To further illustrate the evolutionary history of the tea genome, a model-based clustering algorithm implemented in Admixture was used to estimate the relative genome composition for each accession. The clustering algorithm analysis indicated that the three populations fit the best model for all 139 accessions (Supplementary Fig. 13). When k was 3, the *C. sinensis* var. *sinensis* (CSS), *C. sinensis* var. *assamica*

(Masters) Chang (CSA), and *C. sinensis*-related species (CSR) could be distinguished; this was consistent with the principal component analysis (PCA) result (Fig. 2d). When k from 3 to 4, most of new accessions collected out of China arose from CSA and CSS (yellow color, marked with arrow), indicating their high diversity.

##### 4.3 Historically effective population size

Contigs longer than 100 kb were analyzed using Multiple Sequentially Markovian Coalescent (MSMC)[53] to infer the historically effective population size from multiple individuals of the same population. BEAGLE (version 4.1) was used to phase the genotype calls as the MSMC method was better suited for phased data. The phased VCF file was filtered with base quality over 20 and mapping quality over 30 and the depth was between 1/2 and twice of the mean depth. This mask file could be generated via maskBed.pipeline.sh in MSMC-tools. The mappability mask file was generated via the pipeline documented in maize. Four/Six samples from each group with high sequencing depth were selected and each sample was treated as a haploid. We also used MSMC for the timing and nature of population separations between two populations.

##### 4.4 High heterozygosity maintain tea plants adaptability

Tea has self-incompatibility and high heterozygosity. We wanted to find the regions which tend to maintain heterozygosity in tea, and what are the advantages of high heterozygosity. We examined the high heterozygosity and high deviation ratio region by calculating the heterozygosity and deviation ratio in sliding window 20 kb by steps 2 kb, respectively. The deviation ratio was calculated by (Ho-He)/He (Ho:average observed heterozygosity, He:average expected heterozygosity). We collected the intersection of top 1% of deviation ratio regions and top 1% of heterozygosity regions. We obtained 655 genes in the region. The genes that may keep tea heterozygosity were collected (Supplementary Table 25). The genes about disease resistance, growth and development, self-incompatibility, terpene synthase, and flavanone metabolism were in high heterozygosity and high deviation ratio region. These genes may be closely related to the adaptability of tea. We further examined the high heterozygosity points of CDS regions and found that half of genes had high heterozygosity points in these CDS regions, and about 2/3 of those points could result in nonsynonymous mutations.

##### 4.5 Gene flow of tea

To test whether admixture is confounding the phylogeny, the population allele frequency-based model Treemix was applied to account for variance arisen by secondary migration events. The groups were split by phylogenetic tree (Supplementary Table 26). When up to six migration events were included in the model, the major branching patterns in our tree remained largely unchanged. The results showed a great admixture among the populations (Supplementary Fig. 19). The results of some accessions in the phylogenetic tree were against the traditional classification. F4-test and F3-test of the accessions were examined by treemix (Supplementary Table 27). When the Z-score was greater than 3 or less than -3, the results of F4-test indicated there was gene flow between the samples[52]. The gene flow affected the position of HZ114, HZ104, HZ122, and HZ050 in the phylogenetic tree.

We also randomly generated 1000 groups for F4-test. Every group contain three randomly individuals and CM-1 (out group). Gene flow was detected in 979 groups. This result shown that the wide gene flow was among tea accessions. The detail results were supply in attachment (Supplementary Table 28).

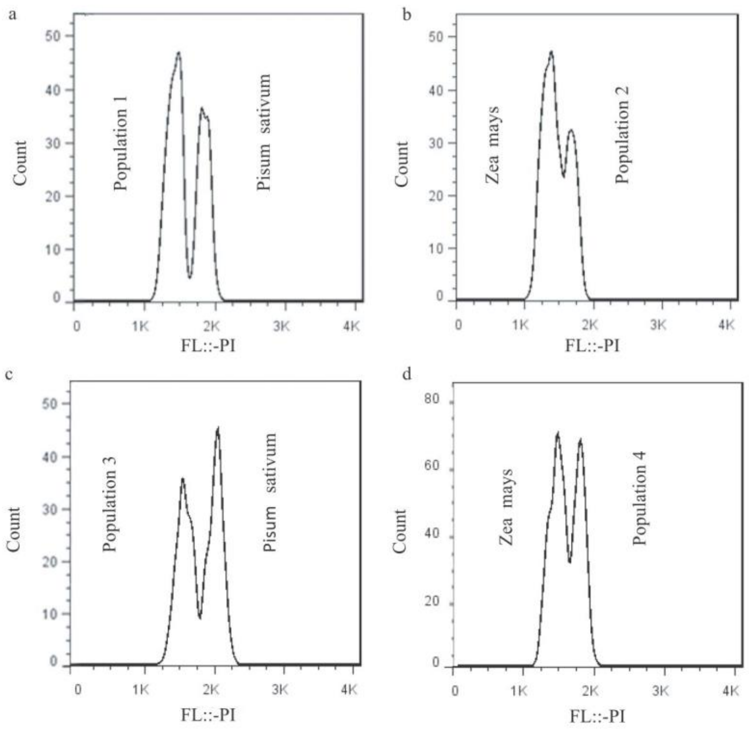

Supplementary Fig. 1: The evaluation of the genome size of LJ43 by flow cytometry. The four populations of LJ43 were used to determinate the genome size. Two populations were estimated with *Pisum sativum* as internal standard, and the other were estimated with *Zea mays* as internal standard. Population1, population3 and population4 contained 3 repeat leaf samples, population2 contained 4 repeat leaf samples. The mean of all was 3295.397087 Mb ± 179.5980113 Mb.

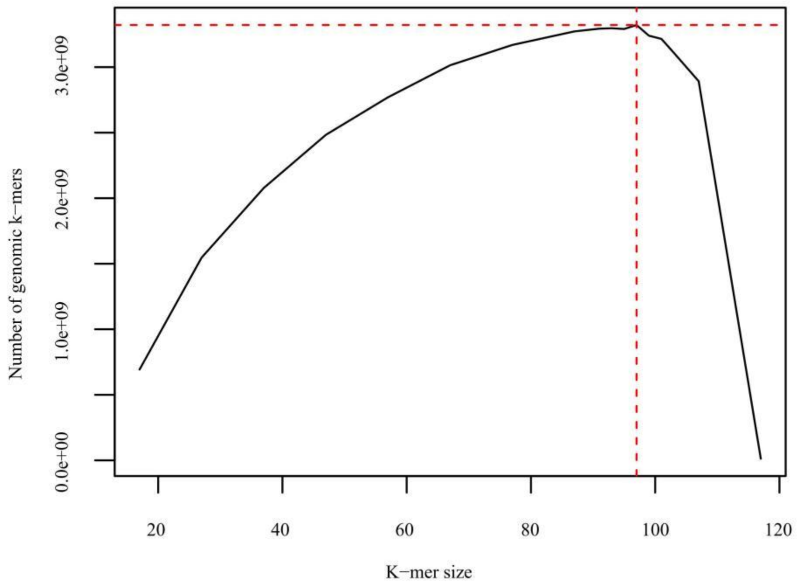

Supplementary Fig. 2: The genome size of LJ43 estimated by K-mer analysis. The best k was 97 by KmerGenie. The predicted genome size was about 3,321,109,494 bp.

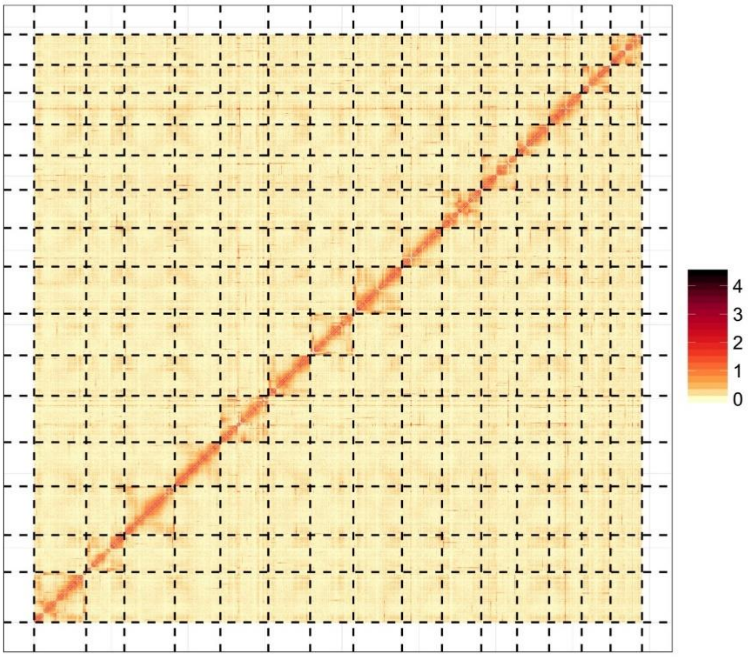

Supplementary Fig. 3: Genome-wide all-by-all Hi-C interaction of scaffold Hi-C genome. The fraction of contigs in orderings with high orientation quality: 12288 (59.31%) with length 2840497749 (92.13%). The scaffold genome has 205 collinear blocks containing 3,053 genes.

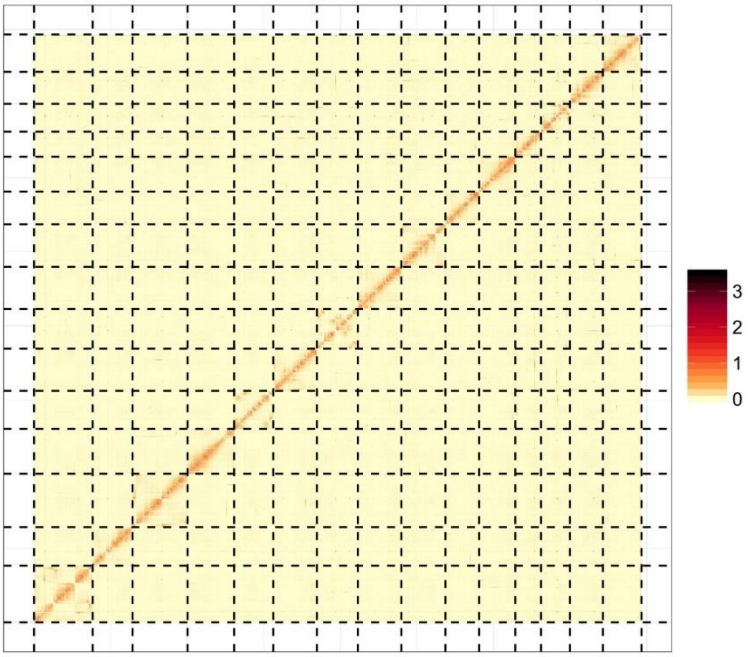

Supplementary Fig.4: Genome-wide all-by-all Hi-C interaction of contig Hi-C genome. The fraction of contigs in orderings with high orientation quality: 17715 (76.47%), with length 2801038355 (93.56%). The contig Hi-C genome comprises 208 collinear blocks containing 3,205 genes.

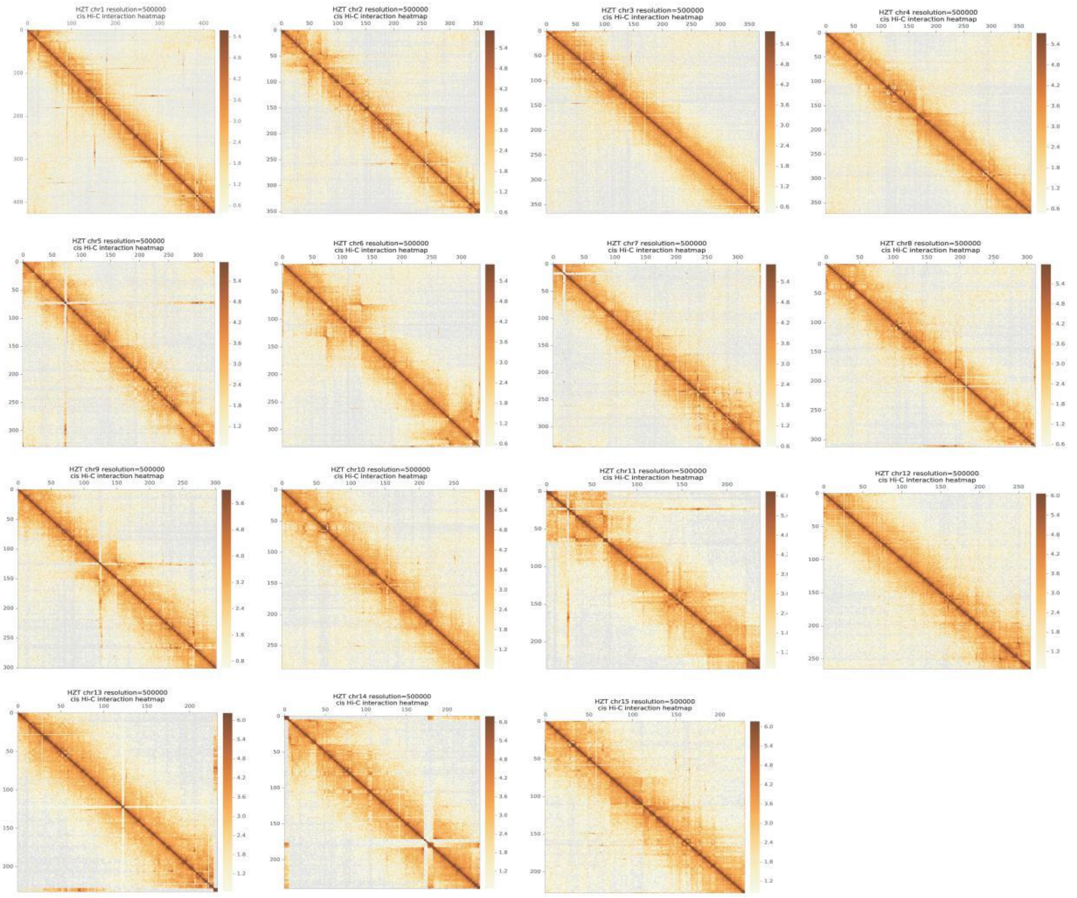

Supplementary Fig. 5: Hi-C interaction of Chromosome-inside. Resolution is 500kp.

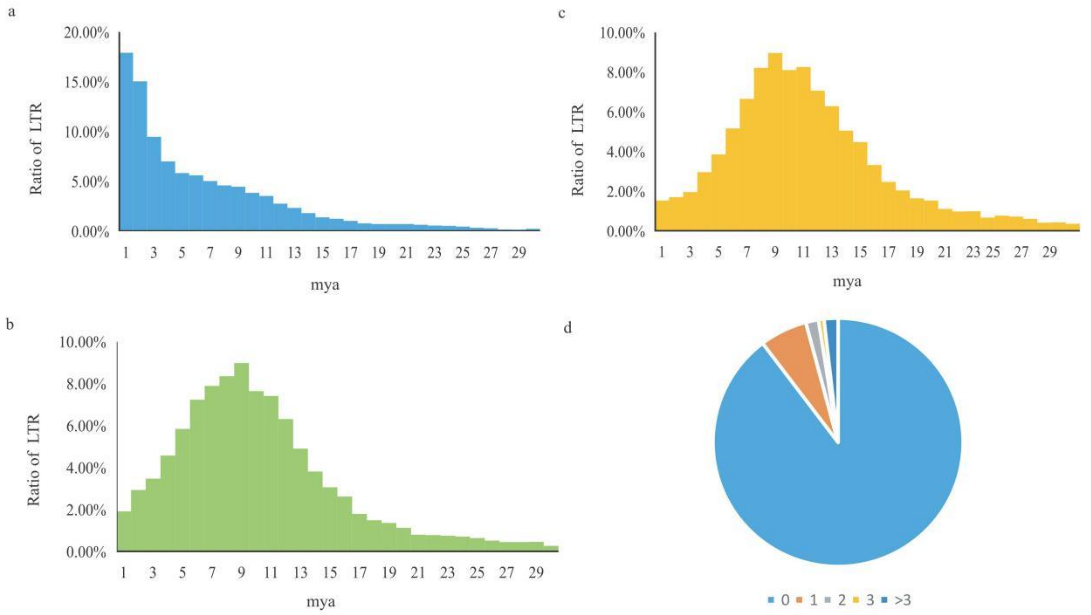

Supplementary Fig. 6: Insertion time of the annotated LTR in LJ43 (*Camellia sinensis* var. *sinensis*), Shuchazao (*Camellia sinensis* var. *sinensis*) and YK10 (*Camellia sinensis* var. *assamica*) genome and the number of corrected bases of LTR terminal sequence in LJ43. a, b, and c are the insertion time of the LTR in the genome of LJ43, SCZ, and YK10, respectively. The abscissa was million years (mya), the ordinate was percentage of LTR. d. The number of bases corrected by Illumina read in complete LTR terminal sequence. The number of corrected bases of most LTR terminal sequence were 0, and the number of corrected bases (≤3) was about 98.19%. The results show that the recent LTR of LJ43 were true and not introduced by error correction. The genome of LJ43 had more recent LTR, implying that its genome is more complete.

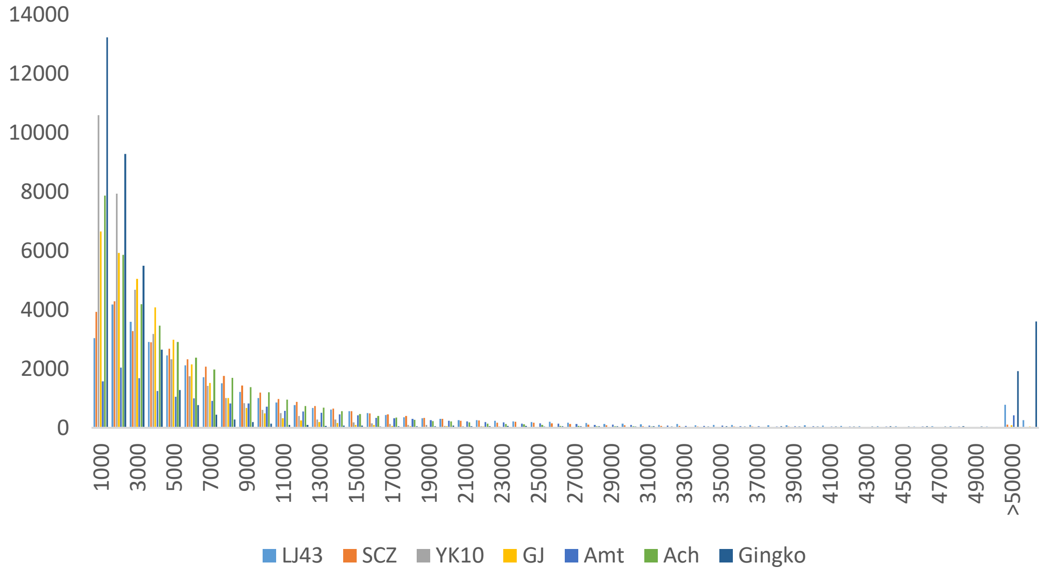

Supplementary Fig. 7: The gene length distribution of seven species: LJ43, SCZ, YK10, GJ, Amt, Ach, and Gingko were Longjing43, Shuchazao, Yunkang 10, citrus[33], *Amborella trichopoda*, *Actinidia Chinensis*, *and Gingko biloba*. X-axis was genes length, y-axis was genes number. LJ43 and SCZ had more genes with length 10kb-50kb, LJ43, Amt and ginkgo had more genes with length ≥ 50kb. The average gene length of LJ43, SCZ, YK10, GJ, Amt, Ach, and Gingko were 10,816bp, 7,386bp, 3,549bp, 4,061bp, 11,053bp, 5,388bp, and 25,619bp, respectively.

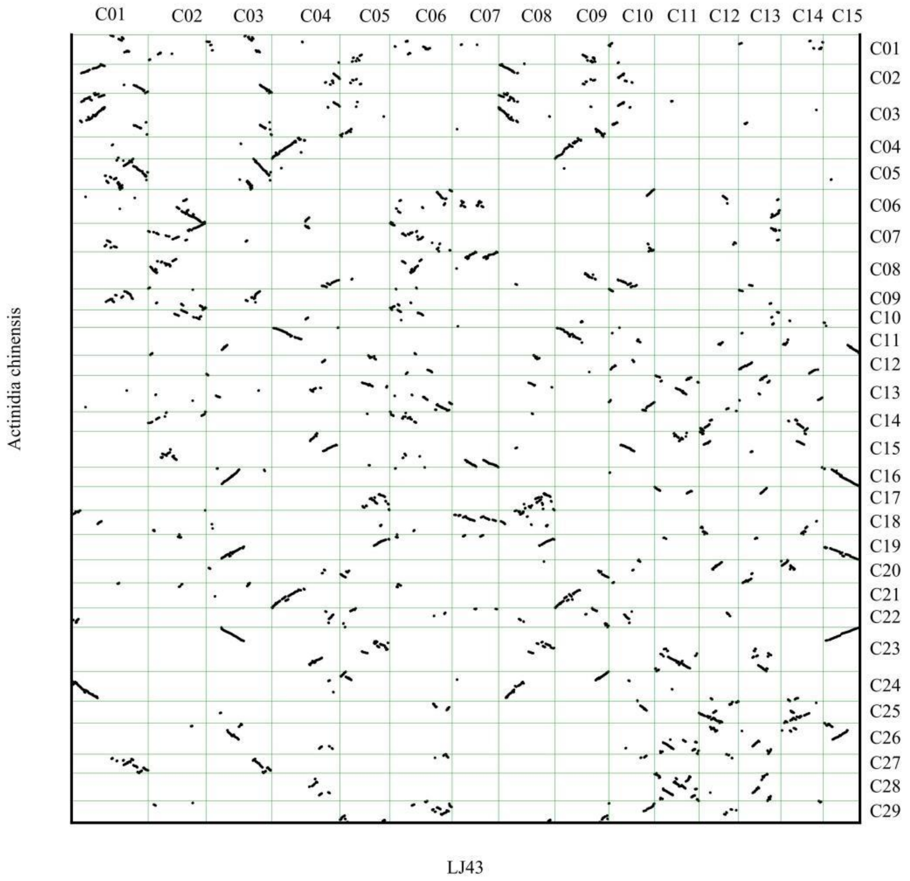

Supplementary Fig. 8: The collinearity of *Actinidia chinensis* and LJ43. The chromosome-level *Actinidia chinensis* genome assembly (y-axis) aligned to the chromosome-level of LJ43 genome assembly (x-axis). C was chromosome.

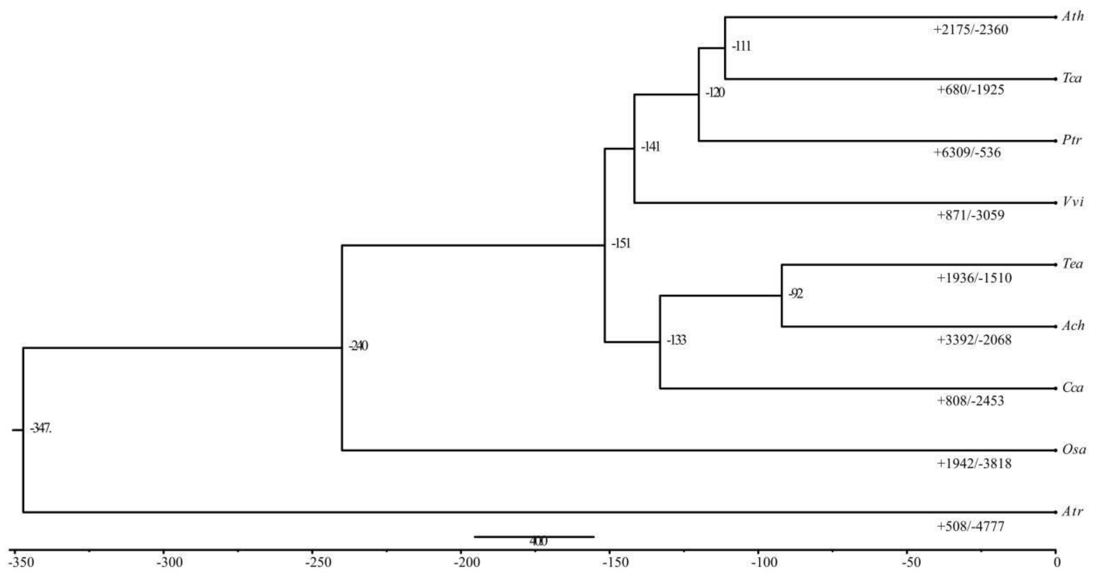

Supplementary Fig. 9: Expansion and contraction of gene families of LJ43 and 8 other plant species. The divergence time is shown beside each node, the unit is million years. Ath, Tca, Ptr, Vvi, Tea, Ach, Cca, Osa, Atr represented *Arabidopsis thaliana, Theobroma cacao, Populus trichocarpa, Vitis vinifera,* LJ43*, Actinidia Chinensis, Coffea, Oryza sativa subsp. Geng* and *Amborella trichopoda. Amborella trichopoda* was the outgroup. The "+" indicates expansion of gene families and the "-" indicates contraction of gene families.

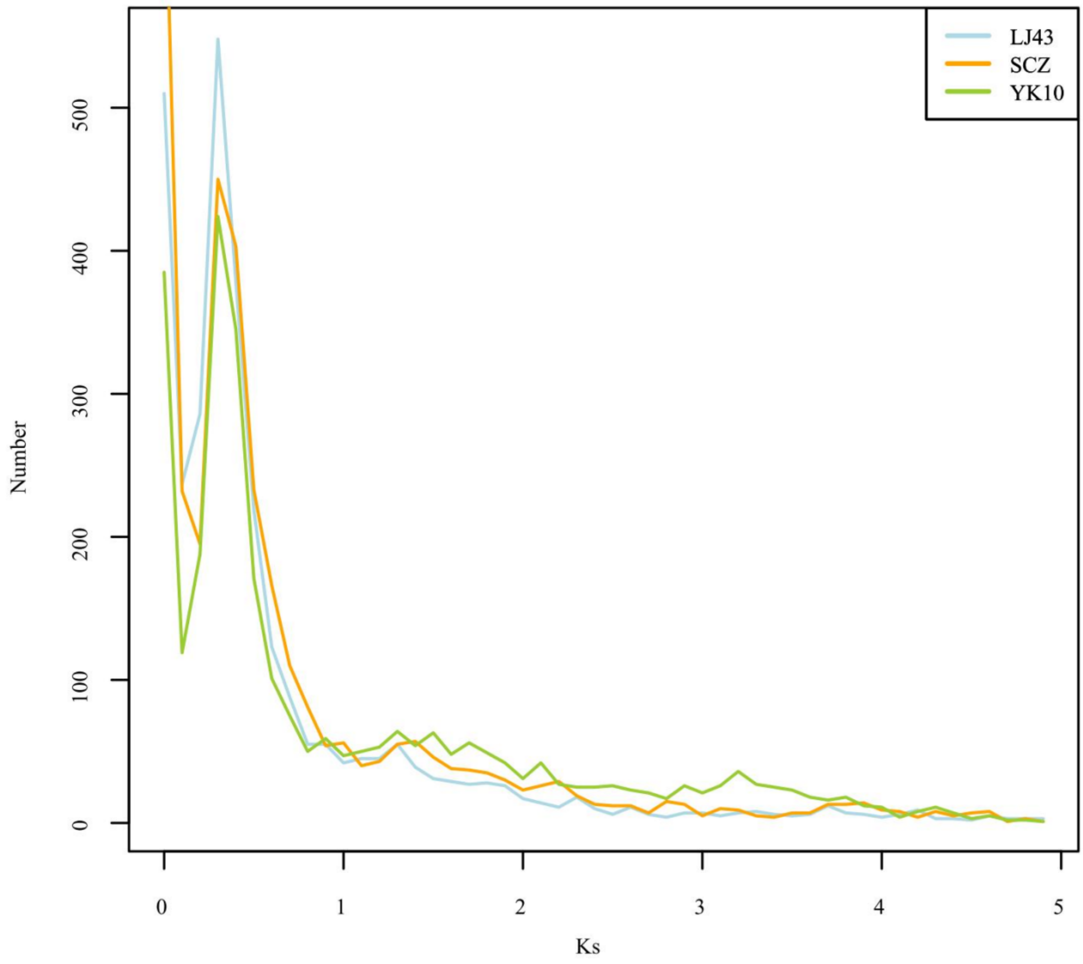

Supplementary Fig. 10: Whole genome duplication in LJ43, SCZ and YK10. The gene pairs were selected from 2-member gene groups. The x axis is Ks. The y axis is the number of gene clusters with this degree of divergence.

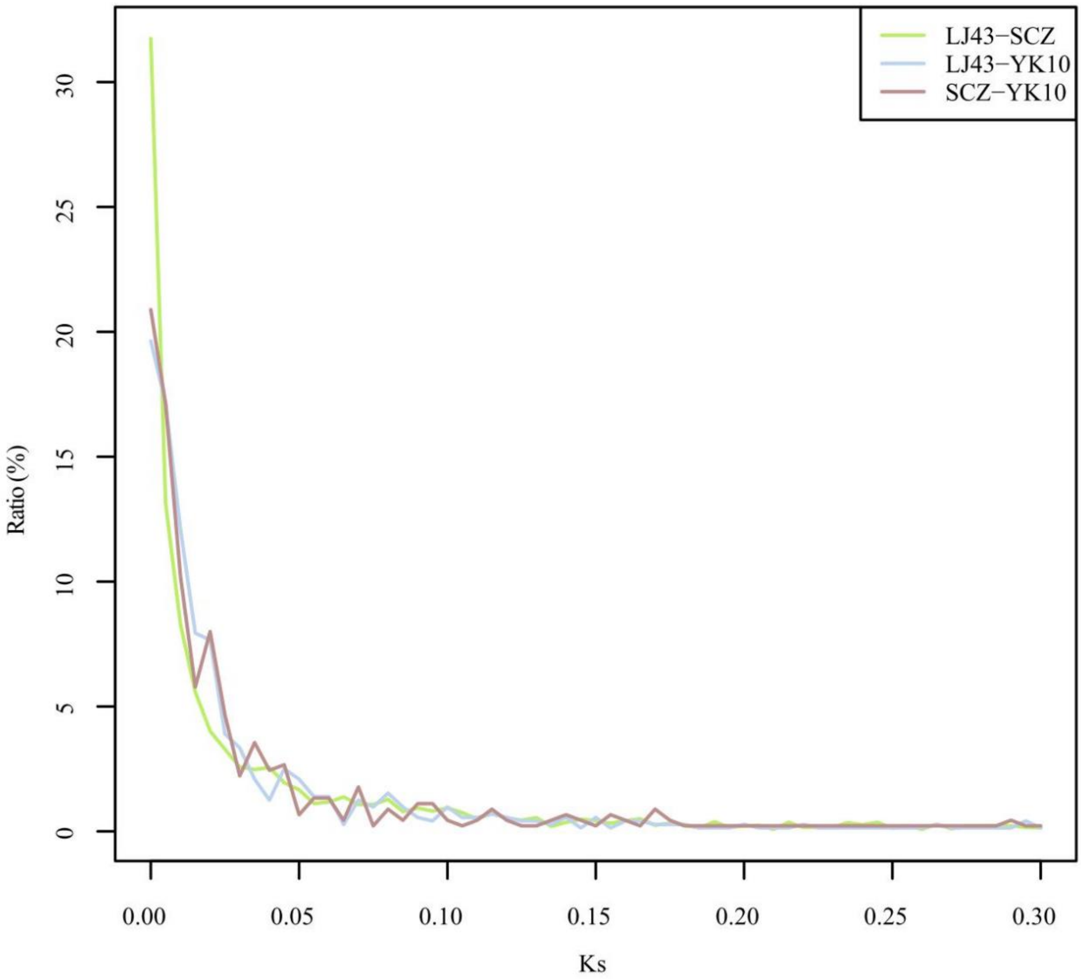

Supplementary Fig. 11: The diversity of LJ43, SCZ and YK10. The x axis is the Ks of collinearity genes of two tea genomes. The y axis is the number of gene pairs with this degree of divergence.

Supplementary Fig, 12: The phylogenetic tree of all genome region with bootstrap. We constructed the phylogenetic tree by all genome SNPs.

The figure is supplied in supplementary figure 12.pdf.

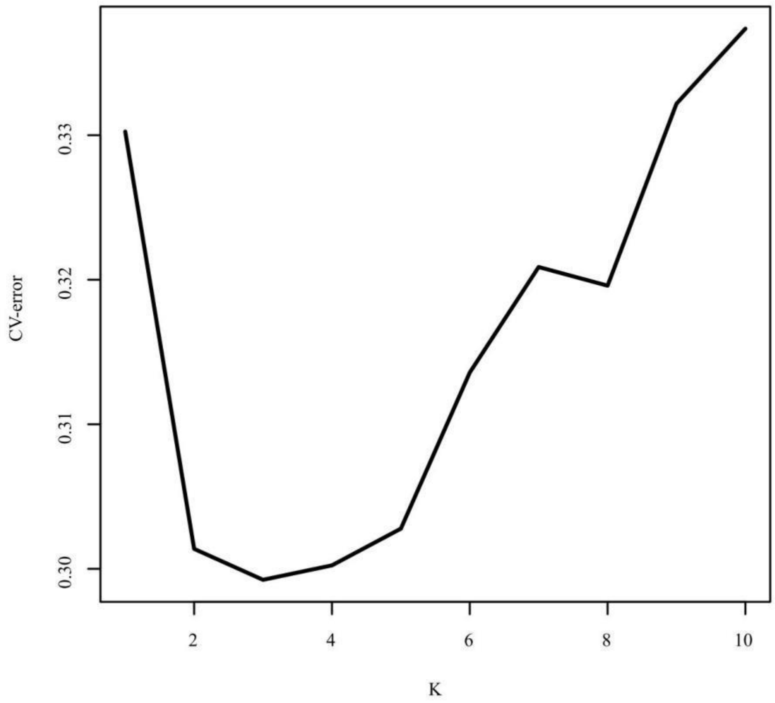

Supplementary Fig. 13: The CV-error of tea population. The CV-error was calculated by Admixture, k=3 had the lowest CV-error.

Supplementary Fig. 14: The phylogenetic tree of inter gene region with bootstrap. We consider the selection status of SNPs, and the phylogenetic tree was constructed by inter gene region SNPs.

The figure is supplied in supplementary figure 14.pdf.

Supplementary Fig. 15: The phylogenetic tree of 4DTV. We constructed the phylogenetic tree by 4DTV sites.

The figure is supplied in supplementary figure 15.pdf.

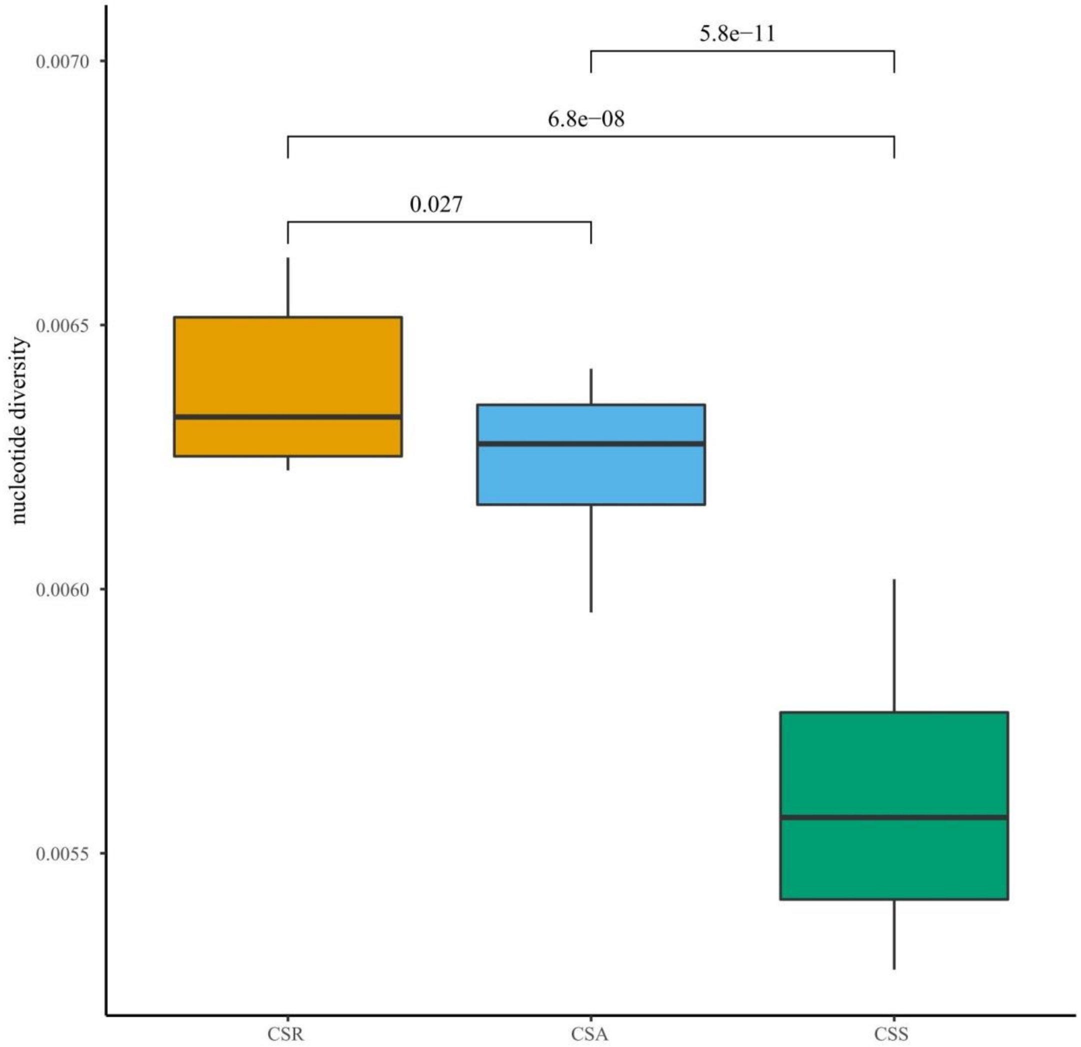

Supplementary Fig.16: The nucleotide diversity of CSA, CSS, and CSR. The boxes from left to right are CSR, CSA, and CSS, respectively. The P-values of all were less than 0.05.

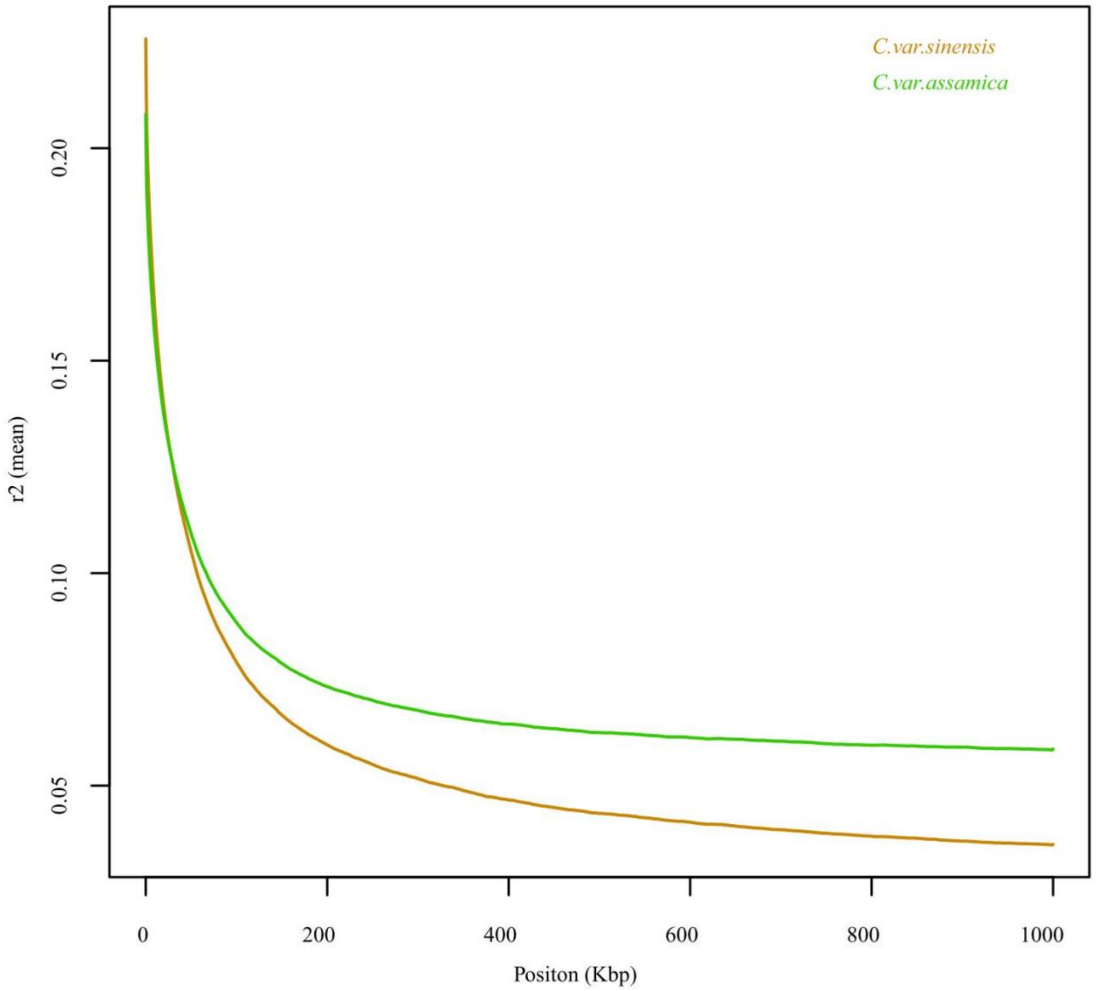

Supplementary Fig. 17: The LD of tea population. Squared correlation coefficient (r2) of linkage disequilibrium (LD) between the SNPs called in CSS and CSA populations. The average $r^2$ among SNPs decayed to approximately 50% of its maximum value at approximately 41 kb and 59 kb, respectively. These values thus indicate that the tea genomes have relatively long LD distances and slow LD decays.

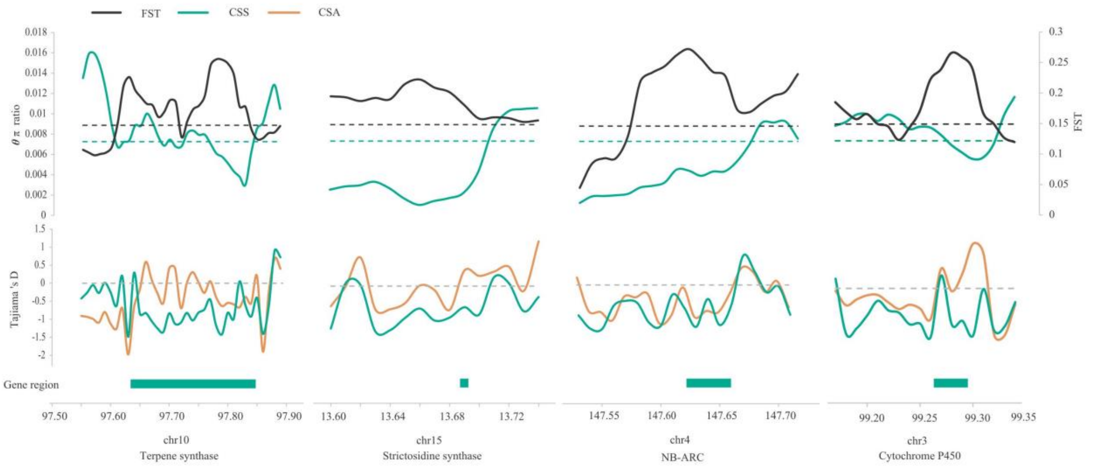

Supplementary Fig. 18: The Fst, θπ ratio and Tajiama 's D of some selection genes. The region of selective genes had high *Fst* value, low $\theta_\pi$ value and below zero Tajima's D.

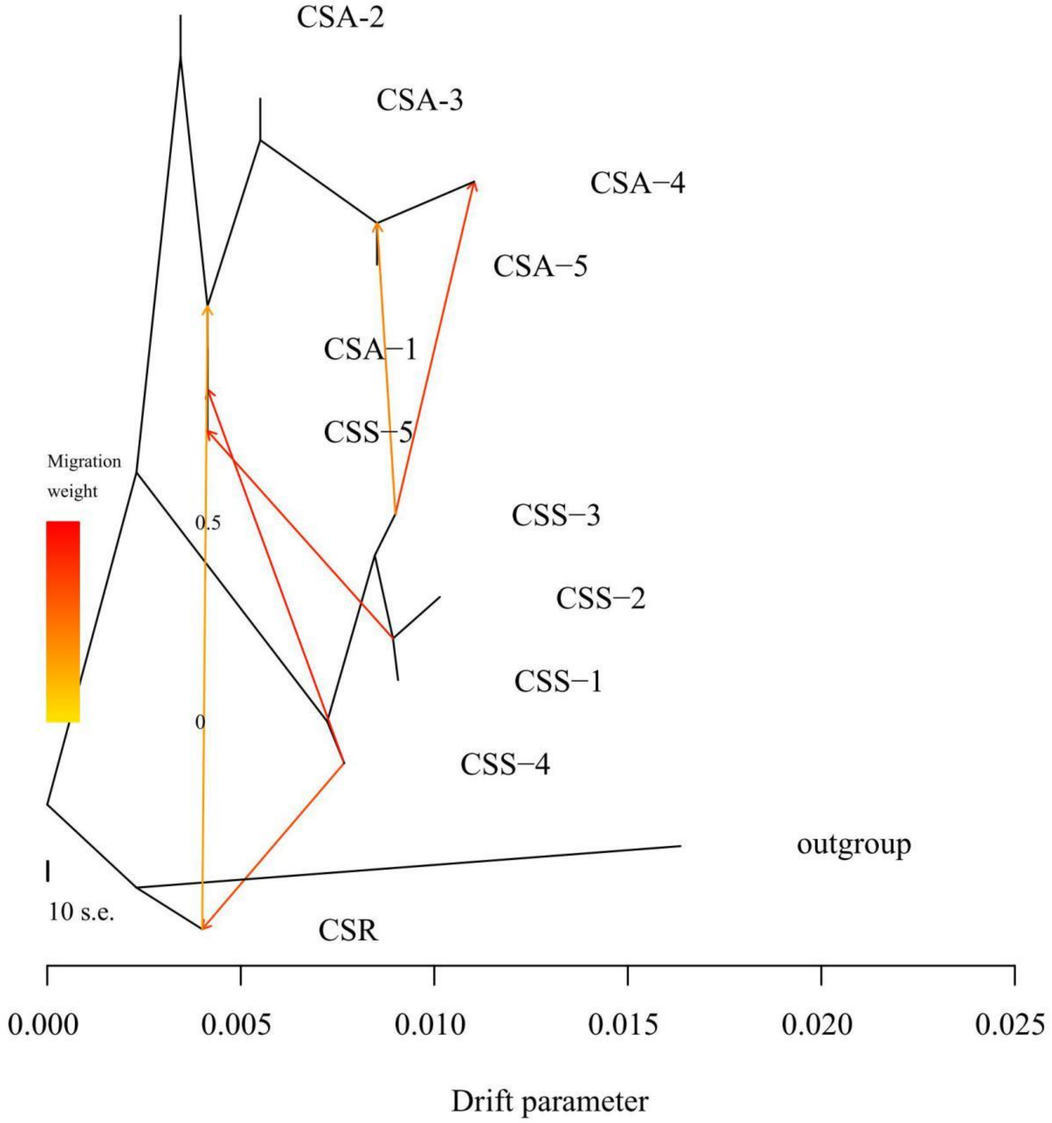

Supplementary Fig. 19: The gene flow in tea population. The direction of arrow represents the direction of gene flow. The information of groups is supplied in supplementary Table 27.

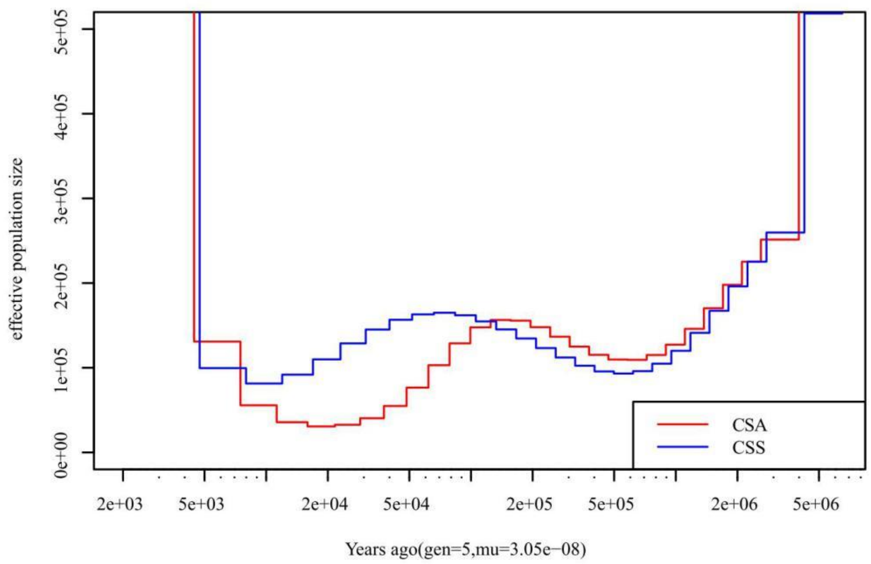

Supplementary Fig. 20: The history population of tea population. The generation was 5 years.

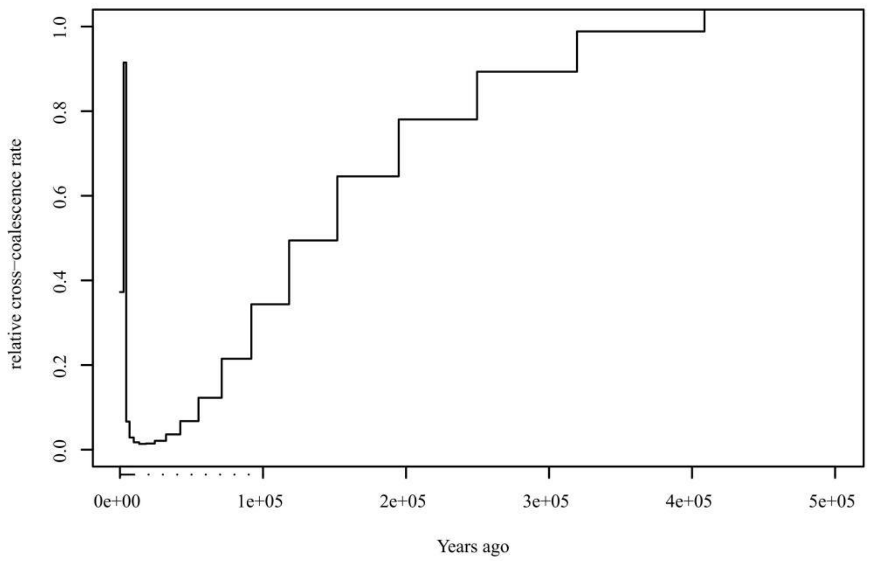

Supplementary Fig. 21: The history population of CSA and CSS.

Supplementary Table 1: Statistics of sequence data.

| Library | Clean Data |
|---|---|
| PE | ~214 Gb |
| PacBio | ~196 Gb |
| RNA-seq | ~340 Gb |
| Hic(PE150) | ~263 Gb |
| 10X(PE150) | ~247 Gb |
| BIONANO | ~445 Gb |

Supplementary Table 2: State of tea genome

| Items | Contig_len (bp) | Scaffold_len (bp) | Contig_num | Scaffold_num |
|---|---|---|---|---|
| Total | 3,259,965,435 | 3,260,671,035 | 37,600 | 30,544 |
| Max_length | 2,426,329 | 212,836,541 | - | - |
| Number>=2000bp | - | - | 37,600 | 30,544 |
| N50 | 271,332 | 143,847,529 | 3,214 | 10 |
| N60 | 195,993 | 120,058,857 | 4,625 | 12 |
| N70 | 134,207 | 114,499,148 | 6,629 | 15 |
| N80 | 80,686 | 81,145 | 9,749 | 2,695 |
| N90 | 35,684 | 35,688 | 15,729 | 8,671 |

Note: “-” indicates missing data.

Supplementary Table 3: State of chromosome length and coefficient of determination.

| Pseudomolecule | Scaffold Number | Length (bp) | Coefficient of determination ($R^2$) |
|---|---|---|---|
| chr1 | 593 | 212,836,541 | 0.96 |
| chr2 | 614 | 176,915,850 | 0.96 |
| chr3 | 568 | 183,890,167 | 0.94 |
| chr4 | 545 | 186,373,698 | 0.90 |
| chr5 | 529 | 164,367,897 | 0.89 |
| chr6 | 498 | 165,692,707 | 0.97 |
| chr7 | 517 | 168,674,417 | 0.88 |
| chr8 | 476 | 156,400,541 | 0.98 |
| chr9 | 456 | 150,683,510 | 0.98 |
| chr10 | 453 | 143,847,529 | 0.91 |
| chr11 | 401 | 118,392,857 | 0.92 |
| chr12 | 401 | 132,670,986 | 0.85 |
| chr13 | 348 | 116,245,087 | 0.96 |
| chr14 | 366 | 120,058,857 | 0.98 |
| chr15 | 306 | 114,499,148 | 0.84 |
| Total anchored | 7,071 | 2,311,549,792 | - |
| Unanchored | 30,529 | 949,121,243 | - |

Note: “-” indicates missing data.

Supplementary Table 4: The variation and INDEL of LJ43.

| Type | Number | Ratio |
|---|---|---|
| variation (heterozygous) | 19,753,302 | 0.606% |
| variations (homogeneous) | 7,307 | 0.000224% |
| INDEL (heterozygous) | 2,264,855 | 0.0695% |
| INDEL (homogeneous) | 18,525 | 0.000568% |

Supplementary Table 5: Repetitive sequence annotation of RepeatMasker.

| Elements | Number | Length (bp) |
|---|---|---|
| DNA_elements | 342,783 | 166,374,592 |
| ERV_classI | 5,144 | 3,432,108 |
| ERV_classII | 474 | 344,906 |
| hAT-Charlie | 340 | 97,911 |
| L3/CR1 | 5,031 | 1,160,068 |
| LINE1 | 73,377 | 57,624,329 |
| LINE2 | 9,844 | 3,445,887 |
| LINEs | 121,399 | 76,115,524 |
| Low_complexity | 123,740 | 6,408,906 |
| LTR_elements | 767,684 | 1,417,058,775 |
| Satellites | 11,812 | 32,810,223 |
| Simple_repeats | 869,486 | 105,405,729 |
| SINEs | 16,385 | 3,174,787 |
| Small_RNA | 17,992 | 5,066,448 |
| Total_interspersed_repeats | | 2,448,318,482 |
| bases_masked | | 2,381,432,965 |

Supplementary Table 6: The TE of LJ43.

| | Repbase TEs | | TE proteins | | *De novo* | | Combined TEs | |
|---|---|---|---|---|---|---|---|---|
| Type | Length (Bp) | % of genome | Length (Bp) | % in genome | Length (Bp) | % in genome | Length (Bp) | % in genome |
| DNA TE | 20,381,582 | 0.63 | 13,417,865 | 0.41 | 197,436,270 | 6.06 | 209,854,625 | 6.44 |
| LINE | 634,240 | 0.02 | 21,127,314 | 0.65 | 94,769,667 | 2.91 | 101,288,844 | 3.11 |
| SINE | 115,772 | 0.00 | 0 | 0.00 | 4,014,840 | 0.12 | 4,057,783 | 0.12 |
| LTR-retro | 366,724,615 | 11.25 | 464,352,782 | 14.24 | 1,964,618,648 | 60.27 | 1,981,304,145 | 60.77 |
| Total | 387,854,727 | 11.90 | 497,370,701 | 15.26 | 2,260,839,425 | 69.36 | 2,302,505,397 | 70.44 |

Supplementary Table 7: Transcriptomes sequence of LJ43.

| Name | Tissue type | Clean Reads reapeat1 | Clean data (bp) repeat1 | Clean Reads reapeat2 | Clean data (bp) repeat2 | Clean Reads reapeat3 | Clean data (bp) repeat3 |
|---|---|---|---|---|---|---|---|
| leaf summer | mature leaves | 204,015,168 | 7,509,164,019 | 185,432,168 | 6,848,866,161 | 196,054,632 | 7,225,952,895 |
| leaf autumn | mature leaves | 134,536,224 | 4,960,492,402 | 148,311,944 | 5,345,086,650 | 199,661,960 | 7,312,697,492 |
| leaf winter | mature leaves | 133,755,424 | 4,816,136,977 | 135,592,000 | 4,890,266,334 | 142,236,600 | 5,124,935,707 |
| leaf spring | mature leaves | 172,916,408 | 6,344,736,952 | 212,148,920 | 7,786,169,626 | 215,747,576 | 7,929,395,621 |
| root autumn | roots | 164,994,272 | 5,980,998,613 | 143,619,888 | 5,211,047,770 | 200,609,576 | 7,321,219,607 |
| root summer | roots | 150,369,688 | 5,490,083,176 | 126,524,824 | 4,620,519,432 | 159,517,416 | 5,818,447,162 |
| root winter | roots | 118,735,432 | 4,273,133,810 | 146,046,888 | 5,255,500,404 | 132,702,536 | 4,782,931,542 |
| root spring | roots | 155,342,192 | 5,660,781,661 | 129,662,080 | 4,757,721,164 | 130,815,552 | 4,806,046,193 |
| flower winter | flowers | 154,929,968 | 5,579,880,513 | 146,720,784 | 5,289,249,128 | 125,384,736 | 4,524,026,055 |
| flower autumn | flowers | 115,593,368 | 4,152,920,821 | 154,444,920 | 5,547,858,358 | 191,818,400 | 7,003,668,404 |
| flower(young fruit) spring | young fruit | 217,866,304 | 8,002,440,801 | 159,085,104 | 5,844,261,629 | 185,071,736 | 6,776,869,697 |
| stem autumn | young stems | 136,883,416 | 4,929,267,997 | 128,878,040 | 4,636,834,266 | 198,809,720 | 7,251,424,376 |
| stem summer | young stems | 192,130,136 | 7,065,984,704 | 184,804,800 | 6,796,434,130 | 195,752,824 | 7,197,948,142 |
| stem winter | young stems | 127,212,360 | 4,586,059,072 | 161,788,424 | 5,821,583,681 | 152,565,648 | 5,620,530,317 |
| stem spring | young stems | 132,921,200 | 4,880,375,054 | 111,220,840 | 4,086,690,304 | 127,033,040 | 4,664,850,247 |
| bud autumn | axillary buds | 192,373,512 | 7,093,878,955 | 197,798,048 | 7,283,645,328 | 175,308,432 | 6,451,941,331 |
| bud winter | axillary buds | 147,912,040 | 5,324,502,387 | 115,592,528 | 4,167,683,576 | 134,121,152 | 4,826,824,784 |
| bud summer | axillary buds | 234,540,448 | 8,622,286,106 | 210,273,200 | 7,733,356,033 | 185,376,328 | 6,842,916,444 |
| bud spring | one bud and two leaves | 177,138,808 | 6,495,895,305 | 143,576,968 | 5,184,808,153 | 263,048,184 | 9,662,721,977 |

Supplementary Table 8: The genes function annotation of LJ43.

| Database | Number (ratio) |
|---|---|
| Swissprot | 26,373 (78.59%) |
| InterPro | 30,655 (91.35%) |
| KEGG | 8,643 (25.76%) |
| GO | 20,035 (59.71%) |
| Combined total annotated | 31,437 (93.69%) |
| Unannotated | 3,502 (6.31%) |

Supplementary Table 9: GO enrichment of expansion genes.

| GO | All genes | Expansion genes | P-value | FDR | Function |
|---|---|---|---|---|---|
| GO:0006376 | 4 | 4 | 2.83E-03 | 4.41E-02 | Luc7-related |
| GO:0006421 | 4 | 4 | 2.83E-03 | 4.41E-02 | AsparaginE-tRNA ligase |
| GO:0008146 | 39 | 25 | 5.33E-08 | 2.18E-06 | Sulfotransferase domain |
| GO:0008234 | 66 | 31 | 1.71E-05 | 5.09E-04 | Ulp1 protease family, C-terminal catalytic domain |
| GO:0009607 | 28 | 15 | 4.49E-04 | 8.64E-03 | Bet v I/Major latex protein |
| GO:0016758 | 311 | 176 | 2.20E-16 | 2.40E-14 | UDP-glucuronosyl/UDP-glucosyltransferase |
| GO:0016891 | 7 | 6 | 8.46E-04 | 1.54E-02 | Dicer dimerisation domain |
| GO:0016998 | 25 | 14 | 0.38E-03 | 7.77E-03 | Glycoside hydrolase, family 19, catalytic |
| GO:0030246 | 177 | 67 | 6.80E-06 | 2.47E-04 | Galactose mutarotase-like domain |
| GO:0030247 | 73 | 42 | 2.44E-10 | 1.33E-08 | Wall-associated receptor kinase, galacturonan-binding domain |
| GO:0031683 | 8 | 7 | 2.22E-04 | 4.84E-03 | Guanine nucleotide binding protein (G-protein), alpha subunit |
| GO:0042545 | 61 | 27 | 2.05E-04 | 4.78E-03 | Pectinesterase, catalytic |
| GO:0043531 | 436 | 268 | 2.20E-16 | 2.40E-14 | NB-ARC |
| GO:0045735 | 60 | 29 | 1.60E-05 | 5.09E-04 | Cupin 1 |
| GO:0046488 | 19 | 13 | 3.35E-05 | 9.14E-04 | Phosphatidylinositol-4-phosphate 5-kinase, core |
| GO:0048268 | 15 | 11 | 5.20E-05 | 1.31E-03 | Phosphoinositide-binding clathrin adaptor, domain 2 |
| GO:0048544 | 152 | 123 | 2.20E-16 | 2.40E-14 | S-locus glycoprotein domain |
| GO:0051740 | 4 | 4 | 0.28E-02 | 0.44E-01 | Ethylene receptor |
| GO:0055085 | 770 | 245 | 1.18E-08 | 5.52E-07 | ABC transporter type 1, transmembrane domain |
| GO:0070588 | 23 | 23 | 2.19E-15 | 1.79E-13 | P-type ATPase, subfamily IIB |
| GO:0071805 | 34 | 26 | 6.55E-11 | 4.28E-09 | Potassium transporter |

Supplementary Table 10: KEGG enrichment of expansion genes.

| KEGG | All genes | Expansion genes | P-value | FDR | Function |
|---|---|---|---|---|---|
| K01183 | 34 | 18 | 1.49E-04 | 0.71E-02 | chitinase |
| K05391 | 14 | 11 | 1.75E-05 | 0.17E-02 | CNGC; cyclic nucleotide gated channel, plant |
| K06617 | 7 | 7 | 3.47E-05 | 0.22E-02 | raffinose synthase [EC:2.4.1.82] |
| K07437 | 7 | 7 | 3.47E-05 | 0.22E-02 | CYP26A; cytochrome P450 family 26 subfamily A |
| K08237 | 9 | 9 | 1.85E-06 | 0.26E-03 | Glycosyltransferases Metabolism |
| K11835 | 8 | 7 | 0.22E-03 | 0.92E-02 | Ubiquitin system Genetic Information Processing |
| K11844 | 7 | 6 | 0.85E-03 | 0.03 | Ubiquitin system Genetic Information Processing |
| K13260 | 6 | 6 | 0.15E-03 | 0.71E-02 | CYP81E1_7; isoflavone/4'-methoxyisoflavone 2'-hydroxylase |
| K13457 | 56 | 43 | 2.20E-16 | 1.25E-13 | disease resistance protein RPM1 |
| K13459 | 16 | 14 | 8.88E-08 | 2.51E-05 | disease resistance protein RPS2 |
| K13691 | 11 | 9 | 6.38E-05 | 0.36E-02 | Glycosyltransferases Metabolism |
| K15095 | 10 | 8 | 2.28E-04 | 0.92E-02 | (+)-neomenthol dehydrogenase |
| K15639 | 10 | 10 | 4.26E-07 | 8.03E-05 | CYP734A1, BAS1; PHYB activation tagged suppressor 1 [EC:1.14.-.-] |
| K15803 | 8 | 8 | 8.01E-06 | 0.91E-03 | GERD; (-)-germacrene D synthase |
| K18819 | 7 | 7 | 3.47E-05 | 0.22E-02 | GOLS; inositol 3-alpha-galactosyltransferase [EC:2.4.1.123] |

Supplementary Table 11: (InterPro) IPR enrichment of expansion genes.
The table is supplied in supplementary table 11.excel.

Supplementary Table 12: The IPR enrichment of special groups of LJ43.

| IPR | All genes | Special genes | P-value | FDR | Function |
|---|---|---|---|---|---|
| IPR000163 | 13 | 5 | 0.19E-02 | 0.49E-01 | Prohibitin |
| IPR002182 | 436 | 52 | 0.81E-03 | 0.32E-01 | NB-ARC |
| IPR004045 | 86 | 17 | 0.22E-03 | 0.13E-01 | Glutathione S-transferase, N-terminal |
| IPR008218 | 7 | 5 | 4.60E-05 | 0.36E-02 | ATPase, V1 complex, subunit F |
| IPR009600 | 4 | 3 | 0.16E-02 | 0.48E-01 | GPI transamidase subunit PIG-U |
| IPR016088 | 7 | 4 | 0.96E-03 | 0.32E-01 | Chalcone isomerase, 3-layer sandwich |
| IPR016363 | 4 | 4 | 3.30E-05 | 0.36E-02 | Legume lectin |
| IPR017989 | 18 | 17 | 2.20E-16 | 5.17E-14 | Ribosome-inactivating protein type 1/2 |
| IPR021113 | 10 | 5 | 0.45E-03 | 0.21E-01 | Acyl-ACP-thioesterase, N-terminal |

Supplementary Table 13: The GO enrichment of special groups of LJ43.

| GO | All genes | Special genes | P-value | FDR | Function |
|---|---|---|---|---|---|
| GO:0030598 | 100 | 27 | 4.03E-09 | 5.24E-07 | Ribosome-inactivating protein |
| GO:0043531 | 436 | 52 | 0.81E-03 | 0.31E-01 | NB-ARC |
| GO:0045430 | 7 | 4 | 0.96E-03 | 0.31E-01 | Chalcone isomerase, 3-layer sandwich |
| GO:0046961 | 10 | 5 | 0.45E-03 | 0.30E-01 | ATPase, V1 complex, subunit H |

Supplementary Table 14: The KEGG enrichment of special groups of LJ43.

| KEGG | All genes | Special genes | P-value | FDR | Function |
|---|---|---|---|---|---|
| K01859 | 7 | 4 | 0.96E-03 | 0.04 | chalcone isomerase |
| K14153 | 8 | 5 | 0.12E-03 | 0.98E-02 | hydroxymethylpyrimidine kinase<br>phosphomethylpyrimidine kinase<br>thiamine-phosphate diphosphorylase |
| K14305 | 16 | 9 | 5.71E-07 | 9.77E-05 | nuclear pore complex protein Nup43 |
| K14595 | 6 | 4 | 0.44E-03 | 0.02 | abscisate beta-glucosyltransferase |

Supplementary Table 15: The positive Darwinian selection genes of LJ43.
The table is supplied in supplementary table 15.excel.

Supplementary Table 16: The information of tea population.
The table is supplied in supplementary table 16.excel.

Supplementary Table 17: The SNP Results.

| | Number | ts/tv | ts/tv(1st ALT) |
|---|---|---|---|
| INDEL (only Hard-filter) | 34,850,045 | - | - |
| SNP (only Hard-filter) | 387,042,351 | 2.27 | 2.66 |
| SNP with multialleles | 52,270,113 (13.51%) | 0.96 | 1.41 |
| SNP with indel | 18,118,594 (4.68%) | 2.07 | 2.51 |
| SNP (gap 5 bp,qual>=40,maf>=0.01,dp(2.5%~97.5%),biallele SNP) | 218,870,098 (56.55%) | 3.56 | 3.56 |

Note: "-" indicates missing data.

Supplementary Table 18: Counts of the variant types of all tea samples.

| Type (alphabetical order) | Count | Ratio |
|---|---|---|
| initiator_codon_variant | 297 | 0% |
| intergenic_region | 188,323,429 | 89.37% |
| intron_variant | 20,506,072 | 9.73% |
| missense_variant | 995,686 | 0.47% |
| non_canonical_start_codon | 4 | 0% |
| splice_acceptor_variant | 7,777 | 0.00% |
| splice_donor_variant | 6,369 | 0.00% |
| splice_region_variant | 117,681 | 0.06% |
| start_lost | 2,517 | 0.00% |
| stop_gained | 33,580 | 0.02% |
| stop_lost | 3,155 | 0.00% |
| stop_retained_variant | 1,722 | 0.00% |
| synonymous_variant | 734,138 | 0.35% |
| exon | 1,756,690 | 0.834% |
| intergenic | 188,323,429 | 89.422% |
| intron | 20,410,690 | 9.692% |
| splice_site_acceptor | 7,777 | 0.004% |
| splice_site_donor | 6,369 | 0.003% |
| splice_site_region | 95,641 | 0.045% |

Supplementary Table 19: The selection genes in CSS.
The table is supplied in supplementary table 19.excel.

Supplementary Table 20: The selection genes in CSA.
The table is supplied in supplementary table 20.excel.

Supplementary Table 21: The GO enrichment of sweepfinder results of CSA.

| GO | Number | P-value | FDR | Function |
|---|---|---|---|---|
| GO:0000027 | 2 | 1.58E-03 | 3.88E-02 | Midasin |
| GO:0000166 | 40 | 2.40E-04 | 1.18E-02 | Nucleotide-binding alpha-beta plait domain |
| GO:0003676 | 63 | 2.60E-04 | 1.18E-02 | Ribonuclease H-like domain |
| GO:0004379 | 2 | 1.58E-03 | 3.88E-02 | Myristoyl-CoA:protein N-myristoyltransferase, N-terminal |
| GO:0004672 | 96 | 3.91E-05 | 3.93E-03 | Protein kinase domain |
| GO:0005488 | 38 | 1.63E-05 | 2.40E-03 | Armadillo-type fold |
| GO:0005515 | 222 | 2.54E-07 | 7.47E-05 | Ankyrin repeat |
| GO:0005524 | 178 | 1.02E-11 | 6.03E-09 | Protein kinase domain |
| GO:0006468 | 96 | 4.01E-05 | 3.93E-03 | Protein kinase domain |
| GO:0008270 | 72 | 2.49E-04 | 1.18E-02 | Transcription factor TFIIB |
| GO:0009234 | 2 | 1.58E-03 | 3.88E-02 | Menaquinone biosynthesis protein MenD |
| GO:0016020 | 67 | 1.89E-04 | 1.11E-02 | Pyrophosphate-energised proton pump |
| GO:0016021 | 62 | 3.05E-04 | 1.28E-02 | Cornichon |
| GO:0016301 | 14 | 5.58E-07 | 1.10E-04 | Diacylglycerol kinase, catalytic domain |
| GO:0016491 | 55 | 1.60E-04 | 1.05E-02 | Polyketide synthase, enoylreductase domain |
| GO:0016638 | 3 | 1.15E-03 | 3.88E-02 | Pyridoxine 5'-phosphate oxidase, dimerisation, C-terminal |
| GO:0016887 | 18 | 3.46E-04 | 1.36E-02 | ABC transporter-like |
| GO:0031491 | 2 | 1.58E-03 | 3.88E-02 | ISWI, HAND domain |
| GO:0042626 | 10 | 8.43E-04 | 3.10E-02 | ABC transporter type 1, transmembrane domain |
| GO:0043044 | 2 | 1.58E-03 | 3.88E-02 | ISWI, HAND domain |
| GO:0048278 | 2 | 1.58E-03 | 3.88E-02 | Exocyst complex component Sec10-like |
| GO:0050242 | 2 | 1.58E-03 | 3.88E-02 | Pyruvate, phosphate dikinase |
| GO:0055085 | 44 | 7.86E-05 | 5.78E-03 | Sugar transporter, conserved site |
| GO:0055114 | 95 | 6.76E-05 | 5.69E-03 | Alcohol dehydrogenase, C-terminal |

Supplementary Table 22: The GO enrichment of sweepfinder results of CSS.

| GO | Number | P-value | FDR | Function |
|---|---|---|---|---|
| GO:0000166 | 31 | 1.06E-03 | 3.56E-02 | P-type ATPase, A domain |
| GO:0000178 | 2 | 9.38E-04 | 3.35E-02 | Exosome complex component RRP45 |
| GO:0000287 | 16 | 4.96E-06 | 8.86E-04 | Terpene synthase, metal-binding domain |
| GO:0003910 | 3 | 1.43E-03 | 4.51E-02 | DNA ligase, ATP-dependent, N-terminal |
| GO:0004185 | 11 | 4.56E-05 | 4.07E-03 | Peptidase S10, serine carboxypeptidase |
| GO:0004373 | 3 | 9.15E-04 | 3.35E-02 | Bacterial/plant glycogen synthase |
| GO:0005488 | 27 | 9.33E-04 | 3.35E-02 | Armadillo-type fold |
| GO:0005515 | 160 | 3.46E-04 | 1.69E-02 | Ankyrin repeat |
| GO:0005524 | 129 | 2.56E-07 | 9.02E-05 | Protein kinase domain |
| GO:0006281 | 10 | 7.89E-04 | 3.35E-02 | XPG/Rad2 endonuclease |
| GO:0006508 | 28 | 2.18E-04 | 1.30E-02 | Peptidase S8/S53 domain |
| GO:0010333 | 9 | 1.32E-05 | 1.76E-03 | Terpene synthase, N-terminal domain |
| GO:0016491 | 47 | 4.39E-05 | 4.07E-03 | Oxoglutarate/iron-dependent dioxygenase |
| GO:0016829 | 9 | 5.51E-05 | 4.22E-03 | Terpene synthase, N-terminal domain |
| GO:0016844 | 4 | 2.43E-04 | 1.30E-02 | Strictosidine synthase, conserved region |
| GO:0043531 | 29 | 9.30E-05 | 6.23E-03 | NB-ARC |
| GO:0055114 | 86 | 3.37E-07 | 9.02E-05 | Cytochrome P450, E-class, group I |

Supplementary Table 23: The average expression of terpene selection genes in CSS

| Gene | Bud | Flower | Leaf | Stem | Root |
|---|---|---|---|---|---|
| Cha6.1001 | 0.407081 | 0.257628 | 0.0419781 | 0.120084 | 0.494778 |
| Cha12.1130 | 14.1382 | 0.384967 | 2.49478 | 0.441107 | 0.414616 |
| Cha2.1172 | 0 | 0 | 0 | 10.0337 | 1.63255 |
| Cha5.764 | 0.074889 | 0 | 0 | 0.15892 | 0 |
| Cha5.908* | 5.54577 | 0.704239 | 0.091366 | 0.182698 | 0.666819 |
| Cha5.693 | 1.40924 | 1.57527 | 0 | 0.371166 | 0.380853 |
| Cha5.401* | 34.2759 | 19.6657 | 159.187 | 70.1075 | 16.6851 |
| Cha10.244 | 8.16136 | 6.08036 | 22.9976 | 0.00750407 | 20.0126 |
| Cha9.1387* | 6.23429 | 10.8835 | 29.8996 | 0.0307483 | 6.57594 |
| Cha4.48 | 0.0467892 | 1.4549 | 0 | 0.0198882 | 0.238279 |
| Cha12.700 | 0.947587 | 0.45224 | 0.894742 | 0.460001 | 0.827391 |
| Cha4.1595 | 10.9641 | 0.955113 | 0 | 0.494315 | 0.0972183 |
| Cha6.581* | 60.7087 | 0.173015 | 0.112903 | 1.51996 | 1.70406 |
| Cha2.1198 | 1.96537 | 4.41655 | 13.1417 | 35.7095 | 2.82229 |
| Cha2.1199* | 0.417747 | 1.39067 | 11.0507 | 0.0579967 | 1.25263 |

Note: The average genes expression of tissue was calculated, while t.test was used to identify significant difference. "*" indicates that the gene expression of bud or leaf was significantly higher than that in other tissues.

Supplementary Table 24: The average expression of NB-ARC genes in CSS.

| Gene | Spring | Summer | Autumn | Winter |
|---|---|---|---|---|
| Cha1.388 | 0.385611 | 0.396025 | 0.493984 | 0.675391 |
| Cha8.782 | 0.0720208 | 0 | 0.0810365 | 0.0179926 |
| Cha10.62 | 3.20988 | 2.88009 | 1.96424 | 1.90201 |
| Cha8.1593 | 4.36668 | 3.83948 | 3.99813 | 4.42616 |
| Cha3.216 | 0.0299588 | 0.154957 | 0.0776982 | 0.0200274 |
| Cha13.297* | 12.0024 | 7.37154 | 9.10494 | 12.3034 |
| Cha10.1031 | 0.291862 | 0.442867 | 0.313353 | 0.112577 |
| Cha3.408* | 3.68761 | 4.74686 | 2.65007 | 1.90488 |
| Cha5.927* | 1.15157 | 1.00833 | 0.515501 | 0.291326 |
| Cha5.933* | 4.21901 | 0.945205 | 2.58612 | 2.58561 |
| Cha13.203 | 0.437237 | 0.464759 | 0.835618 | 0.206354 |
| Cha15.199 | 0.0729581 | 1.01925 | 0.155762 | 0.0641214 |
| Cha15.196 | 9.69115 | 2.97109 | 4.58856 | 8.01155 |
| Cha3.1486 | 0.214578 | 0.230928 | 0.237035 | 0.129093 |
| Cha2.260 | 0.137786 | 0.303307 | 0.132342 | 0.0669902 |
| Cha15.711 | 1.21677 | 0.608785 | 1.08043 | 1.13186 |
| Cha1.2079 | 6.37278 | 3.40941 | 4.19738 | 6.20538 |
| Cha10.265 | 0.753061 | 1.92344 | 0.588219 | 0.385155 |
| Cha7.622 | 0.565678 | 0.62565 | 0.528498 | 0.0950264 |
| Cha8.1550 | 0.0950757 | 0.0418575 | 0.189681 | 0.209651 |
| Cha14.438* | 6.82429 | 3.48099 | 8.11467 | 4.44938 |
| Cha5.1205 | 2.89219 | 1.9246 | 3.30116 | 2.86623 |
| Cha10.659 | 0.251886 | 0.0882413 | 0.0810854 | 0.0168054 |
| Cha9.1005* | 3.37949 | 1.30861 | 2.70701 | 1.95418 |
| Cha2.488 | 2.95852 | 2.46588 | 2.26903 | 3.20311 |
| Cha13.137* | 0.0120879 | 0.0417264 | 0.0832469 | 0.111354 |
| Cha2.712 | 0.795715 | 1.33952 | 0.957988 | 0.476364 |
| Cha13.226 | 4.25625 | 4.1153 | 4.29495 | 4.67308 |
| Cha5.1382 | 0.938097 | 6.38732 | 1.10065 | 0.615442 |

Note: The average genes expression of season was calculated, while t-test was used to identify significant difference. "*" indicates that the gene expression in the summer was significantly lower than that in other seasons.

Supplementary Table 25: The high heterozygosity genes keep tea.

| GO | Number |
|---|---|
| Cytochrome P450 | 12 |
| UDP-glucuronosyl/UDP-glucosyltransferase | 13 |
| Small auxin-up RNA | 2 |
| Terpene synthase | 3 |
| Cytokinin dehydrogenase | 1 |
| Multi antimicrobial extrusion protein | 5 |
| NB-ARC | 10 |
| S-locus | 10 |
| AP2/ERF domain | 3 |
| Malic oxidoreductase | 1 |
| NAC domain | 4 |
| WD40 | 8 |

Supplementary Table 26: The groups of the treemix.

The table is supplied in supplementary table 26.excel.

Supplementary Table 27: The results of F4-test and F3-test

| Samples | Statistic | Standard error | Z-score |
|---|---|---|---|
| DBZ-1,HZ002;HSKC,CM-1 | -0.000260712 | 5.76778e-05 | -4.52014 |
| CSA2, CSR; CSA-test, CM-1 | 0.00696553 | 3.23781e-05 | 215.131 |
| CSA-test,CSA2;CSR,CM-1 | -0.0004924 | 1.89843e-05 | -25.9373 |
| HZ100,HZ118;HZ122,CM-1 | -0.00312965 | 6.77548e-05 | -46.1909 |
| HZ100,HZ122;HZ118,CM-1 | -0.00241358 | 6.89268e-05 - | 35.0166 |
| CSA,CSS2;CSS,CM-1 | -0.0129479 | 5.63863e-05 | -229.628 |
| CSA,CSS;CSS2,CM-1 | -0.013012 | 5.56352e-05 | -233.88 |
| CSA,CSA2;CSR,CM-1 | -0.0004924 | 2.18646e-05 | -22.5204 |
| CSA2,CSR;CSA,CM-1 | 0.00696553 | 3.85973e-05 | 180.467 |
| HZ104,HZ114;HZ117,CM-1 | 0.00619503 | 5.12426e-05 | 120.896 |
| CSA,CSR;CSA2,CM-1 | 0.00647313 | 3.82574e-05 | 169.199 |
| CSA2,CSA;CSR,CM-1 | 0.0004924 | 2.18646e-05 | 22.5204 |
| HZ104,HZ117;HZ114,CM-1 | 0.00858967 | 5.77297e-05 | 148.791 |
| DBZ-1,HSKC;HZ002,CM-1 | 0.00255663 | 6.59185e-05 | 38.7847 |
| HZ039,HZ074;HZ092,CM-1 | -0.000293896 | 5.69659e-05 | -5.15916 |
| HZ021,YNLDP1;HZ050,CM-1 | 0.00881408 | 5.79118e-05 | 152.198 |
| HZ021,HZ050;YNLDP1,CM-1 | 0.00694945 | 5.04036e-05 | 137.876 |
| HZ114,HZ117;HZ104,CM-1 | 0.00239463 | 7.2061e-05 | 33.2306 |
| CSA2;CSA,CSR | 0.00865398 | 3.81241e-05 | 226.995 |
| CSA;CSA2,CSR | 0.00669717 | 4.09978e-05 | 163.354 |
| CSR;CSA,CSA2 | 0.0179839 | 4.66329e-05 | 385.648 |
| CSA;CSS,CSS2 | 0.023809 | 6.07489e-05 | 391.924 |
| CSS2;CSA,CSS | 0.00712921 | 4.19279e-05 | 170.035 |
| CSS;CSA,CSS2 | 0.00476459 | 4.15159e-05 | 114.766 |

Note: CSA-test contained HZ114, HZ119, and HZ104; CSR contained NC, HZ084, HZ001, XYDCS, LBDCS, and HZ027; CSA contained HZ104, HZ114, and HZ119; CSA2 contained HZ118, HZ122, HZ100, HZ072, HZ123, and HZ117; CSS contained HZ050, HZ021, QXDM-1, and QXDM-2; CSS2 contained HZ016, HZ036_HZ036, HZ008, and HZ041; CSA contained XH-1, ASM, YH-2, and HZ118. The A,B;C,D was F4-test, and the A;B,C was F3-test.

Supplementary Table 28: F4-test of random individual.

The table is supplied in supplementary table 28.excel.
