## Supplementary figures and images for "Population sequencing enhances understanding of tea plant evolution"

### Supplementary fig 12

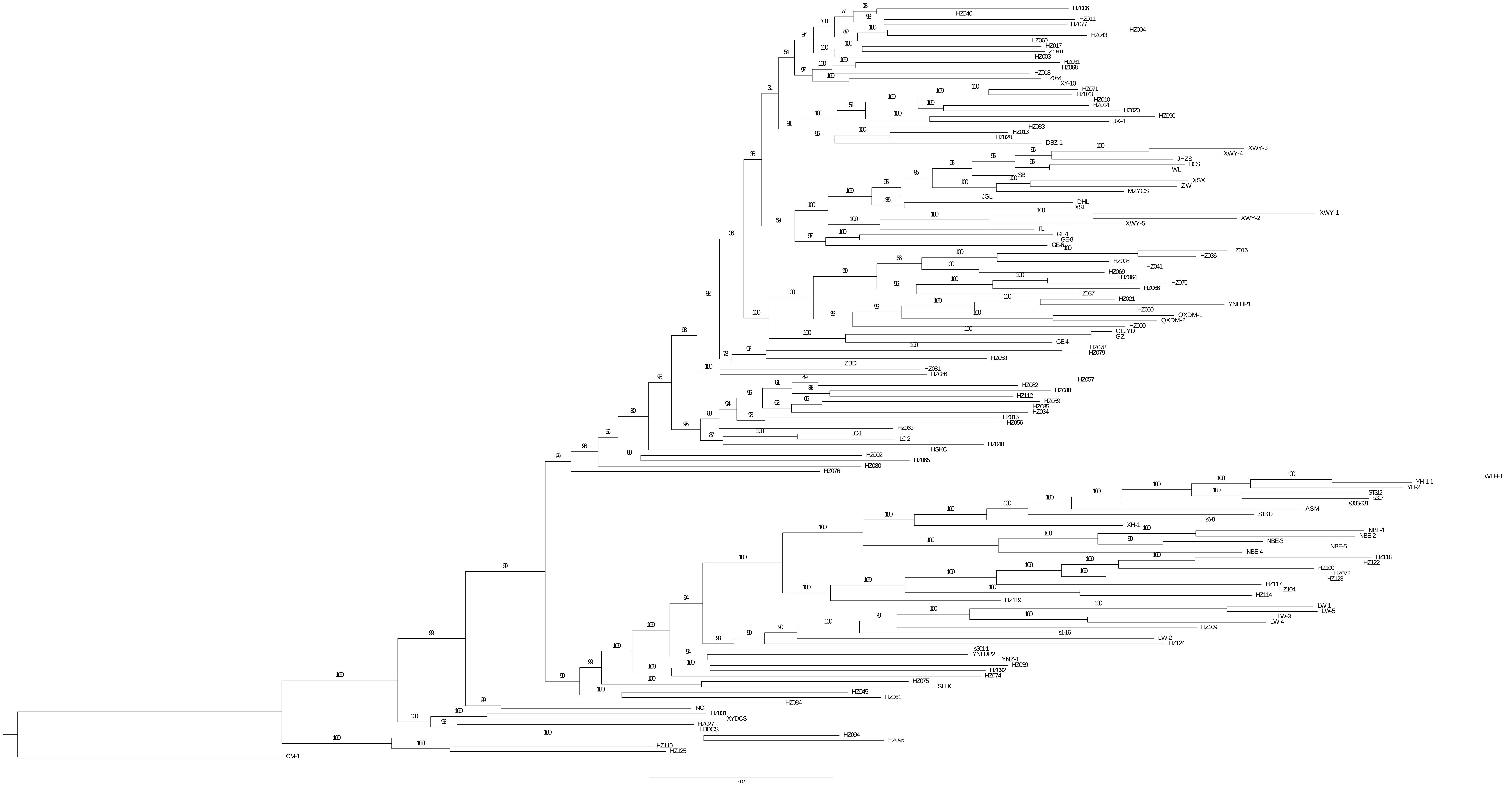

### Supplementary fig 14

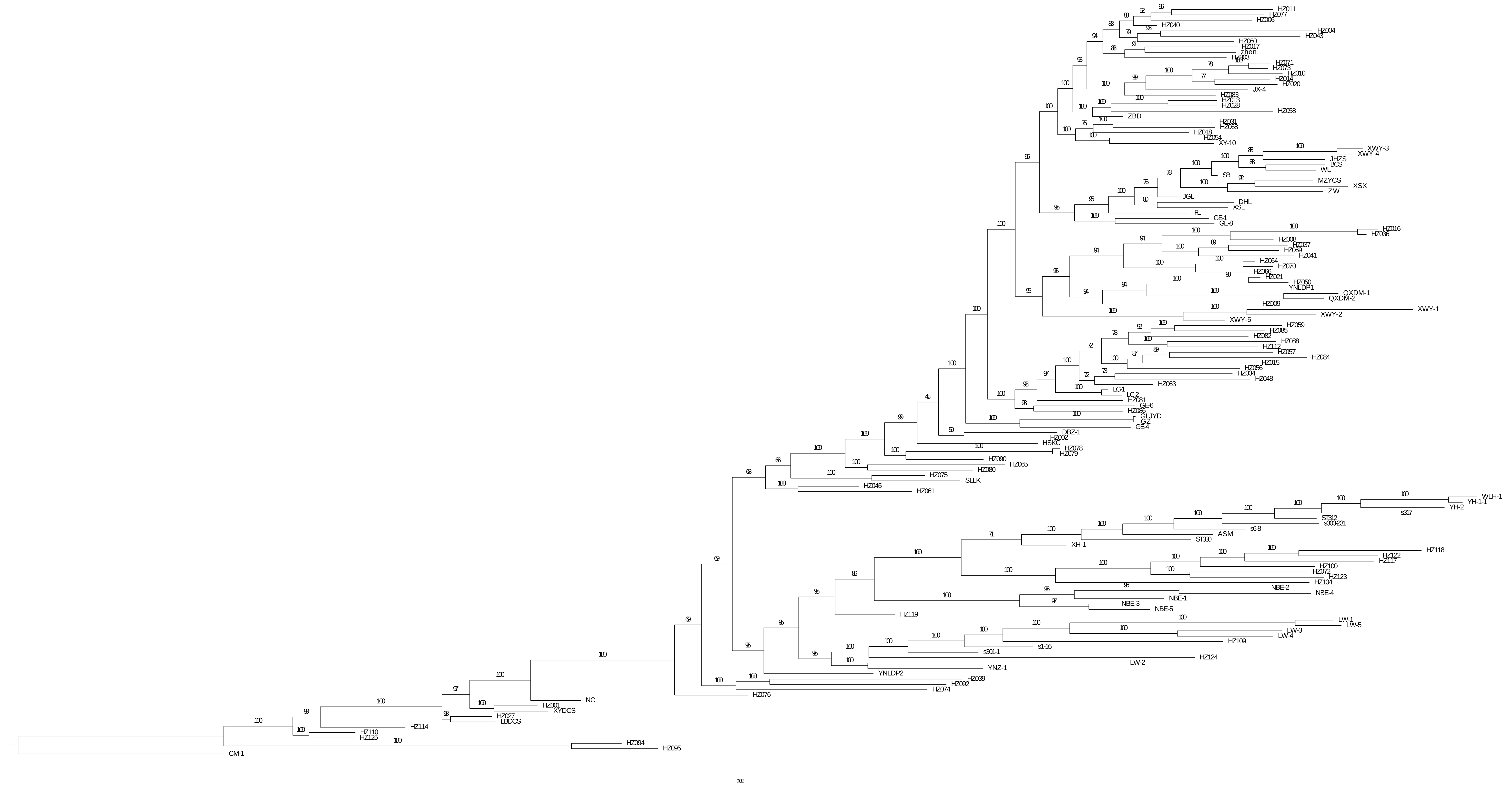

### Supplementary fig 15

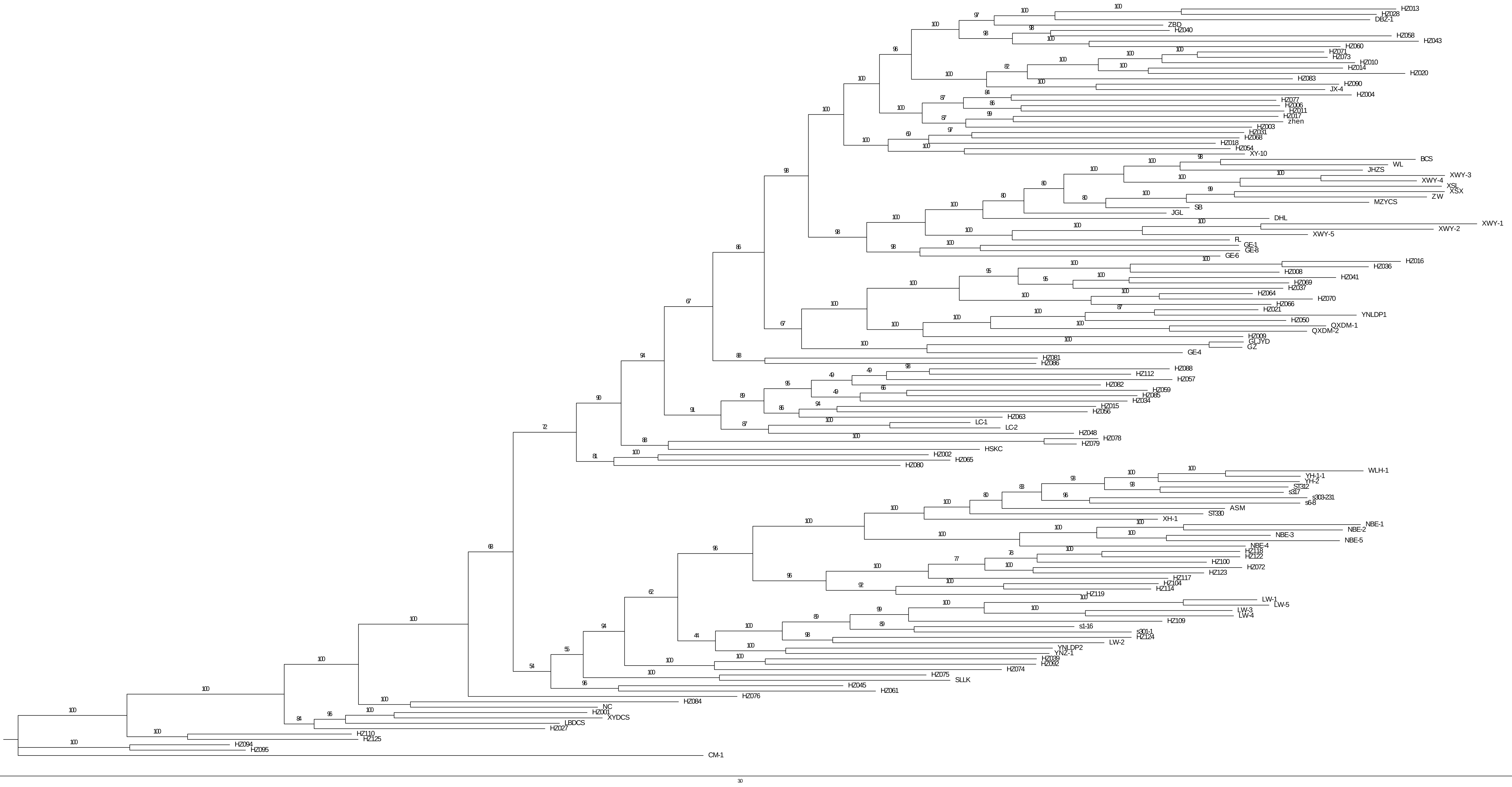
