## Supplementary tables for "Population sequencing enhances understanding of tea plant evolution"

Supplementary Table 11: IPR enrichment of expansion genes.

| IPR | All genes | Expansion genes | P-value | FDR |
|---|---|---|---|---|
| IPR000209 | 85 | 55 | 3.20E-16 | 3.86E-14 |
| IPR000528 | 19 | 12 | 2.16E-04 | 3.92E-03 |
| IPR000668 | 44 | 19 | 2.45E-03 | 2.74E-02 |
| IPR000863 | 39 | 24 | 3.03E-07 | 1.37E-05 |
| IPR000916 | 28 | 15 | 4.49E-04 | 7.40E-03 |
| IPR001046 | 16 | 11 | 1.31E-04 | 2.80E-03 |
| IPR001781 | 13 | 11 | 4.76E-06 | 1.57E-04 |
| IPR002110 | 254 | 85 | 9.91E-05 | 2.18E-03 |
| IPR002182 | 436 | 263 | 2.20E-16 | 3.19E-14 |
| IPR002213 | 290 | 175 | 2.20E-16 | 3.19E-14 |
| IPR002528 | 72 | 32 | 4.99E-05 | 1.25E-03 |
| IPR002781 | 7 | 6 | 8.46E-04 | 1.06E-02 |
| IPR003480 | 156 | 52 | 2.20E-03 | 2.54E-02 |
| IPR003653 | 22 | 12 | 1.37E-03 | 1.62E-02 |
| IPR003855 | 31 | 25 | 1.91E-11 | 1.30E-09 |
| IPR004148 | 6 | 6 | 1.51E-04 | 3.03E-03 |
| IPR004240 | 29 | 21 | 2.44E-08 | 1.18E-06 |
| IPR004326 | 34 | 19 | 3.64E-05 | 9.43E-04 |
| IPR004487 | 4 | 4 | 2.83E-03 | 2.74E-02 |
| IPR004522 | 4 | 4 | 2.83E-03 | 2.74E-02 |
| IPR004813 | 23 | 12 | 2.26E-03 | 2.56E-02 |
| IPR004859 | 6 | 5 | 3.17E-03 | 2.98E-02 |
| IPR004882 | 4 | 4 | 2.83E-03 | 2.74E-02 |
| IPR005381 | 6 | 5 | 3.17E-03 | 2.98E-02 |
| IPR005484 | 11 | 8 | 6.66E-04 | 9.28E-03 |
| IPR005630 | 38 | 28 | 5.44E-11 | 3.29E-09 |
| IPR005821 | 33 | 26 | 1.98E-11 | 1.30E-09 |
| IPR006685 | 17 | 11 | 2.95E-04 | 5.08E-03 |
| IPR007369 | 9 | 8 | 5.73E-05 | 1.34E-03 |
| IPR008089 | 7 | 6 | 8.46E-04 | 1.06E-02 |
| IPR009014 | 14 | 9 | 1.16E-03 | 1.43E-02 |
| IPR011025 | 8 | 7 | 2.22E-04 | 3.92E-03 |
| IPR011068 | 9 | 7 | 7.98E-04 | 1.05E-02 |
| IPR011356 | 5 | 5 | 6.53E-04 | 9.28E-03 |
| IPR011531 | 9 | 7 | 7.98E-04 | 1.05E-02 |
| IPR011555 | 9 | 7 | 7.98E-04 | 1.05E-02 |
| IPR011701 | 43 | 20 | 6.11E-04 | 9.28E-03 |
| IPR012110 | 5 | 5 | 6.53E-04 | 9.28E-03 |
| IPR012317 | 22 | 11 | 5.24E-03 | 4.75E-02 |
| IPR013094 | 81 | 42 | 1.69E-08 | 8.76E-07 |
| IPR013215 | 8 | 7 | 2.22E-04 | 3.92E-03 |
| IPR014525 | 4 | 4 | 2.83E-03 | 2.74E-02 |
| IPR014712 | 15 | 11 | 5.20E-05 | 1.26E-03 |
| IPR015341 | 7 | 6 | 8.46E-04 | 1.06E-02 |
| IPR017163 | 8 | 8 | 8.01E-06 | 2.52E-04 |

| | | | | |
|---|---|---|---|---|
| IPR017441 | 920 | 528 | 2.20E-16 | 3.19E-14 |
| IPR017761 | 32 | 30 | 2.20E-16 | 3.19E-14 |
| IPR017972 | 211 | 101 | 2.41E-15 | 2.50E-13 |
| IPR017975 | 25 | 13 | 1.55E-03 | 1.81E-02 |
| IPR018045 | 9 | 9 | 1.85E-06 | 6.37E-05 |
| IPR018088 | 5 | 5 | 6.53E-04 | 9.28E-03 |
| IPR018146 | 8 | 7 | 2.22E-04 | 3.92E-03 |
| IPR018167 | 5 | 5 | 6.53E-04 | 9.28E-03 |
| IPR018200 | 43 | 18 | 4.72E-03 | 4.33E-02 |
| IPR018225 | 4 | 4 | 2.83E-03 | 2.74E-02 |
| IPR018259 | 4 | 4 | 2.83E-03 | 2.74E-02 |
| IPR018338 | 10 | 9 | 1.46E-05 | 4.24E-04 |
| IPR018368 | 9 | 9 | 1.85E-06 | 6.37E-05 |
| IPR018371 | 13 | 10 | 6.02E-05 | 1.36E-03 |
| IPR018451 | 28 | 21 | 8.63E-09 | 4.81E-07 |
| IPR018456 | 22 | 15 | 8.64E-06 | 2.61E-04 |
| IPR019780 | 38 | 21 | 1.79E-05 | 4.81E-04 |
| IPR019787 | 80 | 31 | 1.20E-03 | 1.44E-02 |
| IPR019798 | 12 | 10 | 1.76E-05 | 4.81E-04 |
| IPR019801 | 21 | 20 | 2.94E-12 | 2.66E-10 |
| IPR020843 | 39 | 23 | 1.56E-06 | 6.27E-05 |
| IPR020904 | 40 | 20 | 1.82E-04 | 3.56E-03 |
| IPR021792 | 4 | 4 | 2.83E-03 | 2.74E-02 |
| IPR022636 | 5 | 5 | 6.53E-04 | 9.28E-03 |
| IPR022740 | 6 | 6 | 1.51E-04 | 3.03E-03 |
| IPR022893 | 5 | 5 | 6.53E-04 | 9.28E-03 |
| IPR023299 | 75 | 37 | 6.02E-07 | 2.57E-05 |
| IPR024171 | 48 | 47 | 2.20E-16 | 3.19E-14 |
| IPR024750 | 9 | 9 | 1.85E-06 | 6.37E-05 |
| IPR025287 | 71 | 43 | 1.42E-11 | 1.14E-09 |
| IPR027246 | 13 | 8 | 3.30E-03 | 3.07E-02 |
| IPR033122 | 4 | 4 | 2.83E-03 | 2.74E-02 |
| IPR033905 | 78 | 32 | 3.09E-04 | 5.21E-03 |
| IPR033939 | 4 | 4 | 2.83E-03 | 2.74E-02 |
| IPR033950 | 4 | 4 | 2.83E-03 | 2.74E-02 |

| Function |
|---|
| Peptidase S8/S53 domain |
| Plant lipid transfer protein/Par allergen |
| Peptidase C1A, papain C-terminal |
| Sulfotransferase domain |
| Bet v I/Major latex protein |
| NRAMP family |
| Zinc finger, LIM-type |
| Ankyrin repeat |
| NB-ARC |
| UDP-glucuronosyl/UDP-glucosyltransferase |
| Multi antimicrobial extrusion protein |
| Transmembrane protein TauE-like |
| Transferase |
| Ulp1 protease family, C-terminal catalytic domain |
| Potassium transporter |
| BAR domain |
| Nonaspanin (TM9SF) |
| Mlo-related protein |
| Clp protease, ATP-binding subunit ClpX |
| Asparagine-tRNA ligase |
| Oligopeptide transporter, OPT superfamily |
| Putative 5-3 exonuclease |
| Luc7-related |
| Zinc finger-XS domain |
| Ribosomal protein L18 |
| Terpene synthase, metal-binding domain |
| Ion transport domain |
| Mechanosensitive ion channel MscS |
| Peptidase A22B, signal peptide peptidase |
| Nucleotide sugar epimerase |
| Transketolase C-terminal/Pyruvate-ferredoxin oxidoreductase domain II |
| G protein alpha subunit, helical insertion |
| Nucleotidyltransferase, class I, C-terminal-like |
| Peptidase M17, leucine aminopeptidase/peptidase B |
| Bicarbonate transporter, C-terminal |
| V-ATPase proteolipid subunit C, eukaryotic |
| Major facilitator superfamily |
| Thiamine pyrophosphate (TPP)-dependent enzyme |
| Poly(ADP-ribose) polymerase, catalytic domain |
| Alpha/beta hydrolase fold-3 |
| Cobalamin-independent methionine synthase MetE, N-terminal |
| Ethylene receptor |
| Phosphoinositide-binding clathrin adaptor, domain 2 |
| Glycoside hydrolase family 38, central domain |
| Phosphatidylinositol-4-phosphate 5-kinase, plant |

Protein kinase, ATP binding site
Laccase
Cytochrome P450, conserved site
Tubulin, conserved site
Sulphate anion transporter, conserved site
Chalcone/stilbene synthase, active site
Glyoxalase I, conserved site
S-adenosylmethionine decarboxylase subgroup
Ubiquitin specific protease, conserved site
Transaldolase, active site
Ribosomal protein L21e, conserved site
Carbonic anhydrase, alpha-class, conserved site
ClpA/B, conserved site 1
Chitin-binding, type 1, conserved site
NAF/FISL domain
PTR2 family proton/oligopeptide symporter, conserved site
Germin, manganese binding site
Zinc finger, PHD-finger
Serine hydroxymethyltransferase, pyridoxal phosphate binding site
Glycoside hydrolase, family 35, conserved site
Polyketide synthase, enoylreductase domain
Short-chain dehydrogenase/reductase, conserved site
Beta-fructofuranosidase
S-adenosylmethionine synthetase superfamily
Polyphenol oxidase, C-terminal
Shikimate dehydrogenase family
P-type ATPase, cytoplasmic domain N
S-receptor-like serine/threonine-protein kinase
Calcium-transporting P-type ATPase, N-terminal autoinhibitory domain
Wall-associated receptor kinase, galacturonan-binding domain
Eukaryotic porin/Tom40
Letm1 ribosome-binding domain
Secretory peroxidase
Branched-chain aminotransferase
Sumo domain

Supplementary Table 15: The positive Darwinian selection genes of LJ43.

| Gene name |
|---|
| Cha10.476 |
| Cha10.538 |
| Cha10.933 |
| Cha11.1056 |
| Cha11.1272 |
| Cha11.183 |
| Cha1.1446 |
| Cha1.1485 |
| Cha11.967 |
| Cha1.2063 |
| Cha12.1252 |
| Cha12.257 |
| Cha1.2357 |
| Cha1.242 |
| Cha13.423 |
| Cha13.428 |
| Cha13.655 |
| Cha13.690 |
| Cha13.931 |
| Cha14.1115 |
| Cha14.1250 |
| Cha14.159 |
| Cha14.562 |
| Cha14.738 |
| Cha14.979 |
| Cha15.411 |
| Cha1.558 |
| Cha15.953 |
| Cha2.1494 |
| Cha2.1874 |
| Cha2.891 |
| Cha2.938 |
| Cha3.1543 |
| Cha3.1739 |
| Cha3.2114 |
| Cha3.2142 |
| Cha3.25 |
| Cha3.29 |
| Cha3.364 |
| Cha3.366 |
| Cha4.527 |
| Cha4.705 |
| Cha4.863 |
| Cha5.1485 |

| |
|---|
| Cha5.1614 |
| Cha5.171 |
| Cha5.238 |
| Cha5.654 |
| Cha5.670 |
| Cha5.819 |
| Cha6.1760 |
| Cha6.1885 |
| Cha6.844 |
| Cha7.301 |
| Cha7.318 |
| Cha7.460 |
| Cha7.676 |
| Cha7.898 |
| Cha8.1635 |
| Cha8.935 |
| Cha9.1355 |
| Cha9.301 |
| Cha9.5 |
| Cha9.682 |
| Cha9.946 |
| ChaUn10341.2 |
| ChaUn10409.4 |
| ChaUn11121.1 |
| ChaUn11930.3 |
| ChaUn21494.1 |
| ChaUn24968.1 |
| ChaUn4163.1 |
| ChaUn5208.2 |
| ChaUn9520.1 |

Supplementary Table 16: The informations of tea population.

| Sample | Collected site | Species | Group | Tree-type | Leaf Size |
|---|---|---|---|---|---|
| HZ001 | Xuyong Szechwan | *C. sinensis* | CSR | semi-arbor | middle |
| HZ002 | Ziyuan Guangxi | *C. sinensis* var. *pubilimba* | CSS | - | middle |
| HZ003 | Qingdao Shandong | *C. sinensis* | CSS | shrub | middle |
| HZ004 | Tiantai Zhejiang | *C. sinensis* | CSS | shrub | middle |
| HZ006 | Ruian Zhejiang | *C. sinensis* | CSS | shrub | middle |
| HZ008 | Anxi Fujian | *C. sinensis* | CSS | shrub | middle |
| HZ009 | Ziyuan Guangxi | *C. sinensis* | CSS | shrub | middle |
| HZ010 | Changsha Hunan | *C. sinensis* | CSS | shrub | middle |
| HZ011 | Yongjia Zhejiang | *C. sinensis* | CSS | shrub | middle |
| HZ013 | Shucheng Anhui | *C. sinensis* | CSS | shrub | middle |
| HZ014 | Fuan Fujian | *C. sinensis* | CSS | semi-arbor | large |
| HZ015 | Guilin Guangxi | *C. sinensis* | CSS | shrub | middle |
| HZ016 | Fuan Fujian | *C. sinensis* | CSS | shrub | middle |
| HZ017 | Wuxi Jiangshu | *C. sinensis* | CSS | shrub | large |
| HZ018 | Qimen Anhui | *C. sinensis* | CSS | shrub | large |
| HZ020 | Changsha Hunan | *C. sinensis* | CSS | shrub | middle |
| HZ021 | Taiwan | *C. sinensis* | CSS | shrub | middle |
| HZ027 | Wuchuan Guizhou | *C.sp* | CSR | arbor | middle |
| HZ028 | Wuyuan Jiangxi | *C. sinensis* | CSS | shrub | large |
| HZ031 | Mount Wuyi Fujian | *C. sinensis* | CSS | shrub | middle |
| HZ034 | Xianning Hubei | *C. sinensis* | CSS | shrub | middle |
| HZ036 | Anxi Fujian | *C. sinensis* | CSS | shrub | middle |
| HZ037 | Mount Wuyi Fujian | *C. sinensis* | CSS | shrub | small |
| HZ039 | Ruyuan Guangdong | *C. sinensis* var. *pubilimba* | CSA | semi-arbor | middle |
| HZ040 | Anxi Fujian | *C. sinensis* | CSS | shrub | large |
| HZ041 | Anxi Fujian | *C. sinensis* | CSS | semi-arbor | middle |

| | | | | | |
|---|---|---|---|---|---|
| HZ043 | Hefei Anhui | *C. sinensis* | CSS | shrub | middle |
| HZ045 | Menghai Yunnan | *C. sinensis* var. *assamica* | CSA | arbor | large |
| HZ048 | Suichuan Jiangxi | *C. sinensis* | CSS | shrub | middle |
| HZ050 | Menghai Yunnan | *C. sinensis* var. *assamica* | CSS | arbor | large |
| HZ054 | Anqing Anhui | *C. sinensis* | CSS | shrub | middle |
| HZ056 | Meitan Guizhou | *C. sinensis* | CSS | shrub | middle |
| HZ057 | Chongqing Szechwa | *C. sinensis* | CSS | semi-arbor | middle |
| HZ058 | Linhai Zhejiang | *C. sinensis* | CSS | shrub | middle |
| HZ059 | Ankang shaanxi | *C. sinensis* | CSS | shrub | middle |
| HZ060 | Mingshan Szechwan | *C. sinensis* | CSS | shrub | middle |
| HZ061 | Guilin Guangxi | *C. sinensis* var. *assamica* | CSA | semi-arbor | large |
| HZ063 | Taoyuan Hunan | *C. sinensis* | CSS | shrub | large |
| HZ064 | Mount Wuyi Fujian | *C. sinensis* | CSS | shrub | middle |
| HZ065 | Zaoan Fujian | *C. sinensis* | CSS | semi-arbor | large |
| HZ066 | Fuan Fujian | *C. sinensis* | CSS | shrub | middle |
| HZ068 | Mount Wuyi Fujian | *C. sinensis* | CSS | shrub | middle |
| HZ069 | Anxi Fujian | *C. sinensis* | CSS | shrub | middle |
| HZ070 | Fuan Fujian | *C. sinensis* | CSS | shrub | middle |
| HZ071 | Songxi Fujian | *C. sinensis* | CSS | semi-arbor | large |
| HZ072 | Jinghong Yunnan | *C. sinensis* var. *assamica* | CSA | arbor | large |
| HZ073 | Fuding Fujian | *C. sinensis* | CSS | semi-arbor | middle |
| HZ074 | Longsheng Guangxi | *C. sinensis* var. *pubilimba* | CSA | semi-arbor | middle |
| HZ075 | Taiwan | *C. sinensis* var. *assamica* | CSA | semi-arbor | large |
| HZ076 | Hainan | *C. sinensis* var. *assamica* | CSS | arbor | large |
| HZ077 | Qionglai Szechwan | *C. sinensis* | CSS | semi-arbor | middle |
| HZ078 | Jinxiu Guangxi | *C. sinensis* var. *pubilimba* | CSS | shrub | middle |
| HZ079 | Shangsi Guangxi | *C. sinensis* var. *pubilimba* | CSS | arbor | large |
| HZ080 | Jianghua Hunan | *C. sinensis* | CSS | semi-arbor | large |
| HZ081 | Guiyang Guizhou | *C. sinensis* | CSS | semi-arbor | middle |
| HZ082 | Chaozhou Guangdo | *C. sinensis* | CSS | semi-arbor | middle |

| | | | | | |
|---|---|---|---|---|---|
| HZ083 | Anxi Fujian | *C. sinensis* | CSS | shrub | middle |
| HZ084 | Xishui Guizhou | *C.sp* | CSR | semi-arbor | middle |
| HZ085 | Nanjiang Szechwan | *C. sinensis* | CSS | shrub | middle |
| HZ086 | Mount Wuyi Fujian | *C. sinensis* | CSS | shrub | middle |
| HZ088 | Daozhen Guizhou | *C.sp* | CSS | arbor | large |
| HZ090 | Longzhou Guangxi | *C. sinensis* var. *pubilimba* | CSS | arbor | large |
| HZ092 | Longmen Guangdor | *C. sinensis* | CSA | shrub | middle |
| HZ094 | Nayong Guizhou | *C.sp* | CSR | - | - |
| HZ095 | Malipo Yunnan | *C. crassicolumna* | CSR | arbor | large |
| HZ100 | Lincang Yunnan | *C. sinensis* var. *assamica* | CSA | arbor | large |
| HZ104 | Ruili Yunnan | *C. gymnogyna* | CSA | arbor | large |
| HZ109 | Xichou Yunnan | *C. sinensis* | CSA | semi-arbor | large |
| HZ110 | Nanjian Yunnan | *C. atrothea* | CSR | arbor | large |
| HZ112 | Maguan Yunnan | *C. sinensis* | CSS | shrub | middle |
| HZ114 | Longling Yunnan | *C. taliensis* | CSA | arbor | large |
| HZ117 | Fengqing Yunnan | *C. sinensis* var. *assamica* | CSA | arbor | large |
| HZ118 | Menghai Yunnan | *C. sinensis* var. *assamica* | CSA | arbor | large |
| HZ119 | Shuangjiang Yunna | *C. sinensis* var. *assamica* | CSA | arbor | extra large |
| HZ122 | Longchuan Yunnan | *C. gymnogyna* | CSA | arbor | large |
| HZ123 | Mengla Yunnan | *C. sinensis* var. *assamica* | CSA | arbor | large |
| HZ124 | Wenshan Yunnan | *C. sinensis* var. *assamica* | CSA | semi-arbor | large |
| HZ125 | Fengqing Yunnan | *C. taliensis* | CSR | arbor | large |
| NBE-1 | Nepal | *C. sinensis* var. *assamica* | CSA | shrub | middle |
| NBE-2 | Nepal | *C. sinensis* var. *assamica* | CSA | shrub | middle |
| NBE-3 | Nepal | *C. sinensis* var. *assamica* | CSA | shrub | middle |
| NBE-4 | Nepal | *C. sinensis* var. *assamica* | CSA | shrub | middle |
| NBE-5 | Nepal | *C. sinensis* var. *assamica* | CSA | shrub | middle |
| XWY-1 | Hawaii | *C. sinensis* | CSS | shrub | middle |
| XWY-2 | Hawaii | *C. sinensis* | CSS | shrub | middle |
| XWY-3 | Hawaii | *C. sinensis* | CSS | shrub | middle |

| | | | | | |
|---|---|---|---|---|---|
| XWY-4 | Hawaii | *C. sinensis* | CSS | shrub | middle |
| XWY-5 | Hawaii | *C. sinensis* | CSS | shrub | middle |
| GE-1 | Georgia | *C. sinensis* | CSS | shrub | middle |
| GE-4 | Georgia | *C. sinensis* | CSS | shrub | middle |
| GE-6 | Georgia | *C. sinensis* | CSS | shrub | middle |
| GE-8 | Georgia | *C. sinensis* | CSS | shrub | middle |
| LW-1 | Laos | *C. sinensis* var. *assamica* | CSA | semi-arbor | large |
| LW-2 | Laos | *C. sinensis* var. *assamica* | CSA | semi-arbor | large |
| LW-3 | Laos | *C. sinensis* var. *assamica* | CSA | semi-arbor | large |
| LW-4 | Laos | *C. sinensis* var. *assamica* | CSA | semi-arbor | large |
| LW-5 | Laos | *C. sinensis* var. *assamica* | CSA | semi-arbor | large |
| LC-1 | Rizhao Shandong | *C. sinensis* | CSS | shrub | middle |
| LC-2 | Rizhao Shandong | *C. sinensis* | CSS | shrub | middle |
| GZ | Georgia | *C. sinensis* | CSS | shrub | middle |
| DHL | Japan | *C. sinensis* | CSS | shrub | middle |
| ZW | Japan | *C. sinensis* | CSS | shrub | middle |
| WL | Japan | *C. sinensis* | CSS | shrub | middle |
| YNLDP2 | Vietnam | *C. sinensis* var. *assamica* | CSA | semi-arbor | large |
| GLJYD | Georgia | *C. sinensis* | CSS | semi-arbor | large |
| s303-231 | Kenya | *C. sinensis* var. *assamica* | CSA | semi-arbor | large |
| ST312 | Kenya | *C. sinensis* var. *assamica* | CSA | semi-arbor | large |
| BCS | Japan | *C. sinensis* | CSS | shrub | middle |
| s1-16 | Kenya | *C. sinensis* var. *assamica* | CSA | semi-arbor | large |
| XY-10 | Xinyang Henan | *C. sinensis* | CSS | shrub | middle |
| FL | Japan | *C. sinensis* | CSS | shrub | middle |
| JHZS | Japan | *C. sinensis* | CSS | shrub | middle |
| MZYZS | Japan | *C. sinensis* | CSS | shrub | middle |
| XSX | Japan | *C. sinensis* | CSS | shrub | middle |
| ASM | India | *C. sinensis* var. *assamica* | CSA | arbor | large |
| YNZ-1 | Vietnam | *C. sinensis* var. *assamica* | CSA | arbor | large |

| | | | | | |
|---|---|---|---|---|---|
| ST330 | Kenya | *C. sinensis* var. *assamica* | CSA | semi-arbor | large |
| s317 | Kenya | *C. sinensis* var. *assamica* | CSA | semi-arbor | large |
| QXDM-1 | Taiwan | *C. sinensis* | CSS | shrub | small |
| QXDM-2 | Taiwan | *C. sinensis* | CSS | shrub | small |
| DBZ-1 | Burma | *C. sinensis* var. *assamica* | CSS | arbor | large |
| ZBD | Szechwan | *C. sinensis* | CSS | shrub | middle |
| s301-1 | Kenya | *C. sinensis* var. *assamica* | CSA | semi-arbor | large |
| s6-8 | Kenya | *C. sinensis* var. *assamica* | CSA | semi-arbor | large |
| JX-4 | Jinxiu Guangxi | *C. sinensis* var. *pubilimba* | CSS | semi-arbor | middle |
| NC | Nanchuan Chongqir | *C.sp* | CSR | - | - |
| XYDCS | Xuyong Szechwan | *C. sinensis* | CSR | semi-arbor | large |
| HSKC | Yibing Szechwan | *C. sinensis* | CSS | arbor | large |
| LBDCS | Leibo Szechwan | *C.sp* | CSR | semi-arbor | extra large |
| SB | Japan | *C. sinensis* | CSS | shrub | middle |
| YNLDP1 | Vietnam | *C. sinensis* var. *assamica* | CSS | semi-arbor | large |
| JGL | Japan | *C. sinensis* | CSS | shrub | middle |
| zhen | Japan | *C. sinensis* | CSS | shrub | middle |
| SLLK | Sri Lanka | *C. sinensis* var. *assamica* | CSA | arbor | large |
| XSL | Japan | *C. sinensis* | CSS | shrub | middle |
| CM-1 | Hangzhou Zhejiang | *C. sasanqua* | outgroup | arbor | - |
| WLH-1 | Yingde Guangdong | *C. sinensis* var. *assamica* | CSA | semi-arbor | large |
| XH-1 | Yingde Guangdong | *C. sinensis* var. *assamica* | CSA | semi-arbor | large |
| YH-1-1 | Yingde Guangdong | *C. sinensis* var. *assamica* | CSA | arbor | large |
| YH-2 | Yingde Guangdong | *C. sinensis* var. *assamica* | CSA | arbor | large |

Cold tolerance：1>2>3>4.
″C.sp″ was uncertain taxonomy.
″-″ was no information

| Sprouting Date | Suitable tea | Cold tolerance | Location | Clean data reads | Clean data base（bp） | Depth |
|---|---|---|---|---|---|---|
| middle | black tea | 4 | Hangzhou | 282,914,640 | 42,186,021,678 | 12.94 |
| - | black tea | 2 | Hangzhou | 326,312,338 | 48,674,954,636 | 14.93 |
| middle | green tea | 1 | Hangzhou | 345,073,168 | 51,461,351,670 | 15.79 |
| late | green tea | 2 | Hangzhou | 308,929,732 | 46,156,090,908 | 14.16 |
| early | green tea | 2 | Hangzhou | 315,417,330 | 47,144,416,786 | 14.46 |
| middle | oolong tea | 2 | Hangzhou | 320,971,400 | 47,858,322,944 | 14.68 |
| - | green tea | 2 | Hangzhou | 294,102,400 | 43,971,985,144 | 13.49 |
| early | green tea | 1 | Hangzhou | 322,084,788 | 48,141,792,436 | 14.77 |
| extra early | green tea | 2 | Hangzhou | 315,870,352 | 47,231,941,500 | 14.49 |
| early | green tea | 1 | Hangzhou | 303,459,240 | 45,342,752,992 | 13.91 |
| extra early | green tea | 2 | Hangzhou | 322,302,726 | 48,057,050,318 | 14.74 |
| early | green tea | 2 | Hangzhou | 349,726,702 | 52,195,151,934 | 16.01 |
| middle | oolong tea | 2 | Hangzhou | 340,698,278 | 50,812,184,538 | 15.59 |
| middle | green tea | 2 | Hangzhou | 311,793,540 | 46,515,060,032 | 14.27 |
| middle | black tea | 1 | Hangzhou | 361,965,070 | 54,037,605,820 | 16.58 |
| early | black tea | 1 | Hangzhou | 220,650,016 | 32,000,756,616 | 9.82 |
| middle | oolong tea | 1 | Hangzhou | 295,603,130 | 44,114,361,526 | 13.53 |
| early | green tea | 4 | Hangzhou | 319,134,300 | 47,714,639,278 | 14.64 |
| early | green tea | 1 | Hangzhou | 313,874,516 | 46,868,642,600 | 14.38 |
| late | oolong tea | 2 | Hangzhou | 285,490,486 | 42,673,349,816 | 13.09 |
| early | black tea | 2 | Hangzhou | 312,904,194 | 46,786,150,482 | 14.35 |
| middle | oolong tea | 2 | Hangzhou | 325,903,104 | 48,650,792,752 | 14.92 |
| middle | oolong tea | 2 | Hangzhou | 351,100,840 | 52,420,299,432 | 16.08 |
| late | green tea | 3 | Hangzhou | 316,360,432 | 47,258,589,250 | 14.50 |
| middle | oolong tea | 2 | Hangzhou | 326,037,286 | 48,669,931,570 | 14.93 |
| early | oolong tea | 2 | Hangzhou | 282,326,208 | 42,166,896,750 | 12.93 |

| | | | | | | |
|---|---|---|---|---|---|---|
| early | green tea | 2 | Hangzhou | 220,803,842 | 31,844,181,906 | 9.77 |
| early | black tea | 4 | Hangzhou | 343,033,490 | 51,245,896,906 | 15.72 |
| early | green tea | 4 | Hangzhou | 367,612,948 | 54,939,718,290 | 16.85 |
| early | black tea | 4 | Hangzhou | 307,732,342 | 45,992,717,890 | 14.11 |
| middle | black tea | 1 | Hangzhou | 349,127,924 | 52,113,946,450 | 15.99 |
| early | green tea | 2 | Hangzhou | 361,222,450 | 54,003,103,894 | 16.57 |
| early | green tea | 1 | Hangzhou | 319,014,074 | 47,660,607,586 | 14.62 |
| middle | green tea | 2 | Hangzhou | 337,579,884 | 48,640,803,563 | 14.92 |
| early | green tea | 1 | Hangzhou | 345,326,428 | 51,568,041,246 | 15.82 |
| extra early | green tea | 2 | Hangzhou | 269,587,330 | 39,001,745,611 | 11.96 |
| late | black tea | 2 | Hangzhou | 347,393,474 | 51,861,973,538 | 15.91 |
| early | black tea | 2 | Hangzhou | 376,885,364 | 56,312,131,010 | 17.27 |
| late | oolong tea | 2 | Hangzhou | 303,749,086 | 45,373,242,990 | 13.92 |
| extra early | oolong tea | 3 | Fuan | 408,418,066 | 61,066,516,512 | 18.73 |
| late | oolong tea | 1 | Fuan | 369,863,028 | 55,221,598,876 | 16.94 |
| late | oolong tea | 2 | Fuan | 305,267,616 | 45,630,086,926 | 14.00 |
| middle | oolong tea | 2 | Fuan | 383,816,294 | 57,370,092,076 | 17.60 |
| late | oolong tea | 2 | Fuan | 384,882,382 | 57,515,788,396 | 17.64 |
| early | green tea | 2 | Fuan | 372,339,874 | 55,577,962,128 | 17.05 |
| middle | black tea | 4 | Fuan | 277,864,804 | 40,132,141,116 | 12.31 |
| extra early | green tea | 2 | Hangzhou | 371,600,866 | 55,556,353,930 | 17.04 |
| - | green tea | 3 | Hangzhou | 349,016,160 | 52,019,627,462 | 15.96 |
| late | green tea | 4 | Hangzhou | 282,796,480 | 42,252,881,248 | 12.96 |
| early | black tea | 4 | Hangzhou | 307,203,508 | 45,787,030,688 | 14.05 |
| early | green tea | 1 | Hangzhou | 366,097,670 | 54,679,058,396 | 16.77 |
| early | green tea | 3 | Hangzhou | 268,264,366 | 38,551,626,022 | 11.83 |
| - | green tea | 3 | Hangzhou | 303,970,564 | 45,009,083,804 | 13.81 |
| middle | black tea | 3 | Hangzhou | 313,725,362 | 46,760,369,074 | 14.34 |
| early | green tea | 2 | Hangzhou | 344,325,986 | 51,450,457,508 | 15.78 |
| early | oolong tea | 4 | Hangzhou | 363,479,362 | 54,319,594,160 | 16.66 |

| | | | | | | | |
|---|---|---|---|---|---|---|---|
| late | oolong tea | | 2 | Hangzhou | 348,224,256 | 51,989,301,376 | 15.95 |
| late | green tea | | 3 | Hangzhou | 389,105,340 | 58,136,007,214 | 17.83 |
| early | black tea | | 1 | Hangzhou | 329,371,236 | 49,095,744,524 | 15.06 |
| middle | oolong tea | | 2 | Hangzhou | 361,003,502 | 53,468,592,270 | 16.40 |
| late | green tea | | 3 | Hangzhou | 336,884,094 | 50,206,304,100 | 15.40 |
| late | green tea | | 3 | Hangzhou | 312,269,000 | 46,481,933,238 | 14.26 |
| late | | | 4 | Hangzhou | 284,599,068 | 42,401,507,704 | 13.01 |
| - | - | | 3 | Hangzhou | 283,358,930 | 42,261,020,414 | 12.96 |
| - | - | | 4 | Hangzhou | 317,201,744 | 47,150,875,066 | 14.46 |
| late | black tea | | 4 | Menghai | 318,024,926 | 47,403,613,406 | 14.54 |
| - | black tea | | 4 | Menghai | 319,743,676 | 47,358,290,272 | 14.53 |
| - | green tea | | 4 | Menghai | 320,196,852 | 47,705,543,618 | 14.63 |
| - | black tea | | 4 | Menghai | 316,211,534 | 47,240,109,894 | 14.49 |
| - | green tea | | 4 | Menghai | 319,795,330 | 47,686,219,680 | 14.63 |
| - | black tea | | 4 | Menghai | 328,828,360 | 48,694,975,674 | 14.94 |
| middle | black tea | | 4 | Menghai | 327,627,504 | 48,836,971,896 | 14.98 |
| middle | black tea | | 4 | Menghai | 327,771,652 | 48,530,906,440 | 14.89 |
| - | black tea | | 4 | Menghai | 289,928,296 | 43,160,817,668 | 13.24 |
| - | black tea | | 4 | Menghai | 379,577,986 | 56,549,505,676 | 17.35 |
| - | black tea | | 4 | Menghai | 371,178,692 | 55,312,266,970 | 16.97 |
| - | black tea | | 4 | Menghai | 346,267,828 | 51,584,495,878 | 15.82 |
| - | black tea | | 4 | Menghai | 377,840,744 | 55,893,675,528 | 17.15 |
| - | black tea | | 4 | Hangzhou | 243,317,210 | 35,785,236,037 | 10.98 |
| - | black tea | | 4 | Hangzhou | 274,852,042 | 40,402,826,757 | 12.39 |
| - | black tea | | 4 | Hangzhou | 281,132,966 | 41,367,869,749 | 12.69 |
| - | black tea | | 4 | Hangzhou | 251,796,474 | 37,052,396,673 | 11.37 |
| - | black tea | | 4 | Hangzhou | 249,260,512 | 36,665,268,217 | 11.25 |
| - | green tea | - | | Hangzhou | 268,322,748 | 39,435,629,720 | 12.10 |
| - | green tea | - | | Hangzhou | 237,456,460 | 34,959,339,739 | 10.72 |
| - | green tea | - | | Hangzhou | 291,179,568 | 42,840,248,222 | 13.14 |

| | | | | | | |
|---|---|---|---|---|---|---|
| - | green tea | - | Hangzhou | 287,271,432 | 42,241,227,139 | 12.96 |
| - | green tea | - | Hangzhou | 267,156,592 | 39,298,148,591 | 12.05 |
| - | black tea | 2 | Hangzhou | 244,144,564 | 35,929,706,387 | 11.02 |
| - | black tea | 2 | Hangzhou | 265,628,504 | 39,074,224,025 | 11.99 |
| - | black tea | 2 | Hangzhou | 254,052,346 | 37,357,062,143 | 11.46 |
| - | black tea | 2 | Hangzhou | 263,468,456 | 38,744,572,500 | 11.88 |
| - | black tea | 4 | Hangzhou | 294,254,396 | 43,265,731,400 | 13.27 |
| - | black tea | 4 | Hangzhou | 269,417,526 | 39,622,943,579 | 12.15 |
| - | black tea | 4 | Hangzhou | 250,398,330 | 36,790,850,661 | 11.29 |
| - | black tea | 4 | Hangzhou | 257,513,012 | 37,861,143,898 | 11.61 |
| - | black tea | 4 | Hangzhou | 267,676,272 | 39,310,389,003 | 12.06 |
| - | green tea | 1 | Rizhao | 282,626,090 | 41,548,108,920 | 12.74 |
| - | green tea | 1 | Rizhao | 284,630,454 | 41,851,599,193 | 12.84 |
| - | green tea | 2 | Hangzhou | 242,609,892 | 35,691,571,121 | 10.95 |
| - | green tea | 2 | Hangzhou | 245,286,394 | 36,088,120,310 | 11.07 |
| - | green tea | 2 | Hangzhou | 266,325,398 | 39,174,481,138 | 12.02 |
| - | green tea | 2 | Hangzhou | 300,999,018 | 44,285,701,275 | 13.58 |
| - | black tea | 4 | Hangzhou | 297,849,132 | 43,831,422,828 | 13.45 |
| - | green tea | 2 | Hangzhou | 269,286,194 | 39,618,606,669 | 12.15 |
| - | black tea | 4 | Hangzhou | 229,015,414 | 33,688,882,220 | 10.33 |
| - | black tea | 4 | Hangzhou | 276,613,826 | 40,683,892,474 | 12.48 |
| - | green tea | 2 | Hangzhou | 248,664,070 | 36,582,720,327 | 11.22 |
| - | black tea | 4 | Hangzhou | 224,000,636 | 32,922,341,253 | 10.10 |
| - | green tea | 1 | Hangzhou | 296,330,994 | 43,580,995,941 | 13.37 |
| - | green tea | 2 | Hangzhou | 277,939,032 | 40,923,844,510 | 12.55 |
| - | green tea | 2 | Hangzhou | 273,975,934 | 40,324,963,934 | 12.37 |
| - | green tea | 2 | Hangzhou | 267,450,652 | 39,383,240,629 | 12.08 |
| - | green tea | 2 | Hangzhou | 292,282,568 | 43,033,590,048 | 13.20 |
| - | black tea | 4 | Hangzhou | 258,041,314 | 37,994,920,230 | 11.66 |
| - | black tea | 4 | Hangzhou | 262,052,726 | 38,574,170,485 | 11.83 |

| | | | | | | | |
|---|---|---|---|---|---|---|---|
| - | black tea | | 4 | Hangzhou | 290,072,990 | 42,707,090,822 | 13.10 |
| - | black tea | | 4 | Hangzhou | 246,461,662 | 36,272,628,788 | 11.13 |
| middle | oolong tea | - | | Hangzhou | 259,664,782 | 38,228,250,099 | 11.73 |
| middle | oolong tea | - | | Hangzhou | 278,611,068 | 41,016,563,690 | 12.58 |
| - | black tea | | 4 | Hangzhou | 246,889,582 | 35,477,247,329 | 10.88 |
| - | green tea | | 2 | Hangzhou | 251,536,422 | 36,911,827,481 | 11.32 |
| - | black tea | | 4 | Hangzhou | 244,857,494 | 35,914,025,400 | 11.02 |
| - | black tea | | 4 | Hangzhou | 260,560,642 | 38,230,104,679 | 11.73 |
| - | green tea | | 3 | Hangzhou | 258,758,098 | 37,949,039,327 | 11.64 |
| - | green tea | | 4 | Hangzhou | 329,368,212 | 48,312,845,027 | 14.82 |
| - | green tea | | 4 | Hangzhou | 267,195,848 | 39,185,334,700 | 12.02 |
| - | black tea | | 4 | Hangzhou | 300,877,112 | 44,130,510,706 | 13.54 |
| - | green tea | | 4 | Hangzhou | 308,269,998 | 45,199,216,249 | 13.86 |
| - | green tea | | 1 | Hangzhou | 284,773,486 | 41,833,677,572 | 12.83 |
| - | green tea | | 4 | Hangzhou | 304,521,042 | 44,709,011,431 | 13.71 |
| - | green tea | | 2 | Hangzhou | 302,134,810 | 44,358,780,196 | 13.61 |
| - | green tea | | 2 | Hangzhou | 273,099,340 | 40,115,439,689 | 12.31 |
| - | black tea | | 4 | Hangzhou | 266,221,426 | 39,096,567,674 | 11.99 |
| - | green tea | | 2 | Hangzhou | 304,885,442 | 44,784,405,114 | 13.74 |
| - | - | - | | Hangzhou | 328,208,488 | 47,964,816,618 | 14.71 |
| early | black tea | | 4 | Hangzhou | 322,400,978 | 47,343,237,982 | 14.52 |
| early | black tea | | 4 | Hangzhou | 300,889,328 | 44,221,595,906 | 13.57 |
| early | black tea | | 4 | Yingde | 277,201,524 | 40,511,160,584 | 12.43 |
| early | black tea | | 4 | Yingde | 237,805,208 | 34,764,826,634 | 10.66 |

| Mapping ratio | mapping rate | Heterozygosity |
|---|---|---|
| 99.03% | 64.13% | 0.00699579 |
| 99.30% | 69.75% | 0.00639879 |
| 99.15% | 72.43% | 0.00528031 |
| 98.93% | 71.54% | 0.0041217 |
| 97.38% | 70.71% | 0.00493481 |
| 99.15% | 71.60% | 0.0053158 |
| 99.42% | 70.00% | 0.00559139 |
| 97.33% | 69.60% | 0.00515496 |
| 97.88% | 71.34% | 0.0049893 |
| 99.52% | 76.28% | 0.00569506 |
| 99.17% | 69.95% | 0.00566996 |
| 98.63% | 66.12% | 0.00587981 |
| 99.28% | 70.83% | 0.00477937 |
| 99.04% | 72.06% | 0.00507368 |
| 98.52% | 70.96% | 0.00536081 |
| 99.23% | 66.72% | 0.00466801 |
| 99.36% | 72.00% | 0.00577968 |
| 98.63% | 62.16% | 0.00734471 |
| 99.41% | 77.90% | 0.00497362 |
| 99.28% | 72.55% | 0.00546414 |
| 99.13% | 68.59% | 0.00524222 |
| 98.81% | 70.41% | 0.0053625 |
| 98.70% | 70.24% | 0.00566152 |
| 98.32% | 64.57% | 0.00474689 |
| 97.95% | 70.83% | 0.0075656 |
| 98.88% | 70.03% | 0.00510511 |

| | | |
|---|---|---|
| 99.65% | 65.56% | 0.00316521 |
| 99.10% | 66.94% | 0.00753748 |
| 98.45% | 67.87% | 0.00565311 |
| 99.29% | 71.79% | 0.0061897 |
| 97.50% | 69.17% | 0.00523871 |
| 99.18% | 66.86% | 0.00651164 |
| 96.95% | 64.73% | 0.00607626 |
| 99.54% | 60.40% | 0.0040617 |
| 97.92% | 67.86% | 0.00534374 |
| 99.56% | 65.55% | 0.00450992 |
| 97.36% | 65.18% | 0.00658965 |
| 98.65% | 67.53% | 0.00768552 |
| 98.92% | 70.78% | 0.00567122 |
| 98.83% | 68.51% | 0.00647882 |
| 98.23% | 68.87% | 0.00613836 |
| 99.51% | 72.20% | 0.00546126 |
| 98.52% | 69.12% | 0.00571444 |
| 97.80% | 69.45% | 0.00585929 |
| 98.43% | 70.39% | 0.00610155 |
| 99.27% | 60.59% | 0.00503411 |
| 99.16% | 70.96% | 0.00650714 |
| 99.39% | 67.10% | 0.00545066 |
| 99.38% | 66.53% | 0.00683353 |
| 99.29% | 66.87% | 0.00651437 |
| 97.91% | 71.46% | 0.00508533 |
| 99.66% | 59.29% | 0.00496288 |
| 99.26% | 68.92% | 0.00666985 |
| 99.06% | 67.08% | 0.00610439 |
| 98.08% | 68.72% | 0.00586668 |
| 97.41% | 67.20% | 0.00559671 |

| | | |
|---|---|---|
| 98.34% | 71.91% | 0.00526167 |
| 96.98% | 63.28% | 0.00646489 |
| 99.24% | 69.62% | 0.00549738 |
| 98.93% | 71.42% | 0.00594396 |
| 99.04% | 68.41% | 0.00525825 |
| 99.19% | 64.97% | 0.00688611 |
| 99.09% | 64.58% | 0.00450867 |
| 99.03% | 55.65% | 0.00356742 |
| 98.78% | 57.24% | 0.00346362 |
| 98.83% | 62.01% | 0.0051469 |
| 99.29% | 64.08% | 0.00417516 |
| 99.18% | 63.71% | 0.00425199 |
| 99.34% | 61.32% | 0.00714418 |
| 99.25% | 69.75% | 0.00551505 |
| 99.35% | 60.91% | 0.00652333 |
| 98.73% | 64.72% | 0.00580641 |
| 98.31% | 61.77% | 0.00446506 |
| 99.29% | 64.02% | 0.00792963 |
| 99.17% | 63.46% | 0.00536305 |
| 99.34% | 64.19% | 0.00529357 |
| 99.37% | 62.83% | 0.00398965 |
| 98.99% | 58.45% | 0.00718382 |
| 99.59% | 69.21% | 0.00580123 |
| 99.52% | 69.27% | 0.00514486 |
| 99.55% | 69.49% | 0.00699716 |
| 99.54% | 68.96% | 0.0055635 |
| 99.53% | 70.18% | 0.00605682 |
| 99.36% | 69.87% | 0.00495725 |
| 99.56% | 71.41% | 0.00551898 |
| 99.59% | 72.38% | 0.00494698 |

| | | |
|---|---|---|
| 99.63% | 72.94% | 0.00530854 |
| 99.57% | 72.24% | 0.00605179 |
| 99.60% | 72.59% | 0.00548724 |
| 99.59% | 72.14% | 0.00577756 |
| 99.56% | 71.86% | 0.00545354 |
| 99.57% | 72.46% | 0.00514481 |
| 99.54% | 66.26% | 0.00468863 |
| 99.59% | 65.42% | 0.00475426 |
| 99.55% | 66.04% | 0.00462138 |
| 99.53% | 66.41% | 0.0048471 |
| 99.49% | 65.94% | 0.00430429 |
| 99.53% | 72.45% | 0.00826263 |
| 99.47% | 71.25% | 0.00702415 |
| 99.51% | 71.70% | 0.00567871 |
| 99.55% | 73.50% | 0.00512539 |
| 99.60% | 74.00% | 0.00415095 |
| 99.59% | 73.58% | 0.00506754 |
| 99.58% | 67.63% | 0.00677335 |
| 99.46% | 71.32% | 0.00589339 |
| 99.56% | 67.35% | 0.00506837 |
| 99.54% | 66.69% | 0.00591091 |
| 99.59% | 72.65% | 0.00487917 |
| 99.53% | 65.88% | 0.00577489 |
| 98.86% | 71.78% | 0.00510445 |
| 99.59% | 73.38% | 0.00557866 |
| 99.56% | 73.34% | 0.00490714 |
| 99.62% | 73.77% | 0.00499147 |
| 99.41% | 73.66% | 0.00426862 |
| 99.55% | 66.53% | 0.0059154 |
| 99.53% | 67.16% | 0.00606055 |

| | | |
|---|---|---|
| 99.56% | 68.13% | 0.00609811 |
| 99.55% | 65.45% | 0.00406455 |
| 99.56% | 72.60% | 0.0047408 |
| 99.58% | 72.56% | 0.0049164 |
| 99.33% | 73.64% | 0.00562628 |
| 99.13% | 72.93% | 0.00722143 |
| 97.99% | 66.71% | 0.00673293 |
| 98.56% | 67.79% | 0.00633216 |
| 99.66% | 71.00% | 0.00572322 |
| 98.91% | 65.20% | 0.0081459 |
| 99.46% | 65.23% | 0.00696015 |
| 98.81% | 68.53% | 0.00641772 |
| 99.22% | 64.67% | 0.00743851 |
| 99.15% | 73.30% | 0.00724703 |
| 99.19% | 69.82% | 0.00667459 |
| 99.38% | 73.33% | 0.00750356 |
| 99.38% | 72.92% | 0.00510978 |
| 99.53% | 67.62% | 0.00657587 |
| 99.14% | 72.71% | 0.00562825 |
| 98.39% | 46.85% | - |
| 99.36% | 65.38% | 0.00655854 |
| 99.57% | 68.98% | 0.00752249 |
| 99.31% | 64.67% | 0.00409788 |
| 99.37% | 64.87% | 0.00373143 |

| Supplementary Table 19: The selection genes in CSS |
| --- |
| chr2.1198 |
| chr2.1199 |
| chr2.1200 |
| chr7.1284 |
| chr2.1615 |
| chr2.1616 |
| chr12.777 |
| chr9.1718 |
| chr9.1719 |
| chr11.1155 |
| chr11.1156 |
| chr2.571 |
| chr4.48 |
| chr6.410 |
| chr11.241 |
| chr11.242 |
| chr11.1409 |
| chr5.925 |
| chr5.1665 |
| chr8.583 |
| chr10.244 |
| chr9.898 |
| chr3.522 |
| chr3.2096 |
| chr10.1533 |
| chr13.546 |
| chr8.1666 |
| chr14.781 |
| chr11.361 |
| chr14.780 |
| chr9.197 |
| chr1.551 |
| chr11.527 |
| chr5.554 |
| chr5.505 |
| chr10.587 |
| chr7.145 |
| chr1.2453 |
| chr1.1180 |
| chr7.381 |
| chr2.649 |
| chr2.650 |
| chr15.993 |
| chr6.1008 |
| chr5.1061 |
| chr1.1286 |
| chr10.677 |
| chr7.924 |
| chr7.925 |

chr9.660
chr9.661
chr7.1133
chr5.1232
chr11.1168
chr7.410
chr9.1244
chr3.1336
chr9.543
chr7.1189
chr10.141
chr2.463
chr6.1547
chr1.1501
chr13.264
chr8.1664
chr7.693
chr3.1761
chr2.1050
chr3.1946
chr6.1110
chr3.707
chr5.982
chr3.227
chr4.2077
chr1.660
chr4.2166
chr9.1387
chr6.456
chr6.1280
chr4.1156
chr5.1238
chr7.1472
chr6.2090
chr2.256
chr12.1056
chr7.979
chr7.758
chr10.427
chr2.513
chr2.259
chr2.260
chr1.2079
chr6.953
chr10.925
chr4.851
chr6.1822
chr8.1550
chr10.1379
chr10.1040

chr6.737
chr15.593
chr3.658
chr8.186
chr11.278
chr1.1382
chr10.1459
chr11.1066
chr2.1461
chr4.2054
chr5.1527
chr5.1528
chr7.371
chr14.604
chr10.62
chr7.1000
chr2.161
chr10.345
chr7.612
chr10.231
chr2.1578
chr9.1344
chr15.480
chr13.764
chr2.1906
chr2.1176
chr13.203
chr3.1973
chr3.1974
chr7.622
chr7.623
chr3.1486
chr8.1352
chr3.2088
chr11.453
chr2.1652
chr14.1029
chr10.1143
chr1.1167
chr10.202
chr5.693
chr6.1806
chr10.1085
chr11.1441
chr13.1311
chr12.700
chr2.1051
chr10.459
chr5.1499
chr9.1363

chr13.345
chr13.705
chr1.950
chr10.1043
chr6.1087
chr15.113
chr8.1616
chr5.216
chr13.1395
chr3.469
chr9.1267
chr3.755
chr10.82
chr10.344
chr6.2012
chr6.1369
chr15.374
chr5.1619
chr2.668
chr12.1017
chr14.1244
chr14.856
chr13.1101
chr15.199
chr10.11
chr7.1034
chr8.1435
chr14.143
chr1.203
chr6.209
chr8.387
chr8.648
chr12.1294
chr6.179
chr7.868
chr8.572
chr1.704
chr6.1390
chr1.480
chr13.709
chr2.1554
chr1.2286
chr8.1014
chr2.716
chr3.1415
chr5.68
chr13.1041
chr4.1867
chr3.1843
chr6.1927

chr13.235
chr5.280
chr12.1076
chr2.412
chr3.915
chr6.1534
chr5.41
chr2.324
chr10.659
chr4.1894
chr15.1237
chr5.277
chr12.299
chr5.908
chr7.628
chr2.1478
chr4.1709
chr13.610
chr9.98
chr13.1102
chr3.408
chr12.1214
chr9.1688
chr1.639
chr7.375
chr2.1576
chr7.1340
chr11.1285
chr2.933
chr9.580
chr10.891
chr11.1273
chr2.712
chr7.255
chr13.1080
chr2.616
chr10.259
chr2.1862
chr3.1011
chr4.495
chr11.107
chr1.1054
chr14.238
chr5.521
chr1.1564
chr2.381
chr6.608
chr2.1158
chr2.1357
chr7.1338

chr15.588
chr5.1521
chr5.1522
chr10.265
chr8.895
chr6.435
chr2.1836
chr14.1132
chr13.85
chr13.630
chr13.770
chr3.1802
chr11.202
chr5.799
chr9.1002
chr4.1333
chr5.319
chr4.937
chr14.325
chr4.1681
chr9.542
chr5.259
chr1.2116
chr14.1318
chr2.1440
chr4.1522
chr9.1575
chr8.404
chr13.136
chr8.1587
chr5.1355
chr5.322
chr2.1876
chr13.1010
chr11.440
chr8.1138
chr11.751
chr9.1777
chr12.825
chr5.608
chr7.980
chr10.979
chr3.1392
chr7.983
chr11.1382
chr9.1759
chr10.277
chr2.382
chr9.1112
chr7.278

chr1.1913
chr15.306
chr12.364
chr10.955
chr5.1642
chr15.747
chr9.325
chr9.1359
chr6.581
chr1.1122
chr9.855
chr2.1168
chr8.1593
chr8.1755
chr6.1480
chr4.2108
chr13.629
chr13.292
chr3.756
chr2.1868
chr7.277
chr13.1305
chr13.663
chr11.161
chr14.130
chr3.496
chr2.1684
chr13.226
chr10.1146
chr1.1867
chr5.611
chr9.1105
chr1.680
chr15.583
chr11.817
chr1.1966
chr8.1896
chr2.769
chr6.884
chr3.326
chr3.91
chr8.1583
chr15.768
chr6.1301
chr10.666
chr14.886
chr11.1052
chr10.447
chr4.1334
chr6.1360

chr15.455
chr11.836
chr2.1801
chr9.821
chr7.942
chr5.31
chr10.546
chr12.623
chr7.1393
chr7.563
chr7.1254
chr1.954
chr4.926
chr1.1407
chr15.888
chr6.1330
chr11.1105
chr5.1340
chr4.1594
chr7.1528
chr6.1464
chr3.17
chr9.441
chr9.1005
chr6.1532
chr6.1001
chr9.613
chr3.902
chr11.186
chr12.642
chr10.185
chr7.433
chr15.399
chr12.438
chr15.781
chr3.616
chr9.1808
chr1.1585
chr3.879
chr3.751
chr6.1900
chr10.1438
chr6.272
chr13.1150
chr10.947
chr13.137
chr8.945
chr2.662
chr8.1296
chr2.1096

chr1.739
chr7.1060
chr10.989
chr14.727
chr6.275
chr4.1587
chr3.818
chr7.913
chr11.145
chr5.515
chr2.524
chr11.282
chr5.859
chr11.252
chr8.1408
chr1.1189
chr1.228
chr9.985
chr6.1562
chr7.51
chr3.1798
chr9.492
chr6.854
chr5.1485
chr11.321
chr12.295
chr10.290
chr3.1332
chr3.1311
chr12.977
chr11.256
chr7.235
chr13.763
chr10.348
chr10.529
chr3.480
chr9.1102
chr7.1285
chr9.653
chr8.351
chr2.683
chr1.1022
chr10.280
chr6.1933
chr13.1163
chr8.1448
chr3.1659
chr3.80
chr3.505
chr5.448

chr15.196
chr1.2416
chr1.1306
chr15.380
chr12.1304
chr11.1158
chr3.768
chr4.312
chr5.702
chr10.695
chr14.965
chr8.746
chr7.1339
chr13.336
chr14.858
chr4.2109
chr4.701
chr2.1463
chr1.1173
chr1.691
chr9.1485
chr1.561
chr14.392
chr5.1382
chr3.33
chr11.372
chr1.2535
chr10.72
chr13.1417
chr9.391
chr3.2005
chr12.152
chr6.726
chr4.505
chr11.127
chr13.297
chr3.512
chr1.209
chr12.868
chr6.1832
chr12.1196
chr3.1055
chr8.897
chr3.364
chr14.263
chr1.388
chr8.167
chr1.3
chr7.950
chr6.1345

chr15.687
chr10.1274
chr3.2206
chr7.168
chr12.1217
chr1.2447
chr2.903
chr6.755
chr12.239
chr6.1821
chr7.943
chr3.1140
chr11.281
chr2.940
chr5.270
chr9.436
chr11.1063
chr13.115
chr9.977
chr11.1404
chr13.79
chr8.1385
chr1.681
chr2.482
chr3.2086
chr4.1258
chr8.1449
chr15.589
chr8.1791
chr4.1048
chr7.603
chr4.93
chr10.655
chr10.766
chr2.1812
chr6.109
chr10.340
chr3.45
chr11.59
chr14.635
chr1.2503
chr11.524
chr6.656
chr2.1094
chr1.901
chr2.764
chr10.927
chr10.1082
chr2.1138
chr1.2560

chr6.448
chr15.163
chr13.1171
chr2.713
chr8.121
chr11.912
chr5.895
chr2.77
chr5.707
chr15.391
chr3.324
chr1.2546
chr7.387
chr15.926
chr6.49
chr8.436
chr4.401
chr13.715
chr8.1528
chr6.246
chr14.1397
chr1.1654
chr13.547
chr8.305
chr3.216
chr4.450
chr15.282
chr6.1492
chr2.1678
chr14.105
chr5.165
chr12.532
chr2.833
chr11.816
chr12.1301
chr10.834
chr5.933
chr9.119
chr1.1487
chr13.668
chr15.76
chr9.1641
chr6.1463
chr1.935
chr13.370
chr7.1546
chr15.855
chr13.8
chr15.70
chr8.954

chr15.1149
chr3.2110
chr6.1546
chr3.1257
chr10.685
chr15.1062
chr11.1154
chr6.1880
chr10.432
chr5.764
chr7.272
chr13.1031
chr12.242
chr5.974
chr12.1146
chr4.1805
chr2.278
chr9.1707
chr12.1317
chr13.634
chr5.1455
chr4.2123
chr6.417
chr15.180
chr2.328
chr8.782
chr5.1479
chr5.4
chr1.1268
chr4.879
chr5.1686
chr9.329
chr15.382
chr12.915
chr6.605
chr7.1075
chr10.74
chr11.629
chr13.699
chr3.1409
chr4.563
chr9.433
chr15.375
chr6.1745
chr13.1318
chr1.1156
chr15.1040
chr6.449
chr2.717
chr1.1439

chr1.2466
chr13.707
chr2.1388
chr10.402
chr10.122
chr7.984
chr6.837
chr2.1720
chr2.211
chr5.432
chr10.232
chr2.1613
chr5.1565
chr2.733
chr5.401
chr2.1902
chr3.1743
chr10.978
chr3.426
chr3.113
chr14.93
chr7.719
chr13.301
chr5.1673
chr4.2085
chr7.698
chr2.784
chr12.591
chr4.642
chr6.119
chr6.122
chr13.990
chr5.435
chr10.1201
chr9.1422
chr12.1309
chr4.501
chr10.125
chr8.1731
chr8.1315
chr9.1513
chr9.707
chr2.1584
chr2.1585
chr3.1348
chr5.1154
chr4.1595
chr9.1138
chr5.911
chr11.684

chr7.1344
chr3.252
chr11.956
chr11.1438
chr5.1334
chr2.512
chr15.1093
chr7.778
chr11.105
chr1.2488
chr13.788
chr11.1447
chr4.1021
chr10.1305
chr9.407
chr4.1340
chr4.1796
chr9.996
chr5.1040
chr1.1602
chr9.359
chr5.293
chr9.425
chr9.666
chr8.656
chr8.38
chr11.436
chr7.783
chr4.1730
chr2.367
chr5.1367
chr4.1458
chr6.1727
chr3.336
chr12.169
chr1.129
chr2.1875
chr13.586
chr11.932
chr15.25
chr3.418
chr7.1231
chr13.1391
chr9.361
chr1.2435
chr7.933
chr6.313
chr6.870
chr2.176
chr7.891

chr9.346
chr12.106
chr9.602
chr2.1413
chr5.469
chr15.310
chr15.711
chr2.924
chr8.632
chr2.912
chr4.2135
chr3.334
chr3.1526
chr14.772
chr2.590
chr9.14
chr2.771
chr1.1763
chr3.940
chr2.750
chr1.787
chr9.623
chr13.407
chr8.654
chr6.973
chr3.2119
chr11.508
chr10.419
chr12.501
chr5.775
chr5.55
chr12.189
chr14.1262
chr8.396
chr6.249
chr9.1447
chr10.139
chr5.1205
chr6.859
chr11.974
chr1.825
chr10.1427
chr15.547
chr5.258
chr3.2100
chr5.665
chr5.1630
chr6.746
chr14.578
chr5.528

chr3.411
chr6.1524
chr1.1702
chr13.996
chr1.324
chr13.1390
chr12.744
chr2.920
chr10.409
chr15.130
chr1.457
chr10.853
chr12.1057
chr1.386
chr2.133
chr2.1307
chr15.920
chr4.927
chr11.1481
chr3.2205
chr8.770
chr5.1477
chr15.166
chr4.853
chr13.814
chr15.872
chr5.771
chr9.5
chr2.488
chr5.1274
chr2.1878
chr4.660
chr5.927
chr5.800
chr10.1031
chr3.1736
chr5.189
chr3.916
chr4.1917
chr3.955
chr11.1434
chr1.2142
chr14.330
chr13.402
chr4.514
chr6.1998
chr1.653
chr14.51
chr2.342
chr15.41

chr3.74
chr8.102
chr5.290
chr2.1172
chr10.1368
chr15.214
chr13.1412
chr6.2088
chr12.1130
chr5.1048
chr4.2268
chr14.490
chr1.2516
chr3.297
chr9.1058
chr8.105
chr1.677
chr4.433
chr14.1061
chr3.43
chr6.186
chr11.705
chr5.680
chr13.1341
chr14.1072
chr9.282
chr3.449
chr5.359
chr5.1164
chr4.1384
chr6.745
chr1.451
chr3.1157
chr13.992
chr13.795
chr13.926
chr6.1359
chr5.272
chr8.573
chr11.1304
chr8.1167
chr6.1604
chr11.512
chr6.609
chr14.1093
chr11.421
chr6.300
chr2.1625
chr2.1480
chr8.519

chr10.829
chr15.72
chr4.1134
chr1.1760
chr1.1688
chr3.2011
chr4.422
chr10.550
chr10.648
chr14.438
chr2.1345
chr13.53
chr8.693
chr4.1136
chr1.2502
chr2.1592
chr15.493
chr4.253
chr2.355
chr6.399
chr11.1179
chr9.59
chr7.1328
chr2.102
chr1.2298
chr7.1367
chr6.499
chr4.889
chr8.82
chr11.208
chr4.588
chr12.1033
chr9.1673
chr14.1257
chr10.166
chr14.1358
chr5.1687
chr8.633
chr13.704
chr1.2299
chr15.171
chr6.1992
chr1.2493
chr10.952
chr13.889
chr11.1025
chr12.840
chr9.501
chr8.851
chr11.1359

chr2.628
chr8.1411
chr2.1243
chr2.1647
chr12.1190
chr14.1228
chr9.1614
chr3.97
chr6.1227
chr6.1895
chr5.66
chr7.1112
chr6.1677
chr8.1466
chr12.1193
chr7.335
chr11.26
chr3.1376
chr15.814
chr15.1005
chr12.944
chr8.381
chr5.237
chr6.1179
chr6.553
chr13.1114
chr12.220
chr1.2300
chr1.333
chr7.1348
chr5.1461
chr12.312
chr4.2173
chr13.679
chr13.1410
chr9.217
chr2.1921
chr4.2061
chr1.1577
chr14.183
chr3.208
chr9.1706
chr2.103
chr8.1016
chr5.48
chr4.1715
chr4.1124
chr13.352
chr1.919
chr10.580

chr14.502
chr15.999
chr1.1124
chr11.1350
chr5.115
chr8.424
chr12.822
chr10.907
chr1.482
chr8.981
chr15.622
chr15.1180
chr9.750
chr1.822
chr9.802
chr2.1489
chr3.799
chr10.953
chr5.1441
chr6.1907
chr8.1754
chr2.1048
chr1.1485
chr2.457
chr15.898
chr13.1262
chr13.1157
chr13.887
Cha5.766

| Supplementary Table 20: The selection genes in CSA |
|---|
| chr9.1726 |
| chr9.1727 |
| chr9.1728 |
| chr9.1729 |
| chr5.800 |
| chr5.624 |
| chr5.1339 |
| chr9.803 |
| chr9.804 |
| chr9.522 |
| chr9.1582 |
| chr15.431 |
| chr14.1126 |
| chr7.437 |
| chr8.1435 |
| chr9.1543 |
| chr9.1608 |
| chr10.1533 |
| chr9.660 |
| chr9.661 |
| chr9.662 |
| chr12.466 |
| chr15.976 |
| chr15.1032 |
| chr12.1225 |
| chr5.134 |
| chr9.903 |
| chr8.1102 |
| chr9.1813 |
| chr9.217 |
| chr9.218 |
| chr14.1132 |
| chr12.266 |
| chr6.179 |
| chr9.1670 |
| chr5.71 |
| chr9.1795 |
| chr8.186 |
| chr15.1188 |
| chr9.1531 |
| chr9.1532 |
| chr9.1533 |
| chr15.877 |
| chr9.1600 |
| chr9.1601 |
| chr8.276 |
| chr9.651 |
| chr1.2189 |
| chr9.1575 |

chr15.838
chr5.178
chr8.274
chr9.195
chr5.1665
chr5.1565
chr8.1284
chr5.1238
chr12.915
chr2.1357
chr1.46
chr9.359
chr5.375
chr11.36
chr5.189
chr6.1207
chr9.98
chr15.480
chr9.584
chr8.1589
chr14.17
chr13.1341
chr11.242
chr2.1525
chr12.873
chr8.1246
chr9.856
chr15.1031
chr15.929
chr7.457
chr10.1005
chr15.996
chr2.764
chr15.1173
chr3.2113
chr8.567
chr5.505
chr6.77
chr8.1384
chr8.238
chr10.17
chr8.1849
chr3.2096
chr14.1403
chr9.1398
chr5.708
chr10.675
chr12.744
chr8.422
chr8.167

chr8.1779
chr13.675
chr5.38
chr9.1771
chr8.1526
chr8.1294
chr6.1986
chr6.1345
chr5.398
chr8.1518
chr9.497
chr5.1521
chr5.1522
chr9.629
chr14.392
chr14.393
chr15.902
chr9.904
chr8.85
chr2.1047
chr9.275
chr9.1042
chr5.1154
chr12.1160
chr13.235
chr15.1030
chr5.982
chr10.174
chr2.259
chr2.260
chr5.26
chr10.348
chr10.74
chr15.1079
chr7.652
chr5.1355
chr13.510
chr11.1409
chr9.240
chr9.439
chr13.1076
chr8.235
chr8.1896
chr5.344
chr8.824
chr8.1845
chr8.715
chr8.1434
chr8.193
chr4.358

chr14.325
chr9.1367
chr8.1066
chr7.1393
chr14.733
chr1.525
chr1.950
chr5.280
chr7.980
chr2.1749
chr2.1417
chr8.1276
chr7.1551
chr5.48
chr10.580
chr6.688
chr5.881
chr10.1043
chr2.382
chr6.1999
chr3.799
chr9.436
chr1.1796
chr9.1629
chr13.878
chr6.1838
chr9.58
chr3.1761
chr5.383
chr15.985
chr10.11
chr6.1463
chr2.1659
chr7.467
chr3.512
chr9.1777
chr6.243
chr10.933
chr13.1391
chr10.801
chr10.404
chr8.28
chr4.1441
chr9.498
chr12.424
chr12.916
chr1.551
chr15.1119
chr15.954
chr12.54

chr12.55
chr8.168
chr13.764
chr5.229
chr8.842
chr5.268
chr1.993
chr7.313
chr7.381
chr9.1662
chr8.264
chr2.1883
chr12.105
chr9.1599
chr9.1266
chr13.8
chr8.945
chr6.517
chr8.694
chr15.986
chr5.852
chr9.1514
chr5.1490
chr1.1757
chr15.135
chr5.1485
chr14.132
chr11.1119
chr9.613
chr9.1377
chr15.686
chr9.1749
chr15.1178
chr8.1291
chr5.1152
chr2.784
chr2.252
chr14.1028
chr9.943
chr15.473
chr15.1094
chr15.872
chr13.642
chr9.1607
chr1.1119
chr10.1048
chr5.30
chr1.1121
chr8.509
chr8.484

chr8.205
chr2.616
chr3.968
chr8.1016
chr7.979
chr3.483
chr2.1199
chr12.415
chr9.1674
chr13.715
chr4.370
chr5.234
chr7.973
chr2.10
chr9.1138
chr8.1751
chr11.202
chr8.424
chr15.418
chr9.1707
chr6.1179
chr8.640
chr10.373
chr9.1169
chr10.620
chr11.59
chr2.1388
chr8.1488
chr5.528
chr9.1657
chr3.803
chr15.1062
chr9.1328
chr6.272
chr13.1390
chr8.1524
chr2.1131
chr14.175
chr2.1942
chr10.1003
chr11.402
chr4.664
chr9.1290
chr6.1369
chr10.980
chr13.908
chr5.43
chr2.1043
chr6.1534
chr13.1388

chr10.1040
chr2.185
chr14.841
chr9.547
chr15.399
chr15.400
chr11.1155
chr9.537
chr14.1228
chr8.1886
chr9.273
chr9.1239
chr9.1240
chr5.222
chr2.833
chr9.1151
chr9.1544
chr3.673
chr14.462
chr1.954
chr5.693
chr8.1806
chr7.24
chr9.1267
chr6.109
chr7.1321
chr1.325
chr5.1006
chr2.992
chr8.1659
chr10.62
chr4.1681
chr9.653
chr6.594
chr1.786
chr9.1480
chr9.525
chr13.1097
chr15.239
chr9.589
chr9.743
chr9.1344
chr8.895
chr8.896
chr2.1068
chr9.1668
chr8.1505
chr10.184
chr9.1650
chr2.1614

chr11.200
chr9.1736
chr8.57
chr3.522
chr11.985
chr7.889
chr9.1534
chr5.766
chr8.1293
chr12.426
chr15.438
chr15.876
chr1.482
chr15.1190
chr11.1406
chr9.1375
chr9.1074
chr15.656
chr1.697
chr1.327
chr13.517
chr5.682
chr9.1145
chr7.569
chr14.238
chr10.586
chr12.1270
chr9.811
chr14.130
chr5.545
chr8.1750
chr2.828
chr2.1585
chr14.1020
chr2.1903
chr8.1534
chr12.621
chr5.737
chr13.629
chr15.1068
chr2.1720
chr4.571
chr9.1397
chr15.1149
chr1.218
chr2.1653
chr5.1668
chr10.543
chr9.1434
chr15.994

chr7.704
chr7.264
chr12.61
chr9.591
chr9.96
chr8.318
chr14.1411
chr9.731
chr5.1636
chr12.702
chr15.1090
chr13.971
chr4.323
chr5.9
chr5.586
chr15.68
chr2.989
chr2.577
chr13.668
chr2.129
chr9.1642
chr13.763
chr8.230
chr9.229
chr15.955
chr14.1412
chr10.1191
chr1.1763
chr15.1080
chr14.1230
chr8.397
chr5.290
chr5.1356
chr9.248
chr5.680
chr9.1529
chr13.7
chr2.1480
chr10.1218
chr6.1540
chr14.1235
chr5.158
chr9.389
chr7.276
chr4.529
chr5.1171
chr2.821
chr14.273
chr6.619
chr9.1628

chr9.1444
chr3.1013
chr15.940
chr3.342
chr5.155
chr12.760
chr6.1624
chr13.1041
chr2.1573
chr5.1334
chr3.1680
chr8.1323
chr7.1418
chr5.1635
chr12.510
chr7.622
chr8.893
chr2.230
chr5.1024
chr10.105
chr8.833
chr5.53
chr8.434
chr15.1165
chr2.1065
chr8.14
chr4.197
chr3.1333
chr9.1710
chr2.955
chr2.1229
chr8.1374
chr9.1616
chr4.10
chr10.590
chr10.841
chr5.1676
chr5.777
chr12.688
chr8.1077
chr8.1237
chr2.574
chr5.19
chr15.183
chr2.683
chr15.426
chr13.449
chr15.757
chr9.935
chr3.2205

chr8.190
chr8.877
chr12.1075
chr14.542
chr9.868
chr15.1018
chr8.2
chr4.622
chr8.1672
chr9.7
chr12.619
chr8.1289
chr9.1300
chr8.1247
chr9.1120
chr9.1606
chr2.353
chr5.130
chr14.482
chr14.885
chr9.541
chr5.306
chr15.1216
chr7.535
chr8.189
chr10.1274
chr9.567
chr8.1457
chr15.642
chr9.586
chr3.1619
chr15.1127
chr15.1012
chr2.987
chr12.1317
chr14.61
chr5.639
chr2.267
chr8.720
chr7.742
chr9.1481
chr9.1462
chr3.547
chr10.721
chr8.1380
chr14.136
chr11.1416
chr6.783
chr8.1285
chr6.1492

chr9.942
chr8.208
chr2.133
chr12.1246
chr14.1107
chr5.185
chr8.950
chr9.1374
chr6.2016
chr8.147
chr8.899
chr15.189
chr14.1304
chr5.515
chr9.1401
chr15.726
chr1.359
chr10.611
chr13.1246
chr15.1211
chr12.347
chr13.1171
chr6.1642
chr4.936
chr8.762
chr13.1150
chr5.1620
chr15.20
chr8.1415
chr14.693
chr5.818
chr5.653
chr12.7
chr12.239
chr9.308
chr15.573
chr8.1454
chr9.844
chr8.444
chr9.236
chr12.10
chr1.677
chr12.295
chr9.16
chr2.1181
chr10.498
chr9.950
chr2.500
chr5.159
chr15.1088

chr9.686
chr11.216
chr6.624
chr5.1093
chr8.981
chr2.708
chr15.130
chr5.355
chr1.691
chr15.391
chr1.2560
chr1.1232
chr12.777
chr8.1521
chr5.392
chr14.45
chr8.1538
chr5.426
chr2.1179
chr1.631
chr15.1217
chr2.1275
chr9.1352
chr15.168
chr9.1550
chr2.1574
chr13.707
chr9.518
chr5.214
chr9.1558
chr15.888
chr2.1558
chr8.1012
chr8.1833
chr5.85
chr6.1078
chr2.1815
chr12.199
chr9.451
chr9.31
chr10.597
chr9.303
chr6.27
chr2.1290
chr13.770
chr3.1073
chr3.27
chr14.207
chr5.1338
chr15.83

chr6.675
chr2.729
chr6.107
chr8.1639
chr15.655
chr5.139
chr6.1054
chr10.700
chr4.2
chr10.881
chr12.506
chr5.1642
chr5.448
chr14.373
chr2.266
chr9.1559
chr5.707
chr9.831
chr7.785
chr9.1590
chr12.1165
chr9.748
chr1.216
chr9.142
chr9.1630
chr5.146
chr9.1210
chr10.326
chr8.1880
chr3.915
chr8.690
chr9.14
chr8.133
chr6.1222
chr2.1529
chr8.713
chr15.1029
chr9.951
chr5.979
chr5.727
chr13.1000
chr8.725
chr2.1366
chr8.1569
chr14.376
chr15.675
chr10.793
chr1.464
chr7.950
chr15.946

chr3.1157
chr9.551
chr12.810
chr7.1301
chr14.164
chr4.917
chr15.623
chr12.670
chr14.414
chr14.1424
chr15.10
chr9.1322
chr9.1604
chr10.374
chr7.1225
chr8.435
chr15.39
chr8.1780
chr2.578
chr15.737
chr8.1212
chr12.367
chr7.737
chr3.1466
chr8.604
chr1.175
chr15.719
chr5.62
chr2.486
chr5.1314
chr2.1665
chr2.298
chr2.1177
chr12.1285
chr13.1270
chr10.909
chr7.1085
chr10.989
chr7.891
chr7.436
chr8.321
chr14.822
chr6.948
chr11.1283
chr6.730
chr9.870
chr15.1021
chr9.1745
chr3.1055
chr8.1451

chr13.1083
chr5.267
chr15.265
chr10.978
chr5.1244
chr12.473
chr2.311
chr14.208
chr14.222
chr8.1074
chr6.187
chr9.1167
chr5.1164
chr8.322
chr8.1086
chr8.180
chr2.1775
chr6.1147
chr6.65
chr1.543
chr12.407
chr1.1564
chr14.355
chr12.591
chr10.1053
chr6.955
chr5.718
chr8.1026
chr14.441
chr8.804
chr7.965
chr9.337
chr12.1266
chr9.820
chr5.618
chr15.1099
chr6.947
chr10.576
chr2.921
chr7.1348
chr8.1333
chr2.454
chr5.1632
chr7.22
chr15.254
chr5.1657
chr9.848
chr5.1274
chr6.548
chr15.908

chr2.716
chr10.752
chr9.1109
chr15.763
chr5.170
chr15.388
chr6.1450
chr1.141
chr9.989
chr9.59
chr9.1489
chr2.212
chr2.81
chr9.1060
chr8.1730
chr9.550
chr7.405
chr2.783
chr15.1078
chr12.241
chr7.242
chr7.243
chr15.1005
chr2.1617
chr9.1724
chr5.1300
chr8.619
chr15.1117
chr15.1034
chr5.1646
chr9.1482
chr5.1406
chr9.395
chr14.541
chr2.1078
chr9.1256
chr15.1144
chr12.1283
chr5.608
chr2.1302
chr6.1840
chr8.746
chr14.916
chr15.1154
chr13.1314
chr13.1010
chr8.814
chr9.381
chr7.111
chr5.264

chr15.588
chr15.1010
chr15.924
chr8.766
chr8.9
chr9.1244
chr15.382
chr11.572
chr5.547
chr13.75
chr8.1411
chr1.476
chr2.1237
chr8.1300
chr5.527
chr9.805
chr8.1277
chr3.963
chr9.291
chr15.585
chr4.1683
chr15.1093
chr5.319
chr10.986
chr7.1416
chr9.646
chr9.75
chr15.521
chr3.1717
chr14.694
chr8.1792
chr6.1153
chr8.467
chr9.1567
chr8.1437
chr5.407
chr14.1422
chr2.1921
chr9.921
chr12.89
chr12.1
chr3.1664
chr15.166
chr10.1328
chr7.896
chr15.589
chr12.1086
chr13.1025
chr13.738
chr1.2177

chr11.121
chr7.410
chr15.980
chr8.1352
chr10.1031
chr9.46
chr8.387
chr6.845
chr3.1376
chr5.18
chr9.669
chr8.1417
chr15.190
chr10.1308
chr12.495
chr6.353
chr15.464
chr12.310
chr6.1053
chr15.307
chr14.644
chr7.32
chr9.568
chr11.252
chr14.808
chr6.1505
chr1.1934
chr12.605
chr3.1728
chr9.1704
chr12.307
chr2.197
chr5.928
chr8.1887
chr15.506
chr9.94
chr9.542
chr3.310
chr2.876
chr9.344
chr1.731
chr12.703
chr13.1412
chr14.968
chr9.315
chr10.164
chr9.1330
chr9.1631
chr10.1108
chr7.82

chr7.1316
chr5.1079
chr8.1509
chr8.1847
chr5.1032
chr9.1314
chr9.883
chr9.190
chr5.1226
chr2.701
chr6.275
chr4.1404
chr9.1048
chr9.72
chr1.388
chr8.1269
chr2.1889
chr8.252
chr10.288
chr15.742
chr11.776
chr5.1185
chr1.579
chr3.2071
chr13.391
chr10.1029
chr2.349
chr9.1233
chr8.1638
chr14.727
chr7.984
chr14.456
chr4.1726
chr5.84
chr5.1072
chr8.365
chr3.513
chr5.1670
chr4.903
chr3.1502
chr9.970
chr14.851
chr12.779
chr9.1368
chr7.1021
chr8.1149
chr15.975
chr9.1791
chr2.928
chr12.115

chr14.1359
chr8.1601
chr6.186
chr9.543
chr1.1162
chr9.1236
chr9.1663
chr15.429
chr2.1759
chr1.914
chr2.456
chr13.163
chr2.641
chr14.117
chr9.1391
chr8.482
chr9.1403
chr9.578
chr8.206
chr7.531
chr7.1067
chr14.82
chr15.261
chr8.6
chr15.229
chr14.1102
chr15.840
chr9.1202
chr7.1221
chr5.1634
chr2.1508
chr5.90
chr5.1239
chr13.1028
chr2.348
chr4.367
chr8.446
chr10.90
chr14.150
chr12.1092
chr12.455
chr2.98
chr7.282
chr10.1502
chr14.1362
chr8.999
chr5.919
chr2.319
chr9.209
chr2.1172

chr8.507
chr2.829
chr9.1164
chr9.746
chr5.607
chr5.1253
chr8.1197
chr9.855
chr5.988
chr8.669
chr7.177
chr5.33
chr2.1276
chr10.109
chr12.416
chr9.1581
chr4.180
chr15.439
chr8.459
chr5.499
chr2.1745
chr2.200
chr6.1983
chr15.522
chr2.906
chr5.670
chr9.602
chr15.536
chr8.310
chr3.1529
chr15.852
chr8.593
chr5.788
chr15.912
chr3.1484
chr2.1296
chr9.428
chr10.864
chr4.1903
chr15.246
chr2.602
chr11.820
chr9.5
chr2.106
chr8.1045
chr8.734
chr5.1069
chr2.379
chr15.185
chr15.181

chr9.105
chr14.1273
chr2.743
chr5.1436
chr15.583
chr2.1363
chr2.1120
chr5.115
chr14.1413
chr2.264
chr15.396
chr15.860
chr12.192
chr1.1113
chr2.1587
chr9.496
chr9.1366
chr8.1106
chr15.174
chr5.1278
chr2.127
chr5.52
chr5.726
chr13.1311
chr3.1509
chr2.934
chr13.407
chr5.864
chr14.883
chr5.1664
chr5.905
chr14.1128
chr12.251
chr9.442
chr8.1157
chr8.239
chr10.766
chr9.30
chr10.157
chr14.25
chr4.1382
chr14.301
chr9.11
chr15.920
chr6.1998
chr15.25
chr12.16
chr9.1054
chr12.34
chr9.262

chr6.1467
chr15.670
chr5.1211
chr12.268
chr9.1511
chr15.684
chr8.1089
chr10.346
chr12.1042
chr15.167
chr1.864
chr3.73
chr2.1443
chr8.1146
chr5.295
chr15.23
chr15.635
chr5.1680
chr12.489
chr9.1451
chr2.754
chr8.1219
chr5.328
chr6.943
chr9.1469
chr5.502
chr6.1727
chr9.1262
chr2.1935
chr13.926
chr5.565
chr13.725
chr7.758
chr10.427
chr15.1040
chr3.1056
chr15.511
chr14.299
chr12.632
chr2.99
chr5.649
chr5.69
chr2.625
chr5.1025
chr5.783
chr14.52
chr8.508
chr10.1183
chr2.643
chr9.1588

chr1.267
chr2.1584
chr10.1011
chr9.634
chr1.2373
chr10.1480
chr6.1146
chr9.6
chr1.1124
chr4.160
chr5.1340
chr10.103
chr9.1626
chr12.204
chr5.1656
chr15.163
chr2.1941
chr8.488
chr8.249
chr8.1320
chr9.1535
chr7.1409
chr9.1793
chr8.1479
chr9.250
chr4.255
chr13.481
chr14.85
chr14.1311
chr9.1602
chr2.843
chr10.209
chr10.1270
chr5.137
chr15.1180
chr8.648
chr13.656
chr8.270
chr7.118
chr10.548
chr15.534
chr8.1180
chr2.239
chr2.55
chr5.1186
chr1.2454
chr9.433
chr2.684
chr2.731
chr5.399

chr5.1050
chr2.998
chr12.341
chr7.659
chr5.771
chr5.1589
chr2.15
chr9.535
chr5.652
chr2.741
chr12.211
chr2.1906
chr5.984
chr8.1299
chr15.667
chr2.240
chr5.1468
chr5.1375
chr14.381
chr9.237
chr10.956
chr10.550
chr5.969
chr2.808
chr8.301
chr9.280
chr12.699
chr9.464
chr14.64
chr5.175
chr9.1215
chr15.432
chr15.356
chr9.319
chr2.979
chr2.874
chr15.1143
chr2.1477
chr8.1768
chr15.1052
chr13.709
chr7.817
chr10.838
chr5.1669
chr2.1916
chr5.162
chr9.1523
chr15.1164
chr10.1154
chr9.642

chr5.605
chr7.291
chr15.687
chr7.280
chr8.1167
chr5.644
chr10.173
chr8.935
chr2.1306
chr12.840
chr2.295
chr7.1254
chr15.236
chr9.687
chr14.13
chr8.222
chr10.1264
chr9.485
chr2.1336
chr2.318
chr5.604
chr14.515
chr12.40
chr15.390
chr15.1073
chr2.807
chr14.522
chr8.628
chr14.405
chr2.193
chr2.774
chr10.744
chr14.69
chr8.668
chr15.956
Cha14.1418

Supplementary Table 26: The groups of the treemix.

| CSS-1 | CSS-2 | CSS-3 | CSS-4 | CSS-5 | CSA-1 | CSA-2 | CSA-3 |
|---|---|---|---|---|---|---|---|
| ZBD | GE-1 | GE-4 | HZ081 | HZ076 | HZ045 | YNLDP2 | HZ119 |
| HZ043 | GE-8 | GZ | HZ086 | HZ065 | HZ061 | YNZ-1 | HZ114 |
| HZ028 | FL | GLJYD | HZ056 | HZ080 | HZ075 | LW-2 | HZ104 |
| HZ013 | XWY-5 | HZ009 | HZ063 | HZ078 | SLLK | HZ124 | HZ117 |
| HZ083 | XWY-1 | QXDM-2 | HZ048 | ZH079 | HZ074 | s301-1 | HZ123 |
| HZ090 | XWY-2 | QXDM-1 | LC-1 | DBZ-1 | HZ039 | s1-16 | HZ072 |
| JX-4 | DHL | HZ050 | LC-2 | HZ002 | HZ092 | HZ109 | HZ100 |
| HZ014 | XSL | HZ021 | HZ057 | HSKC | | LW-1 | HZ122 |
| HZ020 | JGL | YNLDP1 | HZ015 | | | LW-5 | HZ118 |
| HZ010 | SB | HZ037 | HZ034 | | | LW-3 | |
| HZ073 | MZYCS | HZ066 | HZ085 | | | LW-4 | |
| HZ071 | XSX | HZ070 | HZ059 | | | | |
| HZ018 | ZW | HZ064 | HZ082 | | | | |
| HZ068 | JHZS | HZ041 | HZ088 | | | | |
| HZ031 | XWY-4 | HZ069 | HZ112 | | | | |
| HZ003 | XWY-3 | HZ008 | GE-6 | | | | |
| zhen | WL | HZ016 | | | | | |
| HZ017 | BCS | HZ036 | | | | | |
| HZ077 | | | | | | | |
| HZ011 | | | | | | | |
| HZ006 | | | | | | | |
| HZ040 | | | | | | | |
| HZ004 | | | | | | | |
| HZ058 | | | | | | | |
| HZ060 | | | | | | | |
| XY-10 | | | | | | | |
| HZ054 | | | | | | | |

| CSA-4 | CSA-5 | CSR | outgroup |
|---|---|---|---|
| NBE-4 | ST330 | XYDCS | CM-1 |
| NBE-3 | ASM | HZ001 | |
| NBE-5 | s303-231 | HZ027 | |
| NBE-2 | s6-8 | LBDCS | |
| NBE-1 | YH-2 | NC | |
| | s317 | HZ084 | |
| | ST312 | HZ094 | |
| | XH-1 | HZ095 | |
| | YH-1-1 | HZ110 | |
| | WLH-1 | HZ093 | |
| | | HZ125 | |

Supplementary Table 28: F4-test of random individual.

| Groups | Statistic | Standard error | Z-score |
|---|---|---|---|
| HZ020,HZ085;YNZ-1,CM-1 | 0.000816975 | 0.000210335 | 3.88416 |
| HZ020,YNZ-1;HZ085,CM-1 | 0.00904846 | 0.000303599 | 29.804 |
| MZYCS,HZ016;HZ021,CM-1 | -0.00201348 | 0.000262648 | -7.66608 |
| MZYCS,HZ021;HZ016,CM-1 | -0.00127894 | 0.000278926 | -4.58522 |
| SB,NBE-1;XWY-5,CM-1 | 0.00656082 | 0.000267022 | 24.5703 |
| SB,XWY-5;NBE-1,CM-1 | -0.00102305 | 0.000196753 | -5.19968 |
| QXDM-1,HZ082;JX-4,CM-1 | 0.00104977 | 0.000261805 | 4.00976 |
| QXDM-1,JX-4;HZ082,CM-1 | 0.000200956 | 0.000256976 | 0.782006 |
| HZ110,HZ070;HZ072,CM-1 | -0.00065355 | 0.000264317 | -2.47261 |
| HZ110,HZ072;HZ070,CM-1 | -0.00391372 | 0.000240279 | -16.2882 |
| GE-4,HZ013;YH-1-1,CM-1 | 0.00220029 | 0.000215534 | 10.2086 |
| GE-4,YH-1-1;HZ013,CM-1 | 0.0107975 | 0.000297927 | 36.2422 |
| HZ008,HZ004;HZ006,CM-1 | -0.00158467 | 0.000275097 | -5.76042 |
| HZ008,HZ006;HZ004,CM-1 | -0.00252325 | 0.00026171 | -9.64137 |
| HZ013,GLJYD;XSL,CM-1 | 9.56E-05 | 0.000257881 | 0.370778 |
| HZ013,XSL;GLJYD,CM-1 | -0.00223111 | 0.000247828 | -9.00267 |
| YNLDP2,HZ125;YH-2,CM-1 | 0.00240386 | 0.000252584 | 9.5171 |
| YNLDP2,YH-2;HZ125,CM-1 | -0.00133097 | 0.000228704 | -5.81959 |
| WLH-1,HZ118;LC-2,CM-1 | 0.00186779 | 0.000199769 | 9.34976 |
| WLH-1,LC-2;HZ118,CM-1 | 0.00875356 | 0.000257254 | 34.0268 |
| LBDCS,HZ081;s1-16,CM-1 | -0.00517407 | 0.000239678 | -21.5875 |
| LBDCS,s1-16;HZ081,CM-1 | 0.000579459 | 0.00026789 | 2.16305 |
| HZ010,HZ016;XWY-3,CM-1 | 0.00105568 | 0.000275459 | 3.83243 |
| HZ010,XWY-3;HZ016,CM-1 | 0.00035063 | 0.0002542 | 1.37934 |
| HZ112,HZ088;NBE-2,CM-1 | -0.00017732 | 0.000207379 | -0.85506 |
| HZ112,NBE-2;HZ088,CM-1 | 0.0111183 | 0.000313497 | 35.4655 |
| HZ104,HZ003;HZ118,CM-1 | 0.0075036 | 0.000283476 | 26.47 |
| HZ104,HZ118;HZ003,CM-1 | -0.00056736 | 0.000207513 | -2.7341 |
| NBE-1,BCS;HZ082,CM-1 | -0.00686795 | 0.00027079 | -25.3627 |
| NBE-1,HZ082;BCS,CM-1 | -0.0052316 | 0.000294392 | -17.7709 |
| HZ045,HZ020;HZ021,CM-1 | -0.00543196 | 0.00026401 | -20.5749 |
| HZ045,HZ021;HZ020,CM-1 | -0.0049553 | 0.000264144 | -18.7599 |
| HZ050,HSKC;HZ069,CM-1 | 0.00440209 | 0.000270754 | 16.2586 |
| HZ050,HZ069;HSKC,CM-1 | 0.000313913 | 0.000231877 | 1.35379 |
| HZ066,HZ069;XY-10,CM-1 | 0.000493608 | 0.00024062 | 2.0514 |
| HZ066,XY-10;HZ069,CM-1 | 0.00204321 | 0.00024772 | 8.24808 |
| HZ065,GE-4;HZ074,CM-1 | -0.00032307 | 0.000229072 | -1.41035 |
| HZ065,HZ074;GE-4,CM-1 | 0.0037581 | 0.000275575 | 13.6373 |
| GE-1,SB;XWY-4,CM-1 | -0.00586985 | 0.00023123 | -25.3853 |
| GE-1,XWY-4;SB,CM-1 | -0.00542103 | 0.00023935 | -22.649 |
| HZ119,HZ036;HZ066,CM-1 | -0.0098496 | 0.000265813 | -37.0546 |
| HZ119,HZ066;HZ036,CM-1 | -0.00956249 | 0.000278061 | -34.3898 |
| XWY-5,JGL;YNLDP2,CM-1 | -0.0002289 | 0.000192557 | -1.18875 |
| XWY-5,YNLDP2;JGL,CM-1 | 0.00876416 | 0.000267331 | 32.784 |
| HZ068,HZ041;s1-16,CM-1 | -0.00024571 | 0.000198943 | -1.23507 |
| HZ068,s1-16;HZ041,CM-1 | 0.0108858 | 0.000302986 | 35.9284 |
| HZ040,QXDM-1;SLLK,CM-1 | -4.82E-05 | 0.000201849 | -0.238569 |
| HZ040,SLLK;QXDM-1,CM-1 | 0.00744724 | 0.00028826 | 25.8352 |

| | | | |
|---|---|---|---|
| HZ083,HZ014;ST330,CM-1 | 0.000387708 | 0.000197984 | 1.95828 |
| HZ083,ST330;HZ014,CM-1 | 0.0116414 | 0.000303262 | 38.3873 |
| HZ064,HZ013;WL,CM-1 | 0.00234276 | 0.000262685 | 8.91853 |
| HZ064,WL;HZ013,CM-1 | -0.00074086 | 0.000235355 | -3.14783 |
| GZ,FL;HZ017,CM-1 | -0.00333539 | 0.000251976 | -13.2369 |
| GZ,HZ017;FL,CM-1 | -0.0032265 | 0.000252513 | -12.7775 |
| HZ003,NBE-2;QXDM-1,CM-1 | 0.00500388 | 0.000305904 | 16.3577 |
| HZ003,QXDM-1;NBE-2,CM-1 | -0.00051815 | 0.000247455 | -2.09393 |
| HZ083,GZ;HZ034,CM-1 | 0.00221823 | 0.000256026 | 8.66406 |
| HZ083,HZ034;GZ,CM-1 | 0.00199877 | 0.0002564 | 7.79549 |
| ST330,HZ073;HZ074,CM-1 | -0.00122239 | 0.000238828 | -5.11829 |
| ST330,HZ074;HZ073,CM-1 | 0.00198934 | 0.000270774 | 7.34686 |
| NBE-5,HZ001;HZ034,CM-1 | 0.00297124 | 0.000287811 | 10.3236 |
| NBE-5,HZ034;HZ001,CM-1 | -0.00424374 | 0.000240099 | -17.6749 |
| XSX,HZ006;HZ079,CM-1 | -0.0004004 | 0.000240186 | -1.66702 |
| XSX,HZ079;HZ006,CM-1 | 0.00578944 | 0.000283132 | 20.4478 |
| HZ020,HZ083;XH-1,CM-1 | -0.00115604 | 0.000213033 | -5.42659 |
| HZ020,XH-1;HZ083,CM-1 | 0.0070461 | 0.000269449 | 26.1501 |
| HZ084,HZ119;NBE-3,CM-1 | -0.00309565 | 0.000251439 | -12.3117 |
| HZ084,NBE-3;HZ119,CM-1 | -0.00366614 | 0.000252621 | -14.5125 |
| HZ014,HZ061;QXDM-1,CM-1 | 0.0057223 | 0.00026539 | 21.5619 |
| HZ014,QXDM-1;HZ061,CM-1 | 0.00129619 | 0.000221472 | 5.85262 |
| LBDCS,HZ048;XWY-1,CM-1 | -0.00765709 | 0.000256469 | -29.8558 |
| LBDCS,XWY-1;HZ048,CM-1 | -0.005111 | 0.000289243 | -17.6702 |
| XWY-2,HSKC;HZ075,CM-1 | 0.00239175 | 0.000242411 | 9.86649 |
| XWY-2,HZ075;HSKC,CM-1 | 0.00344921 | 0.00024671 | 13.9808 |
| s317,DHL;WL,CM-1 | -0.0167425 | 0.00034157 | -49.0163 |
| s317,WL;DHL,CM-1 | -0.0166655 | 0.000336426 | -49.5368 |
| HZ092,GZ;HSKC,CM-1 | -0.00714293 | 0.000265221 | -26.932 |
| HZ092,HSKC;GZ,CM-1 | -0.00516038 | 0.000288156 | -17.9083 |
| NBE-3,HZ088;HZ118,CM-1 | 0.0024381 | 0.000242387 | 10.0587 |
| NBE-3,HZ118;HZ088,CM-1 | 0.00529291 | 0.000263915 | 20.0553 |
| XSX,DHL;LW-4,CM-1 | 0.000477723 | 0.000184631 | 2.58745 |
| XSX,LW-4;DHL,CM-1 | 0.0196353 | 0.000336935 | 58.2762 |
| HZ125,HZ072;HZ092,CM-1 | -0.00394979 | 0.000220375 | -17.923 |
| HZ125,HZ092;HZ072,CM-1 | 0.00222071 | 0.000260243 | 8.53322 |
| HZ092,HZ073;XSX,CM-1 | -0.0139383 | 0.000318587 | -43.7504 |
| HZ092,XSX;HZ073,CM-1 | -0.0145723 | 0.000322064 | -45.2464 |
| HZ061,HZ016;XSX,CM-1 | -0.00644611 | 0.000298002 | -21.6311 |
| HZ061,XSX;HZ016,CM-1 | -0.00697323 | 0.000298617 | -23.3518 |
| HZ017,HZ013;QXDM-2,CM-1 | 0.00310136 | 0.000255588 | 12.1342 |
| HZ017,QXDM-2;HZ013,CM-1 | 0.00172929 | 0.000249232 | 6.9385 |
| LW-3,BCS;HZ028,CM-1 | -0.0146147 | 0.000304999 | -47.9173 |
| LW-3,HZ028;BCS,CM-1 | -0.0136954 | 0.000320873 | -42.6816 |
| LW-5,HZ043;HZ117,CM-1 | -0.00088608 | 0.000288276 | -3.07372 |
| LW-5,HZ117;HZ043,CM-1 | -0.00281563 | 0.000276864 | -10.1697 |
| HZ073,HZ122;XWY-2,CM-1 | 0.00871326 | 0.000309923 | 28.1143 |
| HZ073,XWY-2;HZ122,CM-1 | -0.00174491 | 0.000213627 | -8.16803 |
| HZ066,LBDCS;NBE-1,CM-1 | 0.00784336 | 0.000249021 | 31.4969 |
| HZ066,NBE-1;LBDCS,CM-1 | 0.00336571 | 0.000220601 | 15.257 |

| | | | |
|---|---|---|---|
| XYDCS,HZ071;HZ119,CM-1 | -0.0100634 | 0.000251791 | -39.9675 |
| XYDCS,HZ119;HZ071,CM-1 | -0.00693081 | 0.000272606 | -25.4243 |
| HZ028,NBE-4;XYDCS,CM-1 | 0.00204757 | 0.000227527 | 8.99925 |
| HZ028,XYDCS;NBE-4,CM-1 | 0.00691961 | 0.000263357 | 26.2747 |
| HZ013,ST330;XY-10,CM-1 | 0.00762316 | 0.000292813 | 26.0343 |
| HZ013,XY-10;ST330,CM-1 | -0.00165183 | 0.000211909 | -7.79499 |
| HZ002,HZ015;HZ122,CM-1 | -0.0011805 | 0.000233081 | -5.06477 |
| HZ002,HZ122;HZ015,CM-1 | 0.00465616 | 0.000273038 | 17.0532 |
| ST330,NBE-4;s303-231,CM-1 | 0.000611001 | 0.000243383 | 2.51044 |
| ST330,s303-231;NBE-4,CM-1 | -0.00111192 | 0.000238142 | -4.66915 |
| HZ114,HZ015;HZ018,CM-1 | -0.0147464 | 0.000307974 | -47.882 |
| HZ114,HZ018;HZ015,CM-1 | -0.0141609 | 0.000314038 | -45.0929 |
| XWY-4,HZ002;HZ009,CM-1 | 0.00338432 | 0.000247255 | 13.6876 |
| XWY-4,HZ009;HZ002,CM-1 | 0.00137368 | 0.000249583 | 5.50391 |
| LC-2,HZ013;HZ063,CM-1 | 0.00236724 | 0.000236044 | 10.0288 |
| LC-2,HZ063;HZ013,CM-1 | 0.000725532 | 0.000230739 | 3.14438 |
| HZ079,HZ090;XWY-1,CM-1 | 0.000564847 | 0.000234997 | 2.40363 |
| HZ079,XWY-1;HZ090,CM-1 | 0.00114208 | 0.000240602 | 4.74676 |
| NBE-5,HZ039;HZ081,CM-1 | 0.00552762 | 0.000304711 | 18.1405 |
| NBE-5,HZ081;HZ039,CM-1 | -0.00170497 | 0.000233329 | -7.30716 |
| HZ020,ASM;HZ080,CM-1 | 0.00559885 | 0.000262394 | 21.3376 |
| HZ020,HZ080;ASM,CM-1 | 0.000289969 | 0.000224372 | 1.29236 |
| HSKC,HZ011;HZ037,CM-1 | -0.00634155 | 0.000262905 | -24.1211 |
| HSKC,HZ037;HZ011,CM-1 | -0.0050061 | 0.000267812 | -18.6926 |
| LW-3,HZ041;HZ060,CM-1 | -0.0124993 | 0.000317798 | -39.3308 |
| LW-3,HZ060;HZ041,CM-1 | -0.0119997 | 0.000333157 | -36.0182 |
| HZ045,BCS;s1-16,CM-1 | 0.000833832 | 0.00021546 | 3.87001 |
| HZ045,s1-16;BCS,CM-1 | 0.00474834 | 0.000237346 | 20.006 |
| ASM,HZ013;YH-2,CM-1 | 0.0111279 | 0.000284399 | 39.1277 |
| ASM,YH-2;HZ013,CM-1 | 0.00240506 | 0.000205168 | 11.7224 |
| GE-8,DBZ-1;HZ084,CM-1 | 0.00354829 | 0.000240476 | 14.7553 |
| GE-8,HZ084;DBZ-1,CM-1 | 0.00618263 | 0.000264932 | 23.3367 |
| HZ058,HZ083;YH-1-1,CM-1 | 0.000164987 | 0.000213438 | 0.772998 |
| HZ058,YH-1-1;HZ083,CM-1 | 0.0124999 | 0.000342588 | 36.4867 |
| HZ021,DBZ-1;XWY-1,CM-1 | 0.0037274 | 0.000245539 | 15.1805 |
| HZ021,XWY-1;DBZ-1,CM-1 | 0.00240876 | 0.000250486 | 9.61637 |
| HZ076,HZ037;HZ054,CM-1 | -0.00886885 | 0.000286871 | -30.9158 |
| HZ076,HZ054;HZ037,CM-1 | -0.00977946 | 0.000280704 | -34.839 |
| HZ100,GE-6;HZ036,CM-1 | -0.0118425 | 0.000317055 | -37.3515 |
| HZ100,HZ036;GE-6,CM-1 | -0.0116668 | 0.000318328 | -36.6502 |
| XH-1,HZ070;HZ081,CM-1 | -0.00431212 | 0.000261776 | -16.4726 |
| XH-1,HZ081;HZ070,CM-1 | -0.00524694 | 0.000262286 | -20.0047 |
| GE-8,DHL;HZ123,CM-1 | 0.000185503 | 0.000198388 | 0.935051 |
| GE-8,HZ123;DHL,CM-1 | 0.0152185 | 0.000328298 | 46.3556 |
| HZ125,HZ006;HZ008,CM-1 | -0.018128 | 0.000331408 | -54.6999 |
| HZ125,HZ008;HZ006,CM-1 | -0.0181766 | 0.000330878 | -54.9344 |
| GLJYD,HZ084;ST312,CM-1 | 0.00477673 | 0.000238507 | 20.0276 |
| GLJYD,ST312;HZ084,CM-1 | 0.00453395 | 0.00024076 | 18.8318 |
| XSL,HZ003;s303-231,CM-1 | 0.000174757 | 0.000207412 | 0.842556 |
| XSL,s303-231;HZ003,CM-1 | 0.0106062 | 0.00031722 | 33.4348 |

| | | | |
|---|---|---|---|
| ST312,HZ009;HZ043,CM-1 | -0.00723092 | 0.000305187 | -23.6934 |
| ST312,HZ043;HZ009,CM-1 | -0.00732685 | 0.000307945 | -23.7927 |
| GZ,HZ077;SLLK,CM-1 | -0.0005836 | 0.000216811 | -2.69173 |
| GZ,SLLK;HZ077,CM-1 | 0.00582361 | 0.000279878 | 20.8077 |
| s6-8,HZ001;XH-1,CM-1 | 0.00876307 | 0.000258245 | 33.9331 |
| s6-8,XH-1;HZ001,CM-1 | -0.00115928 | 0.000180586 | -6.41952 |
| XY-10,ST312;XYDCS,CM-1 | 0.00510934 | 0.000247816 | 20.6175 |
| XY-10,XYDCS;ST312,CM-1 | 0.00602764 | 0.000254739 | 23.6621 |
| HZ016,NBE-4;ST330,CM-1 | -0.00257929 | 0.000273037 | -9.44665 |
| HZ016,ST330;NBE-4,CM-1 | -0.00014927 | 0.000312335 | -0.477916 |
| NBE-2,GE-4;GE-8,CM-1 | -0.00472684 | 0.00027998 | -16.8828 |
| NBE-2,GE-8;GE-4,CM-1 | -0.00441749 | 0.000273152 | -16.1723 |
| XWY-1,HZ112;NBE-2,CM-1 | 0.00167697 | 0.000265832 | 6.3084 |
| XWY-1,NBE-2;HZ112,CM-1 | 0.00153534 | 0.000267937 | 5.73021 |
| HZ100,XH-1;s301-1,CM-1 | -0.00175321 | 0.000225141 | -7.78715 |
| HZ100,s301-1;XH-1,CM-1 | -0.00146888 | 0.000226046 | -6.49815 |
| LC-1,HZ002;HZ009,CM-1 | 0.00202188 | 0.000228759 | 8.83848 |
| LC-1,HZ009;HZ002,CM-1 | 0.00109206 | 0.0002287 | 4.77509 |
| HZ048,HSKC;HZ031,CM-1 | 0.00199071 | 0.00027288 | 7.29517 |
| HZ048,HZ031;HSKC,CM-1 | -0.00102247 | 0.000247341 | -4.13382 |
| ST330,ASM;GLJYD,CM-1 | 0.00145362 | 0.000227613 | 6.38635 |
| ST330,GLJYD;ASM,CM-1 | 0.00468287 | 0.000262436 | 17.8439 |
| WLH-1,HZ056;HZ064,CM-1 | -0.00643064 | 0.000258529 | -24.874 |
| WLH-1,HZ064;HZ056,CM-1 | -0.00537882 | 0.000277292 | -19.3977 |
| HZ117,HZ068;NC,CM-1 | -0.00305216 | 0.000251077 | -12.1563 |
| HZ117,NC;HZ068,CM-1 | 0.000682654 | 0.000260461 | 2.62094 |
| HZ037,HZ083;XYDCS,CM-1 | -0.00066673 | 0.000210589 | -3.16603 |
| HZ037,XYDCS;HZ083,CM-1 | 0.0124404 | 0.000288996 | 43.0471 |
| HZ068,LC-1;YNZ-1,CM-1 | 0.000486005 | 0.000194391 | 2.50014 |
| HZ068,YNZ-1;LC-1,CM-1 | 0.0101901 | 0.000282337 | 36.0921 |
| HZ034,XSX;s303-231,CM-1 | -0.00112312 | 0.000238804 | -4.70311 |
| HZ034,s303-231;XSX,CM-1 | 0.00701798 | 0.000314539 | 22.3119 |
| HZ014,HZ079;HZ081,CM-1 | 0.00425693 | 0.000261095 | 16.3042 |
| HZ014,HZ081;HZ079,CM-1 | 0.000119256 | 0.000218931 | 0.544717 |
| HZ104,HZ122;WLH-1,CM-1 | -0.00119539 | 0.000233539 | -5.11861 |
| HZ104,WLH-1;HZ122,CM-1 | -0.00114752 | 0.000231321 | -4.96073 |
| HZ112,HZ073;NBE-2,CM-1 | -0.00182965 | 0.00022888 | -7.99391 |
| HZ112,NBE-2;HZ073,CM-1 | 0.00307003 | 0.00029153 | 10.5308 |
| HZ068,HSKC;HZ031,CM-1 | 0.005716 | 0.000271562 | 21.0486 |
| HZ068,HZ031;HSKC,CM-1 | -0.00016466 | 0.000224695 | -0.732799 |
| HZ079,HZ094;XSX,CM-1 | 0.0188294 | 0.000341081 | 55.205 |
| HZ079,XSX;HZ094,CM-1 | 0.000176837 | 0.000189239 | 0.934465 |
| HZ057,GLJYD;LW-2,CM-1 | -0.00171168 | 0.000222125 | -7.70591 |
| HZ057,LW-2;GLJYD,CM-1 | 0.00675929 | 0.000291833 | 23.1615 |
| s301-1,GE-1;GZ,CM-1 | -0.00590619 | 0.000251045 | -23.5264 |
| s301-1,GZ;GE-1,CM-1 | -0.00599302 | 0.000257434 | -23.2798 |
| HZ088,HZ004;HZ094,CM-1 | 0.000502267 | 0.000199443 | 2.51835 |
| HZ088,HZ094;HZ004,CM-1 | 0.0209832 | 0.000379226 | 55.3316 |
| HZ100,HZ072;HZ082,CM-1 | -0.00121629 | 0.000206795 | -5.88162 |
| HZ100,HZ082;HZ072,CM-1 | 0.00774849 | 0.000283579 | 27.3239 |

| | | | |
|---|---|---|---|
| XSX,HZ076;WLH-1,CM-1 | 0.00255753 | 0.000224143 | 11.4102 |
| XSX,WLH-1;HZ076,CM-1 | 0.00522038 | 0.0002527 | 20.6584 |
| HZ090,NBE-3;SLLK,CM-1 | 0.000784005 | 0.000221269 | 3.54322 |
| HZ090,SLLK;NBE-3,CM-1 | 0.00134152 | 0.000230986 | 5.8078 |
| HZ112,ASM;XY-10,CM-1 | 0.00996318 | 0.000301848 | 33.0073 |
| HZ112,XY-10;ASM,CM-1 | -2.38E-05 | 0.000217386 | -0.109663 |
| GE-8,DHL;HZ084,CM-1 | 0.00151524 | 0.000215254 | 7.0393 |
| GE-8,HZ084;DHL,CM-1 | 0.0101851 | 0.000278912 | 36.5172 |
| HZ068,HZ058;HZ074,CM-1 | -0.0005521 | 0.000238605 | -2.31388 |
| HZ068,HZ074;HZ058,CM-1 | 0.00781217 | 0.000310197 | 25.1846 |
| LC-2,HZ043;YH-1-1,CM-1 | 0.000434675 | 0.000222969 | 1.94949 |
| LC-2,YH-1-1;HZ043,CM-1 | 0.0110076 | 0.000319753 | 34.4254 |
| s1-16,HZ010;HZ060,CM-1 | -0.0143084 | 0.000318668 | -44.9006 |
| s1-16,HZ060;HZ010,CM-1 | -0.013972 | 0.000327467 | -42.6667 |
| LC-1,HZ017;XWY-1,CM-1 | -0.00169459 | 0.000216424 | -7.82992 |
| LC-1,XWY-1;HZ017,CM-1 | 0.0035521 | 0.000271582 | 13.0793 |
| HZ088,HZ092;NBE-4,CM-1 | 0.00489434 | 0.000260019 | 18.823 |
| HZ088,NBE-4;HZ092,CM-1 | 0.0015228 | 0.000240643 | 6.32803 |
| HZ004,HZ070;LW-3,CM-1 | -0.00023897 | 0.000211606 | -1.12932 |
| HZ004,LW-3;HZ070,CM-1 | 0.0126769 | 0.000326189 | 38.8636 |
| HZ104,HZ073;HZ081,CM-1 | -0.0135485 | 0.000310061 | -43.6964 |
| HZ104,HZ081;HZ073,CM-1 | -0.014133 | 0.00031931 | -44.2611 |
| HZ086,HZ069;s317,CM-1 | 3.43E-05 | 0.00019885 | 0.172289 |
| HZ086,s317;HZ069,CM-1 | 0.0127351 | 0.00031881 | 39.9456 |
| LW-4,XWY-5;s301-1,CM-1 | -0.00183689 | 0.000266167 | -6.90128 |
| LW-4,s301-1;XWY-5,CM-1 | -0.00432884 | 0.000245127 | -17.6596 |
| HZ117,HZ054;QXDM-1,CM-1 | -0.00919027 | 0.000295805 | -31.0687 |
| HZ117,QXDM-1;HZ054,CM-1 | -0.00839828 | 0.000301239 | -27.8792 |
| HZ073,HZ088;HZ114,CM-1 | -0.00039235 | 0.00020233 | -1.93918 |
| HZ073,HZ114;HZ088,CM-1 | 0.0156204 | 0.000326732 | 47.808 |
| YH-1-1,HZ088;NBE-4,CM-1 | 0.00359267 | 0.000324997 | 11.0545 |
| YH-1-1,NBE-4;HZ088,CM-1 | -0.00458342 | 0.00023349 | -19.6301 |
| ST330,HZ034;HZ118,CM-1 | 0.00343772 | 0.000235687 | 14.5859 |
| ST330,HZ118;HZ034,CM-1 | 0.00364693 | 0.000251273 | 14.5138 |
| JHZS,HZ014;HZ092,CM-1 | 0.000199863 | 0.000198326 | 1.00775 |
| JHZS,HZ092;HZ014,CM-1 | 0.0133684 | 0.000303856 | 43.9959 |
| HZ002,HZ009;HZ122,CM-1 | -0.00046668 | 0.000210696 | -2.21493 |
| HZ002,HZ122;HZ009,CM-1 | 0.0061718 | 0.000275609 | 22.3933 |
| WLH-1,HZ043;HZ079,CM-1 | -0.00463596 | 0.000292397 | -15.855 |
| WLH-1,HZ079;HZ043,CM-1 | -0.00805536 | 0.000273241 | -29.4808 |
| HZ021,HZ018;NBE-1,CM-1 | -0.0003234 | 0.000217084 | -1.48976 |
| HZ021,NBE-1;HZ018,CM-1 | 0.00607492 | 0.000269926 | 22.5058 |
| HZ056,HZ036;JHZS,CM-1 | -0.00548873 | 0.000283632 | -19.3516 |
| HZ056,JHZS;HZ036,CM-1 | -0.00605268 | 0.00026441 | -22.8912 |
| GZ,HZ124;XWY-5,CM-1 | 0.0105835 | 0.000294435 | 35.9453 |
| GZ,XWY-5;HZ124,CM-1 | 4.81E-05 | 0.000209012 | 0.230085 |
| HZ043,HZ011;HZ094,CM-1 | -4.72E-06 | 0.000178609 | -0.0264522 |
| HZ043,HZ094;HZ011,CM-1 | 0.0242571 | 0.000426458 | 56.8805 |
| XH-1,JX-4;s1-16,CM-1 | 4.10E-05 | 0.000210042 | 0.195112 |
| XH-1,s1-16;JX-4,CM-1 | 0.00534335 | 0.000250667 | 21.3165 |

| | | | |
|---|---|---|---|
| HZ081,HZ014;HZ072,CM-1 | 0.000352999 | 0.000205062 | 1.72142 |
| HZ081,HZ072;HZ014,CM-1 | 0.0121333 | 0.000306489 | 39.5881 |
| XY-10,HZ085;SB,CM-1 | 0.00305672 | 0.0002426 | 12.5998 |
| XY-10,SB;HZ085,CM-1 | 0.00148141 | 0.000223378 | 6.63186 |
| HZ034,HZ117;s6-8,CM-1 | -8.03E-05 | 0.000248946 | -0.322451 |
| HZ034,s6-8;HZ117,CM-1 | 0.000925732 | 0.000245054 | 3.77767 |
| HZ119,HZ050;HZ104,CM-1 | 0.00850856 | 0.00023728 | 35.8588 |
| HZ119,HZ104;HZ050,CM-1 | 0.00525282 | 0.000221735 | 23.6896 |
| s6-8,HZ125;NBE-3,CM-1 | 0.0103271 | 0.000266542 | 38.7446 |
| s6-8,NBE-3;HZ125,CM-1 | 0.000521059 | 0.000186864 | 2.78844 |
| HZ075,HSKC;HZ060,CM-1 | -0.0031349 | 0.000280094 | -11.1923 |
| HZ075,HZ060;HSKC,CM-1 | -0.00579664 | 0.000270666 | -21.4162 |
| GE-4,HZ104;XWY-4,CM-1 | 0.0126643 | 0.000328503 | 38.5514 |
| GE-4,XWY-4;HZ104,CM-1 | 0.000539954 | 0.000211812 | 2.54922 |
| XWY-2,HZ041;NC,CM-1 | -0.0007207 | 0.000210826 | -3.41843 |
| XWY-2,NC;HZ041,CM-1 | 0.00839292 | 0.000272486 | 30.8013 |
| HZ013,HZ084;YH-1-1,CM-1 | 0.00120284 | 0.000232772 | 5.16746 |
| HZ013,YH-1-1;HZ084,CM-1 | 0.00710453 | 0.000278454 | 25.5142 |
| HZ070,GE-8;XWY-2,CM-1 | 0.000441767 | 0.000261574 | 1.68888 |
| HZ070,XWY-2;GE-8,CM-1 | 0.000553437 | 0.000264695 | 2.09084 |
| XWY-4,HZ016;NBE-1,CM-1 | -0.00056327 | 0.000238915 | -2.35762 |
| XWY-4,NBE-1;HZ016,CM-1 | 0.00574724 | 0.000284695 | 20.1874 |
| HZ066,ST312;YNLDP1,CM-1 | 0.00276071 | 0.000279195 | 9.88812 |
| HZ066,YNLDP1;ST312,CM-1 | -0.00318629 | 0.000215781 | -14.7663 |
| NBE-1,HZ110;SB,CM-1 | 0.00899558 | 0.000247424 | 36.3569 |
| NBE-1,SB;HZ110,CM-1 | 0.00092585 | 0.000214408 | 4.31816 |
| XWY-2,JHZS;s317,CM-1 | 0.00316511 | 0.000213522 | 14.8234 |
| XWY-2,s317;JHZS,CM-1 | 0.0137077 | 0.000324305 | 42.268 |
| GE-4,HZ059;HZ095,CM-1 | 0.000107921 | 0.000192808 | 0.559735 |
| GE-4,HZ095;HZ059,CM-1 | 0.0209137 | 0.00033946 | 61.6089 |
| HZ095,HZ010;HZ117,CM-1 | -0.0128458 | 0.000322288 | -39.8581 |
| HZ095,HZ117;HZ010,CM-1 | -0.0142311 | 0.000301308 | -47.2312 |
| FL,HZ040;HZ075,CM-1 | 0.00034914 | 0.000195884 | 1.78238 |
| FL,HZ075;HZ040,CM-1 | 0.00886325 | 0.000270819 | 32.7276 |
| NBE-1,HZ027;HZ125,CM-1 | -0.00017293 | 0.000234331 | -0.737949 |
| NBE-1,HZ125;HZ027,CM-1 | 0.00047898 | 0.000227787 | 2.10275 |
| LC-2,HZ123;YNLDP1,CM-1 | 0.00601444 | 0.000287962 | 20.8863 |
| LC-2,YNLDP1;HZ123,CM-1 | -0.00174408 | 0.000212722 | -8.19887 |
| HZ081,HZ117;LC-2,CM-1 | 0.00886817 | 0.000280474 | 31.6185 |
| HZ081,LC-2;HZ117,CM-1 | 9.98E-05 | 0.000203585 | 0.490112 |
| HZ069,HZ061;HZ074,CM-1 | -0.0003752 | 0.000237612 | -1.57905 |
| HZ069,HZ074;HZ061,CM-1 | 0.00273047 | 0.000270431 | 10.0967 |
| LW-3,HSKC;HZ037,CM-1 | -0.00700268 | 0.000287755 | -24.3356 |
| LW-3,HZ037;HSKC,CM-1 | -0.00849832 | 0.000283672 | -29.9583 |
| HZ110,JX-4;XWY-5,CM-1 | -0.0128812 | 0.000291933 | -44.1239 |
| HZ110,XWY-5;JX-4,CM-1 | -0.0122951 | 0.000293115 | -41.9463 |
| NC,GE-6;s1-16,CM-1 | -0.00426716 | 0.000216521 | -19.7078 |
| NC,s1-16;GE-6,CM-1 | 0.00274583 | 0.000269961 | 10.1712 |
| XWY-3,HZ045;ST330,CM-1 | 0.000560449 | 0.00025064 | 2.23607 |
| XWY-3,ST330;HZ045,CM-1 | 0.00174606 | 0.000254037 | 6.87326 |

| | | | |
|---|---|---|---|
| MZYCS,GE-8;HZ090,CM-1 | 0.00112239 | 0.000201908 | 5.55891 |
| MZYCS,HZ090;GE-8,CM-1 | 0.00983528 | 0.000274067 | 35.8864 |
| ZBD,HZ104;HZ122,CM-1 | -0.00654146 | 0.000287083 | -22.786 |
| ZBD,HZ122;HZ104,CM-1 | -0.00810021 | 0.000260524 | -31.092 |
| NBE-5,ASM;HZ081,CM-1 | 0.00520226 | 0.00026684 | 19.4958 |
| NBE-5,HZ081;ASM,CM-1 | 0.00326532 | 0.000241896 | 13.4988 |
| FL,HZ015;HZ123,CM-1 | -0.00044447 | 0.000226286 | -1.9642 |
| FL,HZ123;HZ015,CM-1 | 0.00831989 | 0.000309323 | 26.8971 |
| s317,HZ056;HZ077,CM-1 | -0.00847558 | 0.000292576 | -28.9688 |
| s317,HZ077;HZ056,CM-1 | -0.00771399 | 0.000299841 | -25.727 |
| s6-8,HZ069;HZ080,CM-1 | -0.00399002 | 0.000242617 | -16.4458 |
| s6-8,HZ080;HZ069,CM-1 | -0.00124573 | 0.000265791 | -4.68686 |
| HZ079,HZ117;YNLDP2,CM-1 | 7.89E-05 | 0.000224587 | 0.351338 |
| HZ079,YNLDP2;HZ117,CM-1 | 0.00327441 | 0.000236379 | 13.8524 |
| HZ078,HZ021;HZ039,CM-1 | -0.00074983 | 0.000227377 | -3.29775 |
| HZ078,HZ039;HZ021,CM-1 | 0.00620222 | 0.00029983 | 20.6858 |
| HZ090,HZ009;HZ094,CM-1 | 0.000358235 | 0.000195566 | 1.83179 |
| HZ090,HZ094;HZ009,CM-1 | 0.0152434 | 0.000318051 | 47.9274 |
| GE-1,HZ104;XSL,CM-1 | 0.0143782 | 0.000315921 | 45.512 |
| GE-1,XSL;HZ104,CM-1 | 0.000528816 | 0.000202383 | 2.61295 |
| XSL,MZYCS;NBE-4,CM-1 | 0.000125219 | 0.000211108 | 0.593151 |
| XSL,NBE-4;MZYCS,CM-1 | 0.0117343 | 0.000296614 | 39.5609 |
| HZ016,HZ040;QXDM-2,CM-1 | 0.0051563 | 0.00026281 | 19.6198 |
| HZ016,QXDM-2;HZ040,CM-1 | -2.21E-05 | 0.00022487 | -0.0983396 |
| HZ076,HZ028;HZ104,CM-1 | -0.00029609 | 0.000220722 | -1.34147 |
| HZ076,HZ104;HZ028,CM-1 | 0.00501355 | 0.000273594 | 18.3247 |
| HZ068,JGL;NBE-4,CM-1 | 2.10E-05 | 0.000205737 | 0.101978 |
| HZ068,NBE-4;JGL,CM-1 | 0.00751056 | 0.000279205 | 26.8997 |
| HZ123,HZ076;HZ122,CM-1 | 0.0106069 | 0.000271574 | 39.0573 |
| HZ123,HZ122;HZ076,CM-1 | -0.00039197 | 0.000182099 | -2.15249 |
| HZ057,HZ006;HZ008,CM-1 | -0.00731984 | 0.000265214 | -27.5997 |
| HZ057,HZ008;HZ006,CM-1 | -0.00624844 | 0.000277036 | -22.5546 |
| DHL,HZ071;HZ084,CM-1 | -0.00022786 | 0.000213086 | -1.06933 |
| DHL,HZ084;HZ071,CM-1 | 0.00926208 | 0.000279211 | 33.1723 |
| HZ094,GE-1;HZ092,CM-1 | -0.0106123 | 0.000306637 | -34.6087 |
| HZ094,HZ092;GE-1,CM-1 | -0.0111008 | 0.000301054 | -36.8731 |
| HZ090,BCS;HZ021,CM-1 | -0.00516273 | 0.000256774 | -20.1061 |
| HZ090,HZ021;BCS,CM-1 | -0.00396695 | 0.000269487 | -14.7204 |
| ST312,HZ054;JGL,CM-1 | -0.0114776 | 0.000300289 | -38.2218 |
| ST312,JGL;HZ054,CM-1 | -0.0113008 | 0.000290021 | -38.9655 |
| HZ002,ASM;LW-2,CM-1 | -0.00179378 | 0.000228283 | -7.85771 |
| HZ002,LW-2;ASM,CM-1 | 0.000265671 | 0.000245414 | 1.08254 |
| FL,XWY-2;XY-10,CM-1 | 0.00298711 | 0.000253029 | 11.8054 |
| FL,XY-10;XWY-2,CM-1 | 0.0015551 | 0.000241966 | 6.42693 |
| LC-1,HZ040;HZ063,CM-1 | 0.00192992 | 0.000190732 | 10.1185 |
| LC-1,HZ063;HZ040,CM-1 | 0.00333447 | 0.000215181 | 15.4962 |
| HZ079,HZ009;HZ041,CM-1 | -0.00286995 | 0.000275401 | -10.421 |
| HZ079,HZ041;HZ009,CM-1 | -0.00453286 | 0.000259 | -17.5014 |
| HZ065,HZ016;YNLDP1,CM-1 | -0.00424268 | 0.000259371 | -16.3575 |
| HZ065,YNLDP1;HZ016,CM-1 | -0.00114271 | 0.000302594 | -3.77639 |

| | | | |
|---|---|---|---|
| FL,HZ017;NC,CM-1 | -0.0002195 | 0.000198955 | -1.10327 |
| FL,NC;HZ017,CM-1 | 0.0108441 | 0.000273589 | 39.6366 |
| NBE-5,HZ003;HZ117,CM-1 | 2.75E-05 | 0.000232 | 0.118501 |
| NBE-5,HZ117;HZ003,CM-1 | 0.00458221 | 0.000286211 | 16.0099 |
| HZ040,HZ069;XWY-5,CM-1 | -0.00037635 | 0.000229685 | -1.63853 |
| HZ040,XWY-5;HZ069,CM-1 | 0.00113751 | 0.000240641 | 4.72701 |
| XH-1,HZ021;LW-3,CM-1 | 0.000538936 | 0.000206967 | 2.60397 |
| XH-1,LW-3;HZ021,CM-1 | 0.00594993 | 0.00025652 | 23.1948 |
| HZ061,HZ028;HZ086,CM-1 | -0.00421659 | 0.000270808 | -15.5704 |
| HZ061,HZ086;HZ028,CM-1 | -0.00604546 | 0.000257288 | -23.4969 |
| LC-1,WL;s301-1,CM-1 | -0.0010862 | 0.000186886 | -5.8121 |
| LC-1,s301-1;WL,CM-1 | 0.0078404 | 0.00025342 | 30.9384 |
| BCS,DHL;HZ100,CM-1 | 7.48E-05 | 0.000192684 | 0.388277 |
| BCS,HZ100;DHL,CM-1 | 0.0163241 | 0.000339242 | 48.1195 |
| HZ028,HZ092;LW-5,CM-1 | 0.000318949 | 0.000254013 | 1.25564 |
| HZ028,LW-5;HZ092,CM-1 | 0.00110158 | 0.000278035 | 3.96204 |
| HZ027,HZ014;HZ043,CM-1 | -0.0115709 | 0.000301477 | -38.3806 |
| HZ027,HZ043;HZ014,CM-1 | -0.0109847 | 0.000297448 | -36.9298 |
| GE-8,HZ090;ZW,CM-1 | 0.00835478 | 0.00028748 | 29.0621 |
| GE-8,ZW;HZ090,CM-1 | -0.00135298 | 0.000204563 | -6.61403 |
| HZ060,HZ110;XYDCS,CM-1 | 0.0025931 | 0.000262296 | 9.88616 |
| HZ060,XYDCS;HZ110,CM-1 | -0.00125097 | 0.000248129 | -5.0416 |
| HZ069,HZ018;HZ119,CM-1 | -0.00020039 | 0.000195604 | -1.02448 |
| HZ069,HZ119;HZ018,CM-1 | 0.00797781 | 0.000265034 | 30.1011 |
| FL,HZ037;HZ080,CM-1 | 0.000174558 | 0.000208986 | 0.835261 |
| FL,HZ080;HZ037,CM-1 | 0.00615338 | 0.000266376 | 23.1003 |
| HZ073,HZ077;XYDCS,CM-1 | -0.00064581 | 0.00017829 | -3.62227 |
| HZ073,XYDCS;HZ077,CM-1 | 0.0139212 | 0.000299458 | 46.488 |
| SB,GE-1;LW-4,CM-1 | -0.00031692 | 0.000185388 | -1.70949 |
| SB,LW-4;GE-1,CM-1 | 0.013654 | 0.000301205 | 45.3311 |
| DHL,FL;HZ031,CM-1 | 0.00041265 | 0.000220839 | 1.86856 |
| DHL,HZ031;FL,CM-1 | 0.00342811 | 0.000256926 | 13.3428 |
| HZ082,HZ076;ST330,CM-1 | 0.00339163 | 0.000221353 | 15.3222 |
| HZ082,ST330;HZ076,CM-1 | 0.0053605 | 0.000243842 | 21.9835 |
| HZ064,HZ100;QXDM-2,CM-1 | 0.0146725 | 0.000329318 | 44.5543 |
| HZ064,QXDM-2;HZ100,CM-1 | -0.00019584 | 0.000194195 | -1.00847 |
| HZ123,HZ083;HZ090,CM-1 | -0.00586987 | 0.000301194 | -19.4887 |
| HZ123,HZ090;HZ083,CM-1 | -0.0103131 | 0.000265868 | -38.7902 |
| GE-8,HZ122;SLLK,CM-1 | 0.00359048 | 0.00028833 | 12.4527 |
| GE-8,SLLK;HZ122,CM-1 | -0.00165289 | 0.000244275 | -6.7665 |
| XWY-3,LW-2;NBE-5,CM-1 | 0.00834782 | 0.000303036 | 27.5473 |
| XWY-3,NBE-5;LW-2,CM-1 | -0.00077473 | 0.000224926 | -3.44438 |
| WL,HZ009;HZ123,CM-1 | -0.00037964 | 0.000216451 | -1.75394 |
| WL,HZ123;HZ009,CM-1 | 0.0103778 | 0.000311035 | 33.3653 |
| HZ100,HZ045;HZ082,CM-1 | -0.00810898 | 0.00025017 | -32.4139 |
| HZ100,HZ082;HZ045,CM-1 | -0.00228218 | 0.00028634 | -7.97019 |
| HZ015,HZ068;LW-1,CM-1 | 0.0010267 | 0.000218059 | 4.70837 |
| HZ015,LW-1;HZ068,CM-1 | 0.00873381 | 0.000294106 | 29.6961 |
| LC-1,HZ083;SB,CM-1 | -0.00181651 | 0.000225019 | -8.07267 |
| LC-1,SB;HZ083,CM-1 | -0.0017365 | 0.000219434 | -7.91357 |

| | | | |
|---|---|---|---|
| YNLDP1,HZ125;LW-1,CM-1 | 0.0038064 | 0.00024142 | 15.7667 |
| YNLDP1,LW-1;HZ125,CM-1 | 0.000216005 | 0.000218569 | 0.98827 |
| HZ073,GE-6;SLLK,CM-1 | -0.00123121 | 0.000214054 | -5.75186 |
| HZ073,SLLK;GE-6,CM-1 | 0.00560736 | 0.000284587 | 19.7035 |
| HZ039,HZ004;JX-4,CM-1 | -0.0115309 | 0.000306193 | -37.6588 |
| HZ039,JX-4;HZ004,CM-1 | -0.0112619 | 0.000322826 | -34.8855 |
| ST330,HZ031;s6-8,CM-1 | 0.00361867 | 0.000267378 | 13.5339 |
| ST330,s6-8;HZ031,CM-1 | -0.00038366 | 0.000231832 | -1.65491 |
| HZ063,HZ109;LW-2,CM-1 | -0.00249637 | 0.000245258 | -10.1785 |
| HZ063,LW-2;HZ109,CM-1 | -0.0002623 | 0.000267304 | -0.981261 |
| WL,XH-1;XYDCS,CM-1 | 0.00212435 | 0.000211297 | 10.0538 |
| WL,XYDCS;XH-1,CM-1 | 0.00822563 | 0.000269024 | 30.5758 |
| HZ040,HZ056;HZ090,CM-1 | 0.00267097 | 0.000231382 | 11.5436 |
| HZ040,HZ090;HZ056,CM-1 | 0.00218919 | 0.00023802 | 9.19749 |
| DBZ-1,HZ027;HZ084,CM-1 | -0.00111059 | 0.000271419 | -4.0918 |
| DBZ-1,HZ084;HZ027,CM-1 | -0.00461415 | 0.000253976 | -18.1677 |
| HZ082,LW-4;SB,CM-1 | 0.0112654 | 0.000297969 | 37.8072 |
| HZ082,SB;LW-4,CM-1 | 0.000359057 | 0.000200339 | 1.79225 |
| XWY-1,HZ092;QXDM-2,CM-1 | 0.00922648 | 0.000318634 | 28.9563 |
| XWY-1,QXDM-2;HZ092,CM-1 | -0.00102259 | 0.000217229 | -4.70741 |
| HZ015,HZ013;s6-8,CM-1 | -2.71E-05 | 0.000216834 | -0.124944 |
| HZ015,s6-8;HZ013,CM-1 | 0.00612797 | 0.000281119 | 21.7985 |
| s303-231,HZ016;HZ076,CM-1 | -0.00338287 | 0.000245195 | -13.7967 |
| s303-231,HZ076;HZ016,CM-1 | 0.000534306 | 0.000264373 | 2.02103 |
| HZ021,DBZ-1;YNLDP2,CM-1 | 0.000915634 | 0.000215453 | 4.24981 |
| HZ021,YNLDP2;DBZ-1,CM-1 | 0.00557519 | 0.000256098 | 21.7698 |
| s301-1,HZ001;YH-1-1,CM-1 | 0.00838509 | 0.000255773 | 32.7833 |
| s301-1,YH-1-1;HZ001,CM-1 | 0.00170874 | 0.000194377 | 8.79087 |
| HZ082,HZ074;ZBD,CM-1 | 0.00964179 | 0.000292511 | 32.9622 |
| HZ082,ZBD;HZ074,CM-1 | 0.000602844 | 0.000196955 | 3.06082 |
| HZ008,HZ016;s6-8,CM-1 | 0.000377036 | 0.00017779 | 2.12069 |
| HZ008,s6-8;HZ016,CM-1 | 0.0155572 | 0.000316866 | 49.0971 |
| GE-1,HZ072;LC-2,CM-1 | 0.0105826 | 0.000295396 | 35.8251 |
| GE-1,LC-2;HZ072,CM-1 | -0.00022272 | 0.000202946 | -1.09743 |
| HZ045,GE-1;YNZ-1,CM-1 | -1.74E-05 | 0.000220137 | -0.0788242 |
| HZ045,YNZ-1;GE-1,CM-1 | 0.00432577 | 0.000250947 | 17.2377 |
| LW-5,HZ066;LW-2,CM-1 | 0.00370262 | 0.000273686 | 13.5287 |
| LW-5,LW-2;HZ066,CM-1 | 0.00129676 | 0.000241575 | 5.36794 |
| HZ109,HZ014;HZ123,CM-1 | 0.00473559 | 0.000266082 | 17.7975 |
| HZ109,HZ123;HZ014,CM-1 | 0.000948375 | 0.00022477 | 4.21931 |
| JHZS,HZ039;XH-1,CM-1 | 0.00805706 | 0.000269239 | 29.9253 |
| JHZS,XH-1;HZ039,CM-1 | 0.00167627 | 0.000216599 | 7.73906 |
| HZ085,YNLDP2;s6-8,CM-1 | 0.00167612 | 0.000231632 | 7.23614 |
| HZ085,s6-8;YNLDP2,CM-1 | 0.00181508 | 0.000236364 | 7.67916 |
| HZ083,HZ123;LW-4,CM-1 | -0.00350304 | 0.000251334 | -13.9378 |
| HZ083,LW-4;HZ123,CM-1 | -0.00362002 | 0.000255456 | -14.1708 |
| LC-1,LW-5;YNLDP2,CM-1 | 0.00253355 | 0.000255202 | 9.92762 |
| LC-1,YNLDP2;LW-5,CM-1 | -0.00092061 | 0.000229776 | -4.00653 |
| HZ071,HZ076;ZBD,CM-1 | 0.00937682 | 0.000257635 | 36.3957 |
| HZ071,ZBD;HZ076,CM-1 | -0.00062143 | 0.000190233 | -3.26667 |

| | | | |
|---|---|---|---|
| HZ058,HZ039;HZ041,CM-1 | 0.00931524 | 0.000317383 | 29.3502 |
| HZ058,HZ041;HZ039,CM-1 | -2.78E-05 | 0.000231866 | -0.119969 |
| ST330,HZ016;HZ092,CM-1 | -0.00178596 | 0.000249811 | -7.14922 |
| ST330,HZ092;HZ016,CM-1 | 0.00335825 | 0.000289529 | 11.599 |
| HZ065,HZ017;HZ028,CM-1 | -0.00737366 | 0.000282721 | -26.081 |
| HZ065,HZ028;HZ017,CM-1 | -0.0062056 | 0.000288247 | -21.5287 |
| XWY-4,GE-1;HZ085,CM-1 | -0.0006052 | 0.000238647 | -2.53595 |
| XWY-4,HZ085;GE-1,CM-1 | 0.00162164 | 0.000258405 | 6.27558 |
| BCS,XWY-2;XWY-3,CM-1 | 0.00518121 | 0.000284035 | 18.2414 |
| BCS,XWY-3;XWY-2,CM-1 | -0.00071294 | 0.00022125 | -3.2223 |
| XWY-2,HZ014;NBE-3,CM-1 | 0.00182516 | 0.000243907 | 7.48303 |
| XWY-2,NBE-3;HZ014,CM-1 | 0.00407324 | 0.000266847 | 15.2643 |
| s303-231,HZ064;LW-2,CM-1 | 0.00138905 | 0.000232939 | 5.96317 |
| s303-231,LW-2;HZ064,CM-1 | 0.00497907 | 0.0002604 | 19.1209 |
| HZ118,HZ013;HZ070,CM-1 | -0.0105589 | 0.000313859 | -33.642 |
| HZ118,HZ070;HZ013,CM-1 | -0.0114127 | 0.000310318 | -36.7775 |
| HZ036,HZ112;HZ114,CM-1 | 0.000310122 | 0.000182483 | 1.69946 |
| HZ036,HZ114;HZ112,CM-1 | 0.0154605 | 0.000320506 | 48.2377 |
| HZ094,LW-4;XWY-5,CM-1 | -0.0097901 | 0.000292824 | -33.4334 |
| HZ094,XWY-5;LW-4,CM-1 | -0.00943457 | 0.000301001 | -31.344 |
| HZ063,HZ045;HZ077,CM-1 | 0.00426823 | 0.000247438 | 17.2497 |
| HZ063,HZ077;HZ045,CM-1 | 0.00108405 | 0.000219183 | 4.94586 |
| HZ125,HZ028;YNZ-1,CM-1 | -0.00515066 | 0.00026317 | -19.5716 |
| HZ125,YNZ-1;HZ028,CM-1 | -0.00626333 | 0.000262793 | -23.8337 |
| HZ109,HSKC;HZ060,CM-1 | -0.00754672 | 0.000300803 | -25.0886 |
| HZ109,HZ060;HSKC,CM-1 | -0.00877148 | 0.000286395 | -30.6272 |
| BCS,HZ039;HZ060,CM-1 | 0.0126048 | 0.00032607 | 38.6567 |
| BCS,HZ060;HZ039,CM-1 | 0.000331744 | 0.00021205 | 1.56446 |
| HZ109,HZ059;YH-1-1,CM-1 | 0.00154029 | 0.000243618 | 6.32259 |
| HZ109,YH-1-1;HZ059,CM-1 | 0.00150408 | 0.000251721 | 5.97517 |
| s1-16,HZ072;QXDM-2,CM-1 | -6.42E-05 | 0.000227271 | -0.282678 |
| s1-16,QXDM-2;HZ072,CM-1 | 0.00130125 | 0.000234008 | 5.56071 |
| HZ013,HZ050;HZ076,CM-1 | 0.00272462 | 0.000227111 | 11.9968 |
| HZ013,HZ076;HZ050,CM-1 | 0.0062846 | 0.00025049 | 25.0893 |
| HZ100,HZ123;LW-2,CM-1 | -0.00046671 | 0.000201392 | -2.31743 |
| HZ100,LW-2;HZ123,CM-1 | 0.00777988 | 0.000277062 | 28.0799 |
| XYDCS,HZ045;HZ048,CM-1 | -0.00409404 | 0.000266507 | -15.3619 |
| XYDCS,HZ048;HZ045,CM-1 | -0.00603214 | 0.000269983 | -22.3427 |
| HZ110,HZ034;JGL,CM-1 | -0.0137489 | 0.000294658 | -46.6605 |
| HZ110,JGL;HZ034,CM-1 | -0.0125909 | 0.000300446 | -41.9074 |
| HZ085,HZ054;YNLDP2,CM-1 | -6.16E-05 | 0.00020651 | -0.298488 |
| HZ085,YNLDP2;HZ054,CM-1 | 0.00801537 | 0.000273959 | 29.2576 |
| MZYCS,HZ009;JX-4,CM-1 | 0.00372165 | 0.000269304 | 13.8195 |
| MZYCS,JX-4;HZ009,CM-1 | 0.0018627 | 0.000257137 | 7.24402 |
| XH-1,HZ058;HZ083,CM-1 | -0.00534552 | 0.0002954 | -18.0959 |
| XH-1,HZ083;HZ058,CM-1 | -0.00658075 | 0.000278548 | -23.6252 |
| s301-1,HZ123;XWY-5,CM-1 | 0.00359083 | 0.000238679 | 15.0446 |
| s301-1,XWY-5;HZ123,CM-1 | 0.00184725 | 0.000220384 | 8.38199 |
| HZ043,HZ065;QXDM-2,CM-1 | 0.00458888 | 0.000313313 | 14.6463 |
| HZ043,QXDM-2;HZ065,CM-1 | -0.00016405 | 0.000255862 | -0.641183 |

| | | | |
|---|---|---|---|
| NBE-1,HZ017;HZ095,CM-1 | 0.00131251 | 0.000196766 | 6.67042 |
| NBE-1,HZ095;HZ017,CM-1 | 0.0176503 | 0.000339642 | 51.9674 |
| HZ036,HZ048;HZ083,CM-1 | 0.00369109 | 0.000275081 | 13.4182 |
| HZ036,HZ083;HZ048,CM-1 | -0.0010604 | 0.000228682 | -4.63701 |
| HZ008,HZ082;HZ109,CM-1 | -0.00055265 | 0.00022931 | -2.41003 |
| HZ008,HZ109;HZ082,CM-1 | 0.0090378 | 0.000310835 | 29.0758 |
| HZ112,HZ020;YNLDP2,CM-1 | -0.00264568 | 0.000228004 | -11.6037 |
| HZ112,YNLDP2;HZ020,CM-1 | 0.00537048 | 0.00029071 | 18.4736 |
| HZ009,HZ071;YH-2,CM-1 | 0.00030718 | 0.000203147 | 1.51211 |
| HZ009,YH-2;HZ071,CM-1 | 0.0108859 | 0.000303616 | 35.8541 |
| QXDM-2,HZ110;LW-2,CM-1 | 0.00310097 | 0.000253425 | 12.2362 |
| QXDM-2,LW-2;HZ110,CM-1 | 0.00089604 | 0.000242391 | 3.69667 |
| HZ119,HZ064;LC-2,CM-1 | -0.00666635 | 0.000256702 | -25.9692 |
| HZ119,LC-2;HZ064,CM-1 | -0.00576707 | 0.000257737 | -22.3758 |
| HZ016,HZ092;NBE-3,CM-1 | 0.0080335 | 0.000284538 | 28.2334 |
| HZ016,NBE-3;HZ092,CM-1 | 0.00177836 | 0.000226237 | 7.8606 |
| HZ011,GLJYD;HZ066,CM-1 | 0.0032265 | 0.000252704 | 12.7679 |
| HZ011,HZ066;GLJYD,CM-1 | 0.00152089 | 0.000234269 | 6.49205 |
| HZ011,DBZ-1;HZ054,CM-1 | 0.0053533 | 0.000261045 | 20.5072 |
| HZ011,HZ054;DBZ-1,CM-1 | 0.00067441 | 0.000213964 | 3.15198 |
| ST312,HZ118;XWY-5,CM-1 | 0.0041371 | 0.000241656 | 17.1198 |
| ST312,XWY-5;HZ118,CM-1 | 0.00355426 | 0.000232541 | 15.2844 |
| ZBD,HZ118;s303-231,CM-1 | -0.00088273 | 0.000289147 | -3.05287 |
| ZBD,s303-231;HZ118,CM-1 | -0.00499916 | 0.000247036 | -20.2366 |
| HZ114,HZ065;XSL,CM-1 | -0.0134255 | 0.000307045 | -43.7248 |
| HZ114,XSL;HZ065,CM-1 | -0.0134068 | 0.000307985 | -43.5306 |
| HZ085,HZ013;LC-1,CM-1 | 0.000655964 | 0.00022753 | 2.88298 |
| HZ085,LC-1;HZ013,CM-1 | -0.00024193 | 0.000224063 | -1.07974 |
| s317,HZ031;HZ080,CM-1 | -0.00740704 | 0.000278859 | -26.562 |
| s317,HZ080;HZ031,CM-1 | -0.00719798 | 0.000288387 | -24.9595 |
| HZ040,HZ061;HZ083,CM-1 | 0.00944186 | 0.000263151 | 35.8801 |
| HZ040,HZ083;HZ061,CM-1 | -0.00054119 | 0.000195188 | -2.77264 |
| HZ017,HZ027;HZ065,CM-1 | 0.00933774 | 0.000272914 | 34.2149 |
| HZ017,HZ065;HZ027,CM-1 | 0.00218175 | 0.000226901 | 9.61545 |
| HZ069,HZ066;HZ114,CM-1 | 9.56E-05 | 0.000176821 | 0.54063 |
| HZ069,HZ114;HZ066,CM-1 | 0.0197539 | 0.000328495 | 60.1345 |
| HZ094,HZ009;HZ109,CM-1 | -0.00921472 | 0.000310876 | -29.6411 |
| HZ094,HZ109;HZ009,CM-1 | -0.00992457 | 0.000302326 | -32.8273 |
| HZ043,LW-1;ZW,CM-1 | 0.0150899 | 0.000364896 | 41.354 |
| HZ043,ZW;LW-1,CM-1 | -6.10E-05 | 0.000219813 | -0.277464 |
| s1-16,GE-8;NBE-5,CM-1 | -0.00578317 | 0.000270193 | -21.4039 |
| s1-16,NBE-5;GE-8,CM-1 | -0.00599895 | 0.000270783 | -22.1541 |
| ST312,HZ077;HZ084,CM-1 | -0.00660176 | 0.00027661 | -23.8667 |
| ST312,HZ084;HZ077,CM-1 | -0.00237468 | 0.000304875 | -7.78902 |
| HZ034,HZ009;NBE-5,CM-1 | -0.00107819 | 0.000265554 | -4.06017 |
| HZ034,NBE-5;HZ009,CM-1 | 0.000791773 | 0.000265716 | 2.97977 |
| XWY-1,HZ063;JX-4,CM-1 | 0.000561581 | 0.000267314 | 2.10083 |
| XWY-1,JX-4;HZ063,CM-1 | -0.00138346 | 0.000242711 | -5.7 |
| XSL,GE-1;GZ,CM-1 | 0.000982121 | 0.000244936 | 4.00971 |
| XSL,GZ;GE-1,CM-1 | 0.00326795 | 0.000251346 | 13.0018 |

| | | | |
|---|---|---|---|
| HZ065,HZ014;LW-2,CM-1 | 0.000305232 | 0.000208673 | 1.46273 |
| HZ065,LW-2;HZ014,CM-1 | 0.0087662 | 0.00028764 | 30.4762 |
| NBE-5,HZ057;NBE-4,CM-1 | 0.0085654 | 0.000284363 | 30.1214 |
| NBE-5,NBE-4;HZ057,CM-1 | 0.0010578 | 0.000223895 | 4.72455 |
| ZBD,HZ004;HZ074,CM-1 | 0.000415248 | 0.000191284 | 2.17085 |
| ZBD,HZ074;HZ004,CM-1 | 0.0113567 | 0.000287742 | 39.4682 |
| HZ082,HZ074;HZ112,CM-1 | 0.0122616 | 0.000318367 | 38.5141 |
| HZ082,HZ112;HZ074,CM-1 | -7.30E-05 | 0.000207455 | -0.351777 |
| GE-4,HZ069;JGL,CM-1 | -0.00153963 | 0.000243513 | -6.32257 |
| GE-4,JGL;HZ069,CM-1 | -0.00126698 | 0.000233059 | -5.43632 |
| HZ075,HZ118;NBE-1,CM-1 | 0.000813793 | 0.000254751 | 3.19446 |
| HZ075,NBE-1;HZ118,CM-1 | -0.00159714 | 0.000238314 | -6.70183 |
| HZ119,HZ112;HZ122,CM-1 | 0.00292092 | 0.000231395 | 12.6231 |
| HZ119,HZ122;HZ112,CM-1 | 0.00238376 | 0.000231738 | 10.2865 |
| SB,HZ125;XWY-5,CM-1 | 0.016797 | 0.000294455 | 57.0444 |
| SB,XWY-5;HZ125,CM-1 | -0.00070304 | 0.000166906 | -4.2122 |
| FL,HZ014;JX-4,CM-1 | -0.00183129 | 0.000242036 | -7.56618 |
| FL,JX-4;HZ014,CM-1 | -0.00021587 | 0.000248691 | -0.868029 |
| HZ125,HZ031;ZBD,CM-1 | -0.0161485 | 0.000310574 | -51.9956 |
| HZ125,ZBD;HZ031,CM-1 | -0.0161926 | 0.000298346 | -54.2745 |
| HZ085,DBZ-1;HZ073,CM-1 | -0.00065229 | 0.000254217 | -2.56589 |
| HZ085,HZ073;DBZ-1,CM-1 | -0.00311601 | 0.000247436 | -12.5932 |
| WL,FL;XWY-2,CM-1 | 0.00566747 | 0.000240318 | 23.5832 |
| WL,XWY-2;FL,CM-1 | 0.00439279 | 0.00024328 | 18.0565 |
| HZ073,HZ078;HZ124,CM-1 | -0.00070381 | 0.000219164 | -3.21131 |
| HZ073,HZ124;HZ078,CM-1 | 0.00930119 | 0.000310889 | 29.9181 |
| XWY-1,XWY-5;s1-16,CM-1 | -1.76E-05 | 0.000172372 | -0.102267 |
| XWY-1,s1-16;XWY-5,CM-1 | 0.014858 | 0.000310578 | 47.8397 |
| HSKC,HZ061;HZ069,CM-1 | 0.00110732 | 0.000267701 | 4.1364 |
| HSKC,HZ069;HZ061,CM-1 | -0.0023254 | 0.000240972 | -9.65009 |
| HZ056,HZ006;HZ021,CM-1 | -0.00442039 | 0.000265033 | -16.6787 |
| HZ056,HZ021;HZ006,CM-1 | -0.00407459 | 0.000271874 | -14.9871 |
| XWY-5,ST330;YNLDP2,CM-1 | 0.00159489 | 0.000220659 | 7.22785 |
| XWY-5,YNLDP2;ST330,CM-1 | 0.0040499 | 0.00024048 | 16.8409 |
| HZ027,HZ065;HZ079,CM-1 | -0.005235 | 0.000263889 | -19.8379 |
| HZ027,HZ079;HZ065,CM-1 | -0.00611096 | 0.000248955 | -24.5464 |
| HZ010,LW-1;LW-3,CM-1 | -0.00855338 | 0.000292935 | -29.1989 |
| HZ010,LW-3;LW-1,CM-1 | -0.0087368 | 0.000286272 | -30.5192 |
| HZ078,HZ076;s303-231,CM-1 | 0.00433062 | 0.000235179 | 18.4142 |
| HZ078,s303-231;HZ076,CM-1 | 0.00239653 | 0.000231876 | 10.3354 |
| ZBD,HZ020;XSL,CM-1 | 0.000413678 | 0.00022396 | 1.84711 |
| ZBD,XSL;HZ020,CM-1 | 0.000245378 | 0.000234677 | 1.0456 |
| HZ041,HZ011;LW-5,CM-1 | 0.000327491 | 0.000217724 | 1.50416 |
| HZ041,LW-5;HZ011,CM-1 | 0.0131254 | 0.000317876 | 41.2909 |
| SB,LW-5;YNLDP2,CM-1 | 0.00295529 | 0.000270194 | 10.9377 |
| SB,YNLDP2;LW-5,CM-1 | -0.00083759 | 0.000231748 | -3.61423 |
| HZ085,HZ125;YH-2,CM-1 | 0.0012829 | 0.000261208 | 4.91142 |
| HZ085,YH-2;HZ125,CM-1 | -0.00151186 | 0.000252867 | -5.97889 |
| HZ056,HZ016;HZ070,CM-1 | -0.00720982 | 0.000273956 | -26.3174 |
| HZ056,HZ070;HZ016,CM-1 | -0.00698951 | 0.000273671 | -25.5399 |

| | | | |
|---|---|---|---|
| HZ050,HZ020;HZ074,CM-1 | -0.00041009 | 0.000204092 | -2.00932 |
| HZ050,HZ074;HZ020,CM-1 | 0.00927379 | 0.000298784 | 31.0384 |
| YNLDP1,HZ092;s301-1,CM-1 | 0.00479774 | 0.000247332 | 19.398 |
| YNLDP1,s301-1;HZ092,CM-1 | 0.000801082 | 0.000208412 | 3.84374 |
| HZ041,HZ013;LC-1,CM-1 | -0.00034043 | 0.000246872 | -1.37896 |
| HZ041,LC-1;HZ013,CM-1 | -0.00102006 | 0.000230884 | -4.41807 |
| HZ069,HZ080;SLLK,CM-1 | 0.0010199 | 0.000232993 | 4.37737 |
| HZ069,SLLK;HZ080,CM-1 | 0.00266208 | 0.000255508 | 10.4188 |
| HZ068,HZ060;s301-1,CM-1 | -7.38E-05 | 0.000205874 | -0.358525 |
| HZ068,s301-1;HZ060,CM-1 | 0.00933942 | 0.000292754 | 31.902 |
| HZ008,HSKC;s6-8,CM-1 | 0.00412936 | 0.000231897 | 17.8068 |
| HZ008,s6-8;HSKC,CM-1 | 0.0058405 | 0.000257542 | 22.6778 |
| HZ109,HZ045;LW-2,CM-1 | 0.00108676 | 0.000237085 | 4.58386 |
| HZ109,LW-2;HZ045,CM-1 | 0.00140127 | 0.00023505 | 5.96159 |
| HZ064,DHL;GE-8,CM-1 | -0.00280543 | 0.000249858 | -11.2281 |
| HZ064,GE-8;DHL,CM-1 | -0.00120975 | 0.000269574 | -4.48766 |
| HZ015,GZ;HZ064,CM-1 | -0.00207783 | 0.000254625 | -8.16036 |
| HZ015,HZ064;GZ,CM-1 | -0.00306368 | 0.000249936 | -12.2579 |
| HZ114,HZ074;HZ110,CM-1 | 0.0027305 | 0.00023855 | 11.4462 |
| HZ114,HZ110;HZ074,CM-1 | -0.00077353 | 0.00020363 | -3.79872 |
| HZ094,HZ021;LW-4,CM-1 | -0.00913077 | 0.000307265 | -29.7163 |
| HZ094,LW-4;HZ021,CM-1 | -0.00954531 | 0.000301159 | -31.6953 |
| JHZS,BCS;NBE-1,CM-1 | 0.000295998 | 0.000184127 | 1.60757 |
| JHZS,NBE-1;BCS,CM-1 | 0.0149395 | 0.000304102 | 49.1264 |
| HZ014,SB;YH-1-1,CM-1 | 0.000160571 | 0.000183246 | 0.876256 |
| HZ014,YH-1-1;SB,CM-1 | 0.013985 | 0.000313338 | 44.6323 |
| LW-2,HZ104;XWY-3,CM-1 | 7.00E-05 | 0.000247478 | 0.282954 |
| LW-2,XWY-3;HZ104,CM-1 | 0.00211579 | 0.000270686 | 7.81639 |
| HZ112,HZ014;HZ061,CM-1 | 0.000928768 | 0.000233613 | 3.97567 |
| HZ112,HZ061;HZ014,CM-1 | 0.00459734 | 0.000257516 | 17.8526 |
| YH-1-1,DHL;HZ015,CM-1 | -0.00882852 | 0.000307276 | -28.7316 |
| YH-1-1,HZ015;DHL,CM-1 | -0.0087276 | 0.000304356 | -28.6757 |
| HZ045,GZ;LW-5,CM-1 | 0.00138005 | 0.000222977 | 6.1892 |
| HZ045,LW-5;GZ,CM-1 | 0.00452525 | 0.000256336 | 17.6536 |
| s6-8,HZ072;YNLDP2,CM-1 | -0.00032679 | 0.000219393 | -1.48952 |
| s6-8,YNLDP2;HZ072,CM-1 | 0.000842812 | 0.000229189 | 3.67736 |
| NBE-1,NBE-3;ST312,CM-1 | 8.21E-05 | 0.000201432 | 0.407595 |
| NBE-1,ST312;NBE-3,CM-1 | 0.0036307 | 0.000243734 | 14.8962 |
| XWY-5,HZ040;HZ058,CM-1 | -0.004686 | 0.000244355 | -19.1771 |
| XWY-5,HZ058;HZ040,CM-1 | -0.00341136 | 0.000260414 | -13.0997 |
| s301-1,HZ058;HZ122,CM-1 | 0.00163562 | 0.000249559 | 6.55404 |
| s301-1,HZ122;HZ058,CM-1 | 0.00325227 | 0.000268481 | 12.1136 |
| LC-1,HZ010;HZ064,CM-1 | -0.00264833 | 0.00023607 | -11.2184 |
| LC-1,HZ064;HZ010,CM-1 | -0.00142844 | 0.000246311 | -5.79936 |
| HZ004,GE-4;YH-1-1,CM-1 | -0.00160994 | 0.000231522 | -6.95372 |
| HZ004,YH-1-1;GE-4,CM-1 | 0.0101359 | 0.000355626 | 28.5015 |
| LW-3,HZ003;HZ006,CM-1 | -0.0162399 | 0.000338142 | -48.0269 |
| LW-3,HZ006;HZ003,CM-1 | -0.0162437 | 0.000333317 | -48.7335 |
| HZ011,HZ016;XWY-4,CM-1 | 0.00201267 | 0.000269916 | 7.45666 |
| HZ011,XWY-4;HZ016,CM-1 | 0.000636503 | 0.000252657 | 2.51924 |

| | | | |
|---|---|---|---|
| HZ071,HZ016;HZ122,CM-1 | -4.27E-05 | 0.000204067 | -0.209364 |
| HZ071,HZ122;HZ016,CM-1 | 0.0124342 | 0.000312352 | 39.8084 |
| HZ061,HZ060;ZW,CM-1 | -0.00885115 | 0.000292106 | -30.3012 |
| HZ061,ZW;HZ060,CM-1 | -0.00910657 | 0.000303214 | -30.0335 |
| HZ082,HZ015;HZ104,CM-1 | 0.000120022 | 0.000198005 | 0.606158 |
| HZ082,HZ104;HZ015,CM-1 | 0.0132402 | 0.000316772 | 41.7974 |
| HZ118,HZ001;HZ006,CM-1 | -0.0017549 | 0.000279893 | -6.2699 |
| HZ118,HZ006;HZ001,CM-1 | -0.00559883 | 0.000257194 | -21.7689 |
| GZ,GE-1;HZ039,CM-1 | -0.00029901 | 0.000218828 | -1.36639 |
| GZ,HZ039;GE-1,CM-1 | 0.00848766 | 0.000301885 | 28.1155 |
| HZ006,HZ063;XWY-3,CM-1 | 0.0048876 | 0.000261566 | 18.6859 |
| HZ006,XWY-3;HZ063,CM-1 | 0.000530055 | 0.000232312 | 2.28166 |
| HZ016,HZ057;YNLDP1,CM-1 | 0.00478188 | 0.000256595 | 18.6359 |
| HZ016,YNLDP1;HZ057,CM-1 | 0.0017732 | 0.000247573 | 7.16232 |
| YNZ-1,HZ001;LBDCS,CM-1 | -0.00747715 | 0.000270008 | -27.6924 |
| YNZ-1,LBDCS;HZ001,CM-1 | -0.00712812 | 0.000259961 | -27.42 |
| QXDM-2,YH-1-1;ZBD,CM-1 | 0.0123279 | 0.000310901 | 39.6523 |
| QXDM-2,ZBD;YH-1-1,CM-1 | 0.00072359 | 0.000196585 | 3.6808 |
| HZ071,HZ016;ST312,CM-1 | -0.00020072 | 0.000212531 | -0.944421 |
| HZ071,ST312;HZ016,CM-1 | 0.00985906 | 0.000302143 | 32.6304 |
| HZ114,HZ070;HZ094,CM-1 | 0.00553637 | 0.000232488 | 23.8136 |
| HZ114,HZ094;HZ070,CM-1 | 0.00463998 | 0.000228627 | 20.295 |
| QXDM-1,HZ112;HZ122,CM-1 | -0.00059982 | 0.000217099 | -2.76288 |
| QXDM-1,HZ122;HZ112,CM-1 | 0.00823276 | 0.00030423 | 27.0609 |
| HZ074,ASM;HZ125,CM-1 | -0.00213601 | 0.000222562 | -9.59739 |
| HZ074,HZ125;ASM,CM-1 | 0.00273968 | 0.000270452 | 10.13 |
| HZ063,HZ014;HZ073,CM-1 | -0.00779423 | 0.000232613 | -33.5073 |
| HZ063,HZ073;HZ014,CM-1 | -0.00621157 | 0.000241556 | -25.7149 |
| LBDCS,HZ076;HZ081,CM-1 | -0.002713 | 0.000260837 | -10.4011 |
| LBDCS,HZ081;HZ076,CM-1 | -0.00694014 | 0.000239713 | -28.9518 |
| HZ050,HZ080;LC-2,CM-1 | 0.000286114 | 0.000270231 | 1.05878 |
| HZ050,LC-2;HZ080,CM-1 | -0.00274908 | 0.000238134 | -11.5442 |
| GE-6,HZ104;NBE-4,CM-1 | 0.00196131 | 0.000306227 | 6.40475 |
| GE-6,NBE-4;HZ104,CM-1 | -0.00420411 | 0.000242188 | -17.3588 |
| HZ056,HZ021;HZ080,CM-1 | -0.00032543 | 0.000244808 | -1.32931 |
| HZ056,HZ080;HZ021,CM-1 | 0.000834331 | 0.000249404 | 3.34529 |
| XY-10,HZ064;HZ086,CM-1 | 0.000148209 | 0.000241417 | 0.613913 |
| XY-10,HZ086;HZ064,CM-1 | 0.000135955 | 0.000250799 | 0.542087 |
| HZ090,HZ021;LC-1,CM-1 | -0.00436724 | 0.000241061 | -18.1167 |
| HZ090,LC-1;HZ021,CM-1 | -0.00447497 | 0.000232249 | -19.268 |
| YNLDP1,LW-2;XWY-5,CM-1 | 0.0097128 | 0.000276838 | 35.0848 |
| YNLDP1,XWY-5;LW-2,CM-1 | 0.000265569 | 0.000193722 | 1.37088 |
| HZ109,HZ086;LW-4,CM-1 | 0.00560027 | 0.000271058 | 20.6608 |
| HZ109,LW-4;HZ086,CM-1 | 0.000782634 | 0.000228642 | 3.42296 |
| HZ031,HZ064;HZ123,CM-1 | -0.00030244 | 0.000197694 | -1.52985 |
| HZ031,HZ123;HZ064,CM-1 | 0.0129336 | 0.00030321 | 42.6558 |
| HZ119,HZ079;HZ114,CM-1 | 0.00487238 | 0.000202998 | 24.0021 |
| HZ119,HZ114;HZ079,CM-1 | 0.00789481 | 0.000216009 | 36.5486 |
| HZ072,HZ081;WLH-1,CM-1 | 0.00415862 | 0.000276641 | 15.0326 |
| HZ072,WLH-1;HZ081,CM-1 | -0.00041321 | 0.000218381 | -1.89213 |

| | | | |
|---|---|---|---|
| HZ114,HZ048;YNZ-1,CM-1 | -0.00480166 | 0.00026694 | -17.9878 |
| HZ114,YNZ-1;HZ048,CM-1 | -0.00654796 | 0.000248185 | -26.3834 |
| XYDCS,HZ008;QXDM-1,CM-1 | -0.0164738 | 0.000317572 | -51.8743 |
| XYDCS,QXDM-1;HZ008,CM-1 | -0.0160706 | 0.000312571 | -51.4142 |
| XWY-1,HZ021;HZ075,CM-1 | -0.00090479 | 0.000225928 | -4.00478 |
| XWY-1,HZ075;HZ021,CM-1 | 0.00412461 | 0.00028803 | 14.3201 |
| HZ008,HZ002;HZ074,CM-1 | -0.00317441 | 0.000238146 | -13.3297 |
| HZ008,HZ074;HZ002,CM-1 | 0.00189499 | 0.000274708 | 6.89819 |
| HZ059,ASM;QXDM-2,CM-1 | 0.00750745 | 0.000312134 | 24.052 |
| HZ059,QXDM-2;ASM,CM-1 | -8.72E-05 | 0.000240669 | -0.362245 |
| HZ048,LBDCS;XWY-1,CM-1 | 0.00765709 | 0.000256469 | 29.8558 |
| HZ048,XWY-1;LBDCS,CM-1 | 0.00254609 | 0.000231148 | 11.015 |
| WL,GE-8;s317,CM-1 | 0.000150454 | 0.000208178 | 0.72272 |
| WL,s317;GE-8,CM-1 | 0.0148522 | 0.000319884 | 46.4301 |
| HZ058,GE-6;HZ040,CM-1 | 0.00261831 | 0.00026324 | 9.94646 |
| HZ058,HZ040;GE-6,CM-1 | 5.68E-05 | 0.000229858 | 0.246926 |
| JX-4,HZ002;MZYCS,CM-1 | 0.00354085 | 0.000252798 | 14.0066 |
| JX-4,MZYCS;HZ002,CM-1 | -0.00114907 | 0.000232079 | -4.95119 |
| HZ112,DHL;HZ109,CM-1 | 0.000825241 | 0.000204891 | 4.02771 |
| HZ112,HZ109;DHL,CM-1 | 0.0101717 | 0.000291178 | 34.9329 |
| NBE-2,HZ095;ZW,CM-1 | 0.0183807 | 0.000356241 | 51.5963 |
| NBE-2,ZW;HZ095,CM-1 | 0.00114358 | 0.000206414 | 5.54025 |
| HZ086,GE-4;HZ100,CM-1 | -0.00015015 | 0.000199141 | -0.753997 |
| HZ086,HZ100;GE-4,CM-1 | 0.010176 | 0.000298368 | 34.1055 |
| HZ094,HZ028;WL,CM-1 | -0.0236846 | 0.000385507 | -61.4376 |
| HZ094,WL;HZ028,CM-1 | -0.0244042 | 0.000378157 | -64.5347 |
| HZ045,HZ008;HZ112,CM-1 | -0.00181027 | 0.000266801 | -6.78509 |
| HZ045,HZ112;HZ008,CM-1 | -0.00490536 | 0.000240717 | -20.3782 |
| LC-1,HZ043;NBE-2,CM-1 | -0.00110129 | 0.000227688 | -4.83685 |
| LC-1,NBE-2;HZ043,CM-1 | 0.00535188 | 0.000285497 | 18.7458 |
| HZ066,HZ054;SB,CM-1 | -0.0018831 | 0.000232556 | -8.09737 |
| HZ066,SB;HZ054,CM-1 | -0.00083035 | 0.000233524 | -3.55574 |
| ST330,HZ034;LBDCS,CM-1 | -0.00578066 | 0.00024628 | -23.4719 |
| ST330,LBDCS;HZ034,CM-1 | 0.000513044 | 0.000284393 | 1.804 |
| LW-3,HZ057;HZ119,CM-1 | 0.00106927 | 0.000249016 | 4.29399 |
| LW-3,HZ119;HZ057,CM-1 | -0.0028765 | 0.000224091 | -12.8363 |
| LC-1,HZ123;ST330,CM-1 | 0.0004671 | 0.000259033 | 1.80324 |
| LC-1,ST330;HZ123,CM-1 | -0.00372938 | 0.000223535 | -16.6836 |
| JGL,HZ078;HZ082,CM-1 | 0.00120272 | 0.000252996 | 4.75391 |
| JGL,HZ082;HZ078,CM-1 | 0.000376274 | 0.000237027 | 1.58748 |
| XWY-3,HZ058;s1-16,CM-1 | -0.00072158 | 0.000227708 | -3.1689 |
| XWY-3,s1-16;HZ058,CM-1 | 0.00922955 | 0.000316811 | 29.1327 |
| HZ084,HZ003;ZW,CM-1 | -0.0107097 | 0.000316145 | -33.876 |
| HZ084,ZW;HZ003,CM-1 | -0.0103984 | 0.000304917 | -34.1024 |
| HZ060,HZ027;HZ037,CM-1 | 0.012011 | 0.000288865 | 41.58 |
| HZ060,HZ037;HZ027,CM-1 | 4.76E-05 | 0.000203677 | 0.233834 |
| s1-16,HZ068;HZ124,CM-1 | 0.00259159 | 0.000231887 | 11.1761 |
| s1-16,HZ124;HZ068,CM-1 | 0.00214186 | 0.000230489 | 9.29266 |
| LC-1,HZ004;HZ124,CM-1 | 1.51E-05 | 0.000197511 | 0.0761991 |
| LC-1,HZ124;HZ004,CM-1 | 0.014239 | 0.000301523 | 47.2234 |

| | | | |
|---|---|---|---|
| HZ069,HZ063;HZ125,CM-1 | -0.00034132 | 0.000184819 | -1.84678 |
| HZ069,HZ125;HZ063,CM-1 | 0.0121108 | 0.000286848 | 42.2202 |
| XWY-2,DBZ-1;NC,CM-1 | 0.00113727 | 0.000228456 | 4.97807 |
| XWY-2,NC;DBZ-1,CM-1 | 0.00622106 | 0.000244652 | 25.4282 |
| HZ124,HZ018;JGL,CM-1 | -0.0159922 | 0.00030771 | -51.9718 |
| HZ124,JGL;HZ018,CM-1 | -0.0157569 | 0.000311835 | -50.5297 |
| HZ016,JX-4;XH-1,CM-1 | -0.000146 | 0.000233874 | -0.624251 |
| HZ016,XH-1;JX-4,CM-1 | 0.00391648 | 0.000266045 | 14.7211 |
| HZ061,LW-4;LW-5,CM-1 | -0.00695002 | 0.000282795 | -24.5762 |
| HZ061,LW-5;LW-4,CM-1 | -0.00729191 | 0.000274645 | -26.5503 |
| HZ068,HZ039;WL,CM-1 | 0.0123927 | 0.000326333 | 37.9757 |
| HZ068,WL;HZ039,CM-1 | 0.000253291 | 0.000204165 | 1.24062 |
| HZ043,HZ001;HZ036,CM-1 | 0.0118376 | 0.000313951 | 37.7051 |
| HZ043,HZ036;HZ001,CM-1 | 0.00055502 | 0.000210146 | 2.64111 |
| DBZ-1,WL;s317,CM-1 | -0.00111589 | 0.000218945 | -5.09669 |
| DBZ-1,s317;WL,CM-1 | 0.00886697 | 0.000302057 | 29.3553 |
| HZ090,FL;HZ058,CM-1 | -0.0039816 | 0.000273795 | -14.5423 |
| HZ090,HZ058;FL,CM-1 | -0.00395296 | 0.000271257 | -14.5728 |
| GE-1,NC;ZW,CM-1 | 0.0108109 | 0.000281502 | 38.4042 |
| GE-1,ZW;NC,CM-1 | -0.0004503 | 0.000207593 | -2.16916 |
| NBE-1,HZ083;YNLDP1,CM-1 | -0.0031648 | 0.000268679 | -11.7791 |
| NBE-1,YNLDP1;HZ083,CM-1 | -0.002476 | 0.000268363 | -9.22629 |
| HZ122,HZ068;NC,CM-1 | -0.00563323 | 0.000259189 | -21.7341 |
| HZ122,NC;HZ068,CM-1 | -0.00209832 | 0.000264123 | -7.94449 |
| XWY-4,HZ006;XY-10,CM-1 | -0.00091662 | 0.000242247 | -3.78384 |
| XWY-4,XY-10;HZ006,CM-1 | -0.00012777 | 0.000242738 | -0.526375 |
| HZ010,HZ001;HZ027,CM-1 | -0.0028116 | 0.000251015 | -11.2009 |
| HZ010,HZ027;HZ001,CM-1 | -0.00283135 | 0.000247082 | -11.4592 |
| NBE-2,HZ040;HZ117,CM-1 | 0.00039983 | 0.000232487 | 1.7198 |
| NBE-2,HZ117;HZ040,CM-1 | 0.00433097 | 0.000281714 | 15.3736 |
| DHL,HZ083;ZW,CM-1 | 0.00337899 | 0.000277392 | 12.1813 |
| DHL,ZW;HZ083,CM-1 | -0.00098176 | 0.000224705 | -4.36909 |
| HZ112,DHL;FL,CM-1 | -0.00581291 | 0.000259125 | -22.4328 |
| HZ112,FL;DHL,CM-1 | -0.00535953 | 0.000257508 | -20.8131 |
| HZ003,HZ028;YNLDP2,CM-1 | 0.00103536 | 0.000200576 | 5.16193 |
| HZ003,YNLDP2;HZ028,CM-1 | 0.0116133 | 0.000289879 | 40.0627 |
| XYDCS,HZ034;YNZ-1,CM-1 | -0.00526255 | 0.000243169 | -21.6415 |
| XYDCS,YNZ-1;HZ034,CM-1 | 0.00124223 | 0.00029064 | 4.27412 |
| ST312,HZ071;HZ117,CM-1 | 0.00062088 | 0.000259978 | 2.3882 |
| ST312,HZ117;HZ071,CM-1 | -0.00046787 | 0.000249542 | -1.8749 |
| HZ069,HZ112;XYDCS,CM-1 | -0.00368947 | 0.000230984 | -15.9729 |
| HZ069,XYDCS;HZ112,CM-1 | 0.00541051 | 0.000296875 | 18.2249 |
| HZ092,HZ068;HZ118,CM-1 | -0.00136152 | 0.000235684 | -5.77689 |
| HZ092,HZ118;HZ068,CM-1 | 0.00126785 | 0.000260459 | 4.86778 |
| HZ010,HSKC;HZ100,CM-1 | 0.00124138 | 0.000206668 | 6.00665 |
| HZ010,HZ100;HSKC,CM-1 | 0.0116756 | 0.000302268 | 38.6267 |
| HZ059,HZ081;XSX,CM-1 | -0.00236269 | 0.000255058 | -9.26333 |
| HZ059,XSX;HZ081,CM-1 | -0.00037035 | 0.000284031 | -1.30391 |
| FL,HZ006;NC,CM-1 | -0.00034755 | 0.00020379 | -1.70541 |
| FL,NC;HZ006,CM-1 | 0.0113709 | 0.000272087 | 41.7914 |

| | | | |
|---|---|---|---|
| XSL,HZ008;HZ074,CM-1 | -0.00039281 | 0.000211027 | -1.86143 |
| XSL,HZ074;HZ008,CM-1 | 0.0108691 | 0.000310624 | 34.9911 |
| NBE-3,HZ041;HZ118,CM-1 | 0.00225537 | 0.000229245 | 9.83825 |
| NBE-3,HZ118;HZ041,CM-1 | 0.00791286 | 0.000276315 | 28.6371 |
| XWY-1,HZ117;XWY-2,CM-1 | 0.0138199 | 0.000307942 | 44.8783 |
| XWY-1,XWY-2;HZ117,CM-1 | 0.000413475 | 0.000188183 | 2.1972 |
| HZ036,HZ021;HZ083,CM-1 | 0.00169277 | 0.000239786 | 7.0595 |
| HZ036,HZ083;HZ021,CM-1 | 0.00287419 | 0.000248431 | 11.5694 |
| HZ060,JHZS;XWY-4,CM-1 | -0.00627154 | 0.000258364 | -24.274 |
| HZ060,XWY-4;JHZS,CM-1 | -0.00553709 | 0.00025983 | -21.3104 |
| HZ010,HZ013;YNLDP1,CM-1 | 0.00248629 | 0.000232086 | 10.7128 |
| HZ010,YNLDP1;HZ013,CM-1 | 0.00511083 | 0.000257711 | 19.8316 |
| XSX,GE-1;YNZ-1,CM-1 | 0.000872964 | 0.000205387 | 4.25033 |
| XSX,YNZ-1;GE-1,CM-1 | 0.0129987 | 0.000309503 | 41.9987 |
| HZ110,HZ059;HZ109,CM-1 | -0.00417432 | 0.0002577 | -16.1983 |
| HZ110,HZ109;HZ059,CM-1 | -0.00350113 | 0.000270678 | -12.9347 |
| HZ092,HZ036;HZ124,CM-1 | -0.00065905 | 0.000250501 | -2.63092 |
| HZ092,HZ124;HZ036,CM-1 | 0.00228877 | 0.000268157 | 8.53519 |
| GLJYD,HZ036;SLLK,CM-1 | -0.00020108 | 0.000217608 | -0.924032 |
| GLJYD,SLLK;HZ036,CM-1 | 0.00594679 | 0.000276182 | 21.5322 |
| HZ117,HZ094;s303-231,CM-1 | 0.0129131 | 0.000305817 | 42.225 |
| HZ117,s303-231;HZ094,CM-1 | 7.13E-05 | 0.000183842 | 0.387608 |
| HZ090,LC-1;YNLDP2,CM-1 | 0.00179888 | 0.000216235 | 8.31908 |
| HZ090,YNLDP2;LC-1,CM-1 | 0.00354216 | 0.000229236 | 15.452 |
| MZYCS,HZ036;QXDM-1,CM-1 | -0.00476021 | 0.000271934 | -17.505 |
| MZYCS,QXDM-1;HZ036,CM-1 | -0.00400778 | 0.00027857 | -14.387 |
| ZBD,HZ059;s6-8,CM-1 | 0.000584238 | 0.000199765 | 2.92462 |
| ZBD,s6-8;HZ059,CM-1 | 0.00834168 | 0.000247534 | 33.6991 |
| HZ048,HZ057;HZ085,CM-1 | 0.00115265 | 0.000284704 | 4.0486 |
| HZ048,HZ085;HZ057,CM-1 | -0.00227825 | 0.000253344 | -8.99269 |
| HZ075,HZ048;MZYCS,CM-1 | -0.00485497 | 0.000267192 | -18.1704 |
| HZ075,MZYCS;HZ048,CM-1 | -0.00526203 | 0.000265981 | -19.7834 |
| GZ,HZ013;MZYCS,CM-1 | -0.00114284 | 0.000260766 | -4.38265 |
| GZ,MZYCS;HZ013,CM-1 | -0.00307557 | 0.000250643 | -12.2708 |
| HZ083,HZ090;LW-3,CM-1 | -0.00086746 | 0.000206424 | -4.20231 |
| HZ083,LW-3;HZ090,CM-1 | 0.00918955 | 0.00029034 | 31.6509 |
| XSX,HZ003;NBE-4,CM-1 | 0.00016402 | 0.000212811 | 0.770732 |
| XSX,NBE-4;HZ003,CM-1 | 0.00955717 | 0.000305966 | 31.2361 |
| HZ110,LW-1;NBE-2,CM-1 | -0.00265123 | 0.000273078 | -9.70869 |
| HZ110,NBE-2;LW-1,CM-1 | -0.00390607 | 0.000259168 | -15.0716 |
| YNLDP1,HZ057;NBE-1,CM-1 | 0.00353964 | 0.000250516 | 14.1294 |
| YNLDP1,NBE-1;HZ057,CM-1 | 0.00234605 | 0.000252331 | 9.29751 |
| HZ008,HZ066;SB,CM-1 | 0.000648817 | 0.000222712 | 2.91325 |
| HZ008,SB;HZ066,CM-1 | 0.00261949 | 0.000239079 | 10.9566 |
| HZ063,HZ076;HZ100,CM-1 | 0.00206188 | 0.000213624 | 9.65193 |
| HZ063,HZ100;HZ076,CM-1 | 0.00768051 | 0.000262282 | 29.2835 |
| ZBD,HZ039;ST312,CM-1 | 0.00362606 | 0.000240294 | 15.0901 |
| ZBD,ST312;HZ039,CM-1 | 0.00269702 | 0.000235646 | 11.4452 |
| HZ011,HZ006;HZ056,CM-1 | -0.00027483 | 0.000219165 | -1.25397 |
| HZ011,HZ056;HZ006,CM-1 | 0.00821642 | 0.000272039 | 30.2031 |

| | | | |
|---|---|---|---|
| SB,HZ082;YH-1-1,CM-1 | -0.00012217 | 0.000202334 | -0.603803 |
| SB,YH-1-1;HZ082,CM-1 | 0.0112059 | 0.000303417 | 36.9323 |
| HZ114,HZ008;HZ066,CM-1 | -0.0200677 | 0.00033395 | -60.092 |
| HZ114,HZ066;HZ008,CM-1 | -0.0199321 | 0.000332434 | -59.9582 |
| HZ068,GE-4;HZ054,CM-1 | 0.00210079 | 0.000277759 | 7.56334 |
| HZ068,HZ054;GE-4,CM-1 | -0.00092655 | 0.00023861 | -3.88311 |
| HZ095,HZ036;HZ094,CM-1 | 0.0155694 | 0.000308426 | 50.48 |
| HZ095,HZ094;HZ036,CM-1 | -0.00041682 | 0.000174537 | -2.38814 |
| DBZ-1,HZ118;ZW,CM-1 | 0.0105972 | 0.000317813 | 33.3442 |
| DBZ-1,ZW;HZ118,CM-1 | -0.00059622 | 0.000218982 | -2.72268 |
| XWY-2,HZ040;HZ081,CM-1 | -0.00344933 | 0.000235909 | -14.6215 |
| XWY-2,HZ081;HZ040,CM-1 | -0.00220592 | 0.000253855 | -8.6897 |
| HZ008,HZ056;QXDM-2,CM-1 | 0.0110649 | 0.000284634 | 38.8741 |
| HZ008,QXDM-2;HZ056,CM-1 | -8.51E-05 | 0.000195414 | -0.435592 |
| GE-6,HZ112;HZ114,CM-1 | 0.000182141 | 0.000177955 | 1.02353 |
| GE-6,HZ114;HZ112,CM-1 | 0.0177533 | 0.000328933 | 53.9724 |
| HZ080,HZ104;s6-8,CM-1 | -0.00177617 | 0.000276108 | -6.43287 |
| HZ080,s6-8;HZ104,CM-1 | -0.00505288 | 0.000238199 | -21.2129 |
| HZ095,HZ013;HZ045,CM-1 | -0.0157144 | 0.000322387 | -48.7438 |
| HZ095,HZ045;HZ013,CM-1 | -0.01725 | 0.00030692 | -56.2036 |
| DHL,HZ069;LW-1,CM-1 | -0.00032256 | 0.000203884 | -1.58209 |
| DHL,LW-1;HZ069,CM-1 | 0.0135001 | 0.000331373 | 40.7398 |
| NBE-1,HZ014;HZ059,CM-1 | -0.00733889 | 0.000286376 | -25.6267 |
| NBE-1,HZ059;HZ014,CM-1 | -0.00595677 | 0.000293155 | -20.3195 |
| MZYCS,HZ071;HZ084,CM-1 | 3.21E-05 | 0.000207506 | 0.154549 |
| MZYCS,HZ084;HZ071,CM-1 | 0.0102346 | 0.000284427 | 35.9834 |
| NBE-2,DBZ-1;HZ065,CM-1 | -0.0007373 | 0.000257018 | -2.86866 |
| NBE-2,HZ065;DBZ-1,CM-1 | -0.00119185 | 0.000246321 | -4.83862 |
| HZ011,HZ079;s1-16,CM-1 | -0.00046044 | 0.000208113 | -2.21245 |
| HZ011,s1-16;HZ079,CM-1 | 0.00814497 | 0.000275051 | 29.6126 |
| XYDCS,HSKC;HZ074,CM-1 | -0.002958 | 0.000235829 | -12.543 |
| XYDCS,HZ074;HSKC,CM-1 | 0.00245008 | 0.000281327 | 8.70899 |
| HZ054,HZ037;HZ085,CM-1 | 0.00200926 | 0.000232762 | 8.63222 |
| HZ054,HZ085;HZ037,CM-1 | 0.00306546 | 0.000252399 | 12.1453 |
| HZ119,HZ006;HZ117,CM-1 | 0.000487666 | 0.000246524 | 1.97817 |
| HZ119,HZ117;HZ006,CM-1 | 0.0014692 | 0.000248291 | 5.91728 |
| HZ003,XWY-4;s317,CM-1 | -3.84E-06 | 0.000197092 | -0.0195055 |
| HZ003,s317;XWY-4,CM-1 | 0.0143636 | 0.000330867 | 43.4121 |
| GZ,HZ048;HZ073,CM-1 | 0.00109326 | 0.000252687 | 4.32653 |
| GZ,HZ073;HZ048,CM-1 | -0.00116321 | 0.000240339 | -4.83988 |
| YNLDP2,HZ095;XWY-5,CM-1 | 0.0141596 | 0.000300265 | 47.1569 |
| YNLDP2,XWY-5;HZ095,CM-1 | 0.000279049 | 0.000195143 | 1.42998 |
| HZ079,HZ021;HZ086,CM-1 | -0.00345759 | 0.00025825 | -13.3885 |
| HZ079,HZ086;HZ021,CM-1 | -0.00386936 | 0.000251303 | -15.3972 |
| XY-10,GE-1;HZ011,CM-1 | 0.00230635 | 0.00026493 | 8.70551 |
| XY-10,HZ011;GE-1,CM-1 | 0.000407291 | 0.000250539 | 1.62566 |
| GE-8,HZ041;s6-8,CM-1 | -0.0011321 | 0.000226175 | -5.00542 |
| GE-8,s6-8;HZ041,CM-1 | 0.00631495 | 0.000271665 | 23.2453 |
| HZ013,GZ;HZ050,CM-1 | 8.82E-05 | 0.000239752 | 0.367813 |
| HZ013,HZ050;GZ,CM-1 | -0.00049602 | 0.000244473 | -2.02892 |

| | | | |
|---|---|---|---|
| HZ078,HZ014;XWY-3,CM-1 | -0.00335438 | 0.000263079 | -12.7505 |
| HZ078,XWY-3;HZ014,CM-1 | -0.00286944 | 0.000271826 | -10.5562 |
| QXDM-1,HZ122;LC-2,CM-1 | 0.00914608 | 0.000284666 | 32.1291 |
| QXDM-1,LC-2;HZ122,CM-1 | -0.00013346 | 0.000212244 | -0.628784 |
| YNLDP1,HZ008;HZ009,CM-1 | -0.00387698 | 0.000254437 | -15.2375 |
| YNLDP1,HZ009;HZ008,CM-1 | -0.00214294 | 0.000278825 | -7.68561 |
| HZ095,HZ050;HZ088,CM-1 | -0.0207938 | 0.000361412 | -57.5347 |
| HZ095,HZ088;HZ050,CM-1 | -0.0209144 | 0.000361271 | -57.8912 |
| LC-1,HZ114;XY-10,CM-1 | 0.0187436 | 0.000322803 | 58.065 |
| LC-1,XY-10;HZ114,CM-1 | -0.00054725 | 0.000174544 | -3.13531 |
| HZ077,SB;ST330,CM-1 | 0.000495307 | 0.00019444 | 2.54735 |
| HZ077,ST330;SB,CM-1 | 0.0109335 | 0.000286348 | 38.1827 |
| HZ064,LBDCS;QXDM-2,CM-1 | 0.014132 | 0.000297279 | 47.5379 |
| HZ064,QXDM-2;LBDCS,CM-1 | -0.00035858 | 0.000179344 | -1.99939 |
| HZ057,HZ066;HZ090,CM-1 | -0.00308069 | 0.000232033 | -13.277 |
| HZ057,HZ090;HZ066,CM-1 | 0.00098678 | 0.000273506 | 3.60789 |
| ASM,HZ065;QXDM-2,CM-1 | -0.00634371 | 0.000293998 | -21.5774 |
| ASM,QXDM-2;HZ065,CM-1 | -0.00767955 | 0.000283843 | -27.0556 |
| HZ057,LW-2;NBE-5,CM-1 | 0.00340838 | 0.00027348 | 12.463 |
| HZ057,NBE-5;LW-2,CM-1 | -0.00225524 | 0.000236194 | -9.54825 |
| HZ063,GE-6;HZ104,CM-1 | -0.00018339 | 0.000200849 | -0.913057 |
| HZ063,HZ104;GE-6,CM-1 | 0.0118697 | 0.000315572 | 37.6133 |
| HZ076,HZ004;HZ094,CM-1 | 0.000776212 | 0.000196941 | 3.94135 |
| HZ076,HZ094;HZ004,CM-1 | 0.0139429 | 0.000306815 | 45.4441 |
| HZ002,BCS;HZ118,CM-1 | -8.87E-06 | 0.000215551 | -0.041129 |
| HZ002,HZ118;BCS,CM-1 | 0.00905376 | 0.000290501 | 31.1661 |
| HZ002,HZ071;HZ088,CM-1 | -0.00354796 | 0.000240899 | -14.728 |
| HZ002,HZ088;HZ071,CM-1 | 8.69E-05 | 0.000270467 | 0.321303 |
| HZ083,LW-1;XWY-4,CM-1 | 0.0142098 | 0.000332787 | 42.6995 |
| HZ083,XWY-4;LW-1,CM-1 | -0.00012124 | 0.000209685 | -0.578174 |
| HZ017,JHZS;LW-1,CM-1 | -0.00026172 | 0.000198131 | -1.32094 |
| HZ017,LW-1;JHZS,CM-1 | 0.0149252 | 0.000344603 | 43.3114 |
| LW-3,HZ095;XYDCS,CM-1 | -0.0008985 | 0.00027728 | -3.24041 |
| LW-3,XYDCS;HZ095,CM-1 | -0.00608562 | 0.00025091 | -24.2542 |
| HZ003,HZ015;HZ117,CM-1 | -0.0007323 | 0.000230582 | -3.17587 |
| HZ003,HZ117;HZ015,CM-1 | 0.00531485 | 0.000283209 | 18.7665 |
| HZ061,HZ073;XSL,CM-1 | -0.00781004 | 0.000271285 | -28.7891 |
| HZ061,XSL;HZ073,CM-1 | -0.00766359 | 0.000271877 | -28.1877 |
| HZ059,HZ020;LW-1,CM-1 | -2.13E-05 | 0.000230825 | -0.0923717 |
| HZ059,LW-1;HZ020,CM-1 | 0.0101337 | 0.000319816 | 31.6859 |
| FL,HZ016;HZ084,CM-1 | 0.00051517 | 0.000219057 | 2.35176 |
| FL,HZ084;HZ016,CM-1 | 0.00928295 | 0.000295647 | 31.3988 |
| HZ063,HZ003;HZ048,CM-1 | 6.95E-05 | 0.000249853 | 0.278019 |
| HZ063,HZ048;HZ003,CM-1 | -0.00064586 | 0.000248875 | -2.59512 |
| HZ011,BCS;HZ124,CM-1 | -0.00045428 | 0.000202631 | -2.24192 |
| HZ011,HZ124;BCS,CM-1 | 0.0156824 | 0.000323588 | 48.4642 |
| SLLK,HZ092;NBE-2,CM-1 | 0.00354796 | 0.000271954 | 13.0462 |
| SLLK,NBE-2;HZ092,CM-1 | -0.00058584 | 0.00023588 | -2.48362 |
| ST312,HZ028;HZ080,CM-1 | -0.00508547 | 0.000264456 | -19.2299 |
| ST312,HZ080;HZ028,CM-1 | -0.00427749 | 0.000264877 | -16.149 |

| | | | |
|---|---|---|---|
| HZ045,HZ041;XWY-2,CM-1 | -0.00599931 | 0.000269448 | -22.2652 |
| HZ045,XWY-2;HZ041,CM-1 | -0.00558986 | 0.000269376 | -20.7511 |
| HZ109,HZ123;WLH-1,CM-1 | -0.00484119 | 0.000234686 | -20.6284 |
| HZ109,WLH-1;HZ123,CM-1 | -0.00186423 | 0.000268667 | -6.9388 |
| HZ083,HZ050;HZ094,CM-1 | -0.00010254 | 0.000173249 | -0.591885 |
| HZ083,HZ094;HZ050,CM-1 | 0.0226723 | 0.000379686 | 59.7133 |
| XSL,DHL;HZ008,CM-1 | -0.00018349 | 0.000229268 | -0.800334 |
| XSL,HZ008;DHL,CM-1 | 0.00261577 | 0.000267732 | 9.7701 |
| HZ041,HZ001;HZ100,CM-1 | 0.0042673 | 0.000239401 | 17.8249 |
| HZ041,HZ100;HZ001,CM-1 | 0.00485975 | 0.00024462 | 19.8665 |
| SLLK,HZ013;HZ065,CM-1 | -0.0028484 | 0.000235984 | -12.0703 |
| SLLK,HZ065;HZ013,CM-1 | -0.00114328 | 0.00026463 | -4.32029 |
| HZ008,HZ063;HZ114,CM-1 | 0.000104045 | 0.000187821 | 0.553959 |
| HZ008,HZ114;HZ063,CM-1 | 0.0145845 | 0.000295916 | 49.2859 |
| LW-4,LW-1;NC,CM-1 | -0.00030219 | 0.000188758 | -1.60094 |
| LW-4,NC;LW-1,CM-1 | 0.0121181 | 0.000289056 | 41.923 |
| NBE-1,HZ125;JGL,CM-1 | 0.0102412 | 0.000268506 | 38.1415 |
| NBE-1,JGL;HZ125,CM-1 | 0.00158649 | 0.000204201 | 7.76926 |
| HZ071,XWY-2;XWY-3,CM-1 | -0.00044791 | 0.00028272 | -1.58428 |
| HZ071,XWY-3;XWY-2,CM-1 | -0.00332502 | 0.000249701 | -13.316 |
| XWY-1,HZ124;WLH-1,CM-1 | 0.00506544 | 0.000254952 | 19.8682 |
| XWY-1,WLH-1;HZ124,CM-1 | -0.00016873 | 0.000212471 | -0.79411 |
| HZ123,HZ068;HZ076,CM-1 | -0.00533045 | 0.000254822 | -20.9183 |
| HZ123,HZ076;HZ068,CM-1 | -0.00357356 | 0.00026809 | -13.3297 |
| HZ048,HZ072;WL,CM-1 | 0.00917131 | 0.000295148 | 31.0736 |
| HZ048,WL;HZ072,CM-1 | 6.74E-05 | 0.000219317 | 0.307281 |
| HZ020,GE-4;XWY-5,CM-1 | -7.43E-05 | 0.000253001 | -0.293495 |
| HZ020,XWY-5;GE-4,CM-1 | 0.000406417 | 0.000263799 | 1.54063 |
| GE-6,HZ054;HZ125,CM-1 | 0.000762474 | 0.000194236 | 3.92551 |
| GE-6,HZ125;HZ054,CM-1 | 0.0169247 | 0.000321532 | 52.6377 |
| HZ039,DBZ-1;HZ119,CM-1 | -0.00369164 | 0.000241376 | -15.2942 |
| HZ039,HZ119;DBZ-1,CM-1 | -0.00248132 | 0.00026087 | -9.51172 |
| HZ109,DHL;HZ082,CM-1 | -0.00906967 | 0.000312024 | -29.0672 |
| HZ109,HZ082;DHL,CM-1 | -0.0101127 | 0.000299158 | -33.8039 |
| DHL,XWY-1;XYDCS,CM-1 | 0.00171823 | 0.000218602 | 7.86007 |
| DHL,XYDCS;XWY-1,CM-1 | 0.0110701 | 0.0002832 | 39.0894 |
| GE-6,HZ094;HZ119,CM-1 | 0.0131328 | 0.000311518 | 42.1575 |
| GE-6,HZ119;HZ094,CM-1 | -0.00082237 | 0.000193151 | -4.25764 |
| HZ040,HZ009;SB,CM-1 | 0.00502184 | 0.000247726 | 20.2718 |
| HZ040,SB;HZ009,CM-1 | 0.000515644 | 0.000191299 | 2.69549 |
| XWY-1,HZ006;HZ125,CM-1 | 0.000873837 | 0.000201314 | 4.34067 |
| XWY-1,HZ125;HZ006,CM-1 | 0.0139346 | 0.000316181 | 44.0717 |
| HZ041,JGL;ZW,CM-1 | -0.0054818 | 0.000261585 | -20.9561 |
| HZ041,ZW;JGL,CM-1 | -0.0061505 | 0.000252971 | -24.3131 |
| HZ013,HZ082;s6-8,CM-1 | -0.00098039 | 0.000213548 | -4.59098 |
| HZ013,s6-8;HZ082,CM-1 | 0.00713559 | 0.000274598 | 25.9855 |
| HZ090,JGL;WLH-1,CM-1 | 0.00350237 | 0.000211366 | 16.5701 |
| HZ090,WLH-1;JGL,CM-1 | 0.00709514 | 0.000244335 | 29.0386 |
| NBE-4,HZ037;HZ119,CM-1 | -0.00038151 | 0.000227025 | -1.68046 |
| NBE-4,HZ119;HZ037,CM-1 | 0.00208212 | 0.000253905 | 8.20039 |

| | | | |
|---|---|---|---|
| HZ123,HZ016;HZ020,CM-1 | -0.0107998 | 0.000317202 | -34.047 |
| HZ123,HZ020;HZ016,CM-1 | -0.0111516 | 0.000310233 | -35.9458 |
| HZ069,HZ037;XWY-3,CM-1 | 0.000344911 | 0.000225568 | 1.52907 |
| HZ069,XWY-3;HZ037,CM-1 | 0.0057009 | 0.000268368 | 21.2428 |
| s6-8,HZ002;HZ059,CM-1 | -0.00349531 | 0.000249046 | -14.0348 |
| s6-8,HZ059;HZ002,CM-1 | -0.00411912 | 0.000251736 | -16.3628 |
| XH-1,HZ109;JX-4,CM-1 | 0.00545067 | 0.000266818 | 20.4284 |
| XH-1,JX-4;HZ109,CM-1 | -5.76E-05 | 0.000224156 | -0.25711 |
| HZ008,HZ064;JGL,CM-1 | -4.12E-05 | 0.000199552 | -0.206375 |
| HZ008,JGL;HZ064,CM-1 | 0.00565193 | 0.00024335 | 23.2255 |
| HZ070,HZ114;MZYCS,CM-1 | 0.018068 | 0.000336279 | 53.7293 |
| HZ070,MZYCS;HZ114,CM-1 | -4.75E-05 | 0.000190486 | -0.249508 |
| HZ109,GE-4;HZ081,CM-1 | -0.00949539 | 0.00030568 | -31.0631 |
| HZ109,HZ081;GE-4,CM-1 | -0.00981974 | 0.000309219 | -31.7566 |
| QXDM-2,HZ001;HZ119,CM-1 | 0.00667613 | 0.000256271 | 26.051 |
| QXDM-2,HZ119;HZ001,CM-1 | 0.00339012 | 0.000222438 | 15.2408 |
| FL,HZ003;XH-1,CM-1 | -0.00087143 | 0.000204082 | -4.27001 |
| FL,XH-1;HZ003,CM-1 | 0.00680632 | 0.00026384 | 25.7971 |
| s6-8,HZ041;HZ054,CM-1 | -0.00803937 | 0.000274257 | -29.3133 |
| s6-8,HZ054;HZ041,CM-1 | -0.00745328 | 0.000284123 | -26.2326 |
| HZ064,HZ002;HZ036,CM-1 | 0.00797014 | 0.000268368 | 29.6985 |
| HZ064,HZ036;HZ002,CM-1 | 9.30E-05 | 0.000206647 | 0.450075 |
| NBE-3,HZ008;HZ018,CM-1 | -0.00613902 | 0.000259809 | -23.629 |
| NBE-3,HZ018;HZ008,CM-1 | -0.00500112 | 0.00027434 | -18.2296 |
| JGL,HZ008;HZ112,CM-1 | 0.000247032 | 0.000232763 | 1.0613 |
| JGL,HZ112;HZ008,CM-1 | 0.00280739 | 0.000234634 | 11.965 |
| SB,BCS;s1-16,CM-1 | -0.00034408 | 0.000143488 | -2.398 |
| SB,s1-16;BCS,CM-1 | 0.0181906 | 0.000299034 | 60.8313 |
| HZ020,HZ016;JHZS,CM-1 | -0.00101777 | 0.000263811 | -3.85795 |
| HZ020,JHZS;HZ016,CM-1 | -0.00178437 | 0.000250388 | -7.12644 |
| HZ125,HZ074;SB,CM-1 | -0.00608222 | 0.000269713 | -22.5507 |
| HZ125,SB;HZ074,CM-1 | -0.00647851 | 0.000250455 | -25.867 |
| HZ077,JHZS;NBE-2,CM-1 | -0.00038308 | 0.000224703 | -1.70483 |
| HZ077,NBE-2;JHZS,CM-1 | 0.00746693 | 0.000308031 | 24.2409 |
| GE-1,HZ028;HZ060,CM-1 | -0.00283785 | 0.000258263 | -10.9882 |
| GE-1,HZ060;HZ028,CM-1 | -0.00323334 | 0.000260511 | -12.4115 |
| HZ010,HZ063;LW-2,CM-1 | -0.00028503 | 0.000200891 | -1.41884 |
| HZ010,LW-2;HZ063,CM-1 | 0.0107006 | 0.000305066 | 35.0762 |
| GE-1,HZ021;ZW,CM-1 | 0.00223809 | 0.000265826 | 8.41936 |
| GE-1,ZW;HZ021,CM-1 | -0.00109477 | 0.000242498 | -4.51453 |
| HZ122,HZ070;LC-1,CM-1 | -0.0107153 | 0.00029302 | -36.5683 |
| HZ122,LC-1;HZ070,CM-1 | -0.0104544 | 0.000289106 | -36.1612 |
| YNLDP2,HZ004;LW-4,CM-1 | 0.00179121 | 0.000243847 | 7.34564 |
| YNLDP2,LW-4;HZ004,CM-1 | 0.00436044 | 0.000261615 | 16.6674 |
| HZ061,NBE-1;YH-1-1,CM-1 | -0.00558021 | 0.000263485 | -21.1785 |
| HZ061,YH-1-1;NBE-1,CM-1 | -0.00342354 | 0.000270215 | -12.6697 |
| GE-8,HZ066;HZ075,CM-1 | 7.52E-05 | 0.000215001 | 0.349917 |
| GE-8,HZ075;HZ066,CM-1 | 0.00735401 | 0.000276164 | 26.6291 |
| HZ031,HZ080;HZ088,CM-1 | 0.00128733 | 0.000278889 | 4.61591 |
| HZ031,HZ088;HZ080,CM-1 | -0.00176573 | 0.000238255 | -7.41108 |

| | | | |
|---|---|---|---|
| HZ057,HZ119;s301-1,CM-1 | -0.00282443 | 0.000232022 | -12.1731 |
| HZ057,s301-1;HZ119,CM-1 | -0.00255083 | 0.000231493 | -11.019 |
| HZ118,HZ017;HZ082,CM-1 | -0.0115091 | 0.000323462 | -35.5809 |
| HZ118,HZ082;HZ017,CM-1 | -0.0118419 | 0.000319616 | -37.0504 |
| HZ040,HZ117;HZ122,CM-1 | -0.0110262 | 0.000274212 | -40.2105 |
| HZ040,HZ122;HZ117,CM-1 | -0.00845545 | 0.000293412 | -28.8177 |
| s1-16,HZ045;YNLDP2,CM-1 | -0.00150104 | 0.000211774 | -7.08793 |
| s1-16,YNLDP2;HZ045,CM-1 | -0.00122599 | 0.000225445 | -5.43807 |
| HZ080,HZ013;s317,CM-1 | 0.000651573 | 0.000220698 | 2.95233 |
| HZ080,s317;HZ013,CM-1 | 0.00795447 | 0.00027723 | 28.6927 |
| HZ017,DBZ-1;HZ064,CM-1 | 0.00499612 | 0.000270917 | 18.4416 |
| HZ017,HZ064;DBZ-1,CM-1 | 0.000747142 | 0.000225409 | 3.3146 |
| HZ004,HZ078;s301-1,CM-1 | 3.78E-05 | 0.000229485 | 0.164709 |
| HZ004,s301-1;HZ078,CM-1 | 0.00566368 | 0.00029038 | 19.5044 |
| HZ037,HZ057;ZBD,CM-1 | 0.00283269 | 0.000262031 | 10.8105 |
| HZ037,ZBD;HZ057,CM-1 | -0.00218451 | 0.000209826 | -10.4111 |
| HZ123,HZ013;HZ036,CM-1 | -0.0105174 | 0.000301916 | -34.8357 |
| HZ123,HZ036;HZ013,CM-1 | -0.0115694 | 0.0002999 | -38.5776 |
| HZ020,HZ100;HZ122,CM-1 | -0.00999276 | 0.000306703 | -32.5813 |
| HZ020,HZ122;HZ100,CM-1 | -0.0106779 | 0.000298507 | -35.7711 |
| LW-5,HZ018;NBE-1,CM-1 | -0.00503597 | 0.000281639 | -17.8809 |
| LW-5,NBE-1;HZ018,CM-1 | -0.00578842 | 0.000269781 | -21.456 |
| HZ059,HZ058;s6-8,CM-1 | -0.00110014 | 0.000233536 | -4.7108 |
| HZ059,s6-8;HZ058,CM-1 | 0.00620606 | 0.000307952 | 20.1527 |
| NBE-4,LC-1;NBE-5,CM-1 | 0.00400025 | 0.000277475 | 14.4166 |
| NBE-4,NBE-5;LC-1,CM-1 | -0.0017465 | 0.000233311 | -7.48571 |
| HZ016,HZ119;QXDM-2,CM-1 | 0.0135723 | 0.00030732 | 44.1634 |
| HZ016,QXDM-2;HZ119,CM-1 | -0.00024516 | 0.000193012 | -1.27015 |
| LW-5,HZ057;NBE-5,CM-1 | -0.00247889 | 0.000286275 | -8.65911 |
| LW-5,NBE-5;HZ057,CM-1 | -0.00466372 | 0.000258398 | -18.0485 |
| YNLDP1,HSKC;HZ109,CM-1 | 0.00196414 | 0.000217396 | 9.03482 |
| YNLDP1,HZ109;HSKC,CM-1 | 0.0055329 | 0.000255238 | 21.6774 |
| HZ050,HZ016;HZ074,CM-1 | -9.11E-05 | 0.000212148 | -0.429236 |
| HZ050,HZ074;HZ016,CM-1 | 0.00946383 | 0.000307362 | 30.7905 |
| HSKC,HZ008;HZ060,CM-1 | -0.0049766 | 0.000279526 | -17.8037 |
| HSKC,HZ060;HZ008,CM-1 | -0.00590316 | 0.000275994 | -21.3887 |
| JX-4,HZ080;HZ088,CM-1 | 0.000166902 | 0.000251866 | 0.662664 |
| JX-4,HZ088;HZ080,CM-1 | -0.00269991 | 0.000236307 | -11.4254 |
| HZ008,HZ058;LW-4,CM-1 | -0.0005107 | 0.00022755 | -2.24435 |
| HZ008,LW-4;HZ058,CM-1 | 0.010462 | 0.000318524 | 32.8451 |
| HZ064,HZ045;ST312,CM-1 | -0.00024291 | 0.000236192 | -1.02845 |
| HZ064,ST312;HZ045,CM-1 | 0.00170406 | 0.00024769 | 6.8798 |
| HZ114,HZ064;QXDM-2,CM-1 | -0.0201451 | 0.000357499 | -56.35 |
| HZ114,QXDM-2;HZ064,CM-1 | -0.0201049 | 0.000359727 | -55.8893 |
| HZ014,HZ043;HZ059,CM-1 | 0.000937277 | 0.000258178 | 3.63035 |
| HZ014,HZ059;HZ043,CM-1 | 0.00281035 | 0.000273937 | 10.2591 |
| HZ001,JHZS;XWY-1,CM-1 | -0.0111347 | 0.000273222 | -40.7532 |
| HZ001,XWY-1;JHZS,CM-1 | -0.00928078 | 0.000296698 | -31.2802 |
| HZ021,NBE-2;XWY-1,CM-1 | 0.00233958 | 0.000290879 | 8.04312 |
| HZ021,XWY-1;NBE-2,CM-1 | -0.00039687 | 0.000270175 | -1.46895 |

| | | | |
|---|---|---|---|
| GLJYD,HZ016;HZ084,CM-1 | -0.00067642 | 0.000232756 | -2.90615 |
| GLJYD,HZ084;HZ016,CM-1 | 0.00665725 | 0.000287264 | 23.1747 |
| HZ118,HZ014;WL,CM-1 | -0.0144981 | 0.000333541 | -43.4673 |
| HZ118,WL;HZ014,CM-1 | -0.0144448 | 0.000335949 | -42.997 |
| HZ021,HZ028;HZ094,CM-1 | 0.000910754 | 0.00017631 | 5.16564 |
| HZ021,HZ094;HZ028,CM-1 | 0.0221827 | 0.000356176 | 62.2802 |
| HZ017,HZ045;HZ081,CM-1 | 0.00628026 | 0.000286874 | 21.892 |
| HZ017,HZ081;HZ045,CM-1 | -0.00148203 | 0.000222962 | -6.647 |
| WL,HZ027;XWY-3,CM-1 | 0.0185799 | 0.000323901 | 57.3628 |
| WL,XWY-3;HZ027,CM-1 | -1.37E-05 | 0.000172625 | -0.0790762 |
| HZ054,HZ010;s301-1,CM-1 | -0.00013106 | 0.000203107 | -0.645265 |
| HZ054,s301-1;HZ010,CM-1 | 0.0103167 | 0.000294561 | 35.024 |
| HZ078,GLJYD;s1-16,CM-1 | 0.000719718 | 0.00021917 | 3.28384 |
| HZ078,s1-16;GLJYD,CM-1 | 0.0059178 | 0.000266063 | 22.2421 |
| LW-3,HZ041;XWY-3,CM-1 | -0.0124084 | 0.000326672 | -37.9841 |
| LW-3,XWY-3;HZ041,CM-1 | -0.012558 | 0.00032536 | -38.5972 |
| HZ077,HZ069;XH-1,CM-1 | 0.00109906 | 0.000214215 | 5.13064 |
| HZ077,XH-1;HZ069,CM-1 | 0.00740032 | 0.000260108 | 28.451 |
| ZBD,LW-2;ST312,CM-1 | 0.00196083 | 0.000259766 | 7.54843 |
| ZBD,ST312;LW-2,CM-1 | -0.00187758 | 0.000236731 | -7.93131 |
| HZ088,HZ020;JHZS,CM-1 | -0.00112933 | 0.000272376 | -4.1462 |
| HZ088,JHZS;HZ020,CM-1 | -0.00044151 | 0.000270693 | -1.63104 |
| LBDCS,LW-5;NBE-5,CM-1 | -0.00232018 | 0.000276365 | -8.39534 |
| LBDCS,NBE-5;LW-5,CM-1 | -0.00516371 | 0.000246557 | -20.9433 |
| HZ100,GE-6;HZ060,CM-1 | -0.0122568 | 0.000327497 | -37.4256 |
| HZ100,HZ060;GE-6,CM-1 | -0.0113983 | 0.000336227 | -33.9007 |
| HZ037,HZ013;HZ077,CM-1 | 0.00182983 | 0.000254618 | 7.18657 |
| HZ037,HZ077;HZ013,CM-1 | -0.00199094 | 0.000233297 | -8.53392 |
| HZ114,HSKC;ST312,CM-1 | -0.0007435 | 0.000257553 | -2.88679 |
| HZ114,ST312;HSKC,CM-1 | -0.00493458 | 0.000232973 | -21.1809 |
| NC,GE-8;LW-2,CM-1 | -0.00331657 | 0.00021814 | -15.2039 |
| NC,LW-2;GE-8,CM-1 | 0.00507702 | 0.000282425 | 17.9765 |
| HZ041,HZ028;HZ123,CM-1 | 0.00143306 | 0.000205295 | 6.98048 |
| HZ041,HZ123;HZ028,CM-1 | 0.01351 | 0.00030873 | 43.7598 |
| HZ117,HZ076;s301-1,CM-1 | 0.00313859 | 0.000236136 | 13.2915 |
| HZ117,s301-1;HZ076,CM-1 | 0.000362045 | 0.000211038 | 1.71554 |
| QXDM-2,HZ009;NBE-3,CM-1 | 0.00150137 | 0.000222367 | 6.75176 |
| QXDM-2,NBE-3;HZ009,CM-1 | 0.00678603 | 0.000275274 | 24.6519 |
| HZ071,HZ039;XH-1,CM-1 | 0.00770222 | 0.000259246 | 29.7101 |
| HZ071,XH-1;HZ039,CM-1 | 0.0017273 | 0.000208696 | 8.27664 |
| HZ004,GLJYD;HZ016,CM-1 | 0.00213102 | 0.000270568 | 7.87609 |
| HZ004,HZ016;GLJYD,CM-1 | -0.00034557 | 0.000261004 | -1.32398 |
| HZ084,HZ043;HZ058,CM-1 | -0.00752095 | 0.000318691 | -23.5995 |
| HZ084,HZ058;HZ043,CM-1 | -0.0080756 | 0.000310609 | -25.9993 |
| HZ071,LW-3;WLH-1,CM-1 | -0.0002488 | 0.000239492 | -1.03887 |
| HZ071,WLH-1;LW-3,CM-1 | -0.00151559 | 0.000238714 | -6.34896 |
| LW-4,GE-8;HZ031,CM-1 | -0.0120888 | 0.000309258 | -39.0897 |
| LW-4,HZ031;GE-8,CM-1 | -0.0118036 | 0.000313899 | -37.6031 |
| s317,HZ045;NBE-1,CM-1 | 0.00302468 | 0.000264582 | 11.4319 |
| s317,NBE-1;HZ045,CM-1 | -0.00090777 | 0.000216168 | -4.19937 |

| | | | |
|---|---|---|---|
| HZ094,HZ018;XSL,CM-1 | -0.0231179 | 0.000365297 | -63.2853 |
| HZ094,XSL;HZ018,CM-1 | -0.0231456 | 0.000368051 | -62.887 |
| HZ077,HZ027;LC-1,CM-1 | 0.011138 | 0.000274634 | 40.5558 |
| HZ077,LC-1;HZ027,CM-1 | -0.00056851 | 0.000191829 | -2.96361 |
| NBE-3,HSKC;HZ039,CM-1 | 0.000190787 | 0.000229643 | 0.830799 |
| NBE-3,HZ039;HSKC,CM-1 | 0.00371595 | 0.000261632 | 14.2029 |
| QXDM-1,HZ048;HZ050,CM-1 | 0.00439511 | 0.000270357 | 16.2567 |
| QXDM-1,HZ050;HZ048,CM-1 | 0.000106833 | 0.000238937 | 0.447116 |
| LBDCS,HZ054;XSL,CM-1 | -0.0131422 | 0.000293862 | -44.7225 |
| LBDCS,XSL;HZ054,CM-1 | -0.0122982 | 0.000302035 | -40.718 |
| HZ088,HZ070;XSL,CM-1 | -0.0023869 | 0.000259284 | -9.20574 |
| HZ088,XSL;HZ070,CM-1 | -0.00344047 | 0.000261578 | -13.1528 |
| HZ036,HZ080;LC-1,CM-1 | 0.00311752 | 0.000256374 | 12.1601 |
| HZ036,LC-1;HZ080,CM-1 | -0.00138364 | 0.000214727 | -6.44373 |
| HZ095,HZ057;XWY-4,CM-1 | -0.0182285 | 0.000347279 | -52.4896 |
| HZ095,XWY-4;HZ057,CM-1 | -0.0154446 | 0.000365531 | -42.2525 |
| XWY-5,HZ015;HZ068,CM-1 | 0.00248092 | 0.000255304 | 9.71753 |
| XWY-5,HZ068;HZ015,CM-1 | -0.00116349 | 0.000227519 | -5.11383 |
| YNLDP1,HZ040;HZ065,CM-1 | -0.00251452 | 0.000228108 | -11.0234 |
| YNLDP1,HZ065;HZ040,CM-1 | 0.0010358 | 0.000271509 | 3.81499 |
| WLH-1,HZ112;LW-1,CM-1 | 0.00134605 | 0.000240355 | 5.60026 |
| WLH-1,LW-1;HZ112,CM-1 | 0.00141418 | 0.000235514 | 6.00464 |
| HZ034,NBE-1;YNZ-1,CM-1 | -0.00106777 | 0.000247176 | -4.31989 |
| HZ034,YNZ-1;NBE-1,CM-1 | 0.00196808 | 0.000269936 | 7.29093 |
| SLLK,HZ040;LW-4,CM-1 | 0.00140492 | 0.000215471 | 6.52024 |
| SLLK,LW-4;HZ040,CM-1 | 0.00566581 | 0.00026008 | 21.7849 |
| HZ119,HZ013;HZ125,CM-1 | 0.00239335 | 0.000207206 | 11.5506 |
| HZ119,HZ125;HZ013,CM-1 | 0.00799042 | 0.000240412 | 33.2363 |
| HZ021,GE-6;HZ015,CM-1 | -0.00201237 | 0.000249938 | -8.05149 |
| HZ021,HZ015;GE-6,CM-1 | 0.00088991 | 0.000275179 | 3.23394 |
| HZ081,HZ043;LBDCS,CM-1 | 0.00101778 | 0.000224267 | 4.53823 |
| HZ081,LBDCS;HZ043,CM-1 | 0.0117175 | 0.00030338 | 38.623 |
| HZ066,GE-8;LW-5,CM-1 | 0.000254213 | 0.000206984 | 1.22818 |
| HZ066,LW-5;GE-8,CM-1 | 0.0121791 | 0.000326601 | 37.2903 |
| XWY-3,HZ014;HZ117,CM-1 | -0.00083816 | 0.000227282 | -3.68773 |
| XWY-3,HZ117;HZ014,CM-1 | 0.00878694 | 0.000318266 | 27.6088 |
| HZ080,HZ013;YNLDP1,CM-1 | -0.00135591 | 0.000245686 | -5.51886 |
| HZ080,YNLDP1;HZ013,CM-1 | 0.000302387 | 0.000259088 | 1.16712 |
| QXDM-1,GE-4;LW-2,CM-1 | -0.00016246 | 0.000214026 | -0.75908 |
| QXDM-1,LW-2;GE-4,CM-1 | 0.0114164 | 0.000309848 | 36.8451 |
| HZ082,NBE-1;YH-2,CM-1 | -0.00798237 | 0.000263871 | -30.2511 |
| HZ082,YH-2;NBE-1,CM-1 | -0.00309029 | 0.000309831 | -9.9741 |
| NBE-2,HZ074;LW-4,CM-1 | 0.000470501 | 0.000252914 | 1.86032 |
| NBE-2,LW-4;HZ074,CM-1 | 0.0011034 | 0.00024397 | 4.52267 |
| MZYCS,HZ069;HZ080,CM-1 | 0.000622161 | 0.000216764 | 2.87022 |
| MZYCS,HZ080;HZ069,CM-1 | 0.00622899 | 0.000265507 | 23.4607 |
| HZ063,HZ084;HZ119,CM-1 | 0.00325529 | 0.000217799 | 14.9463 |
| HZ063,HZ119;HZ084,CM-1 | 0.00618881 | 0.000244442 | 25.3181 |
| HZ002,HZ027;JX-4,CM-1 | 0.00734891 | 0.000281839 | 26.0748 |
| HZ002,JX-4;HZ027,CM-1 | -0.00086204 | 0.000205367 | -4.19758 |

| | | | |
|---|---|---|---|
| HZ125,HZ041;HZ072,CM-1 | -0.00054144 | 0.000266307 | -2.03314 |
| HZ125,HZ072;HZ041,CM-1 | -0.00498099 | 0.000244974 | -20.3327 |
| HZ045,HZ058;ZW,CM-1 | -0.00731443 | 0.000296903 | -24.6358 |
| HZ045,ZW;HZ058,CM-1 | -0.00659807 | 0.000293382 | -22.4897 |
| HZ124,HZ085;JHZS,CM-1 | -0.0116833 | 0.00030213 | -38.6697 |
| HZ124,JHZS;HZ085,CM-1 | -0.011385 | 0.000305761 | -37.2349 |
| DHL,HZ021;JGL,CM-1 | 0.00421917 | 0.000224693 | 18.7775 |
| DHL,JGL;HZ021,CM-1 | 0.000299537 | 0.000215182 | 1.39202 |
| ASM,HZ031;SB,CM-1 | -0.0111054 | 0.000283994 | -39.1044 |
| ASM,SB;HZ031,CM-1 | -0.0110239 | 0.000287426 | -38.3539 |
| JX-4,HZ057;XWY-1,CM-1 | 0.00422324 | 0.000266876 | 15.8247 |
| JX-4,XWY-1;HZ057,CM-1 | 0.00188382 | 0.000242029 | 7.78344 |
| XH-1,GZ;HZ110,CM-1 | 0.000942769 | 0.000196248 | 4.80396 |
| XH-1,HZ110;GZ,CM-1 | 0.00892915 | 0.000246595 | 36.2099 |
| HZ063,HZ064;NBE-2,CM-1 | -0.00296125 | 0.000245408 | -12.0666 |
| HZ063,NBE-2;HZ064,CM-1 | 0.00121244 | 0.000273892 | 4.42671 |
| HZ119,HZ048;LW-4,CM-1 | 0.00263696 | 0.000231748 | 11.3786 |
| HZ119,LW-4;HZ048,CM-1 | 0.00334283 | 0.000231391 | 14.4467 |
| DBZ-1,HZ050;s317,CM-1 | -0.00170143 | 0.000207697 | -8.19188 |
| DBZ-1,s317;HZ050,CM-1 | 0.00690631 | 0.000283773 | 24.3374 |
| HZ079,HZ018;HZ076,CM-1 | -0.00169994 | 0.000214938 | -7.90898 |
| HZ079,HZ076;HZ018,CM-1 | 0.00518879 | 0.000256251 | 20.2488 |
| HZ036,HZ124;NBE-5,CM-1 | 0.00957481 | 0.000293494 | 32.6235 |
| HZ036,NBE-5;HZ124,CM-1 | -0.00039953 | 0.000213654 | -1.86998 |
| HZ034,HZ014;LW-3,CM-1 | 1.01E-05 | 0.000202451 | 0.0499649 |
| HZ034,LW-3;HZ014,CM-1 | 0.0120937 | 0.000309355 | 39.0934 |
| GE-1,HZ039;YH-2,CM-1 | 0.00213218 | 0.00024237 | 8.79721 |
| GE-1,YH-2;HZ039,CM-1 | 0.00347194 | 0.00025924 | 13.3927 |
| XYDCS,HZ070;HZ081,CM-1 | -0.00837455 | 0.000289314 | -28.9462 |
| XYDCS,HZ081;HZ070,CM-1 | -0.0107106 | 0.000275438 | -38.8856 |
| HZ056,HZ040;HZ043,CM-1 | -0.00696189 | 0.000280263 | -24.8406 |
| HZ056,HZ043;HZ040,CM-1 | -0.0076003 | 0.000271046 | -28.0406 |
| JX-4,HZ013;XSL,CM-1 | 0.000610236 | 0.00025873 | 2.35858 |
| JX-4,XSL;HZ013,CM-1 | -0.00123642 | 0.000242733 | -5.09374 |
| GE-1,HSKC;SLLK,CM-1 | 0.00257572 | 0.000241742 | 10.6549 |
| GE-1,SLLK;HSKC,CM-1 | 0.00426482 | 0.000256545 | 16.6241 |
| HZ018,XWY-3;YH-2,CM-1 | -0.00046523 | 0.000214835 | -2.16553 |
| HZ018,YH-2;XWY-3,CM-1 | 0.0137827 | 0.000337462 | 40.8422 |
| SLLK,JX-4;NBE-4,CM-1 | -0.00193625 | 0.000255708 | -7.5721 |
| SLLK,NBE-4;JX-4,CM-1 | -0.00102196 | 0.000262099 | -3.89915 |
| HZ054,HZ090;YNLDP1,CM-1 | 0.00291996 | 0.000241957 | 12.0681 |
| HZ054,YNLDP1;HZ090,CM-1 | 0.00198303 | 0.000242462 | 8.17872 |
| HZ015,FL;LW-5,CM-1 | 0.000427675 | 0.000222265 | 1.92417 |
| HZ015,LW-5;FL,CM-1 | 0.00827203 | 0.000303395 | 27.2649 |
| NBE-2,HZ045;HZ095,CM-1 | 0.000144805 | 0.000199198 | 0.72694 |
| NBE-2,HZ095;HZ045,CM-1 | 0.0142133 | 0.000314022 | 45.2622 |
| HZ094,HZ088;NBE-1,CM-1 | -0.0139609 | 0.000339941 | -41.0687 |
| HZ094,NBE-1;HZ088,CM-1 | -0.014616 | 0.000329248 | -44.3921 |
| WL,YH-2;s303-231,CM-1 | -0.00759577 | 0.000322483 | -23.554 |
| WL,s303-231;YH-2,CM-1 | -0.0117388 | 0.000278531 | -42.1454 |

| | | | |
|---|---|---|---|
| HZ085,HZ095;XH-1,CM-1 | 0.0162107 | 0.000327028 | 49.5697 |
| HZ085,XH-1;HZ095,CM-1 | -0.00048918 | 0.000186637 | -2.62104 |
| HZ050,GE-6;HSKC,CM-1 | -0.00092323 | 0.000247492 | -3.73035 |
| HZ050,HSKC;GE-6,CM-1 | 0.00213784 | 0.000276107 | 7.74277 |
| HZ095,GE-1;HSKC,CM-1 | -0.0151874 | 0.000345326 | -43.9799 |
| HZ095,HSKC;GE-1,CM-1 | -0.0184103 | 0.00032299 | -56.9997 |
| HZ122,HZ060;HZ074,CM-1 | -0.00151553 | 0.00027912 | -5.42968 |
| HZ122,HZ074;HZ060,CM-1 | -0.00194814 | 0.000279334 | -6.97422 |
| HZ073,HZ021;ZW,CM-1 | 0.00221939 | 0.000246874 | 8.98995 |
| HZ073,ZW;HZ021,CM-1 | -0.00063978 | 0.000230276 | -2.77832 |
| LBDCS,GE-6;SB,CM-1 | -0.013561 | 0.000264764 | -51.2191 |
| LBDCS,SB;GE-6,CM-1 | -0.0124629 | 0.000276474 | -45.0779 |
| QXDM-1,HZ104;ST312,CM-1 | -0.00315029 | 0.000290751 | -10.835 |
| QXDM-1,ST312;HZ104,CM-1 | -0.00675374 | 0.000262439 | -25.7345 |
| JHZS,XWY-1;XWY-4,CM-1 | 0.0110156 | 0.000279478 | 39.4149 |
| JHZS,XWY-4;XWY-1,CM-1 | 0.000686287 | 0.000204744 | 3.35194 |
| HZ064,XY-10;YNLDP2,CM-1 | -0.00032244 | 0.000212128 | -1.52003 |
| HZ064,YNLDP2;XY-10,CM-1 | 0.00891339 | 0.000296388 | 30.0734 |
| DHL,LW-3;XSX,CM-1 | 0.0190341 | 0.00034171 | 55.7026 |
| DHL,XSX;LW-3,CM-1 | -0.00024321 | 0.000175721 | -1.38407 |
| MZYCS,DHL;YNZ-1,CM-1 | 8.76E-05 | 0.000177008 | 0.495103 |
| MZYCS,YNZ-1;DHL,CM-1 | 0.0169606 | 0.000320489 | 52.9211 |
| HZ085,HZ034;HZ125,CM-1 | 0.000134064 | 0.000186898 | 0.717308 |
| HZ085,HZ125;HZ034,CM-1 | 0.0173145 | 0.000331594 | 52.216 |
| LBDCS,YH-1-1;s303-231,CM-1 | -0.0139124 | 0.000307608 | -45.2276 |
| LBDCS,s303-231;YH-1-1,CM-1 | -0.0154603 | 0.000292345 | -52.8838 |
| HZ031,GE-6;HZ061,CM-1 | -0.00183665 | 0.000223795 | -8.20681 |
| HZ031,HZ061;GE-6,CM-1 | 0.0037399 | 0.000285464 | 13.1011 |
| HZ094,ASM;NBE-2,CM-1 | -0.0142154 | 0.000320047 | -44.4167 |
| HZ094,NBE-2;ASM,CM-1 | -0.0140679 | 0.000321338 | -43.779 |
| DHL,HZ094;LW-4,CM-1 | 0.00890892 | 0.000318522 | 27.9696 |
| DHL,LW-4;HZ094,CM-1 | -0.00065926 | 0.000225474 | -2.92389 |
| LW-4,HZ094;s301-1,CM-1 | 0.0117675 | 0.00031134 | 37.7962 |
| LW-4,s301-1;HZ094,CM-1 | -8.81E-05 | 0.000193592 | -0.455025 |
| ASM,GE-4;HZ045,CM-1 | -0.00224957 | 0.000258547 | -8.70083 |
| ASM,HZ045;GE-4,CM-1 | -0.00271812 | 0.000241331 | -11.263 |
| HZ013,BCS;HZ074,CM-1 | -0.00104182 | 0.000221056 | -4.71292 |
| HZ013,HZ074;BCS,CM-1 | 0.00870501 | 0.000300371 | 28.9809 |
| HZ085,GZ;YH-2,CM-1 | -0.00104295 | 0.000228999 | -4.55439 |
| HZ085,YH-2;GZ,CM-1 | 0.00795177 | 0.000312798 | 25.4214 |
| HZ063,XWY-5;ZW,CM-1 | -0.00400878 | 0.000274992 | -14.5778 |
| HZ063,ZW;XWY-5,CM-1 | -0.00642293 | 0.000239909 | -26.7724 |
| HZ045,QXDM-1;XWY-3,CM-1 | -0.00722203 | 0.000288296 | -25.0507 |
| HZ045,XWY-3;QXDM-1,CM-1 | -0.00792365 | 0.000285502 | -27.7533 |
| HZ048,HZ119;HZ124,CM-1 | -0.0012115 | 0.000221931 | -5.45891 |
| HZ048,HZ124;HZ119,CM-1 | 0.00313134 | 0.000244196 | 12.8231 |
| ASM,HZ073;HZ118,CM-1 | 0.00442238 | 0.000252047 | 17.5458 |
| ASM,HZ118;HZ073,CM-1 | 0.00241461 | 0.00023264 | 10.3792 |
| GZ,HZ043;NBE-5,CM-1 | -0.00019178 | 0.000255101 | -0.751786 |
| GZ,NBE-5;HZ043,CM-1 | 0.00283207 | 0.000289901 | 9.7691 |

| | | | |
|---|---|---|---|
| YNZ-1,HZ069;ST330,CM-1 | -0.0022045 | 0.000248534 | -8.87002 |
| YNZ-1,ST330;HZ069,CM-1 | -0.00226462 | 0.000256725 | -8.82117 |
| LBDCS,HZ080;HZ125,CM-1 | 0.000756258 | 0.000216026 | 3.50077 |
| LBDCS,HZ125;HZ080,CM-1 | 0.00360361 | 0.00023877 | 15.0924 |
| SB,HZ048;s303-231,CM-1 | 0.00110613 | 0.000220539 | 5.01559 |
| SB,s303-231;HZ048,CM-1 | 0.00752121 | 0.000261924 | 28.7152 |
| XWY-2,HZ057;JGL,CM-1 | 0.00512467 | 0.000273747 | 18.7205 |
| XWY-2,JGL;HZ057,CM-1 | -0.00198539 | 0.000214907 | -9.23838 |
| LW-5,HZ013;HZ039,CM-1 | -0.00167021 | 0.000259561 | -6.43473 |
| LW-5,HZ039;HZ013,CM-1 | -0.00104261 | 0.000273639 | -3.81016 |
| NBE-4,HZ058;HZ063,CM-1 | -0.00765468 | 0.000281994 | -27.1448 |
| NBE-4,HZ063;HZ058,CM-1 | -0.00609486 | 0.000291951 | -20.8763 |
| LW-5,HZ117;JHZS,CM-1 | -0.00303739 | 0.000251418 | -12.081 |
| LW-5,JHZS;HZ117,CM-1 | -0.00123344 | 0.000262458 | -4.69959 |
| HZ119,FL;LC-2,CM-1 | -0.00667169 | 0.000254627 | -26.2018 |
| HZ119,LC-2;FL,CM-1 | -0.00604591 | 0.000262074 | -23.0695 |
| YH-2,BCS;XWY-4,CM-1 | -0.0196594 | 0.000349416 | -56.2636 |
| YH-2,XWY-4;BCS,CM-1 | -0.0196839 | 0.000348637 | -56.4597 |
| HZ003,GE-6;HZ077,CM-1 | 0.00310119 | 0.000242383 | 12.7946 |
| HZ003,HZ077;GE-6,CM-1 | 9.34E-05 | 0.000232923 | 0.401169 |
| HZ104,ASM;ST330,CM-1 | -0.00304629 | 0.000237185 | -12.8435 |
| HZ104,ST330;ASM,CM-1 | -0.0022461 | 0.000248699 | -9.03141 |
| ZBD,HZ085;HZ117,CM-1 | -0.00109336 | 0.000205842 | -5.31167 |
| ZBD,HZ117;HZ085,CM-1 | 0.00737694 | 0.000278425 | 26.4952 |
| ZW,HZ073;HZ118,CM-1 | -3.69E-05 | 0.000198033 | -0.186571 |
| ZW,HZ118;HZ073,CM-1 | 0.0154511 | 0.000334931 | 46.1321 |
| HSKC,HZ112;XWY-2,CM-1 | -0.00236104 | 0.000247681 | -9.53258 |
| HSKC,XWY-2;HZ112,CM-1 | 0.00048804 | 0.000290342 | 1.68091 |
| s303-231,HZ068;XSX,CM-1 | -0.00956544 | 0.000320524 | -29.8432 |
| s303-231,XSX;HZ068,CM-1 | -0.0101605 | 0.000314667 | -32.2897 |
| HZ125,HZ100;HZ112,CM-1 | -0.00314886 | 0.000230778 | -13.6445 |
| HZ125,HZ112;HZ100,CM-1 | 0.000997417 | 0.000258645 | 3.85632 |
| HZ056,HZ094;HZ117,CM-1 | 0.0137589 | 0.000325366 | 42.2873 |
| HZ056,HZ117;HZ094,CM-1 | -0.00049186 | 0.00019213 | -2.56001 |
| JHZS,HZ031;HZ078,CM-1 | 0.000711209 | 0.000246002 | 2.89107 |
| JHZS,HZ078;HZ031,CM-1 | 0.00366325 | 0.000268534 | 13.6417 |
| XY-10,HZ021;HZ110,CM-1 | 5.23E-05 | 0.000192127 | 0.2723 |
| XY-10,HZ110;HZ021,CM-1 | 0.0149773 | 0.00032279 | 46.3994 |
| s301-1,HZ010;LW-2,CM-1 | 0.00187713 | 0.000221076 | 8.49089 |
| s301-1,LW-2;HZ010,CM-1 | 0.00526307 | 0.000252576 | 20.8376 |
| HZ110,HZ125;JX-4,CM-1 | 0.00046182 | 0.000198933 | 2.32149 |
| HZ110,JX-4;HZ125,CM-1 | 0.00584139 | 0.000255269 | 22.8833 |
| LC-2,HZ084;HZ112,CM-1 | 0.00546622 | 0.000285333 | 19.1574 |
| LC-2,HZ112;HZ084,CM-1 | -0.00121324 | 0.000231319 | -5.24486 |
| HZ104,HZ083;YNLDP1,CM-1 | -0.00727596 | 0.000314105 | -23.1641 |
| HZ104,YNLDP1;HZ083,CM-1 | -0.0100204 | 0.000296439 | -33.8025 |
| ST330,HZ076;HZ092,CM-1 | 0.000502619 | 0.000232952 | 2.1576 |
| ST330,HZ092;HZ076,CM-1 | 0.00102528 | 0.000235712 | 4.34972 |
| HZ119,HZ028;HZ086,CM-1 | -0.00738056 | 0.000266364 | -27.7086 |
| HZ119,HZ086;HZ028,CM-1 | -0.0070798 | 0.000257454 | -27.4993 |

| | | | |
|---|---|---|---|
| HZ090,HZ048;HZ072,CM-1 | 0.00416776 | 0.000242736 | 17.1699 |
| HZ090,HZ072;HZ048,CM-1 | 0.00476314 | 0.000231848 | 20.5443 |
| HZ056,HZ066;NBE-2,CM-1 | -0.0021233 | 0.000230263 | -9.22121 |
| HZ056,NBE-2;HZ066,CM-1 | 0.00212333 | 0.000271272 | 7.8273 |
| LW-3,FL;HZ122,CM-1 | 0.0018958 | 0.000258204 | 7.34224 |
| LW-3,HZ122;FL,CM-1 | -0.00045959 | 0.000237853 | -1.93223 |
| XYDCS,HZ011;XSX,CM-1 | -0.014129 | 0.000287342 | -49.1713 |
| XYDCS,XSX;HZ011,CM-1 | -0.0146326 | 0.000295708 | -49.4832 |
| HZ066,HZ082;HZ090,CM-1 | 0.000531869 | 0.000228928 | 2.3233 |
| HZ066,HZ090;HZ082,CM-1 | 0.0048826 | 0.000268778 | 18.166 |
| HZ068,HZ036;LW-1,CM-1 | -0.00043368 | 0.000208129 | -2.08368 |
| HZ068,LW-1;HZ036,CM-1 | 0.0125978 | 0.000329308 | 38.2555 |
| DHL,JGL;LW-4,CM-1 | 0.000178783 | 0.000174391 | 1.02518 |
| DHL,LW-4;JGL,CM-1 | 0.0165496 | 0.000306937 | 53.9186 |
| BCS,HZ045;s301-1,CM-1 | 0.00105067 | 0.000226098 | 4.64696 |
| BCS,s301-1;HZ045,CM-1 | 0.00275444 | 0.000241406 | 11.41 |
| HZ056,LBDCS;LW-4,CM-1 | 0.00559578 | 0.000240202 | 23.2961 |
| HZ056,LW-4;LBDCS,CM-1 | 0.00575619 | 0.000239205 | 24.0638 |
| HZ078,HZ110;MZYCS,CM-1 | 0.0119577 | 0.000295885 | 40.4134 |
| HZ078,MZYCS;HZ110,CM-1 | 0.0020371 | 0.000214971 | 9.47614 |
| LW-5,HZ082;HZ086,CM-1 | -0.0116682 | 0.000321558 | -36.2864 |
| LW-5,HZ086;HZ082,CM-1 | -0.0114523 | 0.00031183 | -36.726 |
| HZ069,HZ094;HZ114,CM-1 | -0.00071706 | 0.000278713 | -2.57276 |
| HZ069,HZ114;HZ094,CM-1 | -0.00533503 | 0.000240623 | -22.1717 |
| HZ028,HZ054;LW-2,CM-1 | -0.00097369 | 0.000183657 | -5.30164 |
| HZ028,LW-2;HZ054,CM-1 | 0.0153562 | 0.000337139 | 45.5486 |
| HZ066,NC;YNLDP2,CM-1 | 0.00536705 | 0.000214759 | 24.991 |
| HZ066,YNLDP2;NC,CM-1 | 0.00516815 | 0.000238785 | 21.6435 |
| XYDCS,HZ041;HZ114,CM-1 | 0.000360147 | 0.000224476 | 1.60439 |
| XYDCS,HZ114;HZ041,CM-1 | 0.00585688 | 0.00026347 | 22.2298 |
| LW-3,GE-6;JGL,CM-1 | -0.0138106 | 0.000298875 | -46.2086 |
| LW-3,JGL;GE-6,CM-1 | -0.0129779 | 0.000315394 | -41.1483 |
| HZ082,HZ018;HZ094,CM-1 | 0.000284702 | 0.000184002 | 1.54728 |
| HZ082,HZ094;HZ018,CM-1 | 0.0211067 | 0.000376827 | 56.0117 |
| YNLDP2,HZ037;HZ114,CM-1 | 0.00150835 | 0.000197798 | 7.62574 |
| YNLDP2,HZ114;HZ037,CM-1 | 0.00899913 | 0.0002667 | 33.7425 |
| GE-8,NC;XWY-5,CM-1 | 0.0100204 | 0.000254246 | 39.4121 |
| GE-8,XWY-5;NC,CM-1 | 0.00197998 | 0.000199981 | 9.90088 |
| ZBD,GE-1;NC,CM-1 | 0.000179308 | 0.00020428 | 0.877759 |
| ZBD,NC;GE-1,CM-1 | 0.0085773 | 0.000245145 | 34.9886 |
| HZ078,HZ027;HZ094,CM-1 | -0.00696453 | 0.000226423 | -30.759 |
| HZ078,HZ094;HZ027,CM-1 | 0.00181096 | 0.000294945 | 6.14001 |
| HZ081,JHZS;SB,CM-1 | -0.00767483 | 0.000256114 | -29.9665 |
| HZ081,SB;JHZS,CM-1 | -0.00764085 | 0.000253404 | -30.1528 |
| JGL,HZ010;HZ043,CM-1 | -0.00048001 | 0.00023539 | -2.03919 |
| JGL,HZ043;HZ010,CM-1 | -0.00115434 | 0.000242382 | -4.76249 |
| SLLK,HZ110;MZYCS,CM-1 | 0.00846161 | 0.000274491 | 30.8266 |
| SLLK,MZYCS;HZ110,CM-1 | 0.00111146 | 0.000218846 | 5.07874 |
| HZ122,DBZ-1;LW-5,CM-1 | 0.00311141 | 0.000265961 | 11.6988 |
| HZ122,LW-5;DBZ-1,CM-1 | -8.18E-05 | 0.000231874 | -0.352686 |

| | | | |
|---|---|---|---|
| HZ112,HZ048;HZ059,CM-1 | 0.00247989 | 0.000274441 | 9.03617 |
| HZ112,HZ059;HZ048,CM-1 | 0.000287407 | 0.000257019 | 1.11823 |
| SB,HZ027;WLH-1,CM-1 | 0.00441178 | 0.000220931 | 19.969 |
| SB,WLH-1;HZ027,CM-1 | 0.00431752 | 0.000225672 | 19.1319 |
| HZ060,HZ001;HZ037,CM-1 | 0.0119792 | 0.000285826 | 41.9108 |
| HZ060,HZ037;HZ001,CM-1 | 0.000110713 | 0.000202455 | 0.546852 |
| s1-16,MZYCS;XWY-5,CM-1 | -0.0123574 | 0.000294724 | -41.9288 |
| s1-16,XWY-5;MZYCS,CM-1 | -0.0125054 | 0.000292163 | -42.8029 |
| BCS,HZ002;HZ077,CM-1 | 0.00552935 | 0.000262216 | 21.087 |
| BCS,HZ077;HZ002,CM-1 | -0.00065218 | 0.000227008 | -2.87293 |
| YH-1-1,HZ122;s6-8,CM-1 | 0.00576108 | 0.000238426 | 24.163 |
| YH-1-1,s6-8;HZ122,CM-1 | 0.00227699 | 0.000198368 | 11.4786 |
| HZ015,HZ059;XWY-5,CM-1 | -0.00135644 | 0.000242229 | -5.59982 |
| HZ015,XWY-5;HZ059,CM-1 | 0.00340216 | 0.000282039 | 12.0628 |
| HZ037,HZ040;HZ080,CM-1 | -0.00019399 | 0.0002036 | -0.952821 |
| HZ037,HZ080;HZ040,CM-1 | 0.00622242 | 0.000255496 | 24.3543 |
| HZ031,QXDM-1;ZBD,CM-1 | 0.00095519 | 0.000243646 | 3.9204 |
| HZ031,ZBD;QXDM-1,CM-1 | 0.000599443 | 0.000228726 | 2.62079 |
| JX-4,HZ018;HZ124,CM-1 | 0.000184126 | 0.000200222 | 0.919608 |
| JX-4,HZ124;HZ018,CM-1 | 0.0132375 | 0.000309913 | 42.7135 |
| HZ069,HZ081;JX-4,CM-1 | -0.00035492 | 0.000246521 | -1.43973 |
| HZ069,JX-4;HZ081,CM-1 | 0.000933391 | 0.000251136 | 3.71668 |
| HZ036,HZ084;LC-1,CM-1 | 0.00577568 | 0.000269619 | 21.4216 |
| HZ036,LC-1;HZ084,CM-1 | -0.00094617 | 0.000206919 | -4.57266 |
| DHL,HZ088;s317,CM-1 | 0.000192944 | 0.000217735 | 0.886141 |
| DHL,s317;HZ088,CM-1 | 0.0104727 | 0.000319801 | 32.7476 |
| HZ069,HZ077;LW-2,CM-1 | 0.000337651 | 0.0001981 | 1.70445 |
| HZ069,LW-2;HZ077,CM-1 | 0.0150348 | 0.000326699 | 46.0204 |
| HZ122,HZ058;ZW,CM-1 | -0.0119339 | 0.000347061 | -34.3857 |
| HZ122,ZW;HZ058,CM-1 | -0.0117605 | 0.000334592 | -35.1487 |
| XSL,GE-8;HZ027,CM-1 | -0.00140534 | 0.000191851 | -7.32514 |
| XSL,HZ027;GE-8,CM-1 | 0.0122377 | 0.000293244 | 41.7322 |
| HZ071,HZ028;HZ124,CM-1 | 0.000943274 | 0.000189515 | 4.97731 |
| HZ071,HZ124;HZ028,CM-1 | 0.0172269 | 0.000315581 | 54.5879 |
| HZ017,HZ013;XWY-1,CM-1 | 0.00250978 | 0.000237915 | 10.549 |
| HZ017,XWY-1;HZ013,CM-1 | 0.00480553 | 0.000263489 | 18.2381 |
| HZ048,HZ020;XH-1,CM-1 | -0.0011418 | 0.000242637 | -4.7058 |
| HZ048,XH-1;HZ020,CM-1 | 0.00344358 | 0.00025796 | 13.3493 |
| HZ001,NBE-2;WL,CM-1 | -0.00620015 | 0.000298599 | -20.7642 |
| HZ001,WL;NBE-2,CM-1 | -0.00853609 | 0.000283915 | -30.0657 |
| HZ056,HZ009;HZ085,CM-1 | 0.00243527 | 0.000286637 | 8.49602 |
| HZ056,HZ085;HZ009,CM-1 | -0.00196322 | 0.000244328 | -8.03518 |
| YNLDP1,HZ122;XWY-5,CM-1 | 0.00753953 | 0.000273061 | 27.6112 |
| YNLDP1,XWY-5;HZ122,CM-1 | 0.000643259 | 0.000204956 | 3.13852 |
| YNLDP1,XWY-3;YH-1-1,CM-1 | 0.00452425 | 0.000234132 | 19.3236 |
| YNLDP1,YH-1-1;XWY-3,CM-1 | 0.00886325 | 0.000291326 | 30.4238 |
| GLJYD,HZ011;HZ037,CM-1 | -0.00330595 | 0.000259634 | -12.7331 |
| GLJYD,HZ037;HZ011,CM-1 | -0.00242001 | 0.0002536 | -9.54262 |
| HZ076,GE-1;QXDM-1,CM-1 | -0.00792661 | 0.000277015 | -28.6144 |
| HZ076,QXDM-1;GE-1,CM-1 | -0.00682829 | 0.000285277 | -23.9357 |

| | | | |
|---|---|---|---|
| GE-1,HZ013;WL,CM-1 | 0.00228484 | 0.000249966 | 9.14061 |
| GE-1,WL;HZ013,CM-1 | -0.00034574 | 0.000237072 | -1.45837 |
| MZYCS,HZ016;XSX,CM-1 | 0.0100319 | 0.000298693 | 33.5862 |
| MZYCS,XSX;HZ016,CM-1 | -0.00017404 | 0.000193594 | -0.89901 |
| XWY-4,QXDM-2;XWY-5,CM- | 0.00103697 | 0.000268726 | 3.85885 |
| XWY-4,XWY-5;QXDM-2,CM- | 0.000232222 | 0.000254396 | 0.912836 |
| SLLK,JGL;YH-1-1,CM-1 | 0.0027377 | 0.000226711 | 12.0757 |
| SLLK,YH-1-1;JGL,CM-1 | 0.00539864 | 0.000260748 | 20.7045 |
| HZ002,HZ064;XSL,CM-1 | -0.00520735 | 0.000256164 | -20.3282 |
| HZ002,XSL;HZ064,CM-1 | -0.00551906 | 0.00026543 | -20.7929 |
| GE-8,HZ009;XWY-2,CM-1 | 0.00182425 | 0.000277668 | 6.5699 |
| GE-8,XWY-2;HZ009,CM-1 | 0.000106205 | 0.000258468 | 0.4109 |
| HZ057,SLLK;YH-2,CM-1 | -0.00400254 | 0.000238312 | -16.7954 |
| HZ057,YH-2;SLLK,CM-1 | 0.00249167 | 0.00029641 | 8.40616 |
| HZ045,HZ092;SLLK,CM-1 | 0.00595985 | 0.000237552 | 25.0886 |
| HZ045,SLLK;HZ092,CM-1 | 0.000625987 | 0.000206494 | 3.0315 |
| YH-2,HZ028;HZ074,CM-1 | -0.00230568 | 0.000273049 | -8.44419 |
| YH-2,HZ074;HZ028,CM-1 | -0.00289395 | 0.000269846 | -10.7245 |
| XYDCS,HZ050;HZ075,CM-1 | -0.00670323 | 0.000245148 | -27.3436 |
| XYDCS,HZ075;HZ050,CM-1 | -0.00407136 | 0.000269161 | -15.1261 |
| s303-231,HZ054;s1-16,CM-1 | 0.000129651 | 0.000240029 | 0.540148 |
| s303-231,s1-16;HZ054,CM-1 | 0.0023603 | 0.000255596 | 9.2345 |
| FL,ASM;GE-1,CM-1 | 0.0106427 | 0.000302722 | 35.1566 |
| FL,GE-1;ASM,CM-1 | -0.0003709 | 0.000208832 | -1.77608 |
| GE-8,JX-4;MZYCS,CM-1 | 0.00510379 | 0.000286187 | 17.8338 |
| GE-8,MZYCS;JX-4,CM-1 | -0.00137234 | 0.000223308 | -6.1455 |
| XYDCS,HZ068;SLLK,CM-1 | -0.00501272 | 0.000261189 | -19.192 |
| XYDCS,SLLK;HZ068,CM-1 | -0.00421972 | 0.000265158 | -15.914 |
| XWY-4,MZYCS;NBE-3,CM-1 | -0.00050993 | 0.000201763 | -2.52739 |
| XWY-4,NBE-3;MZYCS,CM-1 | 0.011158 | 0.000280116 | 39.8333 |
| ZBD,HZ061;XWY-1,CM-1 | 0.00370951 | 0.00022824 | 16.2527 |
| ZBD,XWY-1;HZ061,CM-1 | 0.00219929 | 0.000239406 | 9.18643 |
| HZ057,HZ086;XWY-1,CM-1 | -0.00410465 | 0.000257667 | -15.9301 |
| HZ057,XWY-1;HZ086,CM-1 | -2.25E-05 | 0.00029265 | -0.0767229 |
| GE-4,HZ057;HZ104,CM-1 | 0.00181976 | 0.000216035 | 8.42346 |
| GE-4,HZ104;HZ057,CM-1 | 0.00934139 | 0.000305187 | 30.6088 |
| HZ065,HZ117;NBE-3,CM-1 | 0.00101039 | 0.000261527 | 3.86344 |
| HZ065,NBE-3;HZ117,CM-1 | -0.00160222 | 0.00023021 | -6.95981 |
| YH-1-1,HZ063;QXDM-2,CM-1 | -0.00881124 | 0.000297169 | -29.6506 |
| YH-1-1,QXDM-2;HZ063,CM-1 | -0.00916526 | 0.000309002 | -29.6608 |
| HZ092,HSKC;HZ065,CM-1 | -0.00325799 | 0.000261816 | -12.4438 |
| HZ092,HZ065;HSKC,CM-1 | -0.00535189 | 0.000253121 | -21.1436 |
| XY-10,NBE-4;s301-1,CM-1 | -0.00065263 | 0.000257008 | -2.53933 |
| XY-10,s301-1;NBE-4,CM-1 | 0.00136932 | 0.000276749 | 4.94786 |
| HZ070,HZ041;HZ084,CM-1 | -0.00045866 | 0.000209477 | -2.18954 |
| HZ070,HZ084;HZ041,CM-1 | 0.0102976 | 0.000292929 | 35.1541 |
| HZ048,HZ036;HZ077,CM-1 | -0.00344619 | 0.000267886 | -12.8644 |
| HZ048,HZ077;HZ036,CM-1 | -0.004437 | 0.000262796 | -16.8838 |
| HZ066,HZ063;YH-1-1,CM-1 | 0.000267761 | 0.000200116 | 1.33803 |
| HZ066,YH-1-1;HZ063,CM-1 | 0.0109403 | 0.000298203 | 36.6874 |

| | | | |
|---|---|---|---|
| HZ010,HZ090;WL,CM-1 | 0.00617497 | 0.000259597 | 23.7867 |
| HZ010,WL;HZ090,CM-1 | 0.002654 | 0.000233757 | 11.3537 |
| HZ036,GE-6;HZ086,CM-1 | -9.59E-05 | 0.000247905 | -0.386881 |
| HZ036,HZ086;GE-6,CM-1 | -0.00094424 | 0.000248964 | -3.79267 |
| HZ010,HZ027;HZ085,CM-1 | 0.00748814 | 0.000272085 | 27.5213 |
| HZ010,HZ085;HZ027,CM-1 | -0.00240884 | 0.000217442 | -11.0781 |
| YH-1-1,HZ014;XH-1,CM-1 | 0.00609556 | 0.000282938 | 21.5438 |
| YH-1-1,XH-1;HZ014,CM-1 | -0.0066192 | 0.000214609 | -30.8431 |
| HZ082,HZ060;QXDM-2,CM-1 | -0.00281135 | 0.000267346 | -10.5158 |
| HZ082,QXDM-2;HZ060,CM-1 | -0.00212088 | 0.000279448 | -7.58952 |
| HZ061,HZ086;HZ110,CM-1 | 0.00165768 | 0.000211852 | 7.8247 |
| HZ061,HZ110;HZ086,CM-1 | 0.00975917 | 0.000268472 | 36.3507 |
| HZ064,ST330;XSX,CM-1 | 0.00887347 | 0.000298346 | 29.7422 |
| HZ064,XSX;ST330,CM-1 | -0.00022004 | 0.00022007 | -0.999862 |
| HZ114,HZ034;YNZ-1,CM-1 | -0.00532773 | 0.00026872 | -19.8263 |
| HZ114,YNZ-1;HZ034,CM-1 | -0.00685874 | 0.000253595 | -27.046 |
| GLJYD,HZ090;s303-231,CM-1 | 0.000194564 | 0.000238544 | 0.815632 |
| GLJYD,s303-231;HZ090,CM-1 | 0.00189801 | 0.000235155 | 8.07131 |
| HZ028,HZ059;ZW,CM-1 | 0.00388664 | 0.00027282 | 14.2462 |
| HZ028,ZW;HZ059,CM-1 | -0.00129636 | 0.000228834 | -5.66505 |
| GE-6,HZ061;ST312,CM-1 | -0.00025134 | 0.000215541 | -1.16611 |
| GE-6,ST312;HZ061,CM-1 | 0.00482825 | 0.000268422 | 17.9875 |
| HZ075,HZ013;YNZ-1,CM-1 | 0.00244125 | 0.000240713 | 10.1417 |
| HZ075,YNZ-1;HZ013,CM-1 | 0.00313663 | 0.000238367 | 13.1589 |
| HZ104,HZ027;HZ034,CM-1 | -0.00416669 | 0.00028332 | -14.7066 |
| HZ104,HZ034;HZ027,CM-1 | -0.00750866 | 0.000268422 | -27.9733 |
| NC,HZ077;HZ095,CM-1 | 0.00647431 | 0.000231231 | 27.9993 |
| NC,HZ095;HZ077,CM-1 | 0.0139072 | 0.000285497 | 48.7121 |
| HZ050,HZ061;XWY-3,CM-1 | 0.0049625 | 0.000263509 | 18.8324 |
| HZ050,XWY-3;HZ061,CM-1 | 0.000358784 | 0.000234671 | 1.52888 |
| XSX,FL;HZ086,CM-1 | 0.000870474 | 0.00020518 | 4.24248 |
| XSX,HZ086;FL,CM-1 | 0.00723921 | 0.000263824 | 27.4396 |
| HZ068,GE-4;YNZ-1,CM-1 | 0.000494238 | 0.000205947 | 2.39984 |
| HZ068,YNZ-1;GE-4,CM-1 | 0.00878281 | 0.000290344 | 30.2497 |
| SB,HZ008;HZ073,CM-1 | 0.00089755 | 0.000226802 | 3.95742 |
| SB,HZ073;HZ008,CM-1 | -0.00026262 | 0.000214715 | -1.22312 |
| HZ002,HZ016;HZ018,CM-1 | -0.00388212 | 0.000265254 | -14.6355 |
| HZ002,HZ018;HZ016,CM-1 | -0.00464228 | 0.000260955 | -17.7896 |
| HZ109,GE-1;GZ,CM-1 | -0.00932308 | 0.00027865 | -33.458 |
| HZ109,GZ;GE-1,CM-1 | -0.00933701 | 0.000276479 | -33.7711 |
| HZ068,HZ009;HZ100,CM-1 | -0.0004645 | 0.000211381 | -2.19744 |
| HZ068,HZ100;HZ009,CM-1 | 0.00954416 | 0.000315062 | 30.293 |
| s317,DBZ-1;XYDCS,CM-1 | -0.00316993 | 0.000227116 | -13.9573 |
| s317,XYDCS;DBZ-1,CM-1 | 3.86E-06 | 0.000269079 | 0.0143284 |
| QXDM-2,HZ003;HZ114,CM-1 | 0.00022616 | 0.000186341 | 1.21369 |
| QXDM-2,HZ114;HZ003,CM-1 | 0.0187192 | 0.00033715 | 55.5218 |
| HZ045,HZ082;QXDM-1,CM-1 | -0.00487128 | 0.000253343 | -19.228 |
| HZ045,QXDM-1;HZ082,CM-1 | -0.00104143 | 0.000279913 | -3.72056 |
| HZ064,HZ011;LW-4,CM-1 | 0.000358636 | 0.000202356 | 1.7723 |
| HZ064,LW-4;HZ011,CM-1 | 0.0140927 | 0.000308663 | 45.6574 |

| | | | |
|---|---|---|---|
| HZ058,HZ015;HZ017,CM-1 | 0.00343353 | 0.000266932 | 12.8629 |
| HZ058,HZ017;HZ015,CM-1 | 0.00100775 | 0.000256865 | 3.92327 |
| HZ058,HZ016;XWY-3,CM-1 | -0.00066514 | 0.000294969 | -2.25496 |
| HZ058,XWY-3;HZ016,CM-1 | -0.00132881 | 0.000273833 | -4.85263 |
| ZW,XWY-5;s303-231,CM-1 | -0.00186258 | 0.00022512 | -8.27372 |
| ZW,s303-231;XWY-5,CM-1 | 0.00809621 | 0.000310112 | 26.1074 |
| NC,HZ117;JGL,CM-1 | 0.000538501 | 0.000252699 | 2.131 |
| NC,JGL;HZ117,CM-1 | -0.00339953 | 0.000219203 | -15.5086 |
| HZ125,HZ109;s1-16,CM-1 | -0.0108997 | 0.00026948 | -40.4472 |
| HZ125,s1-16;HZ109,CM-1 | -0.0107043 | 0.000267182 | -40.0638 |
| XYDCS,HZ066;HZ084,CM-1 | -0.00172081 | 0.000277815 | -6.19409 |
| XYDCS,HZ084;HZ066,CM-1 | -0.0037225 | 0.000269374 | -13.8191 |
| HZ104,HZ027;HZ125,CM-1 | 0.00136152 | 0.000255009 | 5.33911 |
| HZ104,HZ125;HZ027,CM-1 | -0.0019662 | 0.000217987 | -9.01978 |
| HZ013,HZ004;HZ027,CM-1 | -0.00031141 | 0.000208322 | -1.49485 |
| HZ013,HZ027;HZ004,CM-1 | 0.0109311 | 0.000281071 | 38.891 |
| HZ006,GLJYD;HZ011,CM-1 | 0.00574288 | 0.000291335 | 19.7123 |
| HZ006,HZ011;GLJYD,CM-1 | -0.00027873 | 0.00023033 | -1.21011 |
| WLH-1,HZ020;MZYCS,CM-1 | -0.0107365 | 0.000310518 | -34.576 |
| WLH-1,MZYCS;HZ020,CM-1 | -0.0105848 | 0.00031352 | -33.7611 |
| MZYCS,DBZ-1;GE-1,CM-1 | 0.00591336 | 0.000263111 | 22.4748 |
| MZYCS,GE-1;DBZ-1,CM-1 | 0.00116658 | 0.000223233 | 5.22585 |
| LW-4,HZ014;s303-231,CM-1 | -0.00244724 | 0.000269586 | -9.07777 |
| LW-4,s303-231;HZ014,CM-1 | -0.00384749 | 0.00026392 | -14.5783 |
| NC,HZ065;MZYCS,CM-1 | -0.00408706 | 0.000278961 | -14.651 |
| NC,MZYCS;HZ065,CM-1 | -0.00738592 | 0.000241912 | -30.5315 |
| HZ071,SLLK;YH-1-1,CM-1 | -0.00240833 | 0.000243711 | -9.88192 |
| HZ071,YH-1-1;SLLK,CM-1 | 0.00295365 | 0.000292647 | 10.0929 |
| NBE-1,HZ109;NBE-5,CM-1 | 0.0111299 | 0.000280173 | 39.7249 |
| NBE-1,NBE-5;HZ109,CM-1 | -8.24E-05 | 0.000186629 | -0.441487 |
| HZ079,HZ031;LC-2,CM-1 | -0.00280515 | 0.000252004 | -11.1314 |
| HZ079,LC-2;HZ031,CM-1 | -0.00191796 | 0.000255777 | -7.49856 |
| YNZ-1,HZ090;XY-10,CM-1 | -0.00566657 | 0.000265824 | -21.317 |
| YNZ-1,XY-10;HZ090,CM-1 | -0.00583122 | 0.000267661 | -21.7859 |
| HZ034,LW-1;XWY-2,CM-1 | 0.00788418 | 0.00029967 | 26.3095 |
| HZ034,XWY-2;LW-1,CM-1 | 5.10E-05 | 0.000229738 | 0.221937 |
| HZ057,HZ061;YH-2,CM-1 | -0.00345717 | 0.000220422 | -15.6843 |
| HZ057,YH-2;HZ061,CM-1 | 0.00382756 | 0.000287146 | 13.3296 |
| HZ112,JHZS;LW-1,CM-1 | 0.000263441 | 0.000212994 | 1.23685 |
| HZ112,LW-1;JHZS,CM-1 | 0.0106448 | 0.000313535 | 33.951 |
| HZ020,FL;XY-10,CM-1 | -0.00062523 | 0.000259103 | -2.41306 |
| HZ020,XY-10;FL,CM-1 | -0.00178317 | 0.000243669 | -7.31802 |
| XSX,HZ010;YH-1-1,CM-1 | 7.36E-05 | 0.000206652 | 0.35632 |
| XSX,YH-1-1;HZ010,CM-1 | 0.0157388 | 0.000352553 | 44.6424 |
| FL,HZ090;LW-4,CM-1 | -0.00114002 | 0.000224283 | -5.08296 |
| FL,LW-4;HZ090,CM-1 | 0.00683897 | 0.000279284 | 24.4875 |
| HZ006,HZ092;XWY-1,CM-1 | 0.00974979 | 0.000308906 | 31.5624 |
| HZ006,XWY-1;HZ092,CM-1 | 0.000824386 | 0.000218593 | 3.77133 |
| HZ094,HZ071;HZ075,CM-1 | -0.0148673 | 0.000311505 | -47.7273 |
| HZ094,HZ075;HZ071,CM-1 | -0.015492 | 0.000311896 | -49.6705 |

| | | | |
|---|---|---|---|
| GE-6,GE-4;XWY-1,CM-1 | 8.78E-06 | 0.000246084 | 0.0356938 |
| GE-6,XWY-1;GE-4,CM-1 | 0.00370578 | 0.000264899 | 13.9894 |
| WL,LW-5;NBE-2,CM-1 | 0.0063038 | 0.000308708 | 20.4199 |
| WL,NBE-2;LW-5,CM-1 | -0.00065989 | 0.000228292 | -2.89057 |
| HZ109,HZ068;HZ083,CM-1 | -0.0136852 | 0.000322999 | -42.3692 |
| HZ109,HZ083;HZ068,CM-1 | -0.0136689 | 0.000323193 | -42.2932 |
| LC-1,LW-4;XY-10,CM-1 | 0.0134394 | 0.000303222 | 44.3219 |
| LC-1,XY-10;LW-4,CM-1 | -0.0005897 | 0.00019252 | -3.06306 |
| HZ118,HZ123;XH-1,CM-1 | -6.28E-06 | 0.000193358 | -0.0325006 |
| HZ118,XH-1;HZ123,CM-1 | 0.00681845 | 0.00024735 | 27.5659 |
| LC-1,HZ057;XWY-1,CM-1 | 0.00351658 | 0.000241948 | 14.5344 |
| LC-1,XWY-1;HZ057,CM-1 | 0.00453633 | 0.000238167 | 19.0469 |
| s303-231,HZ068;HZ123,CM-1 | 0.00402301 | 0.000247197 | 16.2745 |
| s303-231,HZ123;HZ068,CM-1 | 0.00366305 | 0.000242182 | 15.1252 |
| HZ045,HZ088;HZ109,CM-1 | 0.00077619 | 0.000215032 | 3.60965 |
| HZ045,HZ109;HZ088,CM-1 | 0.00723108 | 0.000265157 | 27.2709 |
| HZ006,HSKC;HZ109,CM-1 | 0.00166227 | 0.000219386 | 7.57692 |
| HZ006,HZ109;HSKC,CM-1 | 0.00913547 | 0.000288625 | 31.6517 |
| HZ117,HZ010;HZ086,CM-1 | -0.00851735 | 0.000293045 | -29.065 |
| HZ117,HZ086;HZ010,CM-1 | -0.00918669 | 0.000292052 | -31.4557 |
| HZ057,HZ045;NBE-4,CM-1 | -0.00180095 | 0.000232983 | -7.72994 |
| HZ057,NBE-4;HZ045,CM-1 | 0.00144277 | 0.000250222 | 5.76598 |
| HZ065,HZ048;HZ090,CM-1 | -0.00158102 | 0.000237709 | -6.65109 |
| HZ065,HZ090;HZ048,CM-1 | 0.000732819 | 0.000246784 | 2.96947 |
| SB,HZ069;NC,CM-1 | 0.00129173 | 0.000198166 | 6.51844 |
| SB,NC;HZ069,CM-1 | 0.00978222 | 0.000242274 | 40.3767 |
| HZ058,HZ008;HZ016,CM-1 | -0.00917323 | 0.00028958 | -31.6778 |
| HZ058,HZ016;HZ008,CM-1 | -0.00875379 | 0.000295428 | -29.6309 |
| HZ082,ST330;s1-16,CM-1 | 0.000370929 | 0.000232127 | 1.59796 |
| HZ082,s1-16;ST330,CM-1 | 0.00220043 | 0.000239447 | 9.18963 |
| HZ009,NBE-4;XSL,CM-1 | 0.00396349 | 0.000289376 | 13.6967 |
| HZ009,XSL;NBE-4,CM-1 | -0.0012356 | 0.0002384 | -5.18289 |
| HZ020,HZ080;NBE-2,CM-1 | 0.00200694 | 0.000255546 | 7.85353 |
| HZ020,NBE-2;HZ080,CM-1 | 0.00332863 | 0.000255414 | 13.0323 |
| s6-8,HZ040;SB,CM-1 | -0.0100308 | 0.00026078 | -38.4646 |
| s6-8,SB;HZ040,CM-1 | -0.00996554 | 0.000260489 | -38.2571 |
| HZ027,GE-8;HZ004,CM-1 | -0.0125166 | 0.00029631 | -42.2417 |
| HZ027,HZ004;GE-8,CM-1 | -0.0111184 | 0.000303922 | -36.5831 |
| HZ041,HZ048;YH-2,CM-1 | 0.000578741 | 0.0002192 | 2.64024 |
| HZ041,YH-2;HZ048,CM-1 | 0.0101571 | 0.000301193 | 33.7229 |
| HZ076,BCS;HZ094,CM-1 | 0.000343314 | 0.000194666 | 1.76361 |
| HZ076,HZ094;BCS,CM-1 | 0.0139743 | 0.000310363 | 45.0257 |
| HZ048,DBZ-1;HZ073,CM-1 | -0.00161963 | 0.000248781 | -6.51029 |
| HZ048,HZ073;DBZ-1,CM-1 | -0.00377222 | 0.000237456 | -15.886 |
| HZ002,HSKC;WL,CM-1 | 0.000612515 | 0.000255271 | 2.39947 |
| HZ002,WL;HSKC,CM-1 | -0.00293208 | 0.000240438 | -12.1948 |
| HZ082,HZ009;HZ010,CM-1 | 0.000550164 | 0.000274878 | 2.00149 |
| HZ082,HZ010;HZ009,CM-1 | -0.00249677 | 0.000256887 | -9.7193 |
| s301-1,HZ100;LW-1,CM-1 | 0.000487661 | 0.000237333 | 2.05476 |
| s301-1,LW-1;HZ100,CM-1 | -0.00045473 | 0.000236156 | -1.92555 |

| | | | |
|---|---|---|---|
| XWY-1,HZ070;XWY-5,CM-1 | 0.00497251 | 0.000293193 | 16.9598 |
| XWY-1,XWY-5;HZ070,CM-1 | -0.00144354 | 0.00022958 | -6.28776 |
| WL,HZ109;HZ125,CM-1 | -0.00066686 | 0.000246001 | -2.7108 |
| WL,HZ125;HZ109,CM-1 | 0.00361701 | 0.000256523 | 14.1001 |
| HZ027,HZ071;HZ119,CM-1 | -0.0102968 | 0.000249092 | -41.3375 |
| HZ027,HZ119;HZ071,CM-1 | -0.0070459 | 0.000256819 | -27.4353 |
| HZ037,HZ041;HZ074,CM-1 | -0.00051179 | 0.000195007 | -2.62445 |
| HZ037,HZ074;HZ041,CM-1 | 0.013627 | 0.000313613 | 43.4516 |
| YNZ-1,HZ118;MZYCS,CM-1 | 0.0030372 | 0.000264003 | 11.5044 |
| YNZ-1,MZYCS;HZ118,CM-1 | 0.00178824 | 0.000247116 | 7.23647 |
| HZ072,GE-6;HZ118,CM-1 | 0.0103525 | 0.000270095 | 38.329 |
| HZ072,HZ118;GE-6,CM-1 | 0.0015523 | 0.000195159 | 7.95404 |
| HZ076,HZ094;LW-2,CM-1 | 0.00707837 | 0.000275728 | 25.6716 |
| HZ076,LW-2;HZ094,CM-1 | -0.0004079 | 0.000208888 | -1.9527 |
| HZ123,HZ021;LW-1,CM-1 | 0.00387253 | 0.000262032 | 14.7789 |
| HZ123,LW-1;HZ021,CM-1 | -2.92E-05 | 0.000227926 | -0.128166 |
| HZ123,HZ003;HZ036,CM-1 | -0.0137768 | 0.00032792 | -42.0126 |
| HZ123,HZ036;HZ003,CM-1 | -0.0139085 | 0.000315682 | -44.0584 |
| ASM,HZ048;HZ058,CM-1 | -0.00760291 | 0.00030685 | -24.7773 |
| ASM,HZ058;HZ048,CM-1 | -0.00816528 | 0.000303265 | -26.9246 |
| XSL,HZ017;XWY-4,CM-1 | 0.00679392 | 0.000260483 | 26.082 |
| XSL,XWY-4;HZ017,CM-1 | 9.10E-05 | 0.000202078 | 0.450396 |
| s1-16,HZ039;HZ050,CM-1 | 0.000100716 | 0.000252464 | 0.39893 |
| s1-16,HZ050;HZ039,CM-1 | -0.00160891 | 0.000233581 | -6.88802 |
| HZ094,HZ043;LW-2,CM-1 | -0.00706758 | 0.000345974 | -20.4281 |
| HZ094,LW-2;HZ043,CM-1 | -0.0078655 | 0.00033004 | -23.8319 |
| HZ074,DBZ-1;HZ125,CM-1 | 0.000319731 | 0.000212809 | 1.50243 |
| HZ074,HZ125;DBZ-1,CM-1 | 0.00769726 | 0.000275932 | 27.8955 |
| GE-1,HZ020;HZ037,CM-1 | 0.000249462 | 0.000256012 | 0.974415 |
| GE-1,HZ037;HZ020,CM-1 | 0.000384566 | 0.000243088 | 1.582 |
| GE-1,HZ088;YH-1-1,CM-1 | 0.000497939 | 0.000214803 | 2.31812 |
| GE-1,YH-1-1;HZ088,CM-1 | 0.0121362 | 0.000319065 | 38.0368 |
| XSX,HZ057;HZ058,CM-1 | 0.00414913 | 0.00028388 | 14.6158 |
| XSX,HZ058;HZ057,CM-1 | -0.00054692 | 0.000263401 | -2.07636 |
| HZ085,HZ063;QXDM-1,CM-1 | 0.00121134 | 0.000241304 | 5.01996 |
| HZ085,QXDM-1;HZ063,CM-1 | 0.00380937 | 0.000268568 | 14.184 |
| HZ009,HZ020;LC-2,CM-1 | -0.00356126 | 0.000263594 | -13.5104 |
| HZ009,LC-2;HZ020,CM-1 | -0.00264214 | 0.000278548 | -9.48538 |
| HZ002,GZ;YNZ-1,CM-1 | -0.00015143 | 0.000217048 | -0.697689 |
| HZ002,YNZ-1;GZ,CM-1 | 0.00489942 | 0.000249785 | 19.6146 |
| YH-2,HZ010;YH-1-1,CM-1 | 0.0157933 | 0.000308651 | 51.1687 |
| YH-2,YH-1-1;HZ010,CM-1 | -0.00025982 | 0.000167563 | -1.55059 |
| LW-5,HZ074;YNLDP2,CM-1 | -0.00040429 | 0.000255717 | -1.58101 |
| LW-5,YNLDP2;HZ074,CM-1 | -0.00061972 | 0.000253753 | -2.44223 |
| HZ013,XWY-3;YNLDP2,CM-1 | -0.00122353 | 0.000216034 | -5.66361 |
| HZ013,YNLDP2;XWY-3,CM-1 | 0.00715598 | 0.000277856 | 25.7543 |
| HZ068,HZ063;HZ085,CM-1 | -0.00188924 | 0.000262113 | -7.20774 |
| HZ068,HZ085;HZ063,CM-1 | -0.00297279 | 0.000250813 | -11.8526 |
| SB,HZ063;s6-8,CM-1 | 0.00171712 | 0.000195963 | 8.76247 |
| SB,s6-8;HZ063,CM-1 | 0.00672281 | 0.000234476 | 28.6717 |

| | | | |
|---|---|---|---|
| JX-4,HZ065;HZ070,CM-1 | 0.0022978 | 0.000277173 | 8.29014 |
| JX-4,HZ070;HZ065,CM-1 | -0.00162714 | 0.000230217 | -7.06783 |
| HZ088,HZ118;XSL,CM-1 | 0.0112287 | 0.000320388 | 35.0472 |
| HZ088,XSL;HZ118,CM-1 | 0.000325555 | 0.000211254 | 1.54106 |
| HZ075,GE-6;HZ013,CM-1 | -0.00696709 | 0.000260108 | -26.7854 |
| HZ075,HZ013;GE-6,CM-1 | -0.00580346 | 0.000268482 | -21.6158 |
| HZ036,XWY-1;XYDCS,CM-1 | 0.00142279 | 0.00020646 | 6.89135 |
| HZ036,XYDCS;XWY-1,CM-1 | 0.0116635 | 0.000297699 | 39.1788 |
| HZ064,HZ039;MZYCS,CM-1 | 0.0124397 | 0.000327064 | 38.0343 |
| HZ064,MZYCS;HZ039,CM-1 | 0.000514496 | 0.000216037 | 2.38152 |
| HZ054,HZ060;SLLK,CM-1 | 0.000650196 | 0.000213988 | 3.03847 |
| HZ054,SLLK;HZ060,CM-1 | 0.00864688 | 0.000291777 | 29.6352 |
| HZ061,HZ104;XWY-2,CM-1 | 0.00421987 | 0.000261361 | 16.1457 |
| HZ061,XWY-2;HZ104,CM-1 | 0.000771615 | 0.000235574 | 3.27548 |
| LW-3,HZ104;YNLDP2,CM-1 | 0.000722923 | 0.000238874 | 3.02638 |
| LW-3,YNLDP2;HZ104,CM-1 | 0.000673031 | 0.000233361 | 2.88408 |
| HZ031,HZ068;XH-1,CM-1 | -0.0003599 | 0.000219339 | -1.64083 |
| HZ031,XH-1;HZ068,CM-1 | 0.00646463 | 0.000265164 | 24.3798 |
| HZ008,HZ036;LBDCS,CM-1 | 0.000142909 | 0.000152656 | 0.936151 |
| HZ008,LBDCS;HZ036,CM-1 | 0.0212797 | 0.00031683 | 67.1645 |
| HZ075,HZ065;HZ114,CM-1 | 0.00166283 | 0.000208985 | 7.95673 |
| HZ075,HZ114;HZ065,CM-1 | 0.00907618 | 0.000255133 | 35.5743 |
| LW-5,JX-4;NBE-3,CM-1 | -0.00509431 | 0.000274913 | -18.5306 |
| LW-5,NBE-3;JX-4,CM-1 | -0.00540999 | 0.000263341 | -20.5437 |
| GZ,HZ064;YH-2,CM-1 | 0.000970412 | 0.00021195 | 4.57849 |
| GZ,YH-2;HZ064,CM-1 | 0.0106359 | 0.000309767 | 34.3353 |
| HZ057,HZ034;HZ070,CM-1 | -0.00315251 | 0.000248701 | -12.6759 |
| HZ057,HZ070;HZ034,CM-1 | 0.00107308 | 0.000275466 | 3.89549 |
| HZ050,HZ021;LW-1,CM-1 | 0.000148927 | 0.00016771 | 0.888006 |
| HZ050,LW-1;HZ021,CM-1 | 0.0202191 | 0.000317196 | 63.7433 |
| GLJYD,HZ054;YH-1-1,CM-1 | 0.00125332 | 0.000214355 | 5.84693 |
| GLJYD,YH-1-1;HZ054,CM-1 | 0.0108068 | 0.000297232 | 36.3582 |
| HZ109,GLJYD;HZ082,CM-1 | -0.00761586 | 0.000289274 | -26.3275 |
| HZ109,HZ082;GLJYD,CM-1 | -0.00816202 | 0.000292287 | -27.9247 |
| HZ006,HZ061;XWY-3,CM-1 | 0.00829475 | 0.000293558 | 28.2559 |
| HZ006,XWY-3;HZ061,CM-1 | 0.000363399 | 0.000221993 | 1.63699 |
| HZ076,LC-1;XSL,CM-1 | -0.00837307 | 0.000246598 | -33.9544 |
| HZ076,XSL;LC-1,CM-1 | -0.00762045 | 0.000250873 | -30.3758 |
| GLJYD,HZ021;JX-4,CM-1 | -0.00144931 | 0.00025228 | -5.74484 |
| GLJYD,JX-4;HZ021,CM-1 | -0.00038795 | 0.00026596 | -1.45868 |
| HZ125,HZ109;NBE-1,CM-1 | -0.0035953 | 0.000245525 | -14.6433 |
| HZ125,NBE-1;HZ109,CM-1 | -0.00420212 | 0.000246013 | -17.0809 |
| GE-8,HSKC;HZ006,CM-1 | 0.00535775 | 0.000275862 | 19.4218 |
| GE-8,HZ006;HSKC,CM-1 | -0.00021312 | 0.000244449 | -0.871833 |
| HZ068,HZ015;HZ070,CM-1 | 0.00416408 | 0.000277667 | 14.9967 |
| HZ068,HZ070;HZ015,CM-1 | 0.00051269 | 0.000236691 | 2.16607 |
| JGL,GLJYD;ZBD,CM-1 | 0.00390973 | 0.000224384 | 17.4243 |
| JGL,ZBD;GLJYD,CM-1 | 0.000618909 | 0.000201107 | 3.07751 |
| s301-1,HZ004;XYDCS,CM-1 | -0.00339564 | 0.000249602 | -13.6042 |
| s301-1,XYDCS;HZ004,CM-1 | 0.00309472 | 0.000287624 | 10.7596 |

| | | | |
|---|---|---|---|
| HZ084,HZ002;HZ037,CM-1 | -0.00369354 | 0.0002713 | -13.6142 |
| HZ084,HZ037;HZ002,CM-1 | -0.00551521 | 0.000262203 | -21.0341 |
| HZ123,HZ058;HZ064,CM-1 | -0.0111205 | 0.000331162 | -33.5802 |
| HZ123,HZ064;HZ058,CM-1 | -0.0107848 | 0.000328697 | -32.8108 |
| HZ059,HZ075;HZ094,CM-1 | -0.00037335 | 0.00019667 | -1.89837 |
| HZ059,HZ094;HZ075,CM-1 | 0.0138988 | 0.000314575 | 44.1827 |
| YH-2,HZ050;ST330,CM-1 | 0.00579403 | 0.000303696 | 19.0784 |
| YH-2,ST330;HZ050,CM-1 | -0.00335832 | 0.000213673 | -15.7171 |
| HZ037,HZ064;HZ090,CM-1 | -5.99E-05 | 0.000184054 | -0.325198 |
| HZ037,HZ090;HZ064,CM-1 | 0.0105086 | 0.000267342 | 39.3076 |
| HZ081,HZ002;LW-4,CM-1 | 2.75E-05 | 0.000212491 | 0.1294 |
| HZ081,LW-4;HZ002,CM-1 | 0.00814961 | 0.000281621 | 28.9382 |
| HZ095,JGL;NBE-5,CM-1 | -0.0179499 | 0.000348804 | -51.4613 |
| HZ095,NBE-5;JGL,CM-1 | -0.019139 | 0.000330319 | -57.941 |
| GE-4,HZ020;XYDCS,CM-1 | -0.00054682 | 0.00022836 | -2.39455 |
| GE-4,XYDCS;HZ020,CM-1 | 0.00823765 | 0.000290585 | 28.3485 |
| HZ070,HZ110;YH-2,CM-1 | 0.000753517 | 0.000257851 | 2.9223 |
| HZ070,YH-2;HZ110,CM-1 | -0.00175356 | 0.000250512 | -6.9999 |
| ASM,HZ117;QXDM-1,CM-1 | -0.00089433 | 0.000255273 | -3.50341 |
| ASM,QXDM-1;HZ117,CM-1 | 0.000509581 | 0.000267268 | 1.90663 |
| HZ069,NBE-2;s317,CM-1 | -0.00690956 | 0.000251333 | -27.4916 |
| HZ069,s317;NBE-2,CM-1 | 0.000614591 | 0.000335811 | 1.83017 |
| NBE-2,HZ048;NBE-3,CM-1 | 0.0112451 | 0.000285113 | 39.4408 |
| NBE-2,NBE-3;HZ048,CM-1 | 0.000282316 | 0.000188324 | 1.4991 |
| BCS,HZ015;JGL,CM-1 | 0.00892321 | 0.00025085 | 35.572 |
| BCS,JGL;HZ015,CM-1 | 0.00101508 | 0.000192526 | 5.27243 |
| LBDCS,HZ050;HZ061,CM-1 | -0.00663377 | 0.000268929 | -24.6674 |
| LBDCS,HZ061;HZ050,CM-1 | -0.00635249 | 0.000256431 | -24.7727 |
| HZ034,HZ110;XWY-2,CM-1 | 0.0110074 | 0.000303119 | 36.3137 |
| HZ034,XWY-2;HZ110,CM-1 | 0.000410611 | 0.000204756 | 2.00537 |
| HZ122,HZ075;HZ076,CM-1 | -0.00221862 | 0.000221517 | -10.0156 |
| HZ122,HZ076;HZ075,CM-1 | 0.00100205 | 0.000245752 | 4.07747 |
| QXDM-1,HZ085;ZW,CM-1 | 0.0014476 | 0.000292836 | 4.94337 |
| QXDM-1,ZW;HZ085,CM-1 | -0.0016002 | 0.0002598 | -6.15936 |
| FL,HZ041;LW-5,CM-1 | 3.85E-06 | 0.000203438 | 0.0189094 |
| FL,LW-5;HZ041,CM-1 | 0.0127578 | 0.000321432 | 39.6906 |
| ST312,HZ010;NC,CM-1 | -0.00563537 | 0.000246716 | -22.8415 |
| ST312,NC;HZ010,CM-1 | -0.00022697 | 0.000276414 | -0.821129 |
| HZ064,HZ040;NBE-2,CM-1 | 0.000790043 | 0.000216863 | 3.64305 |
| HZ064,NBE-2;HZ040,CM-1 | 0.00638807 | 0.00027569 | 23.1712 |
| HZ059,HZ010;HZ072,CM-1 | 0.000233808 | 0.000224648 | 1.04078 |
| HZ059,HZ072;HZ010,CM-1 | 0.00974978 | 0.000313987 | 31.0515 |
| HZ112,HZ070;HZ125,CM-1 | 0.000604843 | 0.000202869 | 2.98145 |
| HZ112,HZ125;HZ070,CM-1 | 0.0132659 | 0.000307426 | 43.1515 |
| HZ009,GE-1;s317,CM-1 | -0.00057806 | 0.000223369 | -2.58789 |
| HZ009,s317;GE-1,CM-1 | 0.00859387 | 0.000301758 | 28.4793 |
| HZ063,DHL;HZ014,CM-1 | -0.00223971 | 0.000251524 | -8.90456 |
| HZ063,HZ014;DHL,CM-1 | -0.00402781 | 0.000244272 | -16.489 |
| GE-6,HZ001;HZ090,CM-1 | 0.0078976 | 0.000250492 | 31.5283 |
| GE-6,HZ090;HZ001,CM-1 | 0.00305105 | 0.000225759 | 13.5147 |

| | | | |
|---|---|---|---|
| HZ003,HZ010;HZ014,CM-1 | -0.00402607 | 0.000240404 | -16.7471 |
| HZ003,HZ014;HZ010,CM-1 | -0.00250316 | 0.000260383 | -9.61336 |
| ST330,JHZS;LC-1,CM-1 | -0.00899359 | 0.000269645 | -33.3534 |
| ST330,LC-1;JHZS,CM-1 | -0.00767989 | 0.000276341 | -27.7913 |
| HZ071,FL;HZ124,CM-1 | -0.00016841 | 0.000202053 | -0.833511 |
| HZ071,HZ124;FL,CM-1 | 0.0150033 | 0.000311607 | 48.148 |
| NBE-2,HZ079;HZ117,CM-1 | -0.00340297 | 0.000244802 | -13.9009 |
| NBE-2,HZ117;HZ079,CM-1 | -0.00120568 | 0.000272023 | -4.43226 |
| YH-1-1,HZ031;WLH-1,CM-1 | 0.0134768 | 0.00028203 | 47.785 |
| YH-1-1,WLH-1;HZ031,CM-1 | -0.00121838 | 0.000173744 | -7.01253 |
| DBZ-1,NBE-2;SLLK,CM-1 | -2.50E-05 | 0.000251159 | -0.099642 |
| DBZ-1,SLLK;NBE-2,CM-1 | 0.00109867 | 0.000258269 | 4.25396 |
| s1-16,HZ092;ZW,CM-1 | 0.000414644 | 0.000262212 | 1.58133 |
| s1-16,ZW;HZ092,CM-1 | -0.00123751 | 0.000253725 | -4.87735 |
| HZ079,HZ011;LBDCS,CM-1 | -0.00129944 | 0.000217973 | -5.96146 |
| HZ079,LBDCS;HZ011,CM-1 | 0.00916135 | 0.000277883 | 32.9683 |
| ZW,HZ109;YNLDP1,CM-1 | 0.00801018 | 0.000305048 | 26.2587 |
| ZW,YNLDP1;HZ109,CM-1 | -0.00053706 | 0.000212521 | -2.52711 |
| YNLDP2,HZ039;HZ057,CM-1 | 0.0013428 | 0.000256283 | 5.23951 |
| YNLDP2,HZ057;HZ039,CM-1 | -0.00070767 | 0.00024189 | -2.92558 |
| YNLDP2,HZ001;HZ018,CM-1 | 0.00284582 | 0.000264887 | 10.7435 |
| YNLDP2,HZ018;HZ001,CM-1 | -0.00363423 | 0.00021735 | -16.7207 |
| YH-2,HZ014;HZ061,CM-1 | -0.0054854 | 0.000287007 | -19.1124 |
| YH-2,HZ061;HZ014,CM-1 | -0.007461 | 0.00026419 | -28.241 |
| HZ043,HZ010;HZ118,CM-1 | -1.52E-05 | 0.00021725 | -0.0697524 |
| HZ043,HZ118;HZ010,CM-1 | 0.0153613 | 0.000363015 | 42.3159 |
| FL,XWY-2;s301-1,CM-1 | -0.00060719 | 0.000208768 | -2.90843 |
| FL,s301-1;XWY-2,CM-1 | 0.00697467 | 0.000275718 | 25.2964 |
| HZ037,XWY-4;YNLDP1,CM-1 | 0.000210261 | 0.000240064 | 0.875855 |
| HZ037,YNLDP1;XWY-4,CM-1 | 0.00404413 | 0.00026402 | 15.3175 |
| HZ017,HZ061;HZ085,CM-1 | 0.00300725 | 0.000269551 | 11.1565 |
| HZ017,HZ085;HZ061,CM-1 | -0.00168497 | 0.000229861 | -7.33038 |
| LW-2,FL;HZ118,CM-1 | 0.00200322 | 0.000255924 | 7.8274 |
| LW-2,HZ118;FL,CM-1 | -0.00045766 | 0.000228833 | -1.99996 |
| YNZ-1,HZ064;HZ090,CM-1 | -0.00524034 | 0.000259048 | -20.2292 |
| YNZ-1,HZ090;HZ064,CM-1 | -0.00517714 | 0.000261355 | -19.8088 |
| GE-8,HZ021;WLH-1,CM-1 | -8.48E-05 | 0.000213785 | -0.396882 |
| GE-8,WLH-1;HZ021,CM-1 | 0.0103396 | 0.000299944 | 34.4718 |
| HZ011,HZ070;ZW,CM-1 | 0.00324271 | 0.00026881 | 12.0632 |
| HZ011,ZW;HZ070,CM-1 | 0.000212684 | 0.000241032 | 0.882388 |
| JHZS,HZ050;HZ066,CM-1 | 0.00144471 | 0.000247831 | 5.82943 |
| JHZS,HZ066;HZ050,CM-1 | 0.000762313 | 0.000241196 | 3.16056 |
| HZ003,HZ001;XSL,CM-1 | 0.0137449 | 0.000295893 | 46.4522 |
| HZ003,XSL;HZ001,CM-1 | 0.000533108 | 0.000192615 | 2.76775 |
| SLLK,HZ028;JHZS,CM-1 | -0.00857845 | 0.000298865 | -28.7035 |
| SLLK,JHZS;HZ028,CM-1 | -0.00955153 | 0.000284139 | -33.6157 |
| HZ011,HZ078;HZ122,CM-1 | -0.00451729 | 0.000234871 | -19.2331 |
| HZ011,HZ122;HZ078,CM-1 | 0.00469331 | 0.00031661 | 14.8237 |
| HZ085,GE-4;HZ017,CM-1 | -8.65E-05 | 0.000270836 | -0.319466 |
| HZ085,HZ017;GE-4,CM-1 | -0.00114337 | 0.000258754 | -4.41875 |

| | | | |
|---|---|---|---|
| HZ040,DHL;JHZS,CM-1 | -0.0027686 | 0.000242298 | -11.4264 |
| HZ040,JHZS;DHL,CM-1 | -0.00350257 | 0.000232187 | -15.0851 |
| ST312,HZ009;HZ054,CM-1 | -0.00721729 | 0.000289425 | -24.9366 |
| ST312,HZ054;HZ009,CM-1 | -0.00753589 | 0.000285413 | -26.4034 |
| HZ037,HZ011;HZ017,CM-1 | -0.00230181 | 0.000262425 | -8.77129 |
| HZ037,HZ017;HZ011,CM-1 | -0.00199441 | 0.000255551 | -7.80435 |
| XH-1,HZ124;XYDCS,CM-1 | 0.00350443 | 0.000228244 | 15.3539 |
| XH-1,XYDCS;HZ124,CM-1 | 0.00360635 | 0.000242445 | 14.8749 |
| LW-1,HZ109;HZ114,CM-1 | 0.00020178 | 0.000204298 | 0.987673 |
| LW-1,HZ114;HZ109,CM-1 | 0.0100059 | 0.000287714 | 34.7773 |
| HZ119,GE-1;HZ057,CM-1 | -0.00703214 | 0.000257725 | -27.2854 |
| HZ119,HZ057;GE-1,CM-1 | -0.00441261 | 0.000273942 | -16.1079 |
| HZ074,HZ082;XSX,CM-1 | -0.0087405 | 0.000305993 | -28.5644 |
| HZ074,XSX;HZ082,CM-1 | -0.00791591 | 0.000303417 | -26.0892 |
| HZ027,HZ056;NBE-3,CM-1 | -0.00568961 | 0.000238177 | -23.8882 |
| HZ027,NBE-3;HZ056,CM-1 | -0.00201046 | 0.000279133 | -7.20251 |
| HZ017,HZ081;HZ090,CM-1 | 0.00104289 | 0.000224047 | 4.65479 |
| HZ017,HZ090;HZ081,CM-1 | 0.00566594 | 0.000258507 | 21.918 |
| HZ003,HZ001;ZW,CM-1 | 0.0145092 | 0.000312207 | 46.4729 |
| HZ003,ZW;HZ001,CM-1 | 0.000131243 | 0.000196142 | 0.669119 |
| HZ006,ST312;ZW,CM-1 | 0.0129064 | 0.000333128 | 38.7431 |
| HZ006,ZW;ST312,CM-1 | -3.82E-06 | 0.000211965 | -0.018022 |
| JGL,HZ112;ST312,CM-1 | 0.000614937 | 0.000198986 | 3.09035 |
| JGL,ST312;HZ112,CM-1 | 0.00829667 | 0.000277797 | 29.8659 |
| GE-1,HZ104;WLH-1,CM-1 | -0.00481533 | 0.000273608 | -17.5994 |
| GE-1,WLH-1;HZ104,CM-1 | -0.00706916 | 0.000246721 | -28.6524 |
| LW-5,HZ110;XH-1,CM-1 | 0.00328663 | 0.00025213 | 13.0355 |
| LW-5,XH-1;HZ110,CM-1 | -0.00094804 | 0.000217181 | -4.36519 |
| NBE-4,XWY-3;s1-16,CM-1 | 1.91E-05 | 0.000227875 | 0.083721 |
| NBE-4,s1-16;XWY-3,CM-1 | 0.00509602 | 0.000272239 | 18.719 |
| NC,GE-4;HZ070,CM-1 | -0.008235 | 0.000260957 | -31.5569 |
| NC,HZ070;GE-4,CM-1 | -0.00788648 | 0.000273482 | -28.8373 |
| HZ017,HZ020;HZ080,CM-1 | -0.00041111 | 0.000227478 | -1.80727 |
| HZ017,HZ080;HZ020,CM-1 | 0.00470319 | 0.000251199 | 18.723 |
| BCS,LW-1;WL,CM-1 | 0.0206661 | 0.000326389 | 63.3174 |
| BCS,WL;LW-1,CM-1 | 8.24E-05 | 0.000166174 | 0.496154 |
| LW-5,GE-1;XWY-2,CM-1 | -0.00969159 | 0.000306359 | -31.6348 |
| LW-5,XWY-2;GE-1,CM-1 | -0.00989757 | 0.000310841 | -31.8413 |
| HZ077,YH-2;YNZ-1,CM-1 | 0.000247071 | 0.000289943 | 0.852137 |
| HZ077,YNZ-1;YH-2,CM-1 | -0.00253633 | 0.000253937 | -9.98804 |
| XH-1,GE-1;XWY-4,CM-1 | -0.0064804 | 0.000260818 | -24.8464 |
| XH-1,XWY-4;GE-1,CM-1 | -0.00699376 | 0.000269153 | -25.9844 |
| HZ010,HZ014;HZ124,CM-1 | -0.0002065 | 0.000165839 | -1.2452 |
| HZ010,HZ124;HZ014,CM-1 | 0.0192235 | 0.000315317 | 60.9656 |
| GE-1,GE-6;HZ114,CM-1 | -0.00024913 | 0.000183385 | -1.35848 |
| GE-1,HZ114;GE-6,CM-1 | 0.0180088 | 0.000332314 | 54.1921 |
| HZ081,HZ050;HZ082,CM-1 | 0.00369924 | 0.000260728 | 14.1881 |
| HZ081,HZ082;HZ050,CM-1 | 0.00128977 | 0.000231583 | 5.56938 |
| XWY-3,HZ063;ZW,CM-1 | 0.00826014 | 0.000281982 | 29.2931 |
| XWY-3,ZW;HZ063,CM-1 | -0.00044298 | 0.000210013 | -2.10928 |

| | | | |
|---|---|---|---|
| HZ080,HZ017;HZ040,CM-1 | -0.00855497 | 0.000261798 | -32.6777 |
| HZ080,HZ040;HZ017,CM-1 | -0.00812418 | 0.000262782 | -30.9161 |
| HZ075,HZ073;LC-1,CM-1 | -0.00739521 | 0.0002506 | -29.51 |
| HZ075,LC-1;HZ073,CM-1 | -0.0071246 | 0.000249379 | -28.5694 |
| HZ095,ST330;XWY-5,CM-1 | -0.016404 | 0.000328104 | -49.9964 |
| HZ095,XWY-5;ST330,CM-1 | -0.0159359 | 0.000332783 | -47.8869 |
| XYDCS,NBE-5;XWY-2,CM-1 | -0.00868856 | 0.000281838 | -30.8282 |
| XYDCS,XWY-2;NBE-5,CM-1 | -0.00948164 | 0.000277223 | -34.2022 |
| XWY-3,HZ013;HZ071,CM-1 | 0.00240251 | 0.000263878 | 9.10465 |
| XWY-3,HZ071;HZ013,CM-1 | -0.00037791 | 0.000231438 | -1.63288 |
| HZ065,XY-10;s1-16,CM-1 | -0.00110454 | 0.000218716 | -5.0501 |
| HZ065,s1-16;XY-10,CM-1 | 0.00647486 | 0.000282385 | 22.9292 |
| HZ100,GZ;HZ016,CM-1 | -0.0109936 | 0.000320521 | -34.299 |
| HZ100,HZ016;GZ,CM-1 | -0.0104755 | 0.000326891 | -32.0459 |
| NBE-3,HZ117;s1-16,CM-1 | -0.00112396 | 0.000217053 | -5.17826 |
| NBE-3,s1-16;HZ117,CM-1 | 0.00137675 | 0.000222075 | 6.19948 |
| HZ122,HZ112;YNLDP2,CM-1 | -0.00038259 | 0.000275046 | -1.39101 |
| HZ122,YNLDP2;HZ112,CM-1 | -0.00198649 | 0.000250025 | -7.94518 |
| HZ001,GLJYD;HZ075,CM-1 | -0.00631475 | 0.000238968 | -26.4251 |
| HZ001,HZ075;GLJYD,CM-1 | -0.00422839 | 0.000271661 | -15.5649 |
| LC-2,HZ045;ST312,CM-1 | -0.00146313 | 0.000222694 | -6.57013 |
| LC-2,ST312;HZ045,CM-1 | 0.0036313 | 0.000261231 | 13.9007 |
| HZ081,HZ072;HZ122,CM-1 | -0.00870147 | 0.000281608 | -30.8992 |
| HZ081,HZ122;HZ072,CM-1 | -0.00817368 | 0.000283803 | -28.8006 |
| DBZ-1,HZ092;ST312,CM-1 | 0.00269482 | 0.000228257 | 11.8061 |
| DBZ-1,ST312;HZ092,CM-1 | 0.00194767 | 0.00023663 | 8.23087 |
| HZ066,BCS;YH-2,CM-1 | 0.000188984 | 0.000205438 | 0.919909 |
| HZ066,YH-2;BCS,CM-1 | 0.0126783 | 0.000324412 | 39.0808 |
| HZ092,HZ003;HZ054,CM-1 | -0.0142212 | 0.000315747 | -45.04 |
| HZ092,HZ054;HZ003,CM-1 | -0.0139623 | 0.000317973 | -43.9103 |
| HZ017,HZ088;XYDCS,CM-1 | -0.00291149 | 0.00022932 | -12.6962 |
| HZ017,XYDCS;HZ088,CM-1 | 0.00731236 | 0.00030639 | 23.8662 |
| HZ076,GE-4;HZ092,CM-1 | -0.00215576 | 0.000224642 | -9.59645 |
| HZ076,HZ092;GE-4,CM-1 | 0.00235414 | 0.000259981 | 9.05502 |
| YNLDP2,HZ112;s6-8,CM-1 | -0.00149224 | 0.000232304 | -6.42363 |
| YNLDP2,s6-8;HZ112,CM-1 | -0.00020681 | 0.000249009 | -0.830531 |
| HZ066,HZ039;WL,CM-1 | 0.0114772 | 0.000313268 | 36.637 |
| HZ066,WL;HZ039,CM-1 | 0.000589119 | 0.000210436 | 2.79951 |
| HZ077,HZ015;NBE-1,CM-1 | 0.00258292 | 0.000238566 | 10.8269 |
| HZ077,NBE-1;HZ015,CM-1 | 0.00533918 | 0.000259606 | 20.5665 |
| HZ058,HZ084;LC-2,CM-1 | 0.00474539 | 0.000291161 | 16.2982 |
| HZ058,LC-2;HZ084,CM-1 | -0.00116389 | 0.000240489 | -4.8397 |
| HZ061,HZ065;MZYCS,CM-1 | -0.00143971 | 0.000267381 | -5.3845 |
| HZ061,MZYCS;HZ065,CM-1 | -0.00344987 | 0.000253903 | -13.5873 |
| HZ123,HZ045;HZ080,CM-1 | -0.00490907 | 0.000224111 | -21.9047 |
| HZ123,HZ080;HZ045,CM-1 | 0.00020662 | 0.000257914 | 0.801123 |
| SLLK,HZ001;HZ110,CM-1 | -0.00015746 | 0.000229132 | -0.687191 |
| SLLK,HZ110;HZ001,CM-1 | 0.00157703 | 0.000234733 | 6.7184 |
| HZ081,HZ117;HZ125,CM-1 | -0.00450798 | 0.000233375 | -19.3164 |
| HZ081,HZ125;HZ117,CM-1 | 0.00289731 | 0.000285999 | 10.1305 |

| | | | |
|---|---|---|---|
| HZ041,HZ060;HZ117,CM-1 | 0.00050188 | 0.00022131 | 2.26777 |
| HZ041,HZ117;HZ060,CM-1 | 0.00956225 | 0.000304042 | 31.4504 |
| NBE-3,HZ086;HZ110,CM-1 | -0.00012576 | 0.000197801 | -0.635784 |
| NBE-3,HZ110;HZ086,CM-1 | 0.00860274 | 0.000263653 | 32.629 |
| HZ070,NBE-5;s6-8,CM-1 | -0.00221196 | 0.000237649 | -9.30768 |
| HZ070,s6-8;NBE-5,CM-1 | 0.000812279 | 0.000281822 | 2.88224 |
| HZ058,HZ036;HZ037,CM-1 | -0.00364494 | 0.000280639 | -12.988 |
| HZ058,HZ037;HZ036,CM-1 | -0.00342953 | 0.000276897 | -12.3855 |
| SB,HZ010;XWY-5,CM-1 | 0.00165638 | 0.000207023 | 8.00097 |
| SB,XWY-5;HZ010,CM-1 | 0.00269695 | 0.000231712 | 11.6392 |
| HSKC,HZ015;LW-5,CM-1 | -0.00228599 | 0.000215221 | -10.6216 |
| HSKC,LW-5;HZ015,CM-1 | 0.00602806 | 0.00029473 | 20.4528 |
| HZ109,DBZ-1;HZ011,CM-1 | -0.0093613 | 0.000284617 | -32.8909 |
| HZ109,HZ011;DBZ-1,CM-1 | -0.0101323 | 0.000291052 | -34.8125 |
| LBDCS,ASM;HZ119,CM-1 | -0.00607709 | 0.000244763 | -24.8285 |
| LBDCS,HZ119;ASM,CM-1 | -0.0066884 | 0.000245408 | -27.2543 |
| HZ122,FL;LW-4,CM-1 | 0.00274261 | 0.000248282 | 11.0463 |
| HZ122,LW-4;FL,CM-1 | 0.000749376 | 0.000238911 | 3.13663 |
| XWY-3,FL;JGL,CM-1 | 0.00233445 | 0.000224409 | 10.4027 |
| XWY-3,JGL;FL,CM-1 | 0.000893565 | 0.000209206 | 4.27122 |
| YNZ-1,JGL;XYDCS,CM-1 | -0.00433917 | 0.000228976 | -18.9504 |
| YNZ-1,XYDCS;JGL,CM-1 | 0.0013458 | 0.000271944 | 4.9488 |
| HZ090,JGL;s301-1,CM-1 | -0.0004059 | 0.000208947 | -1.94261 |
| HZ090,s301-1;JGL,CM-1 | 0.00420599 | 0.000250713 | 16.7761 |
| HZ090,HZ118;XWY-4,CM-1 | 0.00814139 | 0.000253517 | 32.1138 |
| HZ090,XWY-4;HZ118,CM-1 | 0.00686815 | 0.000246598 | 27.8516 |
| GE-8,HZ110;XWY-4,CM-1 | 0.0179119 | 0.00032168 | 55.6823 |
| GE-8,XWY-4;HZ110,CM-1 | 0.000385847 | 0.000178388 | 2.16296 |
| HZ010,HZ028;XWY-2,CM-1 | 0.000898881 | 0.000228939 | 3.92629 |
| HZ010,XWY-2;HZ028,CM-1 | 0.00439428 | 0.000264808 | 16.5942 |
| HZ016,HZ056;HZ078,CM-1 | 0.00188848 | 0.00027268 | 6.92562 |
| HZ016,HZ078;HZ056,CM-1 | 0.000647563 | 0.000264228 | 2.45077 |
| HZ114,ASM;HZ010,CM-1 | -0.00712329 | 0.000254983 | -27.9363 |
| HZ114,HZ010;ASM,CM-1 | -0.00305897 | 0.000286432 | -10.6796 |
| ZBD,NBE-4;XH-1,CM-1 | -0.00116935 | 0.00023911 | -4.89042 |
| ZBD,XH-1;NBE-4,CM-1 | -0.00224916 | 0.000229329 | -9.80759 |
| HZ057,HZ001;ZW,CM-1 | 0.00635343 | 0.000261721 | 24.2756 |
| HZ057,ZW;HZ001,CM-1 | 0.00287071 | 0.000228841 | 12.5445 |
| LBDCS,HZ017;HZ027,CM-1 | 0.00211803 | 0.000252106 | 8.40133 |
| LBDCS,HZ027;HZ017,CM-1 | -0.0005141 | 0.000227946 | -2.25537 |
| HZ076,HZ071;HZ080,CM-1 | -0.00427162 | 0.000252798 | -16.8973 |
| HZ076,HZ080;HZ071,CM-1 | -0.00389481 | 0.000262859 | -14.8171 |
| HZ075,DBZ-1;JX-4,CM-1 | -0.00356642 | 0.000252813 | -14.107 |
| HZ075,JX-4;DBZ-1,CM-1 | -0.00484232 | 0.000258054 | -18.7648 |
| GE-1,LBDCS;YH-2,CM-1 | 0.00486232 | 0.000246307 | 19.7408 |
| GE-1,YH-2;LBDCS,CM-1 | 0.0057732 | 0.000252789 | 22.8381 |
| XWY-3,LC-1;ZW,CM-1 | 0.00518691 | 0.000266269 | 19.48 |
| XWY-3,ZW;LC-1,CM-1 | -0.0007119 | 0.000211577 | -3.36473 |
| QXDM-2,HZ031;HZ125,CM-1 | 0.000324998 | 0.000181876 | 1.78692 |
| QXDM-2,HZ125;HZ031,CM-1 | 0.0163865 | 0.000323682 | 50.6253 |

| | | | |
|---|---|---|---|
| HZ079,DBZ-1;NBE-5,CM-1 | 0.000549627 | 0.000248333 | 2.21326 |
| HZ079,NBE-5;DBZ-1,CM-1 | 0.000292731 | 0.000246631 | 1.18692 |
